# The sequences of 150,119 genomes in the UK biobank

**DOI:** 10.1101/2021.11.16.468246

**Authors:** Bjarni V. Halldorsson, Hannes P. Eggertsson, Kristjan H.S. Moore, Hannes Hauswedell, Ogmundur Eiriksson, Magnus O. Ulfarsson, Gunnar Palsson, Marteinn T. Hardarson, Asmundur Oddsson, Brynjar O. Jensson, Snaedis Kristmundsdottir, Brynja D. Sigurpalsdottir, Olafur A. Stefansson, Doruk Beyter, Guillaume Holley, Vinicius Tragante, Arnaldur Gylfason, Pall I. Olason, Florian Zink, Margret Asgeirsdottir, Sverrir T. Sverrisson, Brynjar Sigurdsson, Sigurjon A. Gudjonsson, Gunnar T. Sigurdsson, Gisli H. Halldorsson, Gardar Sveinbjornsson, Kristjan Norland, Unnur Styrkarsdottir, Droplaug N. Magnusdottir, Steinunn Snorradottir, Kari Kristinsson, Emilia Sobech, Helgi Jonsson, Arni J. Geirsson, Isleifur Olafsson, Palmi Jonsson, Ole Birger Pedersen, Christian Erikstrup, Søren Brunak, Sisse Rye Ostrowski, DBDS Genetic Consortium, Gudmar Thorleifsson, Frosti Jonsson, Pall Melsted, Ingileif Jonsdottir, Thorunn Rafnar, Hilma Holm, Hreinn Stefansson, Jona Saemundsdottir, Daniel F. Gudbjartsson, Olafur T. Magnusson, Gisli Masson, Unnur Thorsteinsdottir, Agnar Helgason, Hakon Jonsson, Patrick Sulem, Kari Stefansson

## Abstract

We describe the analysis of whole genome sequences (WGS) of 150,119 individuals from the UK biobank (UKB). This constitutes a set of high quality variants, including 585,040,410 SNPs, representing 7.0% of all possible human SNPs, and 58,707,036 indels. The large set of variants allows us to characterize selection based on sequence variation within a population through a Depletion Rank (DR) score for windows along the genome. DR analysis shows that coding exons represent a small fraction of regions in the genome subject to strong sequence conservation. We define three cohorts within the UKB, a large British Irish cohort (XBI) and smaller African (XAF) and South Asian (XSA) cohorts. A haplotype reference panel is provided that allows reliable imputation of most variants carried by three or more sequenced individuals. We identified 895,055 structural variants and 2,536,688 microsatellites, groups of variants typically excluded from large scale WGS studies. Using this formidable new resource, we provide several examples of trait associations for rare variants with large effects not found previously through studies based on exome sequencing and/or imputation.

## Introduction

Detailed knowledge of how diversity in the sequence of the human genome affects phenotypic diversity depends on a comprehensive and reliable characterization of both sequences and phenotypic variation. Over the past decade insights into this relationship have been obtained from whole exome (WES) and WGS of large cohorts with rich phenotypic data^1, 2^.

The UK biobank (UKB)^3^ documents phenotypic variation of 500,000 subjects across the United Kingdom, with a healthy volunteer bias^4^. The UKB WGS consortium is sequencing the whole genomes of all the participants to an average depth of at least 23.5x. Here, we report on the first data release consisting of a vast set of sequence variants, including single nucleotide polymorphisms (SNPs), short insertions/deletions (indels), microsatellites and structural variants (SVs), based on WGS of 150,119 individuals. All variant calls were performed jointly across individuals, allowing for consistent comparison of results. The resulting dataset provides an unparalleled opportunity to study sequence diversity in humans and its impact on phenotype variation.

Previous studies of the UKB have produced genomewide SNP array data^5^ and WES data^6, 7^. While SNP arrays typically only capture a small fraction of common variants in the genome, when combined with a reference panel of WGS individuals^8^, a much larger set of variants in these individuals can be surveyed through imputation. Imputation however misses variants private to the individuals typed only on SNP arrays and provides unreliable results for variants with insufficient haplotype sharing between carriers in the reference and imputation sets. Poorly imputed variants are typically rare, highly mutable or in genomic regions with complicated haplotype structure, often due to structural variation.

WES is mainly limited to regions known to be translate and consequently reveals only a small proportion (2-3%) of sequence variation in the human genome. It is relatively straightforward to assign function to variants inside protein coding regions, but there is abundant evidence that variants outside of coding exons are also functionally important^9–11^, explaining a large fraction of the heritability of traits^12, 13^. In particular, numerous variants are known to impact disease and other traits through their effects on non-coding genes or RNA^14^ and protein^15, 16^ expression.

Large scale sequencing efforts have typically focused on identifying SNPs and short indels. While these are the most abundant types of variants in the human genome, other types, including structural variants (SVs) and microsatellites, affect a greater number of base-pairs (bps) and consequently are more likely to have a functional impact^17, 18^. Even the SVs that overlap exons are difficult to ascertain with WES due to the much greater variability in the depth of sequence coverage in WES studies than in WGS due to the capture step of targeted sequencing. Microsatellites, polymorphic tandem repeats of 1 to 6 bps, are also commonly not examined in large scale sequence analysis studies. These variants have a higher mutation rate than SNPs and indels^19^, can affect gene expression^20^ and contribute to a range of diseases^21^.

Here, we highlight some of the insights gained from this vast new resource of WGS data that would be challenging or impossible to ascertain from WES and SNP array datasets. First, we show that exons account for a small fraction of the genomic regions displaying sequence constraint due to functional importance. Second, we describe three ancestry-based cohorts within the UKB; with 431,805, 9,633 and 9,252 individuals with British-Irish, African and South Asian ancestries, respectively. Third, using the rich UKB phenotype collection, we report novel findings from genomewide associations (GWAS) – shedding light on the impact of very rare SNPs, indels, microsatellites and structural variants on diseases and other traits.

## Results

### SNPs and indels

The whole genomes of 150,119 UKB participants were sequenced to an average coverage of 32.5x (at least 23.5x per individual, Fig. S1) using Illumina NovaSeq sequencing machines at deCODE Genetics (90,667 individuals) and the Wellcome Trust Sanger Institute (59,452 individuals). Individuals were pseudorandomly selected from the set of UKB participants and divided between the two sequencing centers. All 150,119 individuals were used in variant discovery, 13 were sequenced in duplicate, 11 individuals withdrew consent from time of sequencing to time of analysis and microarray data were not available to us for 135 individuals, leaving 149,960 individuals for subsequent analysis.

Sequence reads were mapped to human reference genome GRCh38^22^ using BWA^23^. SNPs and short indels were jointly called over all individuals using both GraphTyper^24^ and GATK HaplotypeCaller^25^, resulting in 655,928,639 and 710,913,648 variants, respectively. We used several approaches to compare the accuracy of the two variant callers, including comparison to curated datasets^26^ (Table S1, Fig. S2), transmission of alleles in trios (Table S2, Table S3), comparison of imputation accuracy (Table S4) and comparison to WES data (Table S5). These comparisons suggested that GraphTyper provided more accurate genotype calls. For example, despite there being 7.7% fewer GraphTyper variants, we estimated that GraphTyper called 4.5% more true positive variants in trios and had 9.4% more reliably imputing variants than GATK. We therefore restricted subsequent analyses of short variants to the GraphTyper genotypes, although further insights might be gained from exploring these call sets jointly. To contain the number of false positives, GraphTyper employs a logistic regression model that assigns each variant a score (AAscore) predicting the probability that it is a true positive. We focus on the 643,747,446 (98.14%) high quality GraphTyper variants, indicated by an AAscore above 0.5, hereafter referred to as GraphTyperHQ.

The American College of Medical Genetics and Genomics (ACMG) recommends reporting actionable genotypes in a list of genes associated with diseases that are highly penetrant and for which a well-established intervention is available^27^. We find that 4.1% of the 149,960 individuals carry an actionable genotype in one of 73 genes according to ACMG^27^ v3.0. Using WES^28^ and ACMG v2.0 (59 genes), 2.0% were reported to carry an actionable genotype, when restricting our analysis to ACMG v2.0 and same criteria we find 2.5% based on WGS. Increasing the number of actionable genotypes detected in a large cohort, to the extent that it could have a significant impact on societal disease burden.

The number of variants identified per individual is 40 times larger than the number of variants identified through the WES studies of the same UKB individuals (Table 1, Methods). Although referred to as “whole exome sequencing” we find that WES primarily captures coding exons and misses most variant in exons that are transcribed but not translated, missing 72.2% and 89.4%, of the 5’ and 3’ untranslated region (UTR) variants, respectively. Even inside of coding exons currently curated by Encode^9^, we estimate that 10.7% of variants are missed by WES (Table 1). Manual inspection of the missing variants in WES suggests these are missing due to both missing coverage in some regions as well as genotyping filters. Conversely, almost all variants identified with WES are found by WGS (Table 1).

**Table 1.** Overlap of WES and WGS data. Results are computed for the 109,618 samples present in both datasets and is limited to those variants that are present in at least one individual in either dataset. Numbers refer to number of variants found in dataset. WGS refers to the GraphTyperHQ dataset and WES refers to a set of 200k WES sequenced indivdiduals^76^. Missing and present percentages are computed from the number of variants in the union of the two datasets.

| | WGS | WES | WGS ∩<br>WES | Unique<br>to WES | Present<br>WES | Missing<br>WES | Present<br>WGS | Missing<br>WGS |
| --- | --- | --- | --- | --- | --- | --- | --- | --- |
| coding | 6,380,795 | 5,781,829 | 5,686,934 | 94,895 | 89.29% | 10.71% | 98.53% | 1.47% |
| splice | 445,499 | 397,226 | 388,961 | 8,265 | 87.54% | 12.46% | 98.18% | 1.82% |
| 5utr | 2,125,413 | 590,484 | 572,996 | 17,488 | 27.56% | 72.44% | 99.18% | 0.82% |
| 3utr | 7,214,427 | 764,864 | 743,790 | 21,074 | 10.57% | 89.43% | 99.71% | 0.29% |
| proximal | 249,702,570 | 6,189,465 | 5,952,145 | 237,320 | 2.48% | 97.52% | 99.91% | 0.09% |
| intergenic | 292,259,782 | 91,836 | 83,360 | 8,476 | 0.03% | 99.97% | >99.99% | <0.01% |

### Identification of functionally important regions

The number of SNPs discovered in our study corresponds to an average of one every 4.8 bp, in the regions of the genome that are mappable with short sequence reads. This amounts to detection of 7.0% of all theoretically possible SNPs in these regions (a measure of saturation). We observe 81.5% of all possible autosomal CpG>TpG variants, 11.8% of other transitions and only 4.0% of transversions (Table S6). Restricting the analysis to 17,902,255 autosomal CpG dinucleotides methylated in the germline^10^, we observe transition variants at 89.1% of all methylated CpGs. As CpG mutations are so heavily saturated (Fig. 1) the ratio of transitions to transversions (1.66) is lower than found in smaller WGS sets^1^ and de novo mutation (DNM) studies^29^.

**Fig. 1.**
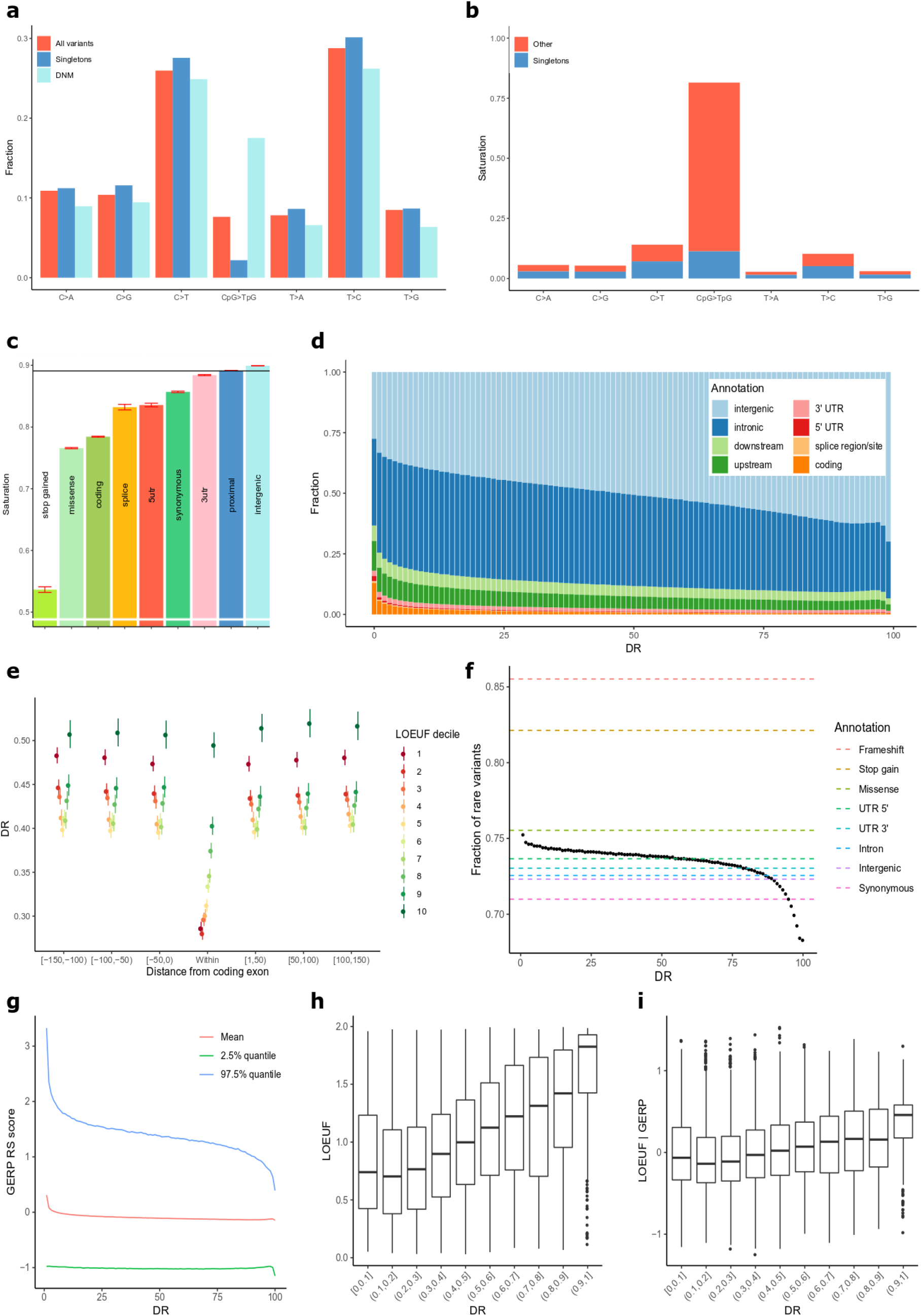
Functionally important regions a) Fraction of SNP’s in each mutation class, for all SNP’s in our dataset, singletons in our dataset, and in an Icelandic set of de novo mutations (DNMs) respectively. b) Saturation levels of mutations in each class, split into singleton variants (blue) and more common variants (red).c) Saturation levels of transitions at methylated CpG sites across genomic annotations and predicted consequence categories. The horizontal line is the average across all methylated CpG-sites.. d) Fraction of regions falling into functional annotation classes, as defined by Ensembl gene map, as a function of DR. e) DR score as a function of distance from exon and LOEUF decile f) Fraction of rare (with 4 or fewer carriers) variants (FRV) as a function of DR. g) Average GERP score in 500bp windows as a function of DR, red line represents average GERP score, blue and green line 95-th percentile. h) LOUEF and i) LOEUFashokGERP as a function of DR.

The vast majority of all variants identified are rare (Table S7), 46.0% and 40.6% of all SNPs and short indels, respectively, are singletons (carried by a single sequenced individual), and 96.6% and 91.7% have frequency below 0.1%. Inference of haplotypes and imputation typically involves identifying variants that are shared due to a common ancestor - are identical by descent. Due to the scale of the UKB WGS data, an observation of the same allele in unrelated individuals does not always imply identity by descent. A clear indication of this is that only 14% of the highly saturated CpG>TpG variants are singletons, in contrast to 47% for other SNPs (Fig. 1b). These recurrence phenomena have been described in other sample sets using sharing of rare variants between different subsets^2, 11^. We used a DNM set from 2,976 trios in Iceland^29^ to assess recurrence directly, as variants present in both that set and the UKB must be derived from at least two mutational events. Out of the 194,687 Icelandic DNMs we find 53,859 (27.7%) in the UKB set providing a direct observation of sequence variants derived from at least two mutational events. As expected, we find that CpG>TpG mutations are the most enriched mutation class in the overlap, due to their high mutation rate^30^ and saturation in the UKB set (Fig. 1b).

The rate and pattern of variants in the genome is informative about the mutation and selection processes that have shaped the genome^31^. The number of sequence variants in the exome has been used to rank genes according to their tolerance of loss-of-function (LoF) and missense variation^11, 32^. The focus on the exome is due to the availability of WES datasets and the relatively straightforward functional interpretation of coding variants. Conservation across a broad range of species^33^ is used to infer the impact of selection beyond the exome, leveraging the extensive accumulation of mutations over millions of years. However, such statistics are only partially informative about sequence conservation specific to humans^34^. Sequence variation in humans^35, 36^ can be used to characterize human specific conservation, but large sample sizes are required for accurate inference, as much fewer mutations separate pairs of humans than different species.

The extensive saturation of CpG>TpG variants at methylated CpGs in large WES cohorts has been used to identify genomic annotation or loci where their absence could be indicative of negative selection^11, 37^. In line with previous reports^11^ we see less saturation of stop-gain CpG>TpG variants than those that are synonymous (Fig. 1c). Synonymous mutations are often assumed to be unaffected by selection (neutral)^37^ however we find that synonymous CpG>TpG mutations are less saturated (85.7%) than those that are intergenic (89.9%), supporting the hypothesis that human codon usage is constrained^38^.

Extending this approach, we used sequence variant counts in the UKB to seek conserved regions in 500bp windows across the human genome. More specifically, we tabulated the number of variants in each window and compared this number to an expected number given the heptamer nucleotide composition of the window and the fraction of heptamers with a sequence variant across the genome and their mutational classes. We then assigned a rank (Depletion Rank, DR) from 0 (most depletion) to 100 (least depletion) for each 500bp window. As expected, coding exons have low DR (mean DR = 28.4), but a large number of non-coding regions show even lower DR (more depletion), including non-coding regulatory elements. Among the 1% of regions with lowest DR, 13.0% are coding and 87.0% are non-coding, with an overrepresentation of splice, UTR, gene upstream and downstream regions (Fig. 1d). DR increases with distance from coding exons (Fig. 1e). After removing coding exons, among the 1% of regions with lowest and highest DR score we see a 3.2 and 0.4-fold overrepresentation of GWAS variants, respectively (Table 2), suggesting that DR score could be a useful prior in GWAS analysis^39^. ENCODE^10^ candidate cis-regulatory elements (cCREs) are more likely than expected by chance to be found in depleted (low DR) regions (Table 2). Notably cCREs located in close proximity to transcription start sites, i.e. proximal enhancer-like and promoter-like sequences (pELS and PLS, respectively), are more enriched among depleted regions than distal enhancer-like sequences (dELSs).

**Table 2.**
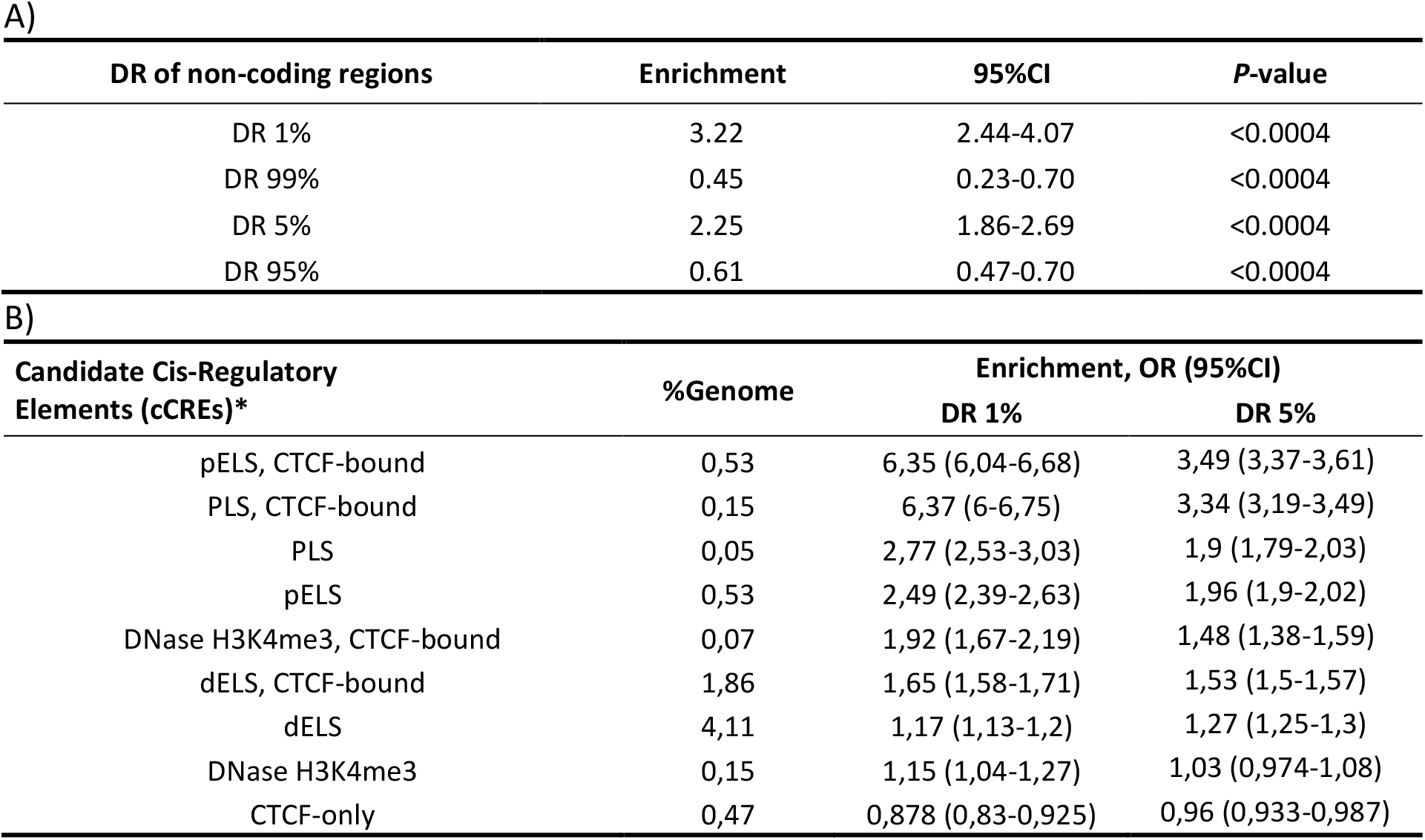
DR enrichment analysis A) Over- and underrepresentation of GWAS variants in low and high DR regions. Windows overlapping coding exons were removed. Lower DR scores indicate greater sequence conservation. B) Enrichment of ENCODÉs candidate cis-regulatory elements (cCREs) among low DR regions defined at the 1st and 5th percentile. The % of the genome covered by cCREs are indicated for each type of cCRE. *Exons of protein coding genes found in overlap with cCRE regions were removed.

| DR of non-coding regions | Enrichment | 95%CI | P-value |
| --- | --- | --- | --- |
| DR 1% | 3.22 | 2.44-4.07 | <0.0004 |
| DR 99% | 0.45 | 0.23-0.70 | <0.0004 |
| DR 5% | 2.25 | 1.86-2.69 | <0.0004 |
| DR 95% | 0.61 | 0.47-0.70 | <0.0004 |

Table 2 DR enrichment analysis A) Over- and underrepresentation of GWAS variants in low and high DR regions.
| Candidate Cis-Regulatory Elements (cCREs)* | %Genome | Enrichment, OR (95%CI) |  |
| --- | --- | --- | --- |
|  |  | DR 1% | DR 5% |
| pELS, CTCF-bound | 0,53 | 6,35 (6,04-6,68) | 3,49 (3,37-3,61) |
| PLS, CTCF-bound | 0,15 | 6,37 (6-6,75) | 3,34 (3,19-3,49) |
| PLS | 0,05 | 2,77 (2,53-3,03) | 1,9 (1,79-2,03) |
| pELS | 0,53 | 2,49 (2,39-2,63) | 1,96 (1,9-2,02) |
| DNase H3K4me3, CTCF-bound | 0,07 | 1,92 (1,67-2,19) | 1,48 (1,38-1,59) |
| dELS, CTCF-bound | 1,86 | 1,65 (1,58-1,71) | 1,53 (1,5-1,57) |
| dELS | 4,11 | 1,17 (1,13-1,2) | 1,27 (1,25-1,3) |
| DNase H3K4me3 | 0,15 | 1,15 (1,04-1,27) | 1,03 (0,974-1,08) |
| CTCF-only | 0,47 | 0,878 (0,83-0,925) | 0,96 (0,933-0,987) |

Regions under strong negative selection are expected to have a greater fraction of rare variants (FRV, defined here as variants carried by at most 4 WGS individuals) than the rest of the genome^36^. We observe a greater FRV in the most depleted regions (DR<5) than in the least depleted regions (DR>95) 74.8% vs 69.1% (Fig. 1f, Fig. S3). This is also seen when limiting to only non-coding regions (74.6% vs 69.2%). Using the FRV of annotated coding variants as a reference (Fig. 1f) we found the most depleted regions (DR < 1) to have a FRV comparable to missense mutations (75.5%).

Overall there is a weak correlation between DR and interspecies conservation as measured by GERP^33^ (linear regression (lr) r^2^ = 0.0050, two-sided (2s) p < 2.2·10^-308^, Fig. 1g). Interestingly, we find a stronger correlation between DR and GERP within coding exons (lr r^2^ = 0.0498, 2s p < 2.2·10^-308^) than outside them (lr r^2^ = 0.0012, 2s p < 2.2·10^-308^). Indicating that the correlation between DR and GERP is mostly due to the most highly conserved elements, such as coding exons, in the 36 mammalian species used to calculate GERP, with much weaker correlation in less conserved regions.

To determine whether DR reflects human specific negative selection that is not captured by GERP, we aggregated DR across the exons and compared it to the LOEUF metric from Gnomad^11^ (Fig. 1h), which measures intolerance to loss-of-function mutations. We found that DR is correlated with LOEUF (lr r^2^=0.085, 2s p < 2.2·10^-16^). LOEUF is correlated with genes demonstrating autosomal dominant inheritance^11^, in line with this we find that DR is correlated with autosomal dominant genes as reported by OMIM^40^ (Table S8). Modelling the LOEUF metric as a function of GERP and extracting the residuals from a linear fit, we obtain a measure human specific loss-of-function intolerance (LOEUF|GERP). We find DR is correlated with LOEUF|GERP (lr r^2^=0.024, 2s p < 2.2·10^-16^, Fig. 1i), indicating that DR measures human specific sequence constraint not captured by GERP. We compared DR with CDTS^35^, a measure of sequence constraint analogous to the one presented here and CADD^41^, Eigen^42^ and LINSIGHT^43^, measures of functional impact that incorporate interspecies conservation (Fig. S4). The constraint metrics that use interspecies conservation form one correlation block (GERP, CADD, Eigen and LINSIGHT) that is less correlated with the DR and CDTS correlation block (Table S9). The regions with the lowest DR score show similar enrichment across all metrics (Fig. S4). Overall, our results show that DR can be used to help identify genomic regions under constraint across the entire genome and as such provides a valuable resource for identifying non-coding sequence of functional importance.

### Multiple cohorts within UKB

Many GWAS^44^ using the UKB data have been based on a prescribed^5^ Caucasian subset of 409,559 participants who self-identified as “White British”. To better leverage the value of a wider range of of UKB participants, we defined three cohorts encompassing 450,690 individuals (Table S10), based on genetic clustering of microarray genotypes informed by self-described ethnicity and supervised ancestry inference (Methods). The largest cohort, XBI (Fig. S6), contains 431,805 individuals, including 99.6% of the 409,559 prescribed Caucasian set, along with around 23,900 additional individuals previously excluded because they did not identify as “White British” (thereof 13,000 who identified as “White Irish”). A principal components analysis (PCA) of the 132,000 XBI individuals with WGS data (Methods), based on 4.6 million loci, reveals an extraordinarily fine-scaled differentiation by geography in the British–Irish Isles gene pool (Fig. S5).

We defined two other cohorts based on ancestry: African (XAF, N=9,633,Fig. S7) and South Asian (XSA, N=9,252, Fig. S8) (Fig. 2a,b,c). The 37,598 UKB individuals who do not belong to XBI, XAF or XSA were assigned to the cohort OTH (others). The WGS data of the XAF cohort represents one of the most comprehensive surveys of African sequence variation to date, with reported birthplaces of its members covering 31 of the 44 countries on mainland sub-Saharan Africa (Fig. S7). Due to the considerable genetic diversity of African populations, and resultant differences in patterns of linkage disequilibrium, the XAF cohort may prove valuable for fine-mapping association signals due to multiple strongly correlated variants identified in XBI or other non-African populations.

**Fig. 2.**
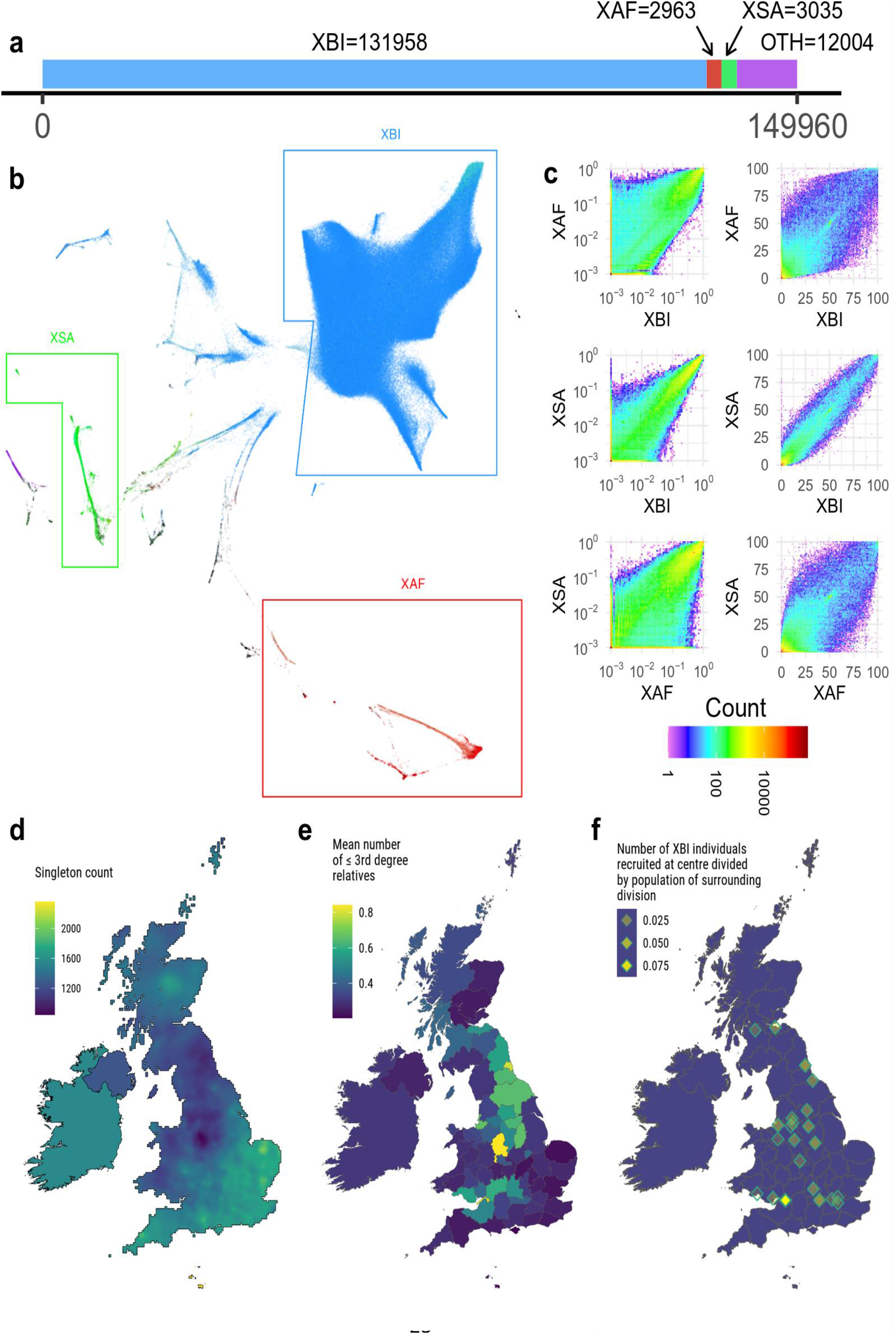
Cohort characteristics a) The number of WGS samples analyzed for phenotypes in our study. b) UMAP plot generated from the first 40 principal components of all UKB participants, colored by self-reported ethnicity: blue shades for ethnic labels under the White category, red shades for Black, and green shades for South Asian; for full color legend see Fig. S28. c) Joint frequency spectrum of variants on chr20 between all pairs of populations. Panels d, e and f show characteristic of XBI cohort across Great Britain and Ireland d) Number of singletons carried by individuals in the XBI cohort as a function of place of birth. e) Mean number of 3^rd^ degree relatives by administrative division f) Location of UKB assessment centers and estimated fraction of surrounding population recruited to the UKB. Differences in singleton counts and number of third relatives are likely a result of denser sampling of individuals living near UKB assessment centers.

We crossed GraphTyperHQ variants with exon annotations and found that on average around one in thirty individuals is homozygous for rare (minor allele frequency, MAF < 1%) LoF mutations in the homozygous state and the median number of heterozygous rare LoF is 24 per individual. We detect rare LoF variants in 19,105 genes, whereof 2,017 genes had homozygous carriers of rare LoFs (n individuals = 5,102). A marked difference in the number of homozygous LoFs carriers was found between the cohorts, with XSA having the largest fraction of homozygous LoF carriers (Fig. S9b). A notable feature of the XSA cohort is elevated genomic inbreeding, likely due to endogamy^45^, particularly among self-identified Pakistanis^46^ (Fig. S9a).

On average, individuals carried alternative alleles for 3,410,510 SNPs and indels (Fig. 3a), per haploid genome. A greater number of variants are generally found in individuals born outside of Europe (Fig. S10), because the human reference genome is primarily derived from individuals of European ancestry^22^. XAF individuals carry the greatest number of alternative alleles (Fig. 3a). We constructed cohort specific DRs and find that XAF shows greater depletion around exons than XBI and XSA (Fig. S11). Largely due to variation in the number of individuals sampled, the average number of singletons per individual varies considerably by ancestry (Fig. 3a). Thus, individuals from the XBI, XAF and XSA cohorts have an average of 1,330, 9623 and 8340 singleton variants, respectively. In XBI, singleton counts (Fig. 2d) indicate that the expected number of new variants discovered per genome is still substantial, but varies geographically, averaging around 1,000 in Northern England and 2,000 South-Eastern England. This pattern is largely explained by denser sampling of some regions (Fig. 2e,f) rather than regional ancestry differences.

**Fig. 3.**
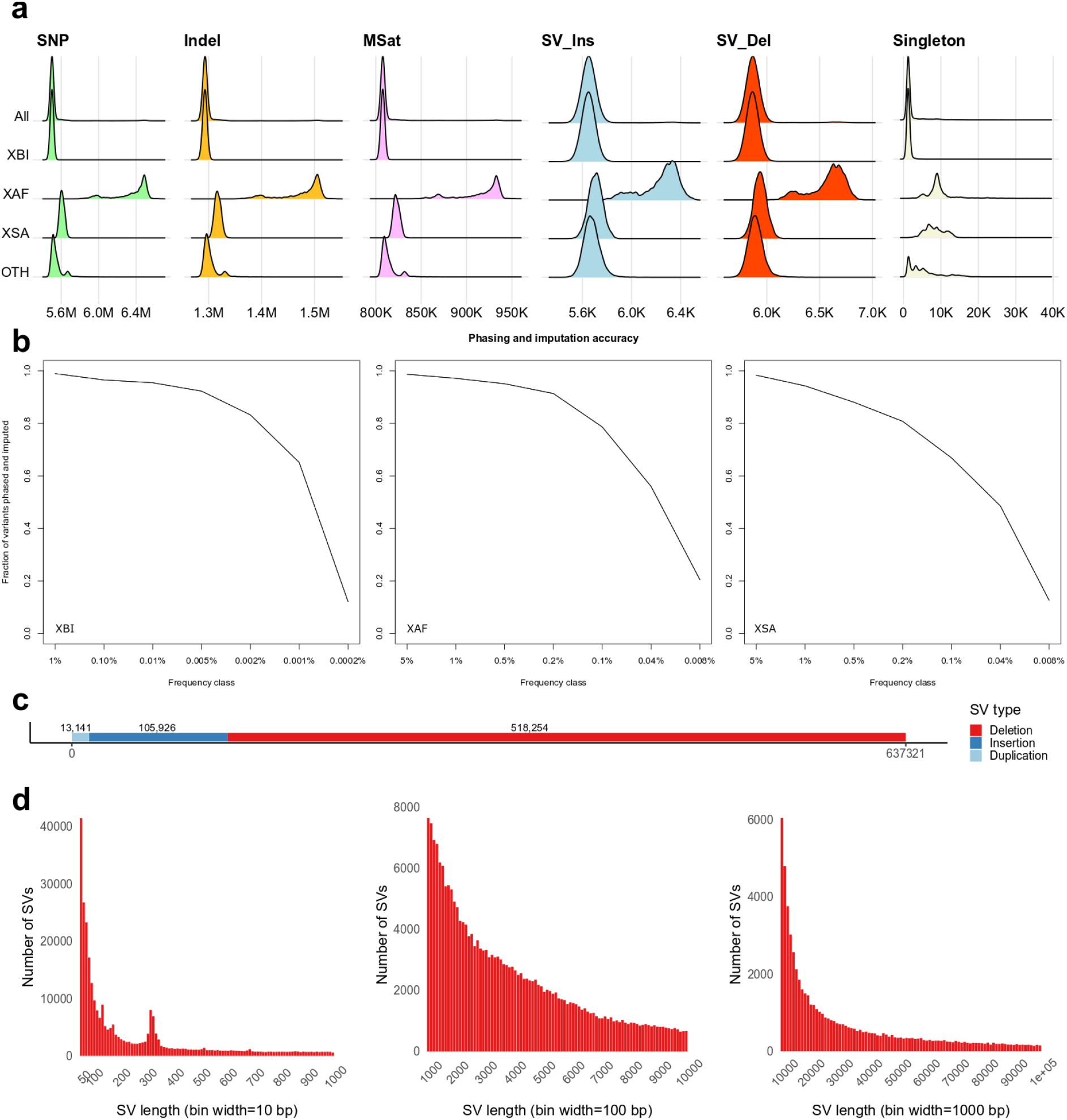
Variant call set a) Number of SNPs, Indels, microsatellites, SV insertions, SV deletions and singleton SNPs carried per diploid genome of individuals in the overall set and partitioned by population. b) Imputation accuracy in the three populations, XBI, XAF and XSA. A variant was considered imputed if “Leave one out r2” of phasing was greater than 0.5 and imputation information was greater than 0.8. x-axis splits variants into frequency classes based on the number of carriers in the sequence dataset. Variants are split by variant type. c) Number of structural variants (SVs) discovered in the dataset by variant type. d) Length distribution of SVs, from 50-1,000 bp, 1,000-10,000bp and 10,000-100,000bp.

### Imputation

We were able to reliably impute variants into the entire UKB sample set down to very low frequency (Fig. 3b). We imputed phased genotypes which permit analysis that depend on phase such as identification of compound LoF heterozygotes. A single reference panel was used to impute into the genomes of all participants in UKB, but results are presented separately for the three cohorts (Table S11). This reference panel can be used for accurate imputation in individuals from the UK and many other populations. In the XBI cohort, 98.5% of variants with frequency above 0.1% and 65.8% of variants in the frequency category of 0.001-0.002% (representing 3-5 WGS carriers) could be reliably imputed (Fig. 3b). Variants were also imputed with high accuracy in XAF and XSA (Fig. 3b), where 97.5% and 94.9% of variants in frequencies 1-5% and 56.6% and 48.9% of variants carried by 3-5 sequenced individuals could be imputed, respectively. A larger number of variants, particularly rare ones, are imputed for all cohorts than when using a alternate imputation panel^5^ (Table S12). It is thus likely that the UKB reference panel provides one of the best available option for imputing genotypes into population samples from Africa and South Asia.

We found a number of clinically important variants that can now be imputed from the dataset. These include rs63750205 (NM_000518.5(HBB):c.*110_*111del) in the 3‘ UTR of HBB, a variant that has been annotated in ClinVar^47^ as likely pathogenic for beta Thalassemia. rs63750205-TTA has 0.005% frequency (freq) in the imputed XBI cohort (imputation information (imp info) 0.98) and is associated with lower mean corpuscular volume by 2.88 s.d. (95% CI 2.43-3.33, 2s p = 1.5·10^-36^, χ^2^).

In the XSA cohort we found rs563555492-G, a previously reported^48^ missense variant in *PIEZO1* (freq = 3.65% XSA, 0.046% XAF, 0.0022% XBI) associated with higher haemoglobin concentration, effect 0.36 s.d. (95% CI 0.28-0.44, 2s p = 8.9·10^-19^, χ^2^). The variant can be imputed into the XSA population with imp info of 0.99.

In the XAF cohort we found the stop gain variant rs28362286-C (p.Cys679Ter) in *PCSK9* (freq = 0.93% XAF, 0.00016% XBI, 0.0070% XSA) imputed in the XAF cohort with imp info 0.93. The variant lowers non-HDL cholesterol by 0.92 s.d. (95% CI 0.75-1.09, 2s p = 2.3·10^-26^, χ^2^). We found a single homozygous carrier of this variant, which has 2.5 s.d. lower non-HDL cholesterol than the population mean, is 61 years old and appears to be healthy.

### SNP and indel associations not present in WES data

We highlight three examples of associations of SNPs and indels associated with traits in the XBI cohort that could not be easily identified in WES or SNP array data.

The first is an association in the XBI cohort between a rare variant rs117919628-A (freq = 0.32%; imp info = 0.90) in the promoter region of GHRH, encoding the growth hormone-releasing hormone close to one of its TSS (Transcription start site) and less height (effect = - 0.32 s.d. (95% CI 0.27-0.36), 2s p = 1.6·10^-39^, χ^2^). GHRH is a neuropeptide secreted by the hypothalamus to stimulate the synthesis of growth hormone (GH). We note that the effect (-0.32 s.d. or -3cm) of rs117919628 is greater than any variants reported in large height GWAS (∼1200 associated variants)^49–51^. In addition to reducing height, rs117919628-A is associated with lower IGF-1 serum levels (Insulin-growth factor 1, effect = -0.36 s.d. (95% CI 0.32-0.40), 2s p = 3.2·10^-58^, χ^2^). The production of IGF-1 is stimulated by GH and mediates the effect of GH on childhood growth, further supporting *GHRH* being the gene mediating the effects of rs117919628-A. Due to its location around 50 bp upstream of the *GHRH* 5‘UTR, this variant is not targeted by the UKB WES, and neither is the only strongly correlated variant rs372043631 (intronic). The height associations of these two variants have not been reported, presumably because they are absent from all versions of the 1,000 Genomes data^52^ and in imputations based on the haplotype reference consortium/UK 10K^53^ (HRC/UK10K) these two variants have low imp info (0.54) and may thus fail quality checks. rs117919628-A is not correlated with rs763014119-C (no individuals carry the minor allele of both variants), a previously reported^54^ very rare frameshift deletion in *GHRH* (Phe7Leufster2; freq = 0.0092%), associated with reduced height and IGF-1 levels (height effect = -0.63 s.d (95% CI 0.36-0.89), 2s p = 4.6·10^-6^; IGF-1 effect = -0.74 s.d. (95% CI 0.49- 0.99), 2s p = 4.9·10^-9^, χ^2^).

The second example is rs939016030-A a rare 3‘ UTR essential splice acceptor variant in the gene encoding tachykinin 3 (*TAC3*; freq = 0.033%; c.*2-1G>T in NM_001178054.1 and NM_013251.3). The XBI cohort has 89 WGS carriers and 281 in the imputation set. This variant is not found in WES of the UKB^53^ and neither are the two highly correlated variants, one intronic (rs34711498) and one intergenic (rs368268673). These 3 variants were absent from the HRC/UK10K^55^ imputation, and are only present in Europeans, with highest frequency in the UK according to Gnomad^11^. The minor allele of this 3‘UTR essential splice variant rs939016030-A is associated with later age of menarche, with an effect of 0.57 s.d. (95% CI 0.41-0.74) or 11 months (2s p = 1.0·10^-11^, χ^2^). Rare coding variants in *TAC3* and its receptor *TACR3* are reported to cause hypogonadotropic hypogonadism^56^ under autosomal recessive inheritance. However, in the UKB, the association of the 3’UTR splice acceptor variant, is only driven by heterozygotes (∼ 1 in 1500 individuals) with no homozygotes detected. We replicated this finding in a set of 39,360 Danes, with an effect of 0.70 s.d. (95% CI 0.34-1.06, freq = 0.05%, 2s p = 0.00014, χ^2^).

The third example is a rare variant (rs1383914144-A; freq = 0.40%) near the centromere of chromosome 1 (start of 1q), that associates with lower uric acid (UA) levels (effect = -0.43 s.d. (95% CI 0.40-0.46) or -0.58 mg/dL (95% CI 0.54-0.62), 2s p = 8.1·10^-170^, χ^2^) and protection against gout (OR = 0.36 (95% CI 0.28-0.46), 2s p = 4.2·10^-15^, χ^2^). A second variant rs1189542743, 4Mb downstream at the end of 1p is strongly correlated with rs1383914144 (r^2^ = 0.68) and yields a similar association with uric acid. Neither variant is targeted by UKB WES nor imputed by the HRC/UK10K and no association was reported in this region in the uric acid GWAS^57^. The effect of rs1383914144-A on uric acid is larger than for any variant reported in the latest GWAS meta-analysis of this trait. We replicate these findings in Iceland (rs1383914144-A, freq = 0.47%; 2s p (UA) = 8.0·10^-37^, χ^2^ and effect (UA) = - 0.51 s.d. (95% CI 0.43-0.59), 2s p (Gout) = 0.0018, χ^2^, OR (Gout) 0.31 (95% CI 0.15-0.64)) and (rs1383914144-A, freq = 0.47%; 2s p (UA) = 1.1·10^-36^, χ^2^ and effect (UA) = - 0.51 s.d. (95% CI 0.43-0.59), 2s p (Gout) = 0.0018, χ^2^, OR (Gout) 0.31 (95% CI 0.15-0.64)).

### Structural variants play an important role in human genetics

We identified structural variants (SVs) in each individual using Manta^58^ and combined these with variants from a long read study^59^ and the assemblies of seven individuals^60^. We genotyped the resulting 895,055 SVs (Fig. 3c) with GraphTyper^60^, of which 637,321 were considered reliable.

On average we identified 7,963 reliable SVs per individual, 4,185 deletions and 3,778 insertion (Fig. 3a). These numbers are comparable to the 7,439 SVs per individual found by Gnomad-SV^61^, another short read study, but considerably smaller than the 22,636 high quality SVs found in a long read sequencing study^59^, mostly due to an underrepresentation of insertions and SVs in repetitive regions. SVs show a similar frequency distribution as SNPs and indels and a similar distribution of variants across cohorts (Fig. 3a).

We present four examples of phenotype associations with structural variants, not easily found in WES data. First, a rare (freq=0.037%) 14,154 bp deletion that removes the first exon in *PCSK9*, previously discovered using long read sequencing in the Icelandic population and is associated with lower non-HDL cholesterol levels^59^. There were thirty two WGS carriers in the XBI cohort (freq 0.012%) and 72 carriers in the XBI imputed set (freq 0.0087%) who had 1.22 s.d. (95% CI 0.90-1.55) lower non-HDL cholesterol levels than non-carriers (2s p = 1.2·10^-13^, χ^2^).

The second example is a 4,160 bp deletion, (freq = 0.037% in XBI), that removes the promoter region from 4,300 to 140 bp upstream of the *ALB* gene that encodes Albumin. Not surprisingly, carriers of this deletion have markedly lower serum albumin levels (effect 1.50 s.d. (95% CI 1.35-1.62) 2s p = 9.5·10^-118^, χ^2^). The variant is also associated with traits correlated with albumin levels; carriers had lower calcium and cholesterol levels: 0.62 s.d. (95% CI 0.50-0.75, 2s p = 2.9·10^-22^, χ^2^) and 0.45 s.d. (95% CI 0.30-0.59, 2s p = 1.1·10^-9^, χ^2^), respectively.

The third SV example is a 16,411 bp deletion (freq = 0.0090% in XBI) that removes the last two exons (4 and 5) of *GCSH*, that encodes Glycine cleavage system H protein. Carriers of this deletion have markedly higher Glycine levels in the UKB metabolomics dataset (effect 1.45 s.d. (95% CI 1.01-1.86), 2s p = 1.2·10^-10^, χ^2^).

The final example is a rare (freq 0.892% in XBI) 754bp deletion overlapping exon 6 of *NMRK2*, encoding nicotinamide riboside kinase 2 that removes 72 bp from the transcribed RNA that corresponds to a 24 amino acid inframe deletion in the translated protein. Carriers of this deletion have a 0.22 s.d. (95% CI 0.18-0.27) earlier age at menopause (2s p = 1.1·10^-26^, χ^2^). Nearby is the variant rs147068659, reported to be associated with this trait^62^, with an effect 0.20 s.d. (95% CI 0.16-0.24) earlier age at menopause (2s p = 2.0·10^-20^, χ^2^) in the XBI cohort. The deletion and rs147068659 are correlated (r^2^ = 0.67), after conditional analysis the deletion remains significant (2s p = 6.4·10^-8^, χ^2^) whereas rs147068659 does not (2s p = 0.39, χ^2^), indicating the deletion is the lead variant for the locus. *NMRK2* is primarily expressed in heart and muscle tissue^63^. In our dataset of right atrium heart tissue, one individual out of a set of 169 RNA sequenced individuals is a carrier of this deletion. As expected we observe decreased expression of exon 6 in this individual (Fig. S12) and an increase in the fraction of transcript fragments skipping exon 6 (Fig. S13).

### Microsatellites are commonly overlooked

We identified 14,321,152 alleles at 2,536,688 microsatellite loci using popSTR^64^ in the 150,119 WGS individuals, who carry on average of 810,606 non-reference microsatellite alleles. The number of non-reference alleles carried per individual shows a similar distribution across the UKB cohorts as other variant types characterized in this study (Fig. 3a). Microsatellites are among the most rapidly mutating variants in the human genome and a source of genetic variation that is usually overlooked in GWAS. Repeat expansions are known to associate with a number of phenotypes, including Fragile X syndrome^65^. We are able to impute microsatellites down to a very low frequency (Fig. S14) in all three cohorts, providing one of the first large scale datasets of imputed microsatellites.

We genotyped a microsatellite within the *CACNA1A* gene that encodes voltage-gated calcium channel subunit alpha 1A. Individuals who have twenty or more repeats of this microsatellite generally suffer from lifelong conditions that affect the brain, including Familial hemiplegic migraine (FHM1), Epilepsy, Episodic Ataxia Type 2 (EA2) and Spinocerebellar ataxia type 6 (SCA6)^66–69^. Carriers in the XBI cohort of 22 copies of the microsattelite repeat were at greater risk for hereditary ataxia (freq = 0.0071%, OR = 304, 2s p = 1.1·10^-31^, χ^2^).

We also confirm an association between a microsatellite within the 3‘ UTR of *DMPK*, encoding DM1 protein kinase, and myotonic dystrophy in the XBI cohort. Expression of *DMPK* is negatively correlated with the number of repeats of the microsatellite^70^. The risk of myotonic dystrophy increases with copy number of the repeats, rising rapidly with the number of repeats carried by an individual up to an odds ratio of 161 for individuals carrying 39 or more repeats (Table S13, Fig. S15).

### Variants that are not imputed

Although the vast majority of WGS variants can be imputed to the larger set of SNP array genotyped individuals it is interesting to examine the variants that are not imputed. A subset of these variants are in regions where there are no nearby variants present in the SNP array data and regions where there are disagreements between the GRCh38^22^ and CHM13^71^ assemblies. Lifting variants over to the CHM13 assembly may allow us to impute a subset of these variants. The failure of those variants to impute on GRCh38 can presumably be attributed to a misassembly on GRCh38. In addition, we identify a number of variants that are most likely recurrently somatic, such as the gain of function mutations in JAK2^72–74^ and CALR^74^ know to be associated with myeloproliferative disorders, including polycythaemia vera and essential thrombocythemia.

## Discussion

The dataset provided by sequencing the whole genomes of 150 thousand UKB participants is unparalleled in its size and provides the most extensive characterization of the sequence diversity in the germline genomes of a single population to date. The UK population is diverse in its genetic ancestry and includes individuals born in countries all over the globe. The African and South Asian ancestry cohorts each number over 9,000 individuals, represent some of the largest available WGS sets of these ancestries and which are likely to have an impact both clinically and in further characterizing the relationship between sequence and traits.

We characterized an extensive set of sequence variants in the WGS individuals, providing two sets of SNP and indel data, as well as microsatellite and SV data, variant classes that are frequently not interrogated in GWAS. We give examples of how these variants play a role in the relationship between sequence and phenotypic variation. Further discoveries may be made by relating the variants presented here to alternate annotations (Table S14), but more importantly we believe there are many other discoveries to be made. The number of SNPs and indels are 40-fold greater than from WES of the same individuals. Even within annotated coding exons WES misses 10.7% of variants, found through WGS. WES misses most of the remainder of the genome, including functionally important UTR, promoter regions and exons yet to be annotated. The importance of these regions is exemplified by the discovery of rare non-coding sequence variants with larger effects on height and menarche than any variants described in GWAS to date.

The DR score presented here is an important resource for identifying genomic regions of functional importance. Although coding exons are clearly under strong purifying selection, as represented by a low DR score, they represent only a small fraction of the regions with low DR score. Clinical geneticists typically focus on coding exons and have only been able to identify the causal variant in fewer than half of clinical cases studied. Currently, 98.4% of variants annotated as pathogenic in the ClinVar^47^ database are within coding exons. Greater attention should be given to other regions of the genome, particularly those with low DR score, where non-coding exons (UTRs), enhancer and promoter regions are overrepresented.

There are still some sequence variants that are not found with short read WGS, including VNTRs, repetitive regions and regions that have only recently been captured by human genome assemblies^71^. Improved assembly^71, 75^, sequencing and representation of the genome and its variation will have important implications for advancing our understanding of the relationship between sequence diversity and human diseases and other traits.

A near complete sequence of the human genome has been known for over twenty years. Genome scientists have yet to assign function to a large fraction of this sequence and have had only partial success in understanding the genetic source of phenotypic diversity. The large-scale sequencing described here, as well as the continued effort in sequencing the entire UKB, promises to vastly increase our understanding of the function and impact of the non-coding genome. When combined with the extensive characterization of phenotypic diversity in the UKB, these data should greatly improve our understanding of the relationship between human genome variation and phenotype diversity.

## Author Contributions

Paper was written by BVH and KS with input from HPE, KHSM, OE, DFG, OTM, GM, UT, AH, HJ and PS. KHSM and AH defined cohorts. OE and HJ identified functionally important regions. FJ and UT were responsible for laboratory operations. DNA sequencing was performed by DNM, SS, KK and OTM. Sample isolation was performed by ES and JS. GM was responsible for the sequence analysis pipeline, developed by BVH, AG, PIO, MA, STS, FZ and SAG, and run by GTS. BVH, HPE, HHa, GP, SK, GH and SAG developed analysis tools. Association analysis was performed by BVH, MOU, AO, BOJ, SK, BDS, DB, VT, US and PS. Phenotypes were defined by MOU, VT, GT, IJ, TR, HHo, HS and PS. SNP and SV genotyping was performed by HPE, PIO and BS. Microsatellite genotyping was performed by SK. Data analysis was performed by BVH, HPE, HHa, GP, AO, OAS, GS and KN. RNA sequence data was analyzed by GHH, supervised by PM. DFG supervised association, data and DR analysis. Figures were drawn by MTH and KHSM. HJ, AJG, IO, PJ collected clinical data in Iceland. OBP, CE, SB, SRP and DBDSGC collected clinical data in Denmark. Study was supervised by BVH and KS. All authors agreed to the final version of the manuscript.

## Data availability

WGS and genotype data can be accessed via the UKB research analysis platform (RAP). DR score will be made available along with final publication of manuscript.

## Code availability

BamQC, https://github.com/DecodeGenetics/BamQC.

GraphTyper, https://github.com/DecodeGenetics/graphtyper.

GATK resource bundle, gs://genomics-public-data/resources/broad/hg38/v0.

Svimmer, https://github.com/DecodeGenetics/svimmer.

popSTR, https://github.com/DecodeGenetics/popSTR.

Dipcall, https://github.com/lh3/dipcall.

RTG Tools, https://github.com/RealTimeGenomics/rtg-tools.

bcl2fastq, https://support.illumina.com/sequencing/sequencing_software/bcl2fastq-conversion-software.html.

Samtools, http://www.htslib.org/.

Samblaster https://github.com/GregoryFaust/samblaster.

## Ethics declaration

A number of authors are employees of deCODE genetics/Amgen.

## Acknowledgements

We thank the participants of the UKB. The sequencing of 450,000 WGS individuals from the UKB, including the 150,119 described here has been funded by the UKB WGS consortium consisting of UK Government’s research and innovation agency, UK Research and Innovation (UKRI), through the Industrial Strategy Challenge Fund, The Wellcome Trust and the pharmaceutical companies Amgen, AstraZeneca, GlaxoSmithKline and Johnson & Johnson. DNA sequenced was performed at the Welcome Trust Sanger Institute and deCODE genetics.

## Supplementary material

### Supplementary Figures

**Fig. S1.**
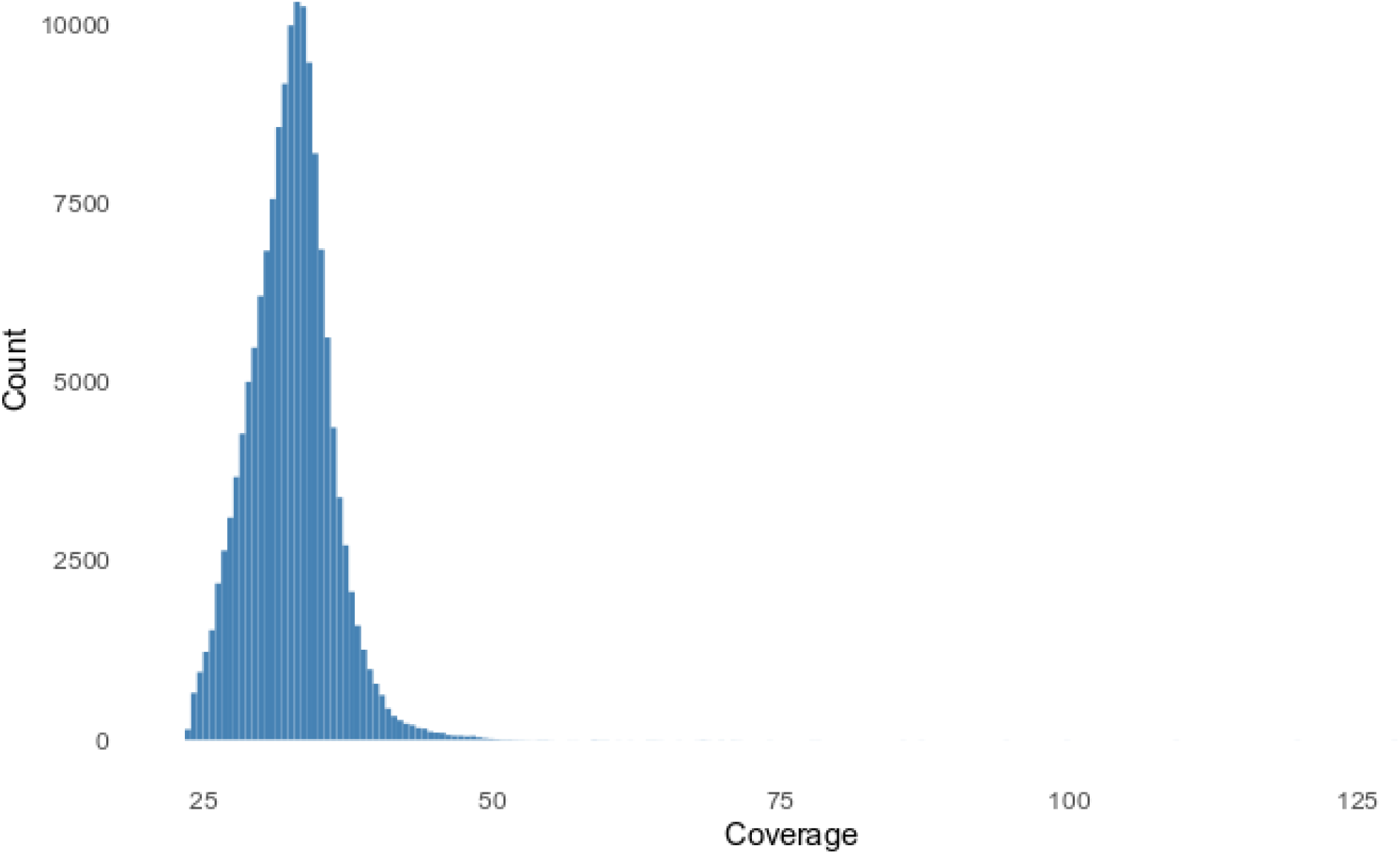
Histogram of average sequence coverage per sample in the 150,119 WGS samples.

**Fig. S2.**
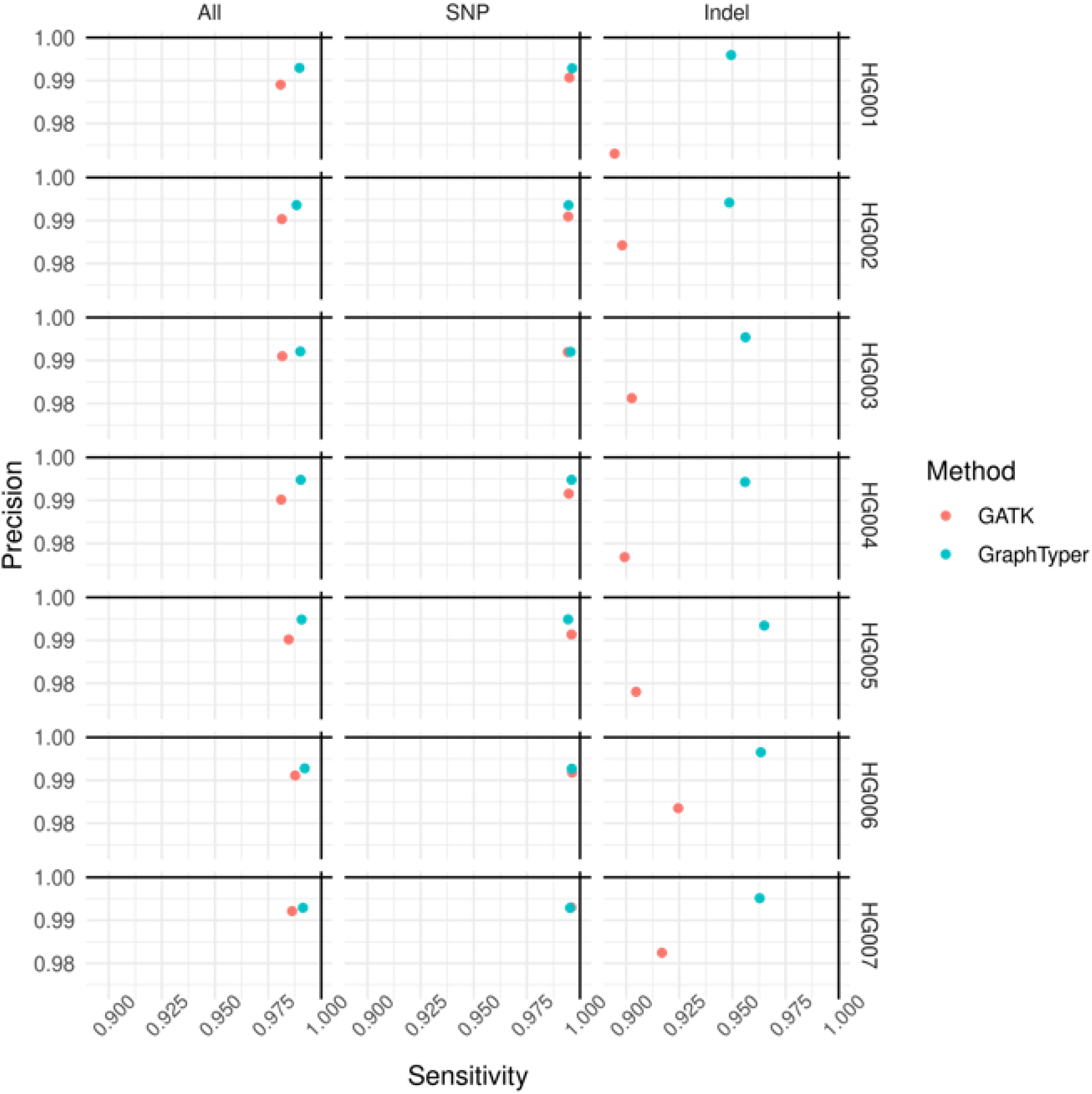
Sensitivity and precision for GATK and GraphTyper callsets in 500 regions benchmarking dataset across the seven Genome in a bottle (GIAB) v3.3.2 truth sets.

**Fig. S3.**
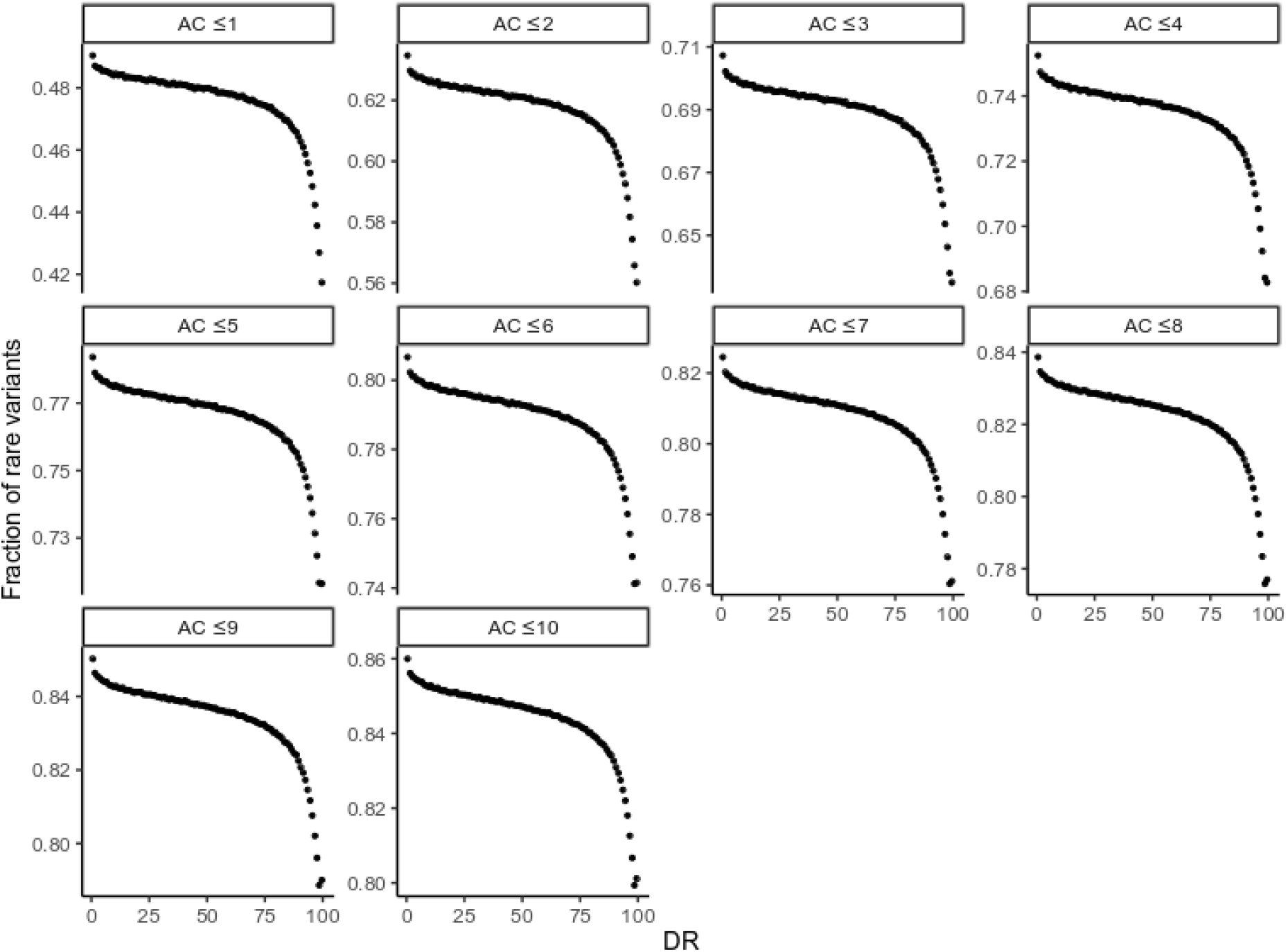
Fraction of rare variants (FRV) as a function of the definition of “rare”, varying the allele count cutoff from at most 1 to at most 10 carriers. Note that homozygous carriers have an allele count of 2.

**Fig. S4.**
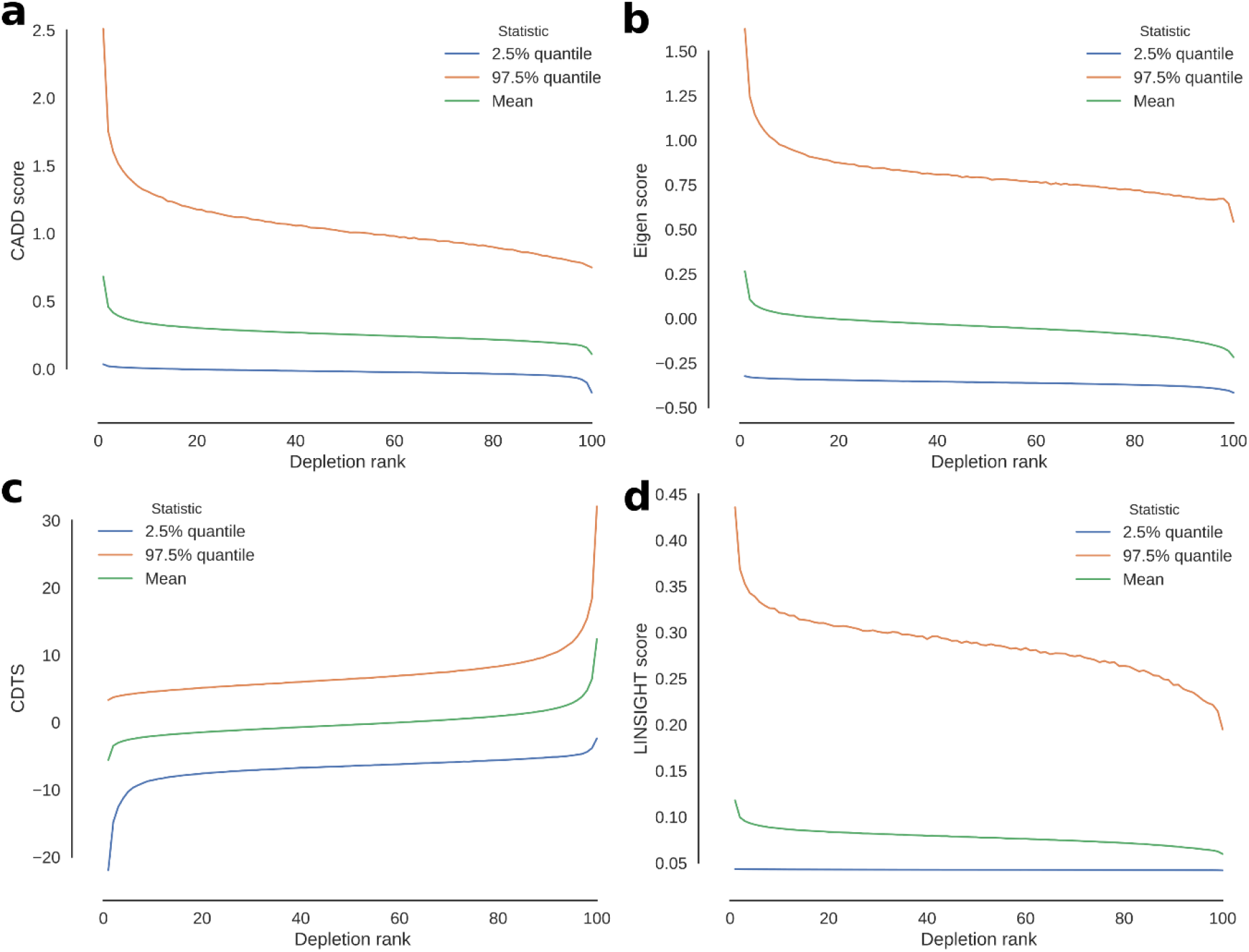
Average score in 500bp windows as a function of Depletion Rank for a) CADD b) Eigen c) CDTS and d) LINSIGHT. Green line represents average score, blue and red line 95-th percentile

**Fig. S5.**
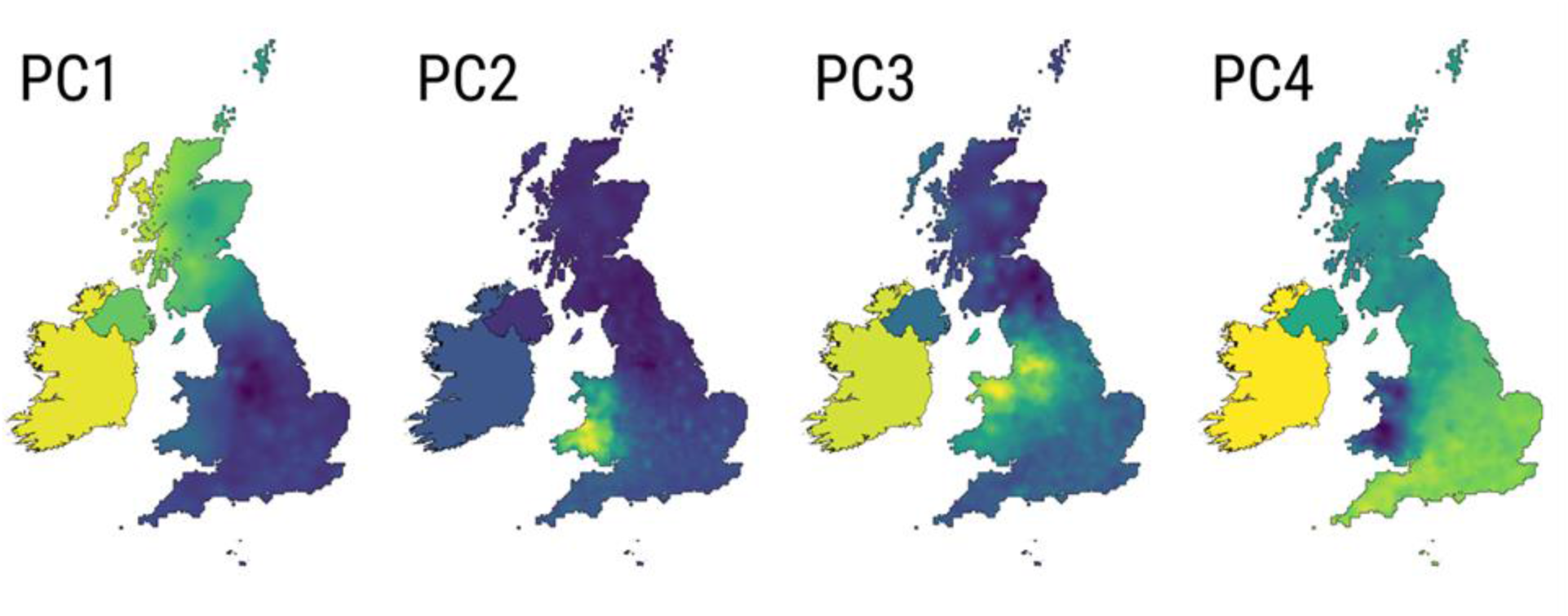
Geographic distribution of the loadings of the first four principal components of a PCA of the XBI population.

**Fig. S6.**
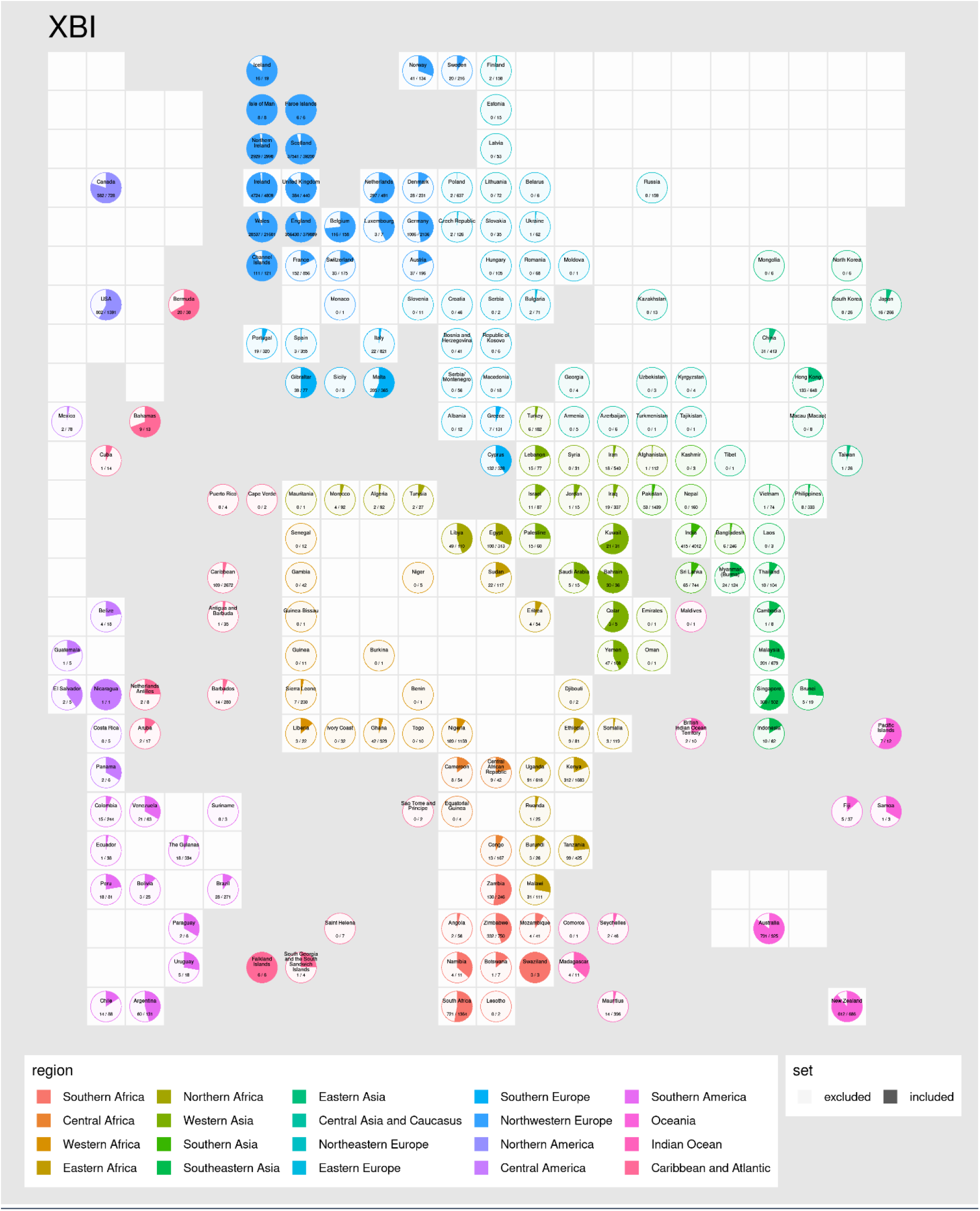
Cartogram-pies indicating the proportion of individuals born in each country (name shown on top of pies) in the XBI cohort. Pies are placed roughly according to their country’s position on a world map. Grey and white squares represent sea and land respectively.

**Fig. S7.**
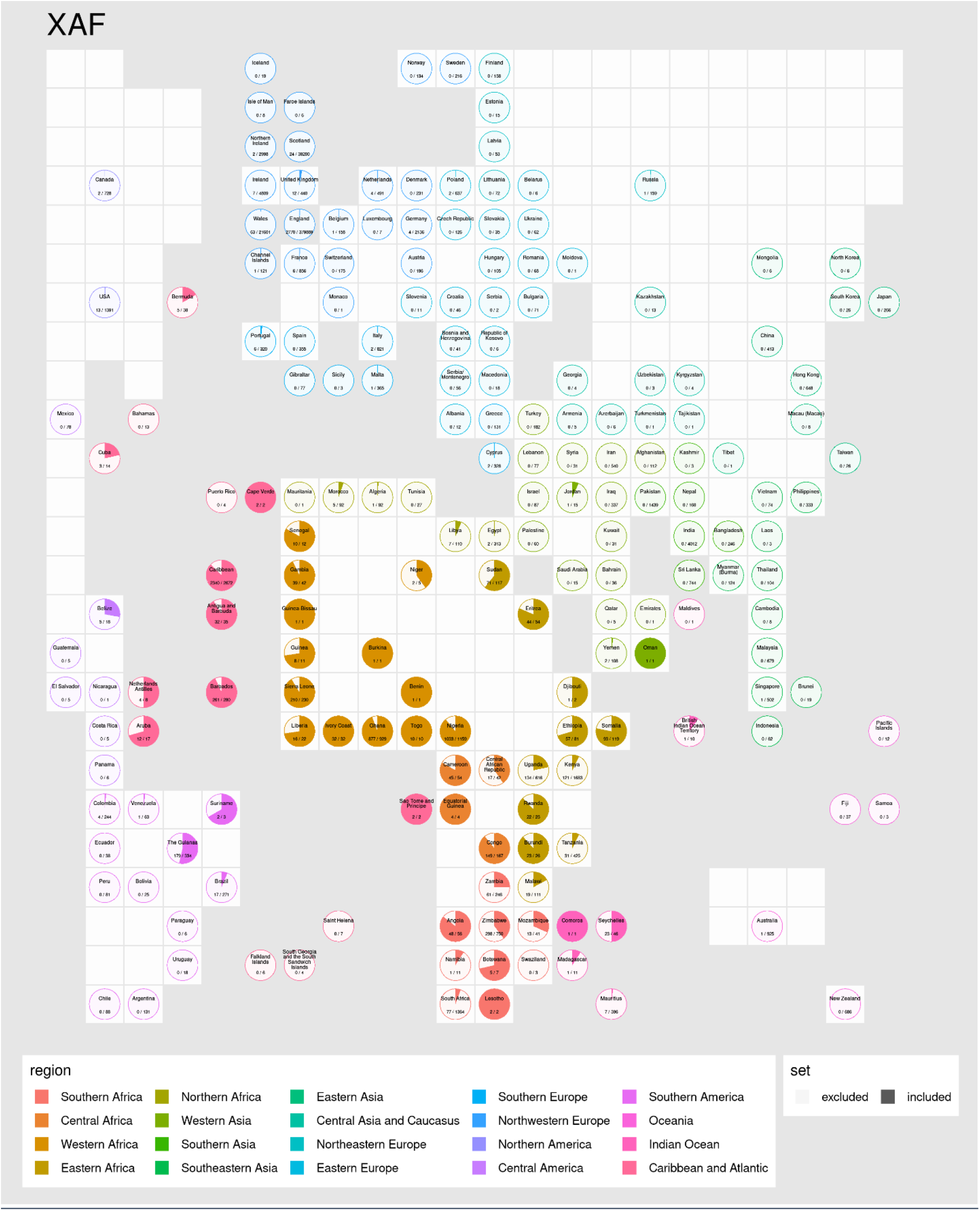
Cartogram-pies indicating the proportion of individuals born in each country (name shown on top of pies) in the XAF cohort. Pies are placed roughly according to their country’s position on a world map. Grey and white squares represent sea and land respectively.

**Fig. S8.**
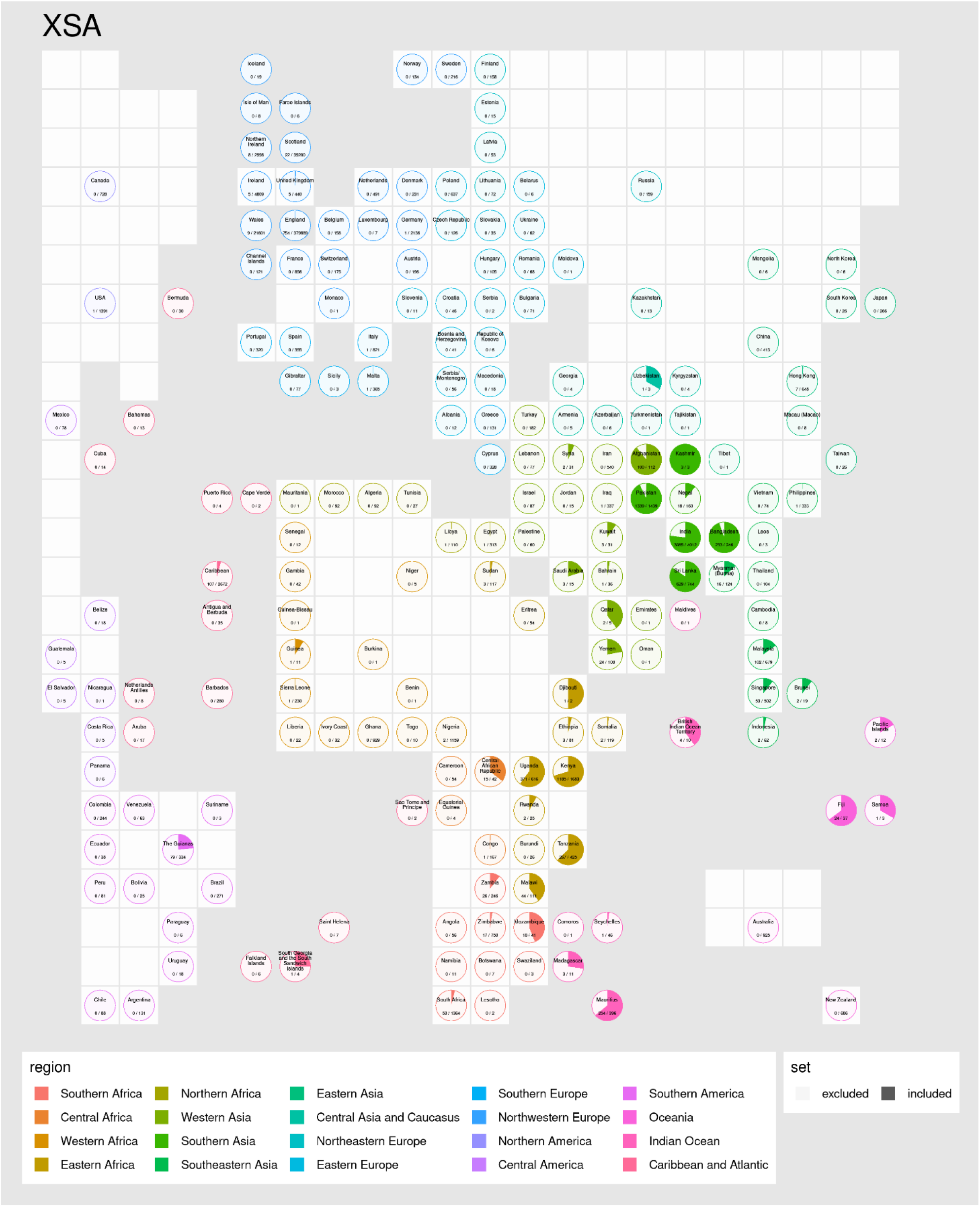
Cartogram-pies indicating the proportion of individuals born in each country (name shown on top of pies) in the XSA cohort. Pies are placed roughly according to their country’s position on a world map. Grey and white squares represent sea and land respectively.

**Fig. S9.**
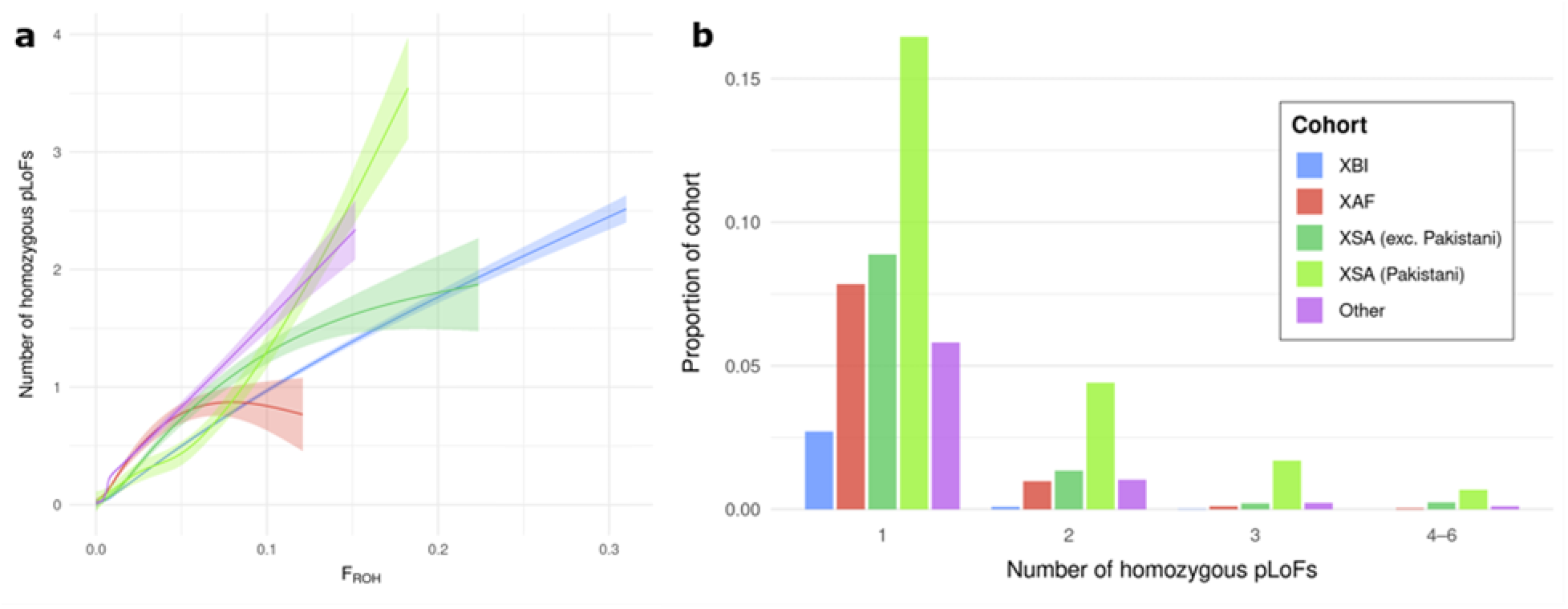
Loss-of-function a) Correlation between the number of LoF genes per sample and fraction of genome with runs of homozygosity. b) Number of homozygous loss-of-function (LoF) genes per sample. Count of homozygous genes annotated as high impact with frequency <1%. Results are presented for XBI, XAF, XSA excluding individuals self-identified as Pakistani, individuals self-identified as Pakistani from the XSA cohort and Others.

**Fig. S10:**
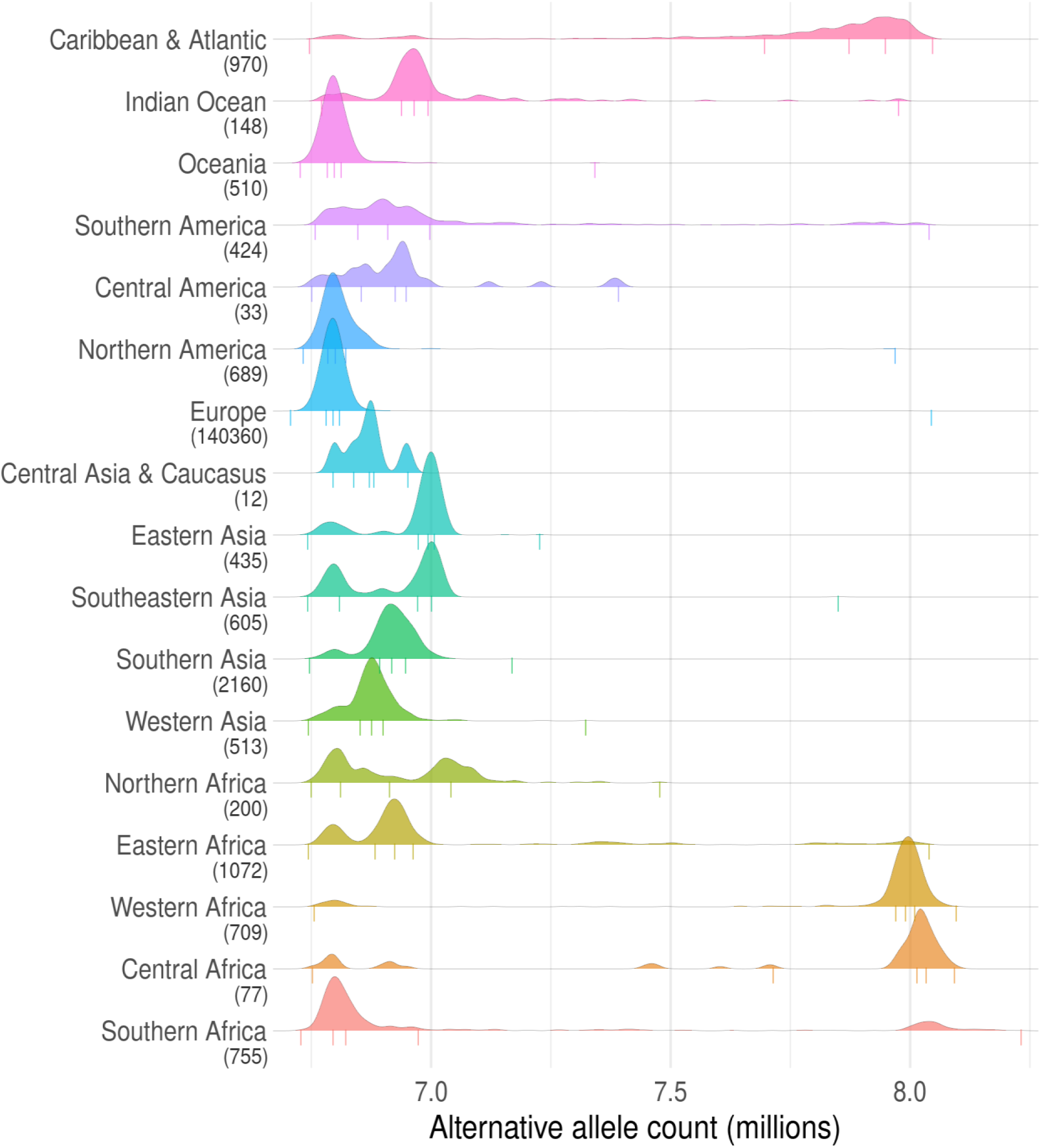
Alternative alleles by region. Numbers in brackets beneath region names indicate count of whole genome sequenced individuals with birthplaces in that region. Assignment of countries to regions is almost identical to the categorization displayed in the cohort cartogram pie figures, with the exception that all European regions are combined into one region in this figure. Vertical lines underneath density curves represent 0th, 25th, 50th, 75th, and 100th percentiles.

**Fig. S11.**
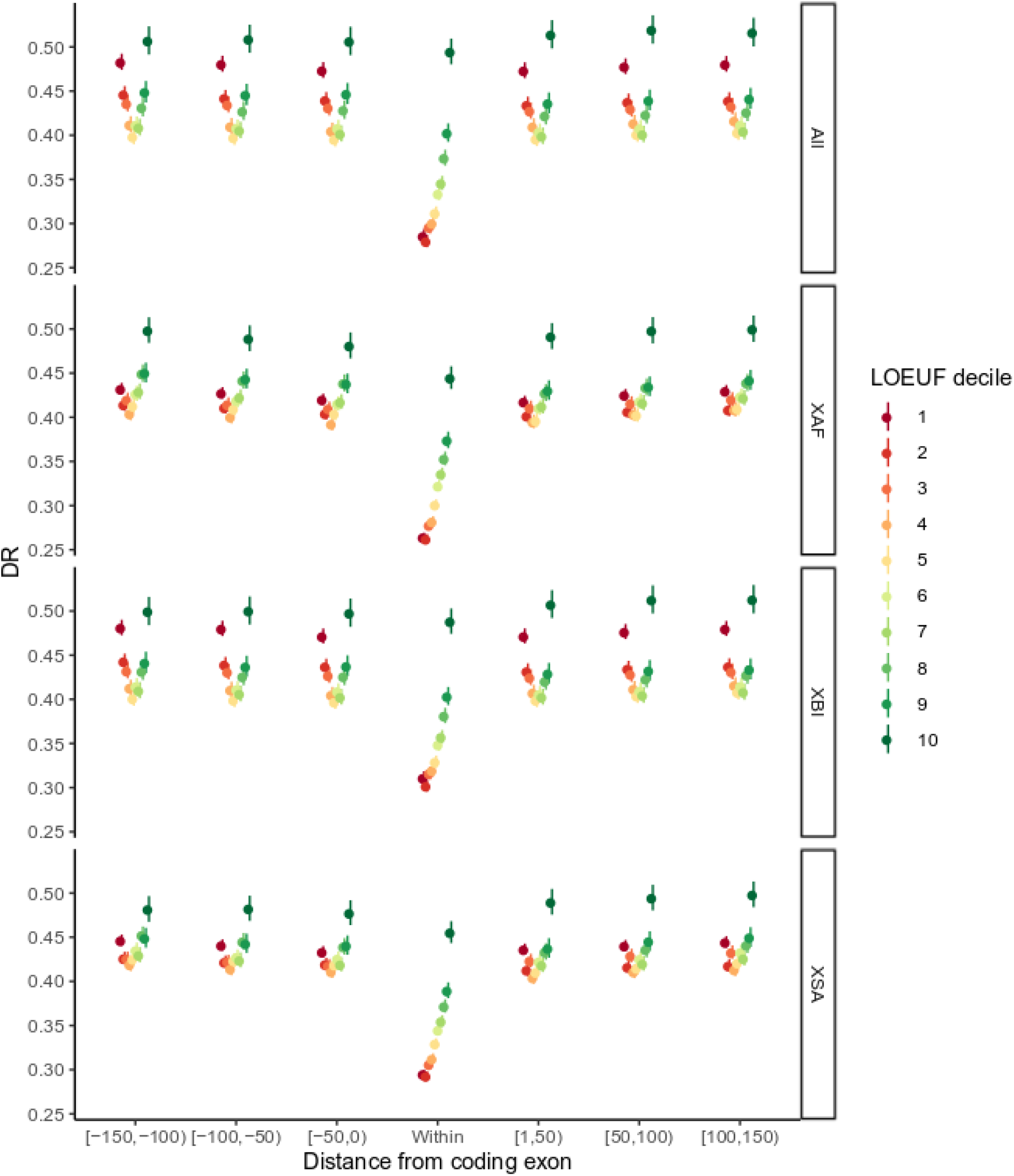
DR as a function of distance from coding exon partitioned by LOEUF^11^ deciles. Results are shown separately for the overall dataset (All) and the individual cohorts, XBI, XAF and XSA.

**Fig. S12.**
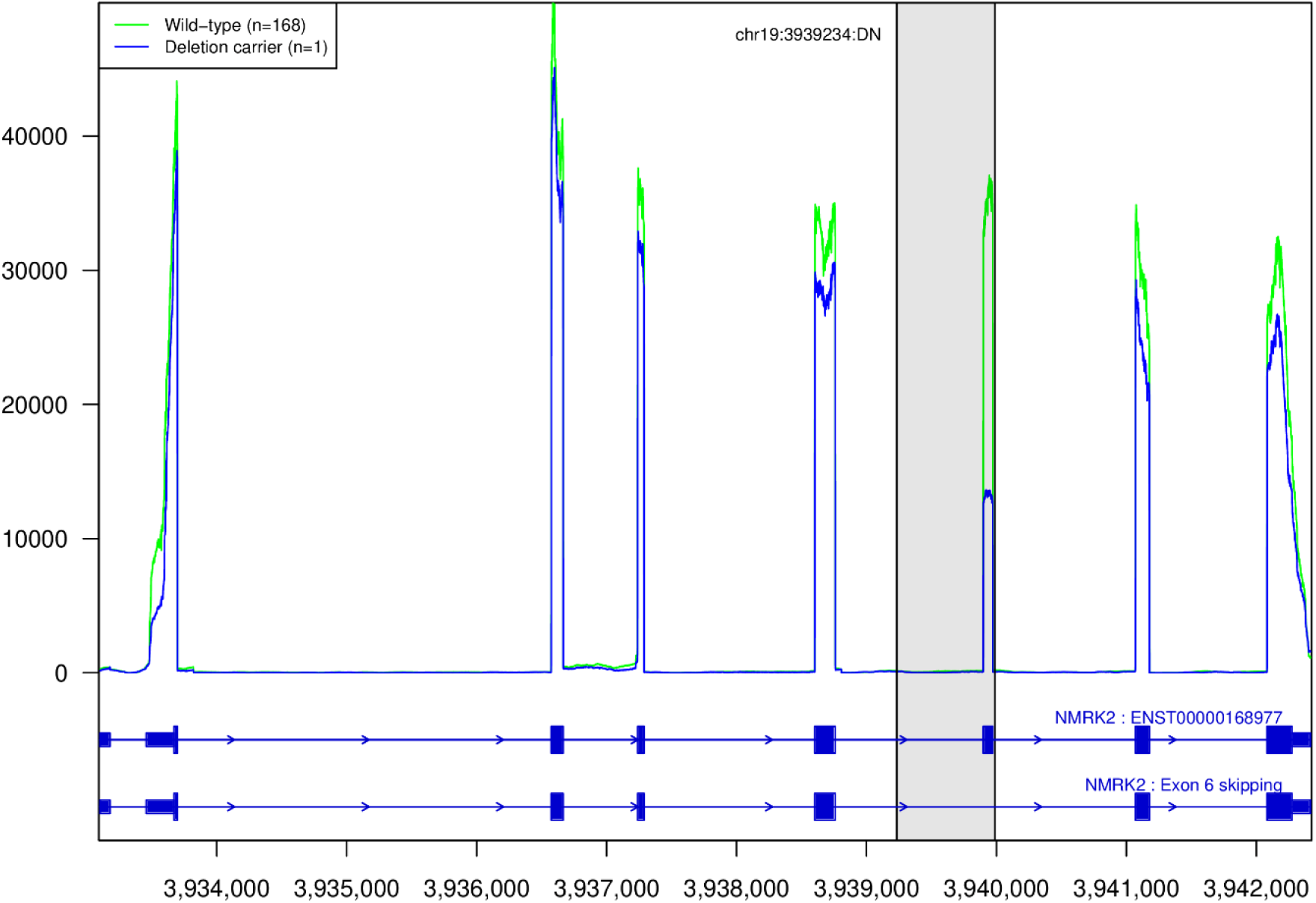
Coverage plot of RNA-sequenced reads from heart tissue from 169 heart tissue samples over the gene NMRK2. One individual is a carrier of a 754bp deletion depicted with gray rectangle that includes exon 6 of NMRK2. The RNA-coverage of the carrier (blue) is lower over exon 6 compared to median coverage of non-carriers (green). Shading marks the deleted region.

**Fig. S13.**
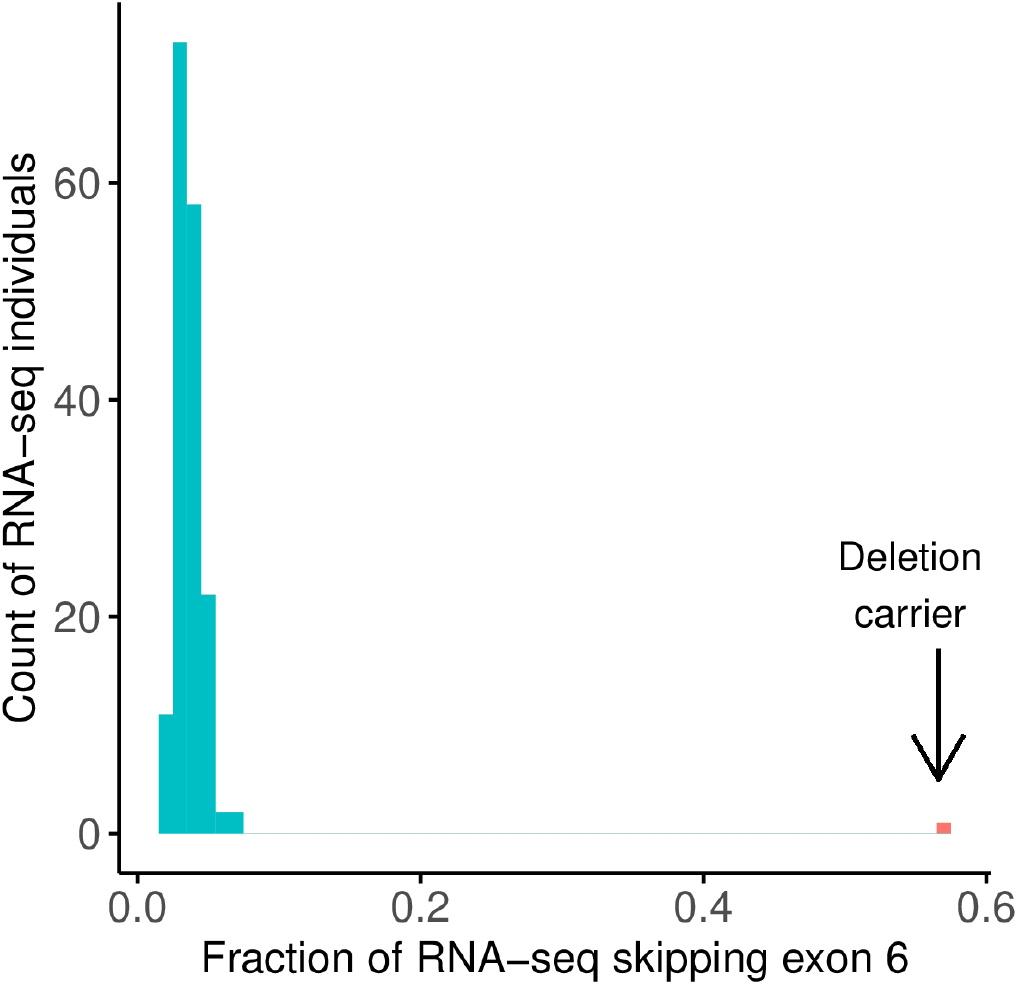
Histogram of fraction of RNA-sequenced fragments skipping exon 6 in NMRK2 out of all fragments aligning from the donor site of exon 5 to either acceptor site of exon 6 or exon 7. The median fraction fragments skipping for wild-type individuals is 0.035 and 0.57 for the carrier of the 754bp deletion.

**Fig. S14.**
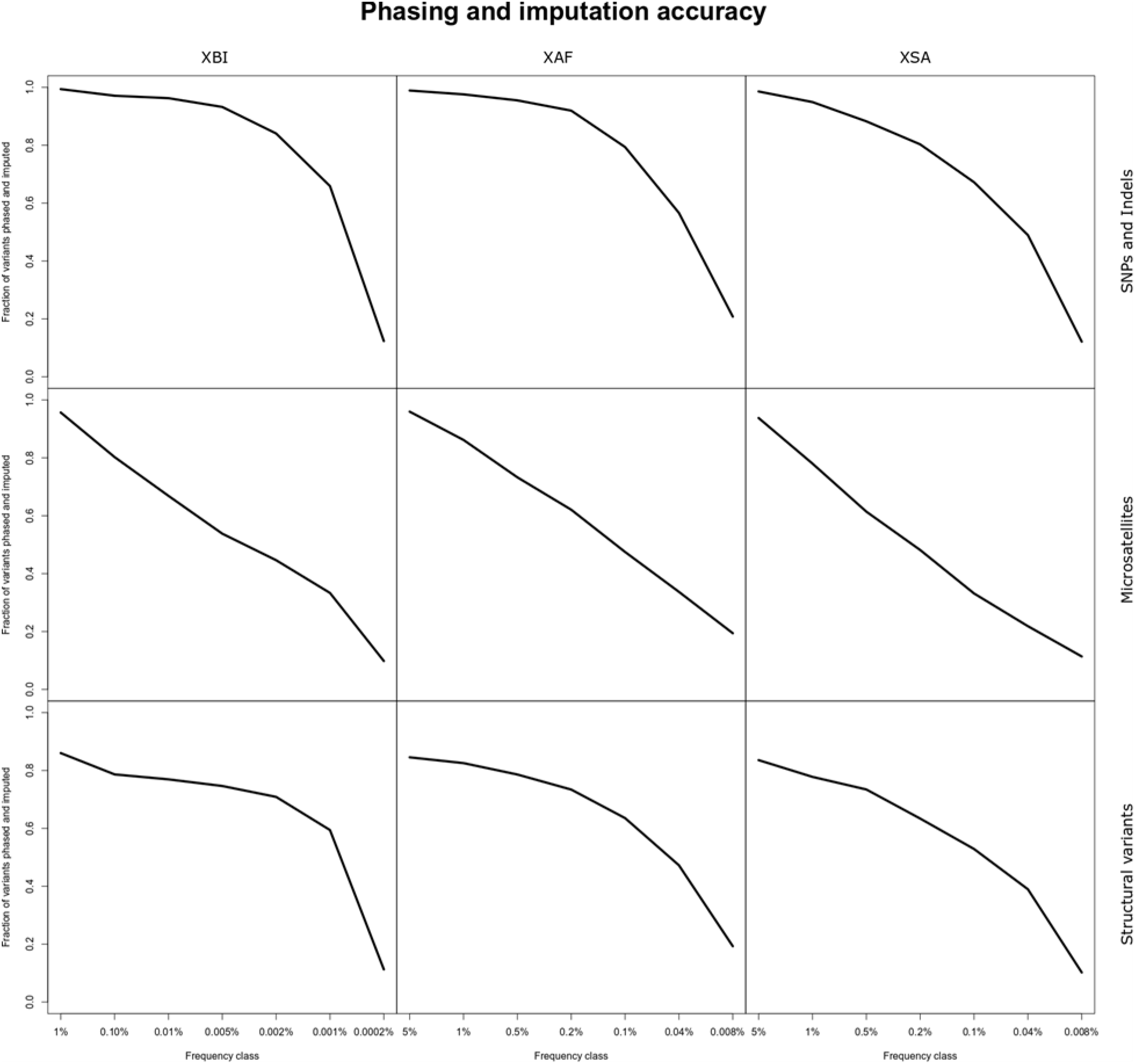
Imputation and phasing accuracy across variant datasets in the three populations. A variant is considered imputed if Leave one out r2 (L1or2) of phasing was greater than 0.5 and imputation information was greater than 0.8. x-axis splits variants into frequency classes based on the frequency in each cohort.

**Fig. S15.**
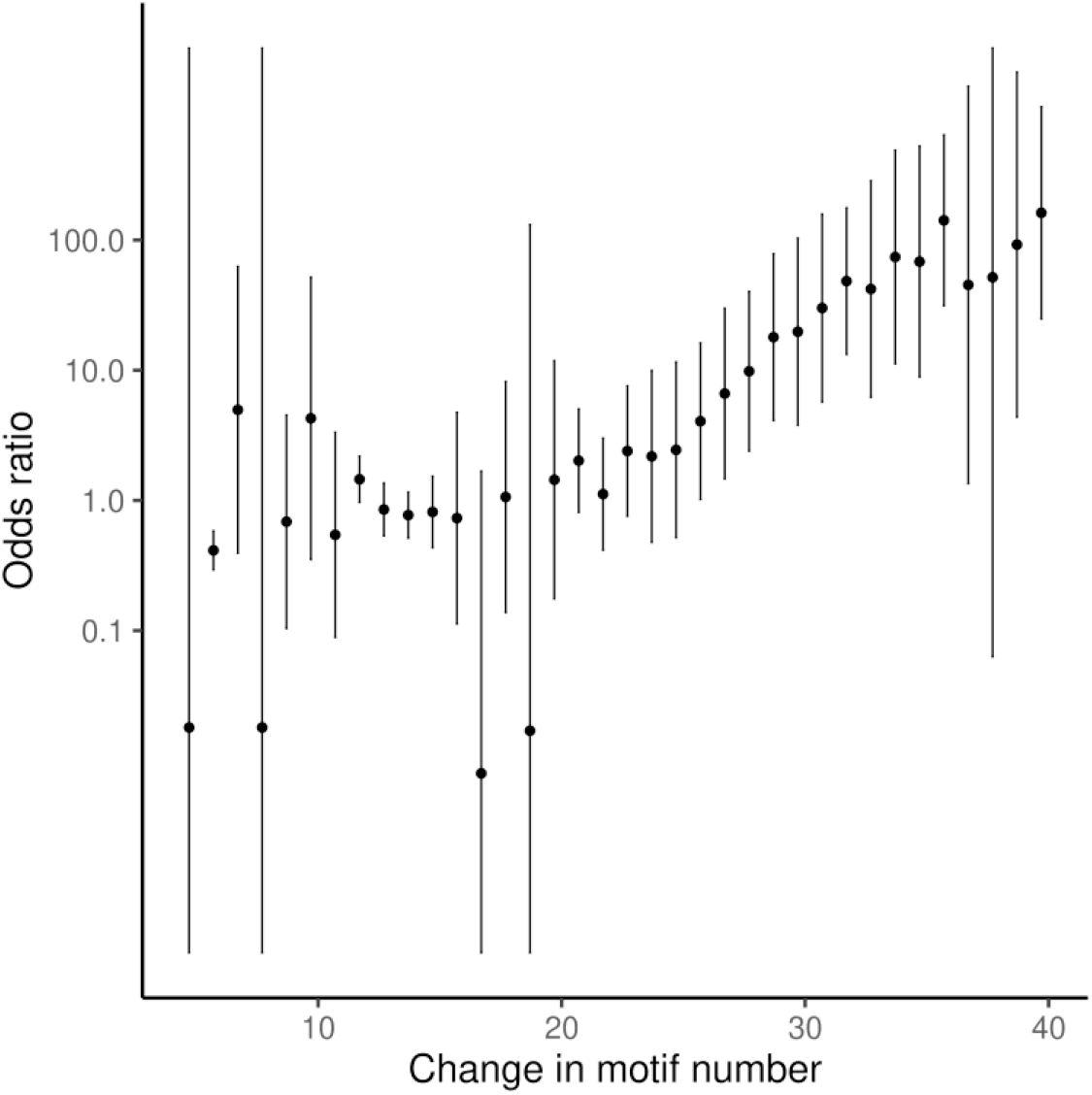
Odds ratio for risk of myotonic dystrophy as a function of repeat length in microsatellite at the 3’ untranslated region of DMPK. Carriers of at least 39.7 copies of the microsatellite repeat motif have a 162-fold increased risk of myotonic dystrophy.

**Fig. S16.**
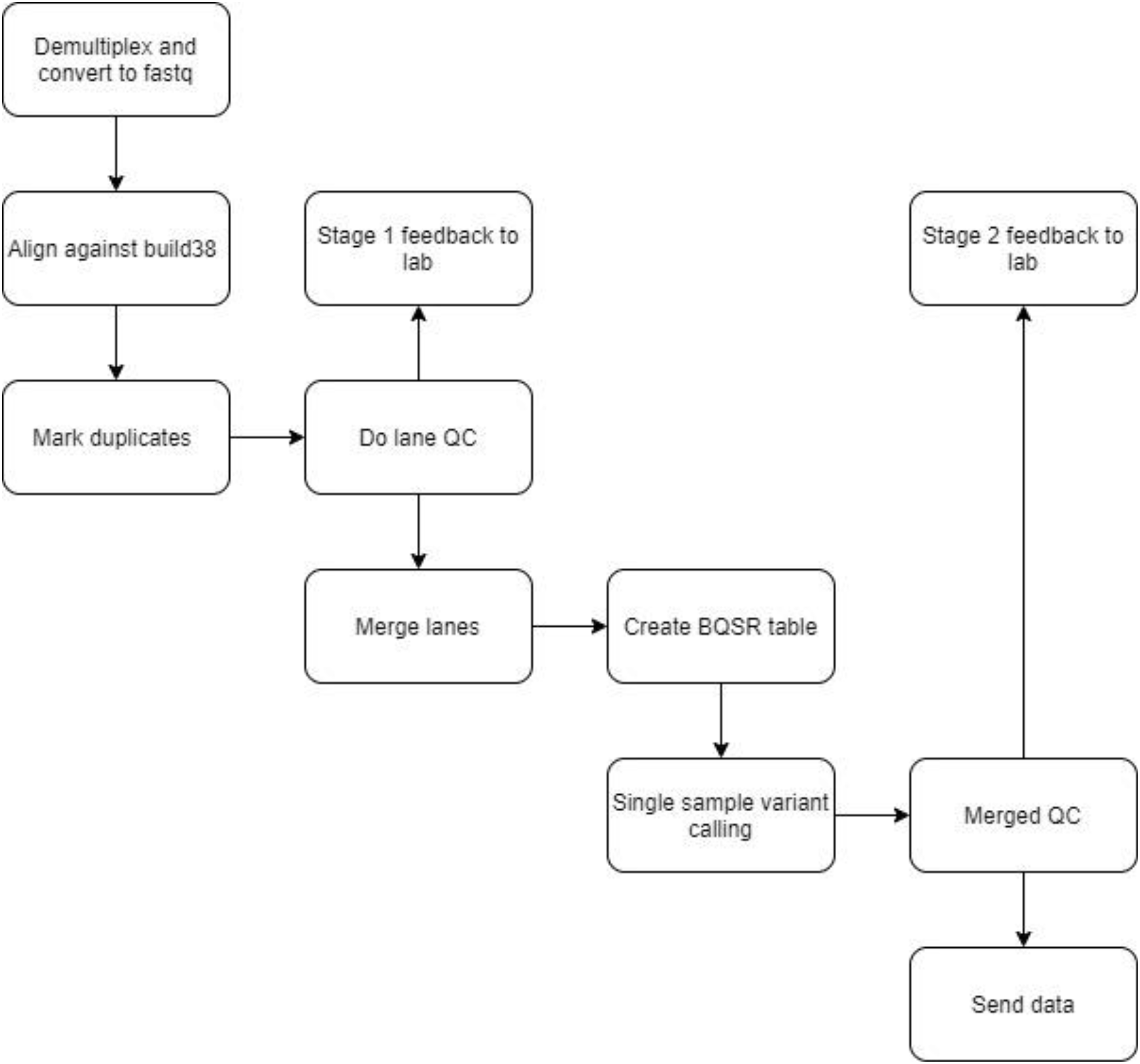
Process outline for UKB sequencing pipeline at deCODE genetics.

**Fig. S17.**
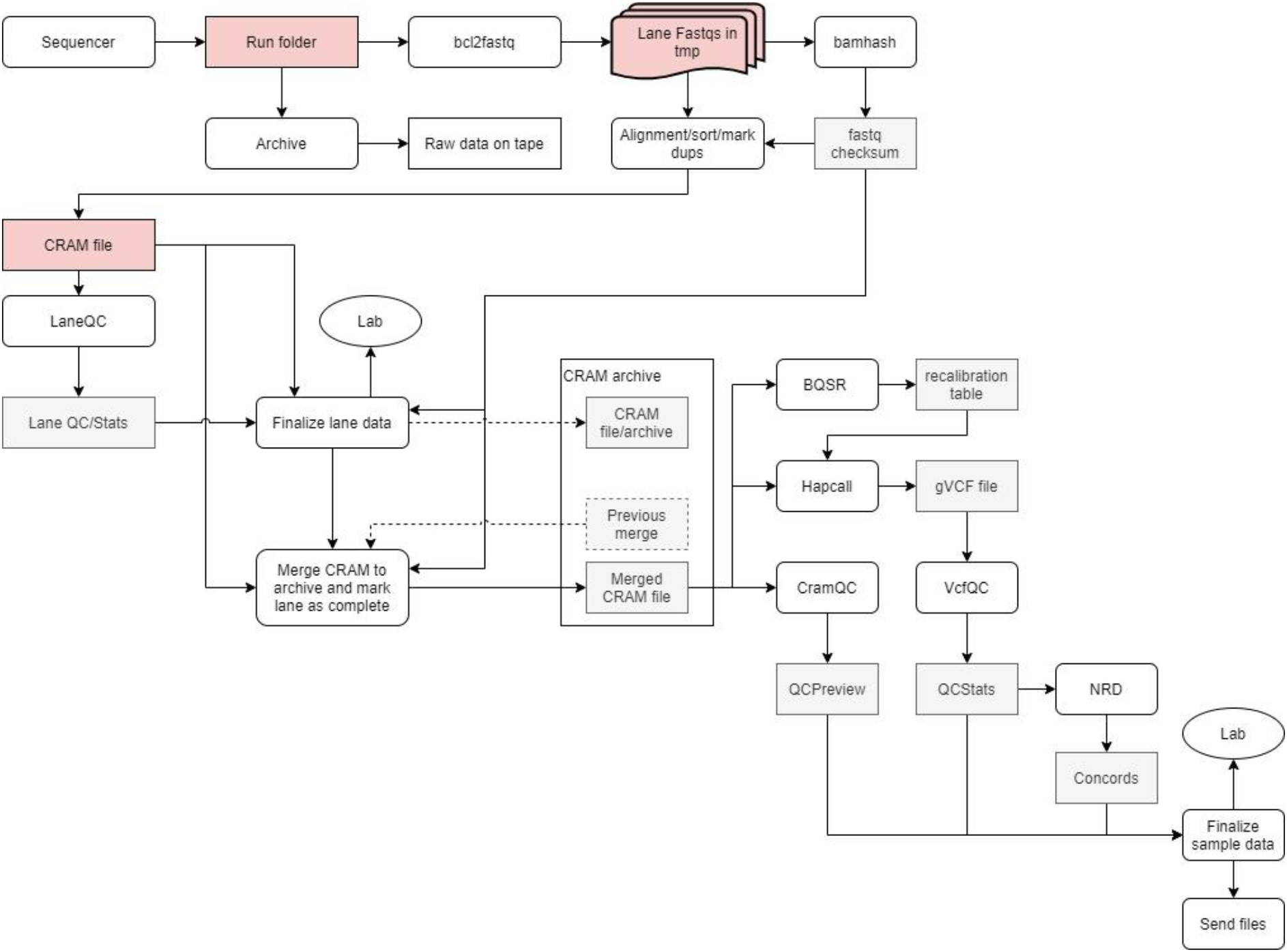
Pipeline for processing of sequence data at deCODE genetics.

**Fig. S18.**
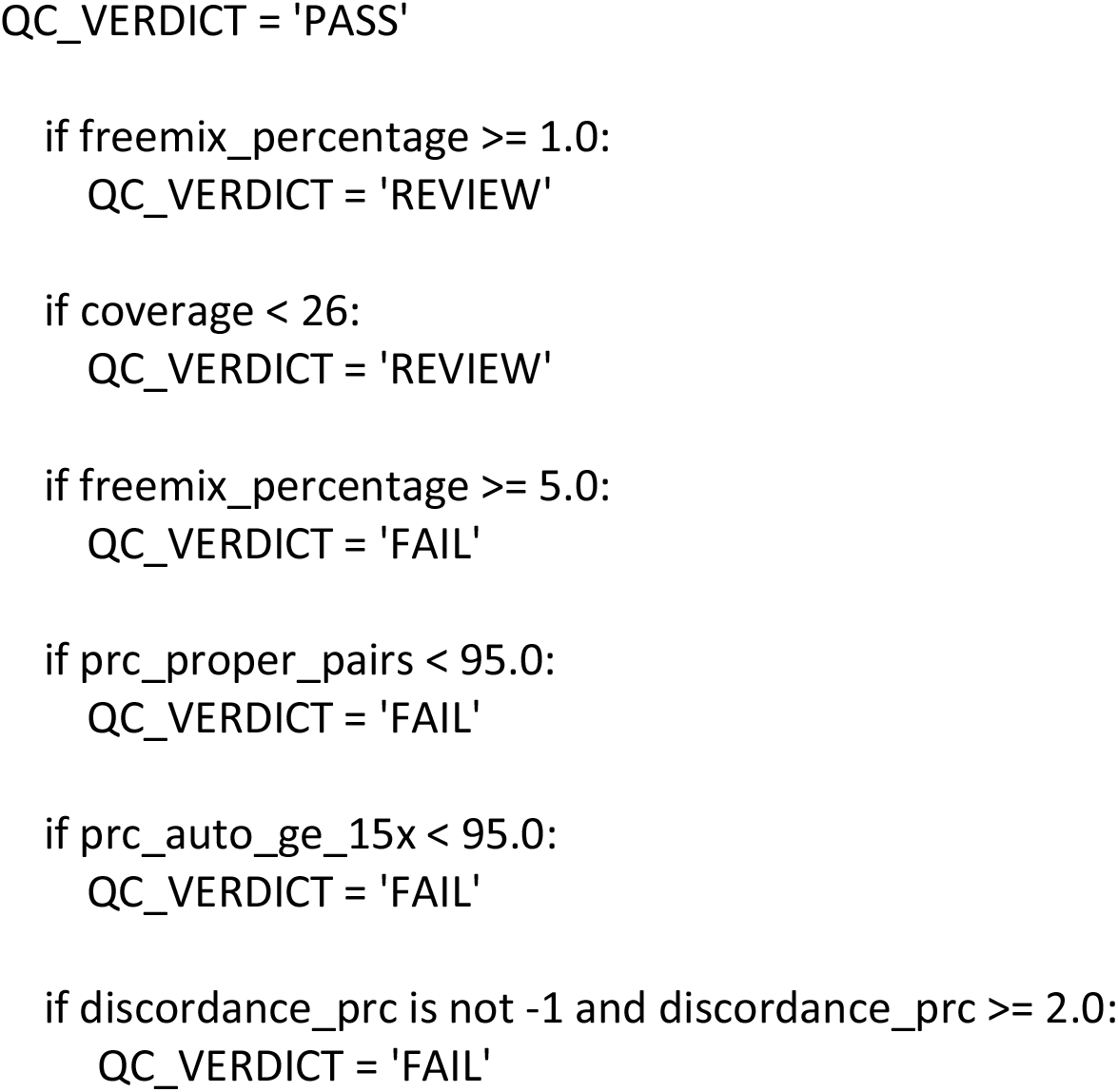
Logic used to compute PASS/FAIL for a WGS cram file.

**Fig. S19.**
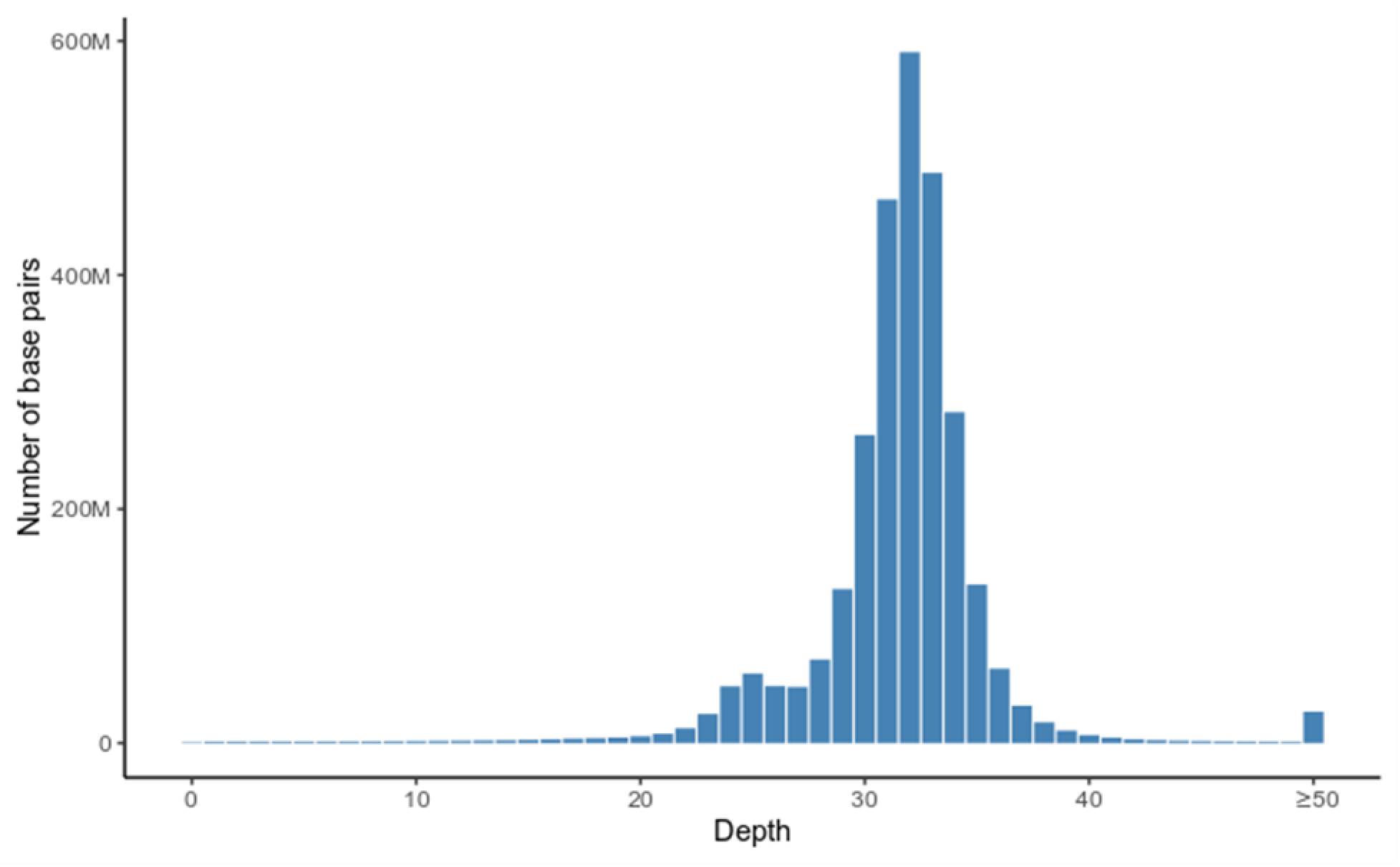
Average sequence coverage per base pair across the genome. The average coverage is computed from 1,000 randomly selected samples.

**Fig. S20.**
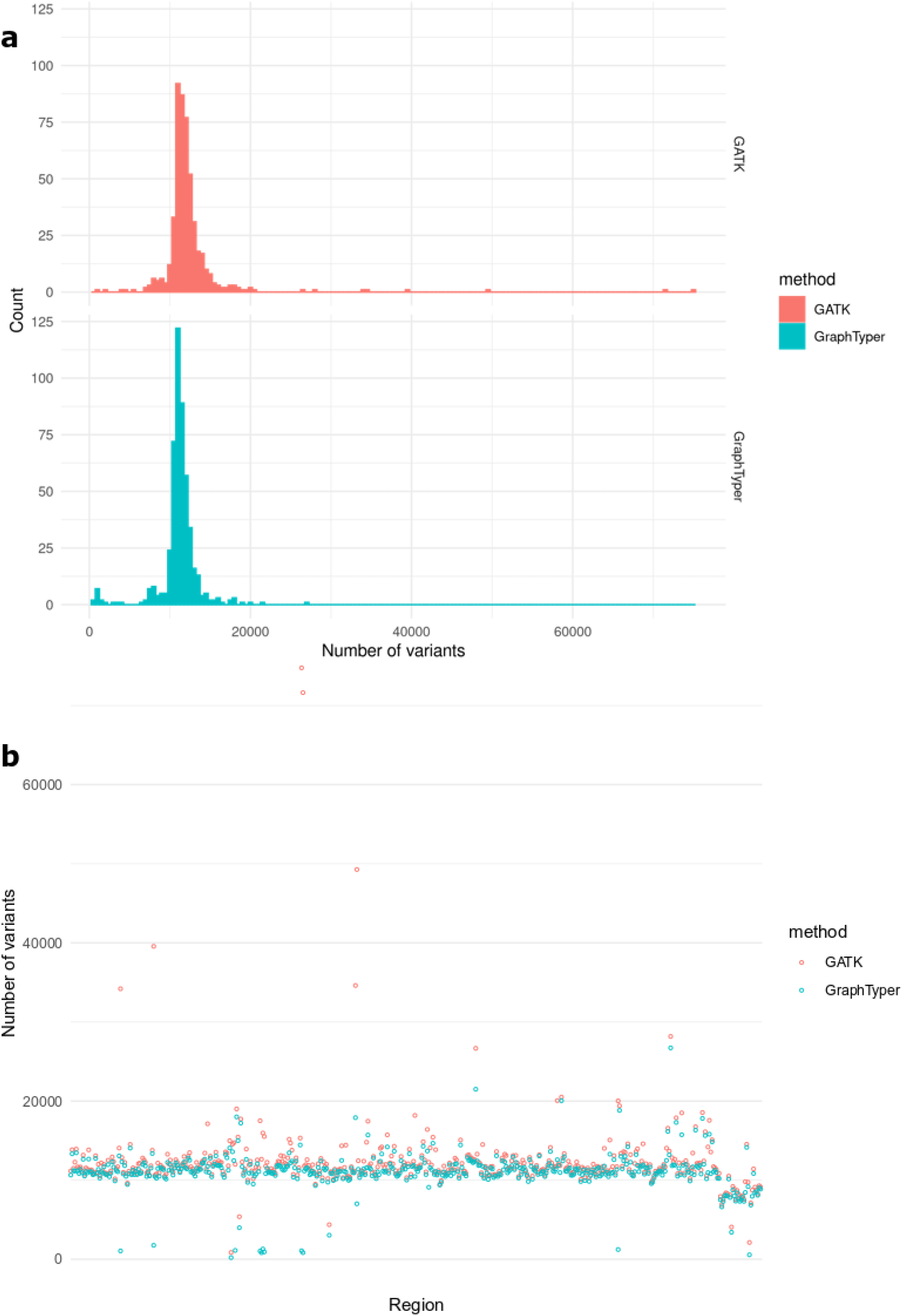
Number of variants per region in the 500 regions test set for the GATK and GraphTyper callsets, presented as a histogram a) and ordered by region b).

**Fig. S21.**
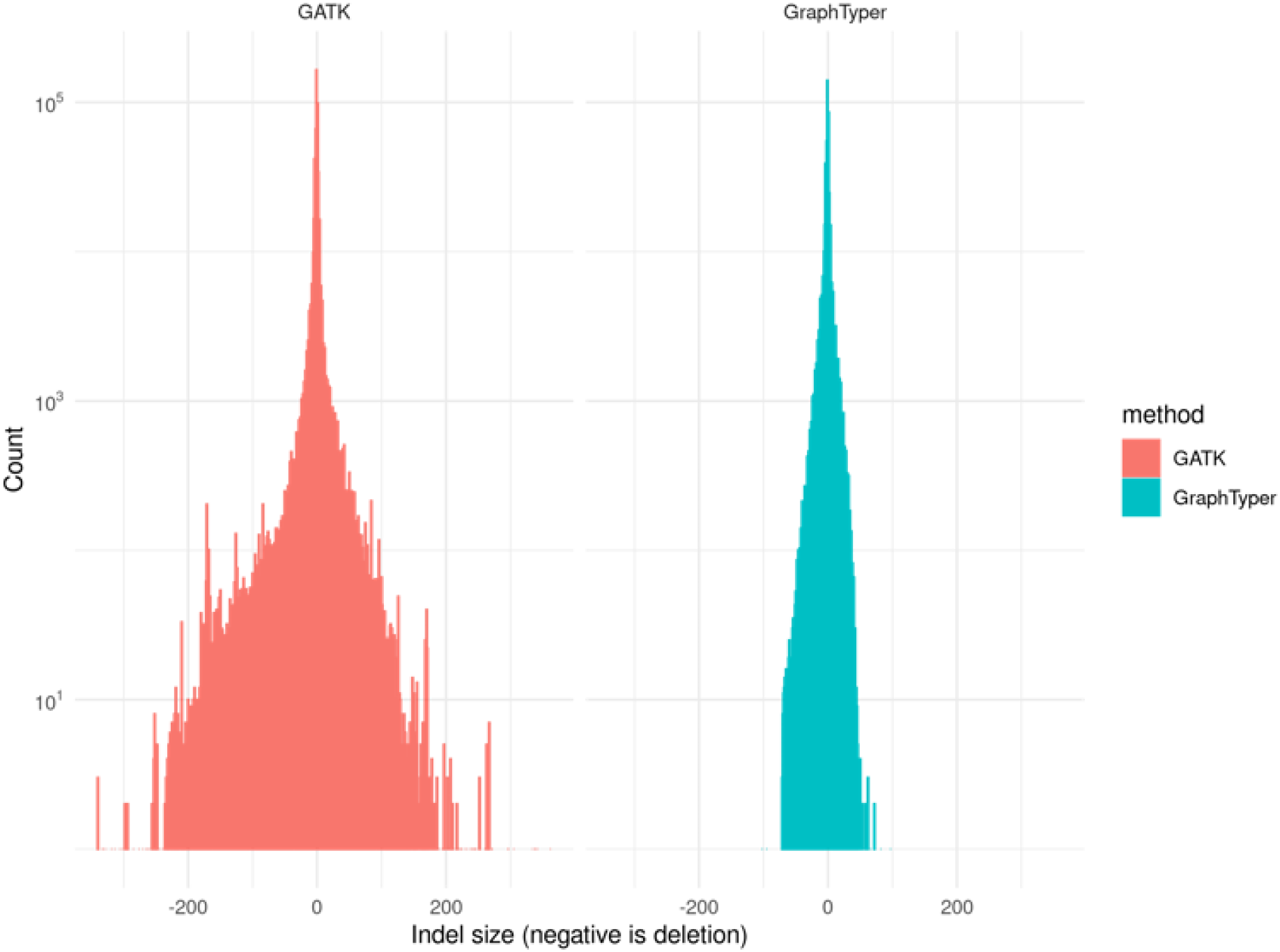
Distribution of indel sizes in GATK and GraphTyper callsets. Negative size indicates a deletion.

**Fig. S22.**
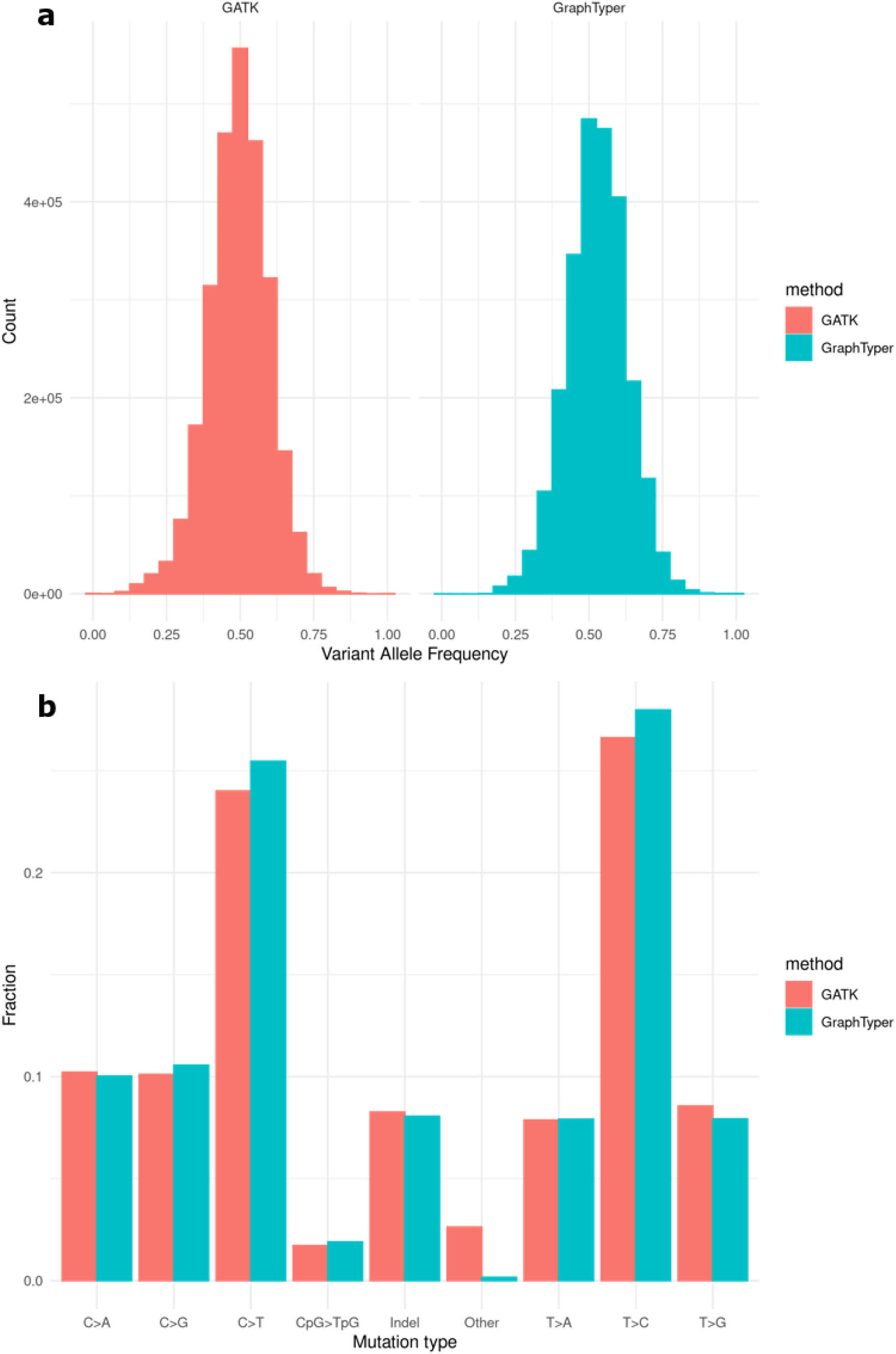
a) Variant allele frequencies (VAF) of singletons. b) Mutation classes of singletons. Results are for the GATK and GraphTyper callsets on 500 randomly selected regions.

**Fig. S23.**
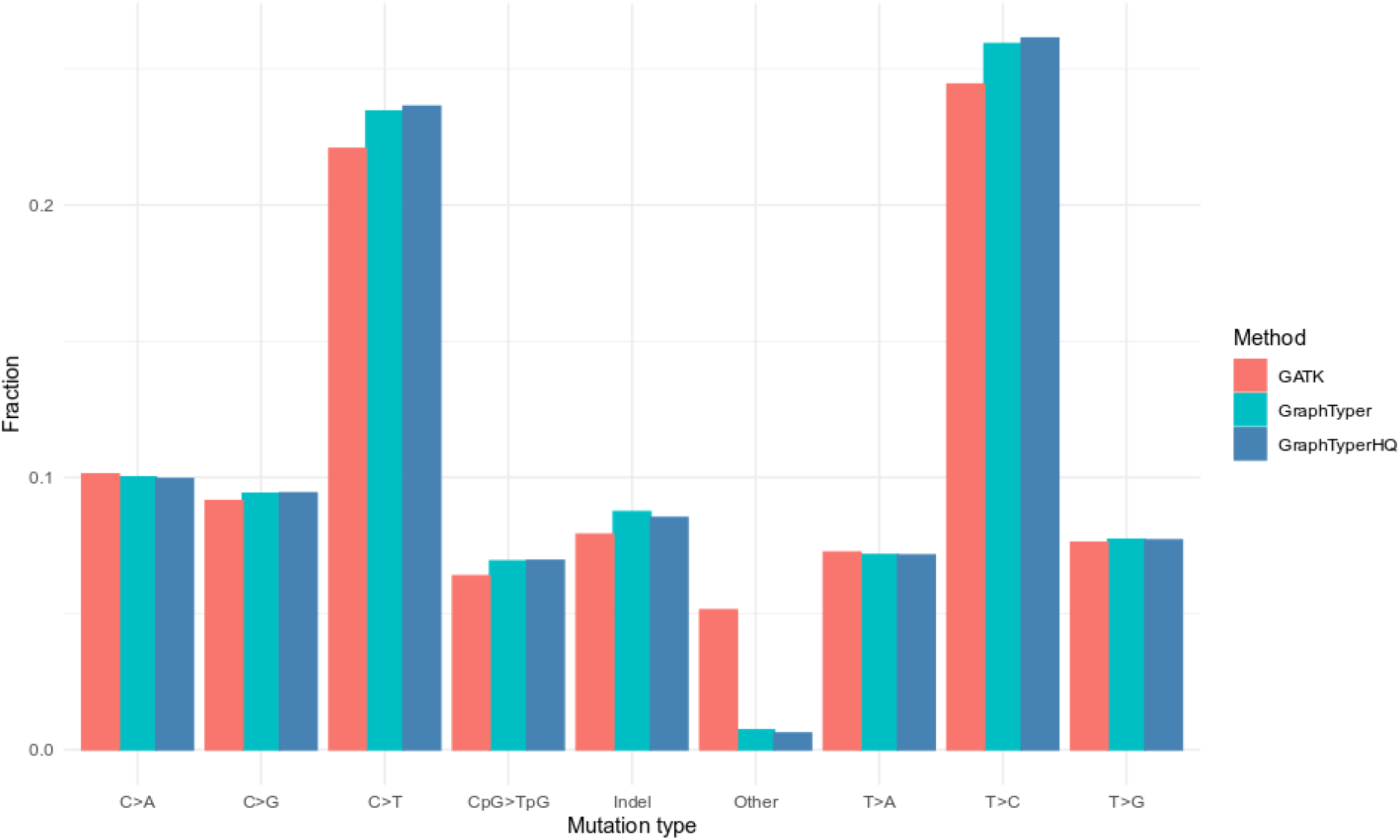
Fraction of variants by mutation type in the GATK, GraphTyper and GraphTyper HQ sets.

**Fig. S24.**
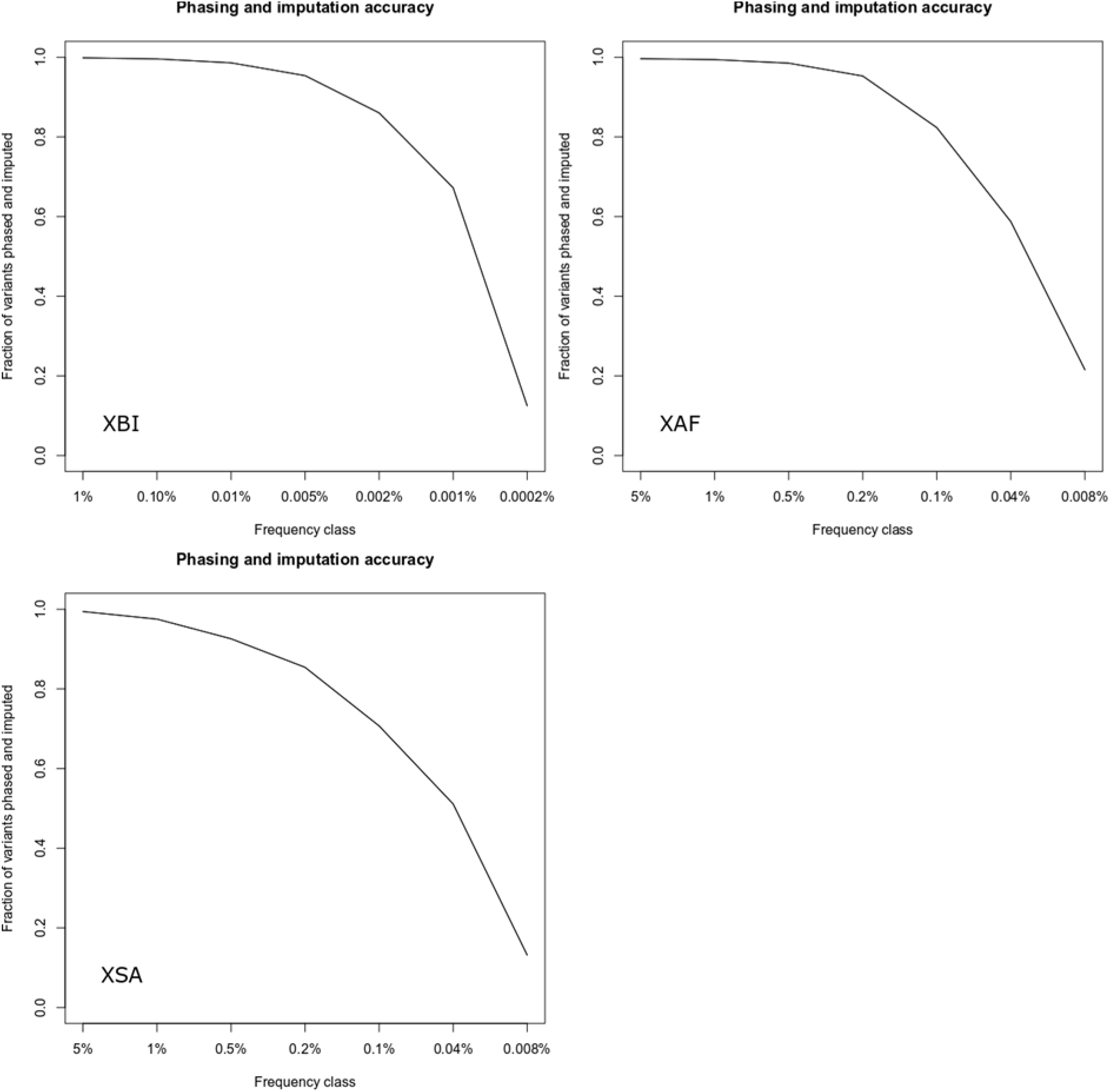
Imputation accuracy for variants with AAscore > 0.9 in the three populations, Top left: XBI, Top Right: XAF, Bottom: XSA. A variant was considered imputed if Leave one out r2 of phasing was greater than 0.5 and imputation information was greater than 0.8. x-axis splits variants into frequency classes based on the number of carriers in the sequence dataset, with the number representing the minimum number of carriers in the frequency class. Variants are split by variant type.

**Fig. S25.**
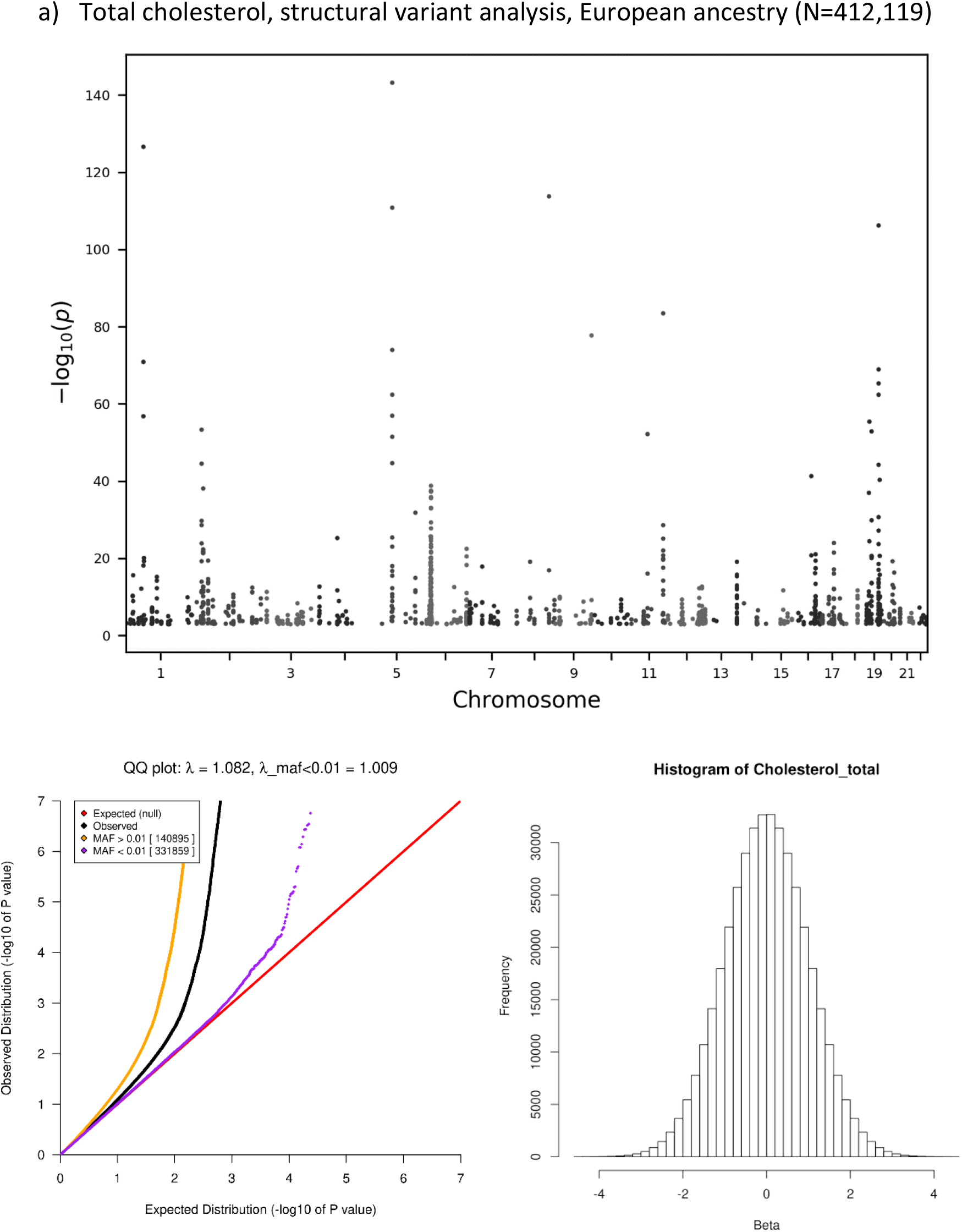
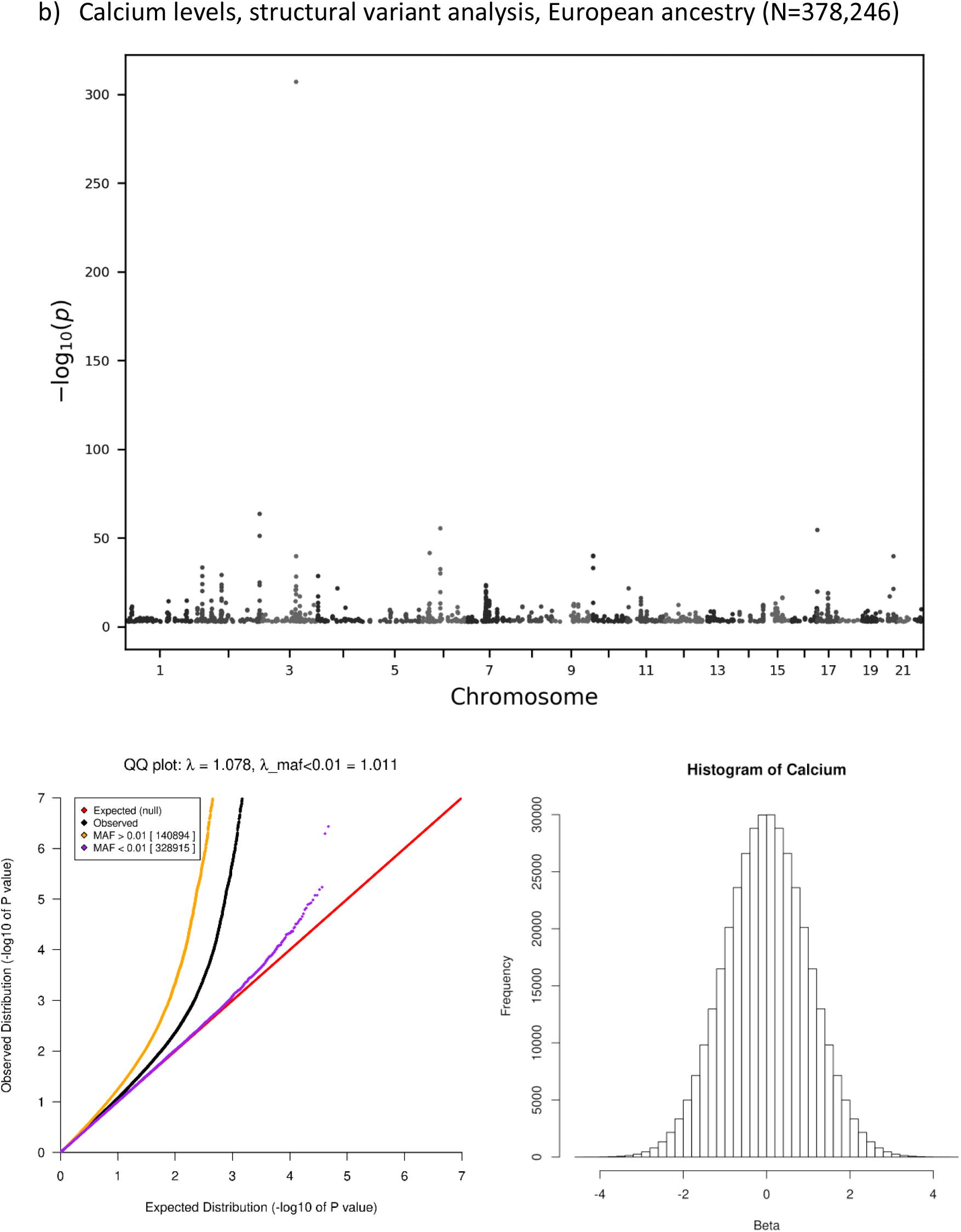
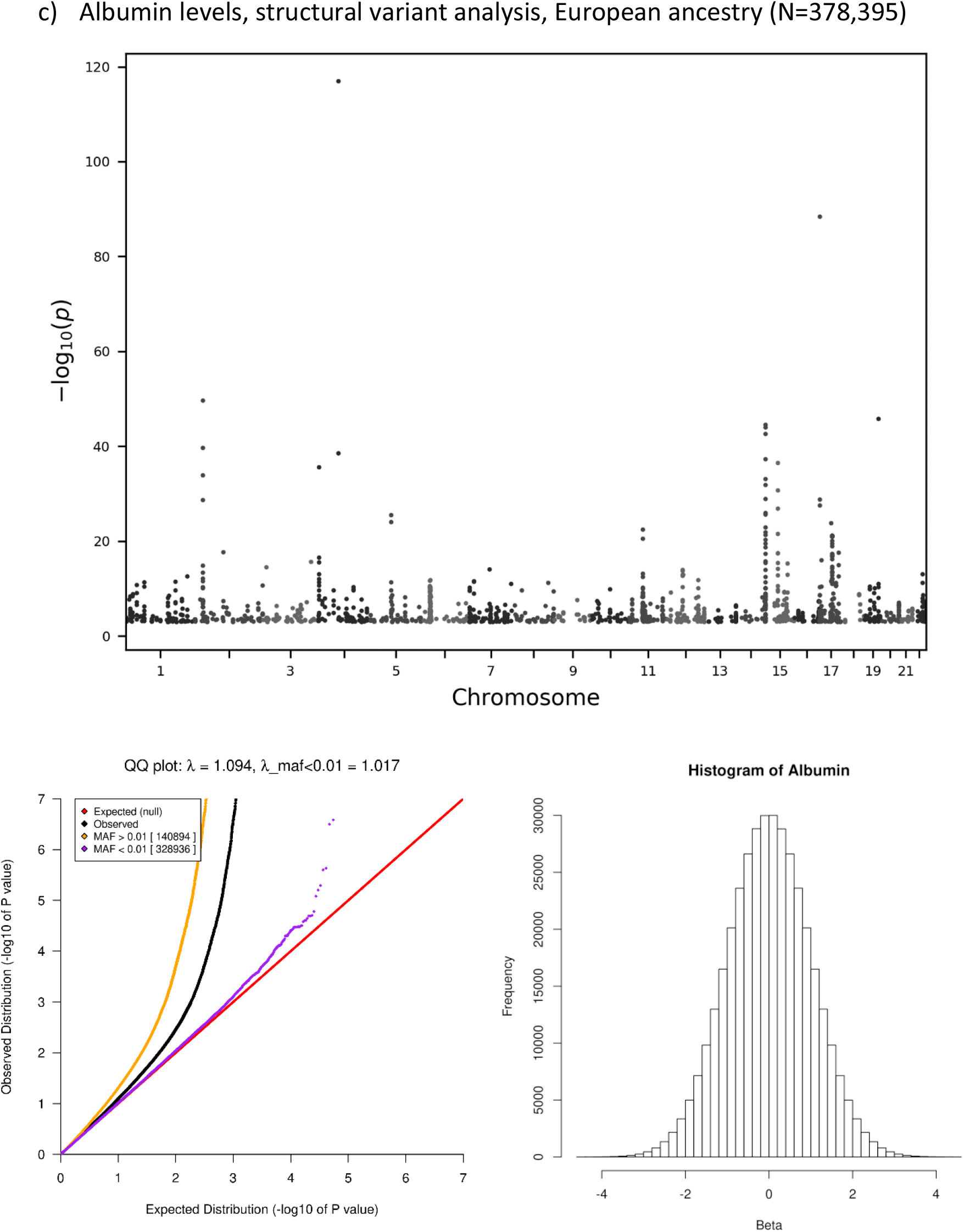
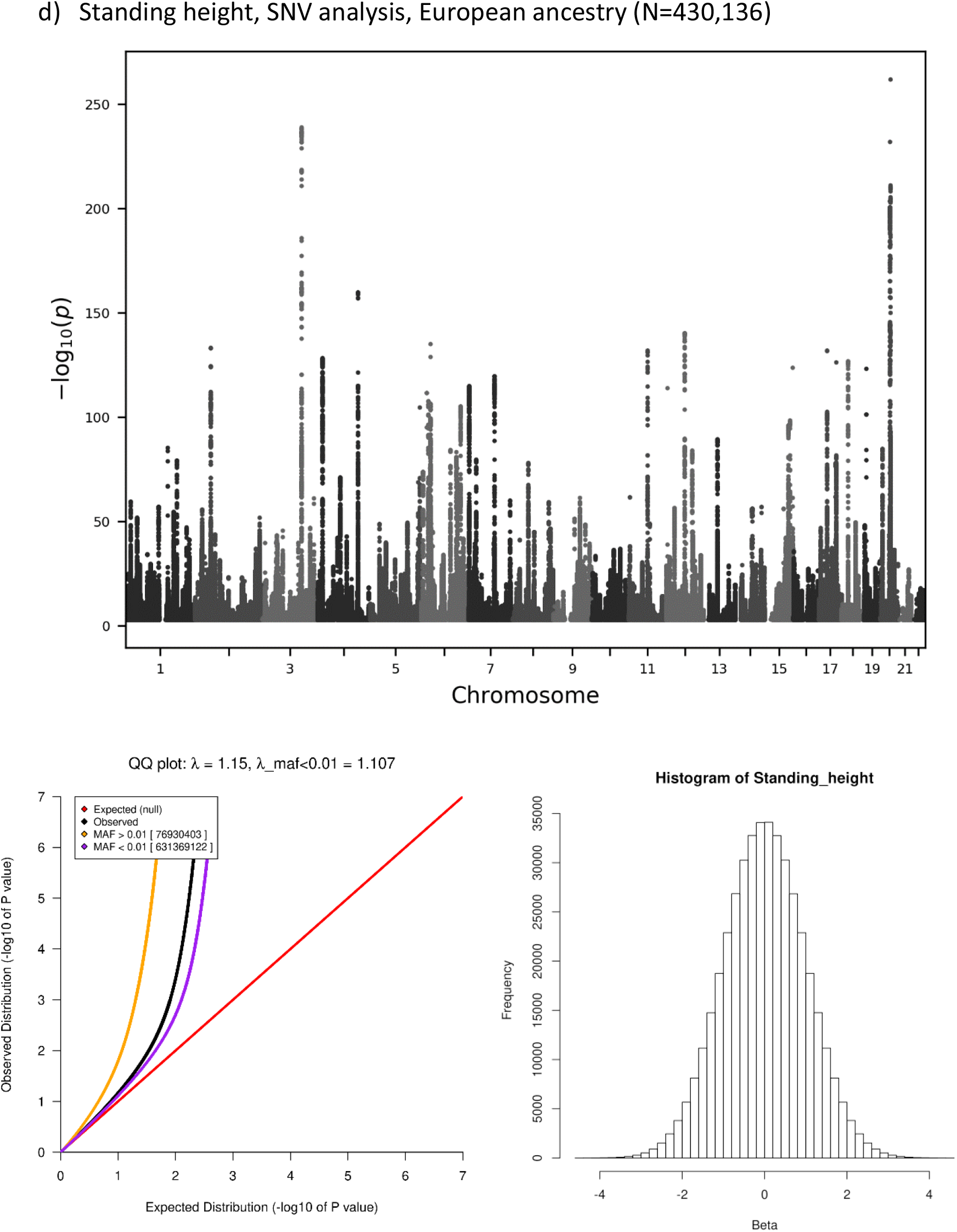
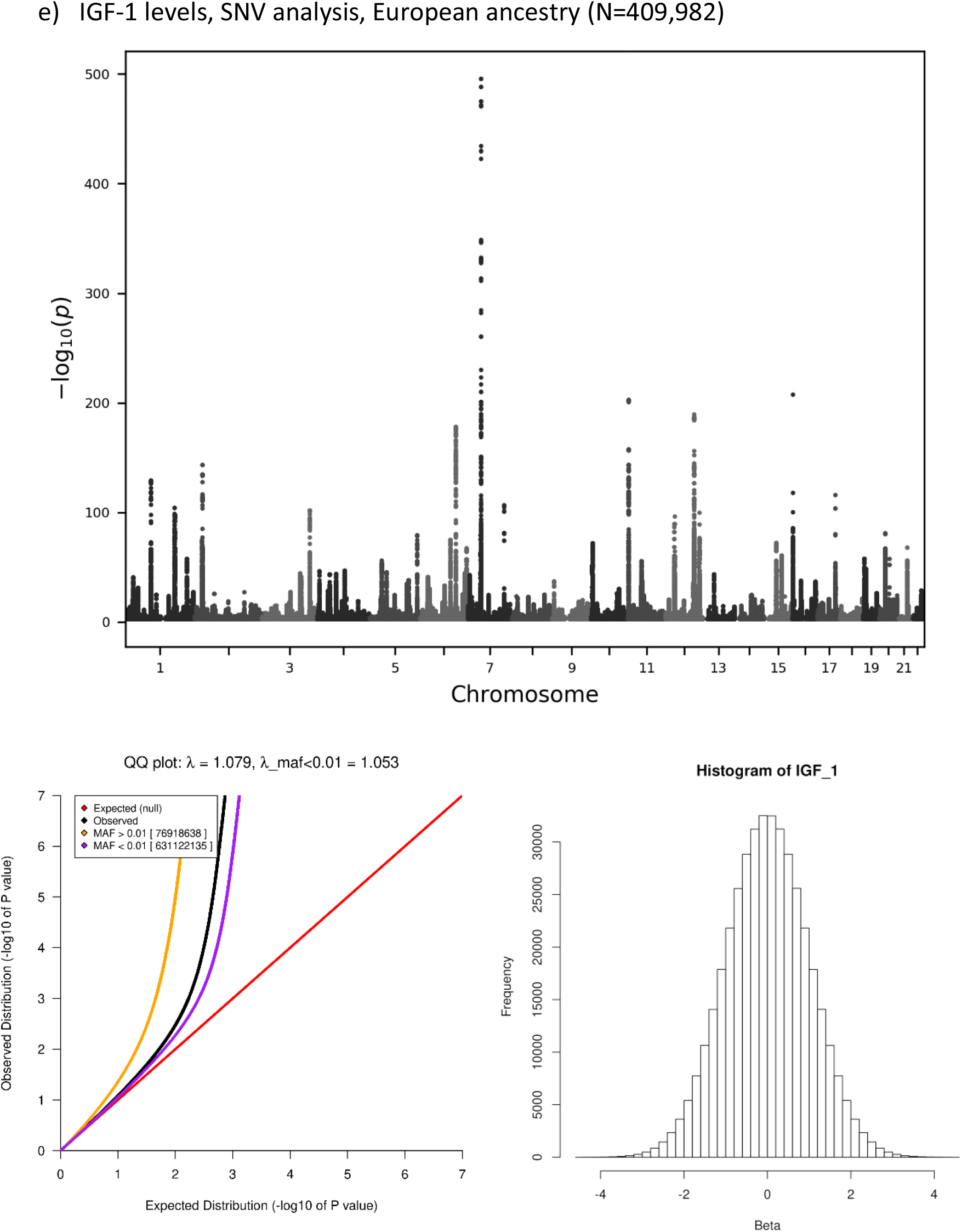
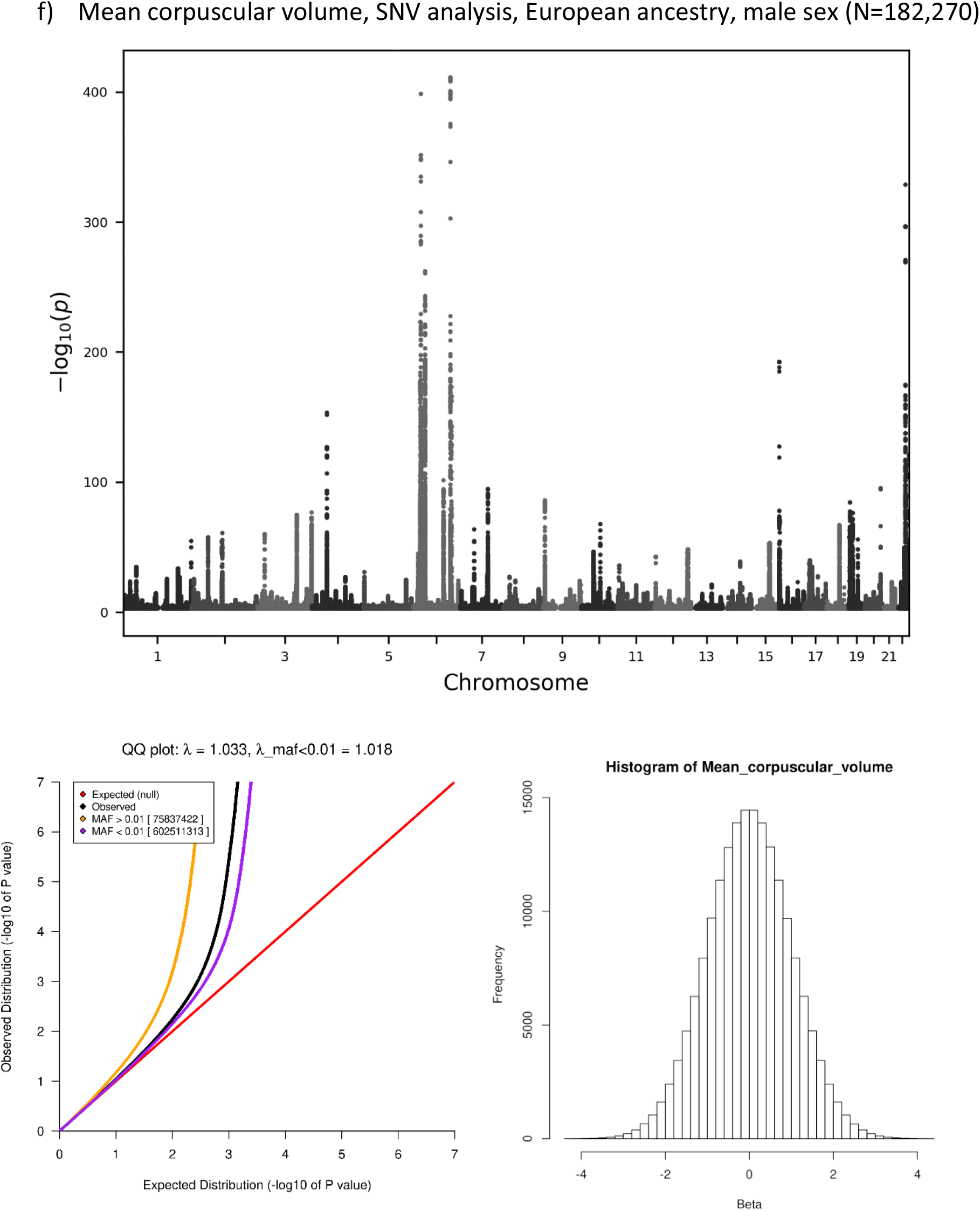
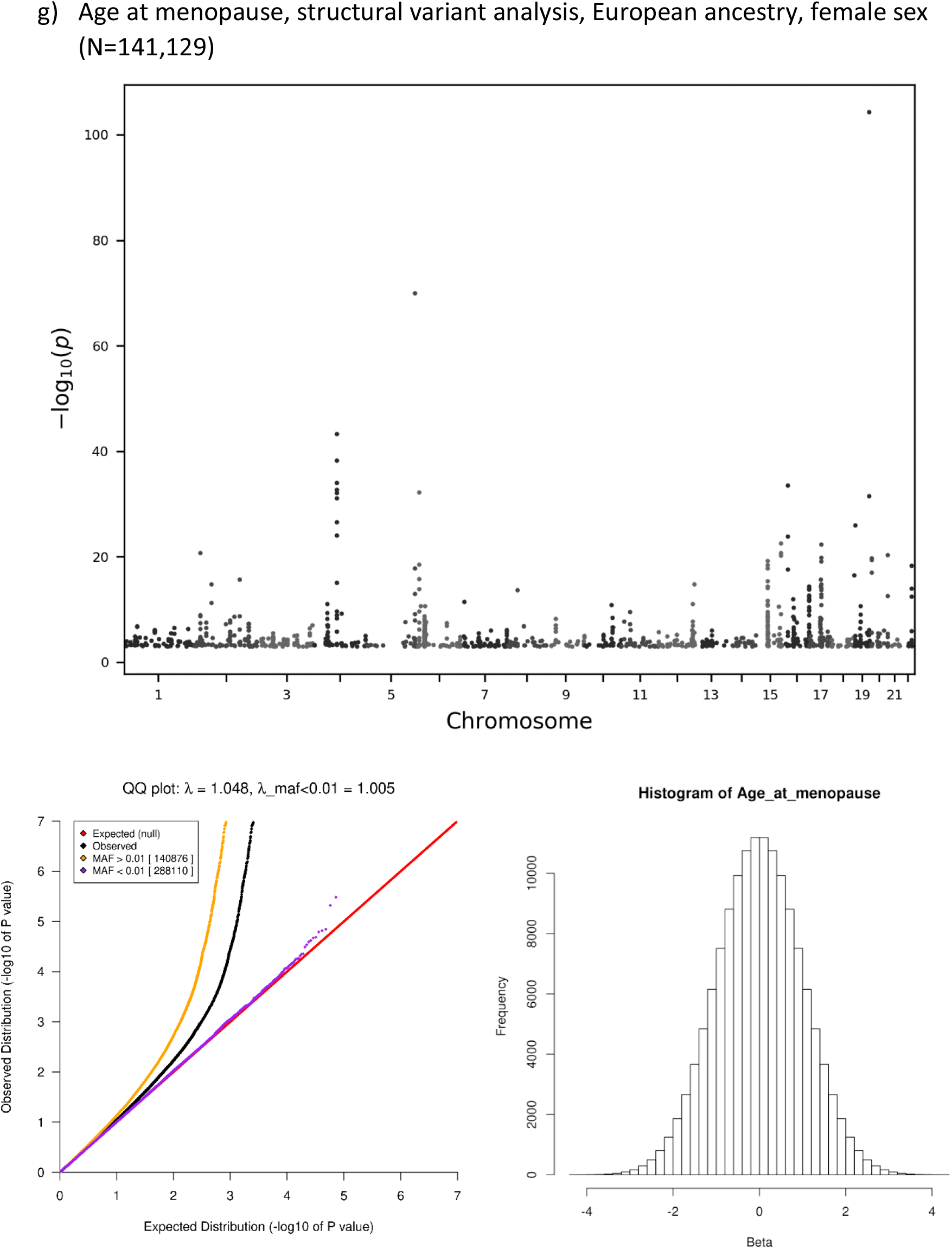
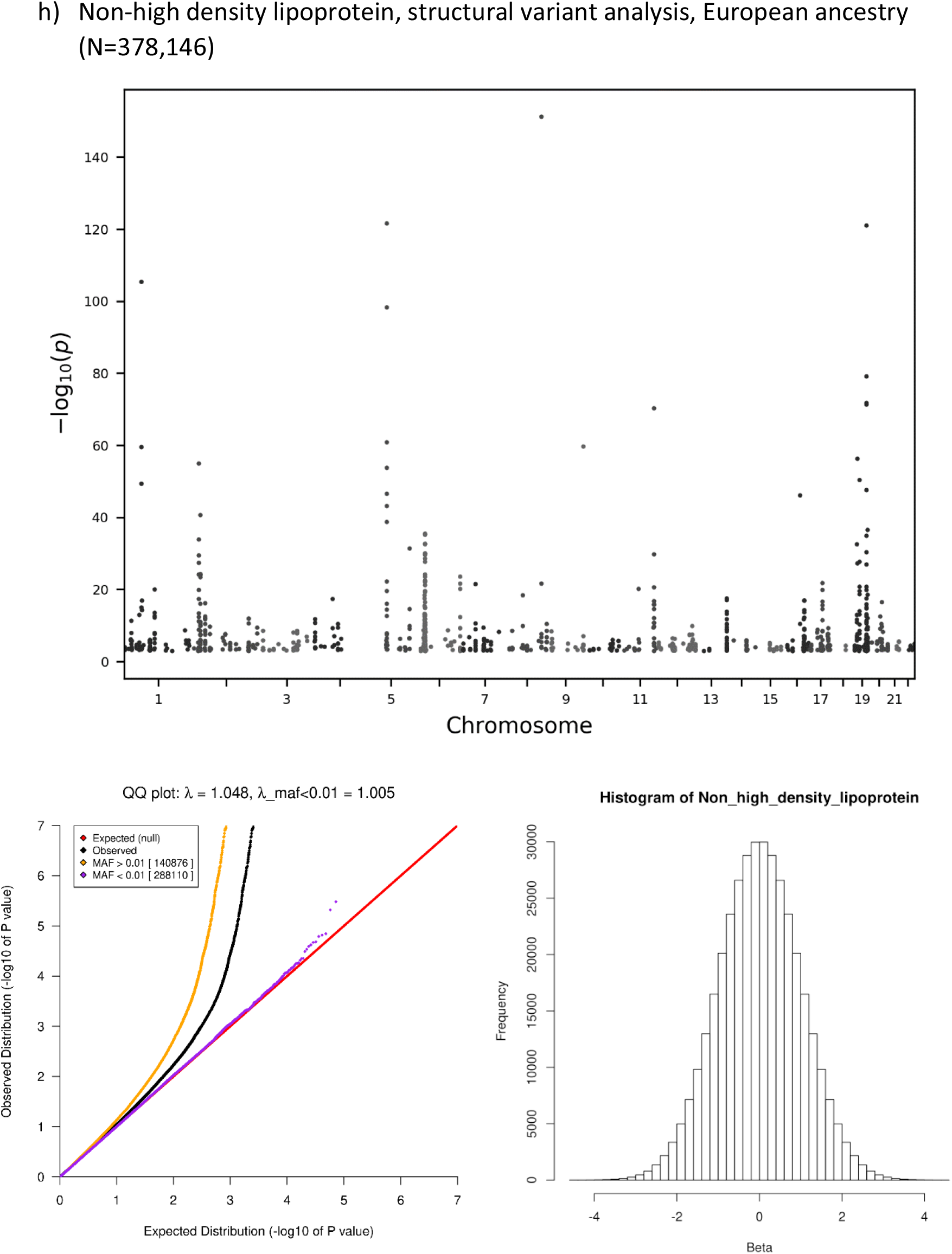
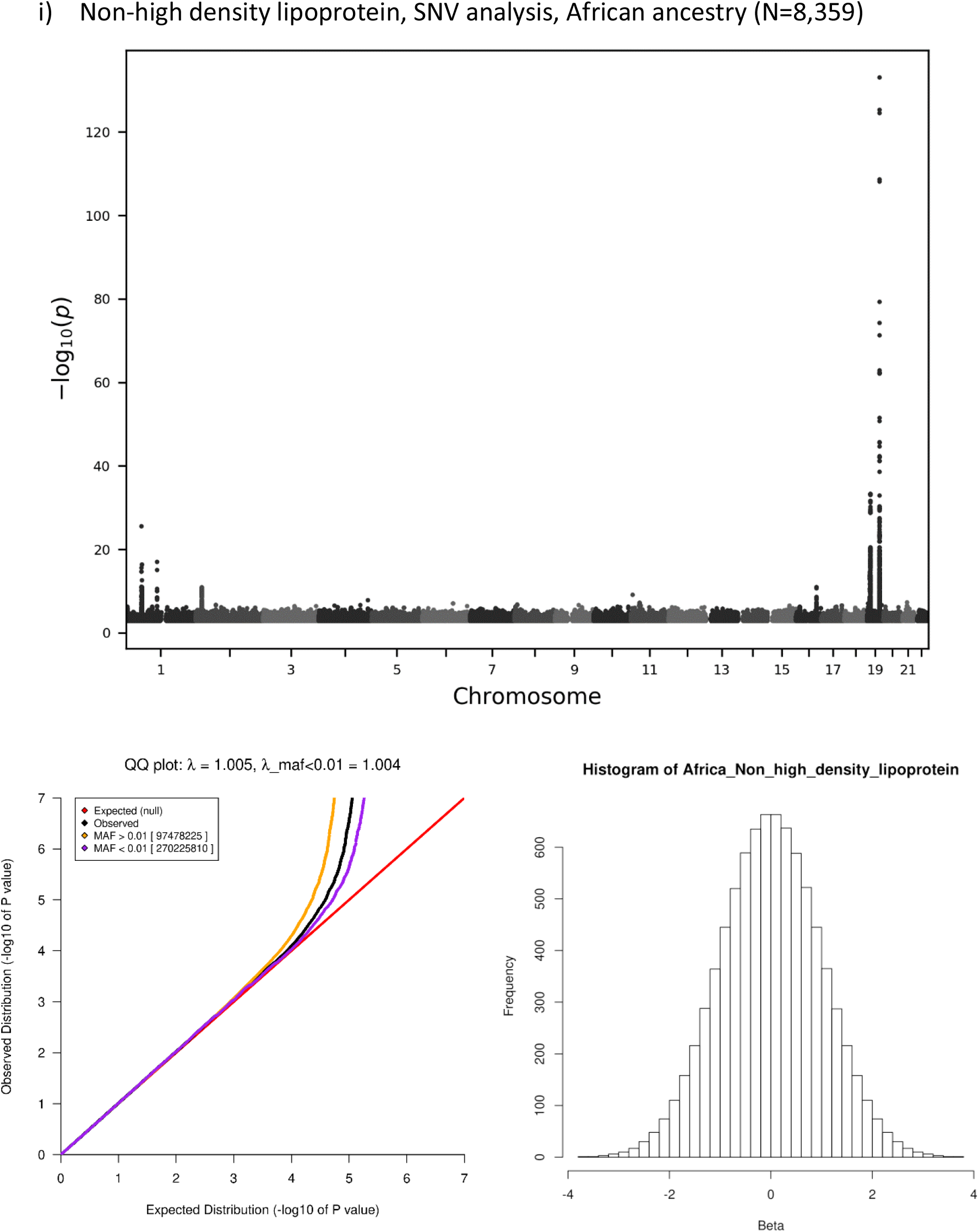
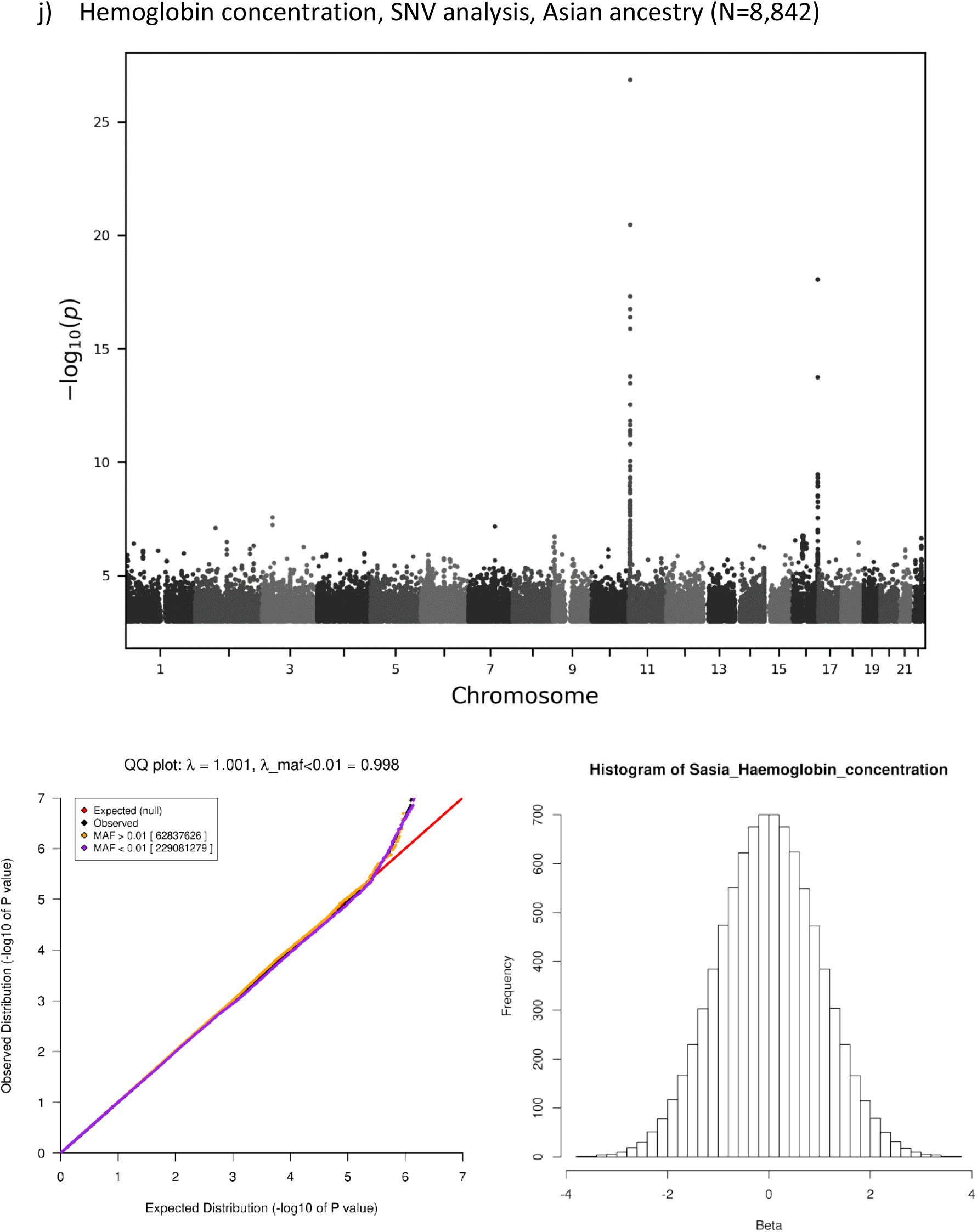
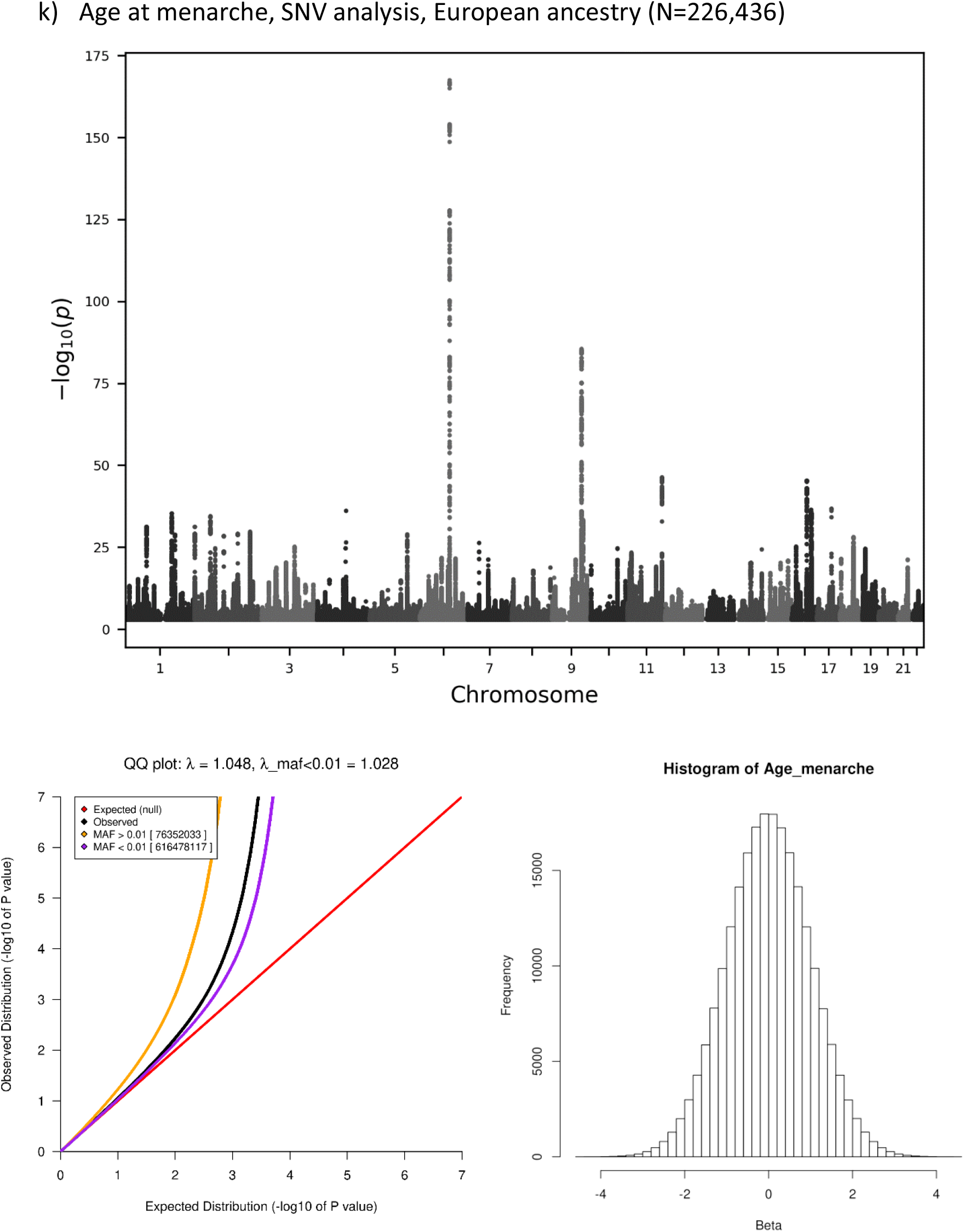
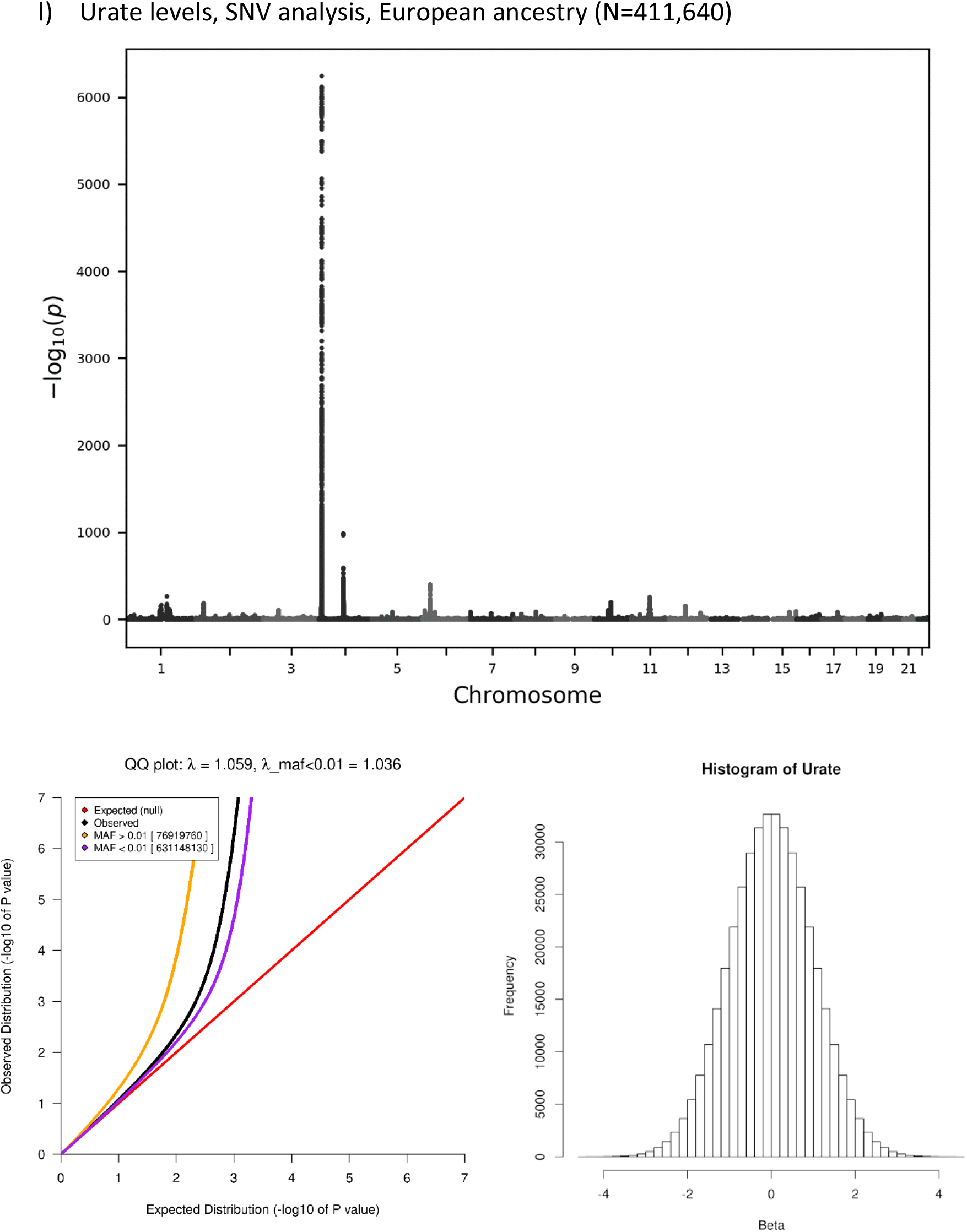
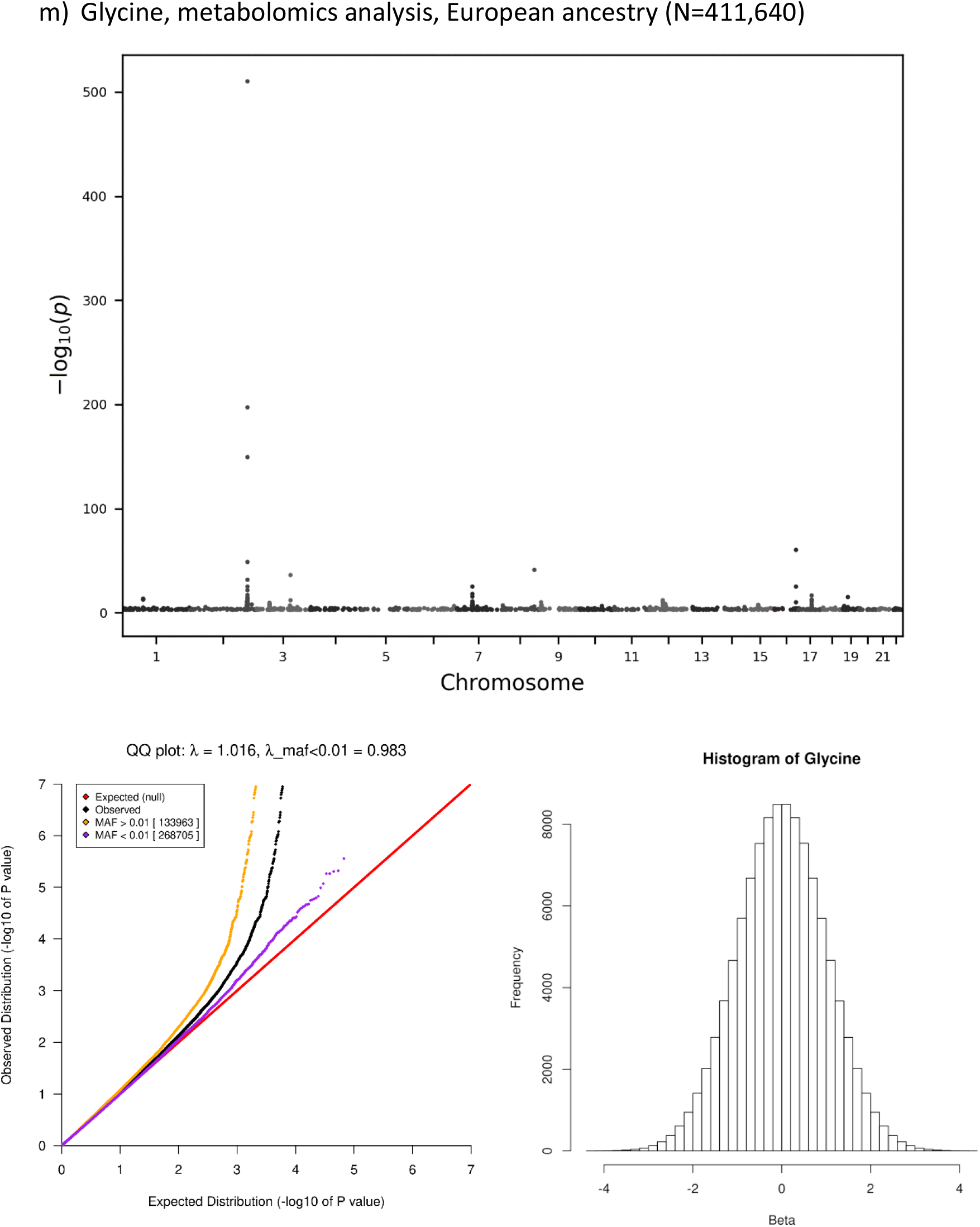
Manhattan plots, quantile-quantile (QQ) plots and histograms of inverse-normal transformed values after adjustment for covariates age, sex and 40 principal components, when applicable, for quantitative traits with significant results reported in this manuscript. For Manhattan plots, the x-axis represents chromosome locations and the y-axis shows the –log10 significance levels of the associations. For QQ plots, the inflation (λ) is shown in the title of each graph, for all variants and for rare variants only (λ_maf<0.01). For the histograms, the x-axis shows the value range of the inverse-normal transformed points and the y-axis shows the count of individuals within value ranges.

**Fig. S26.**
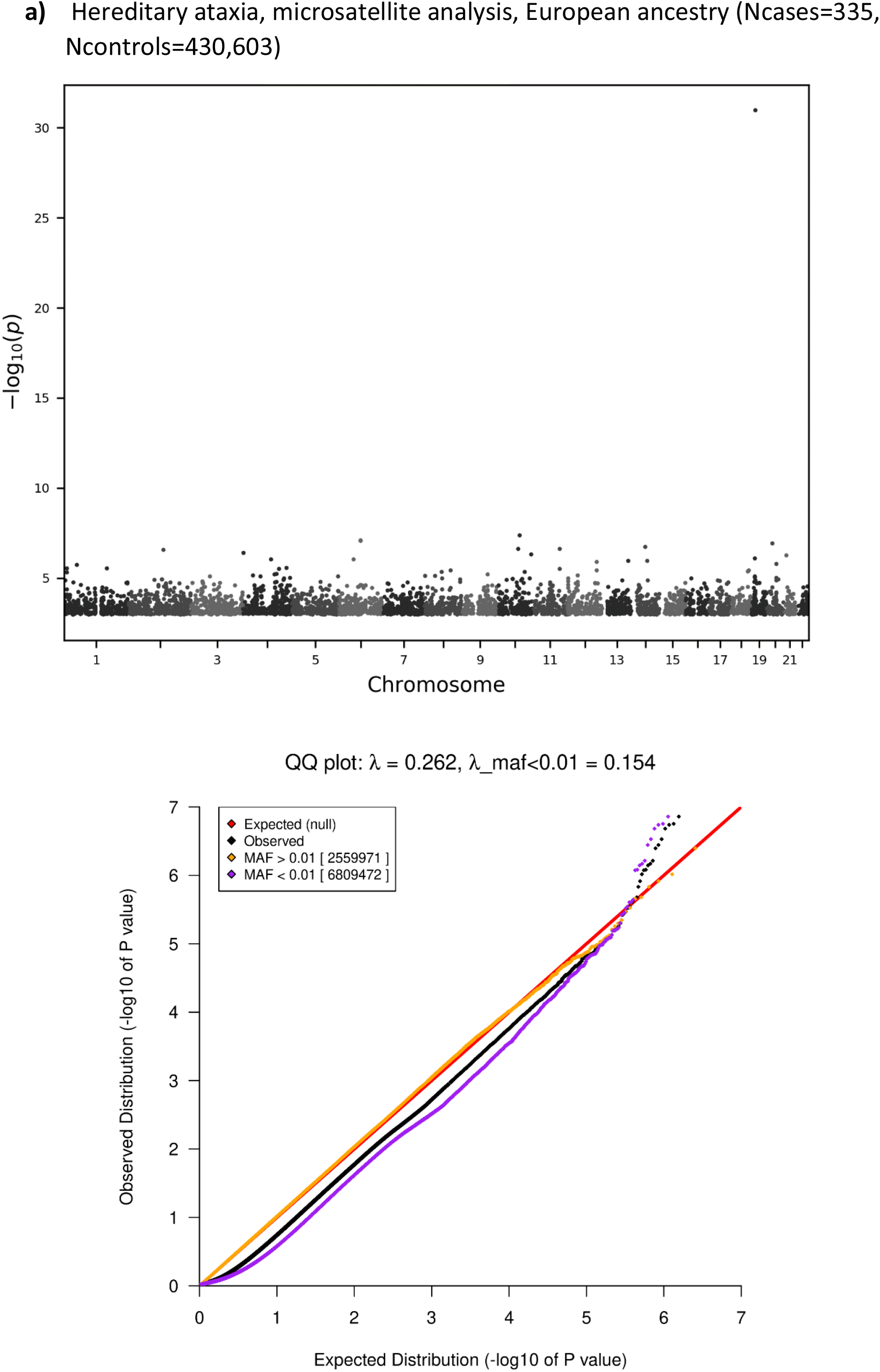
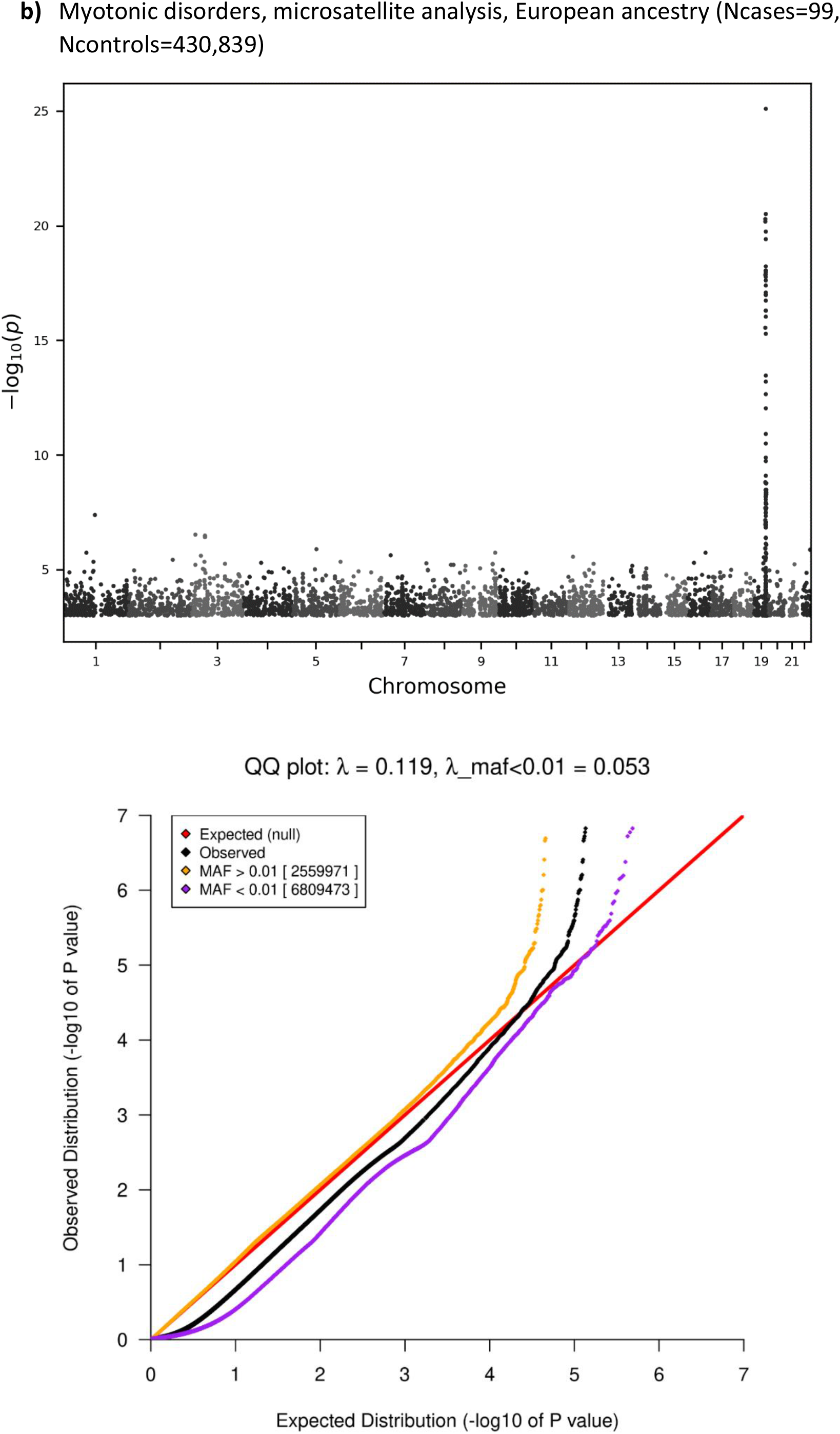
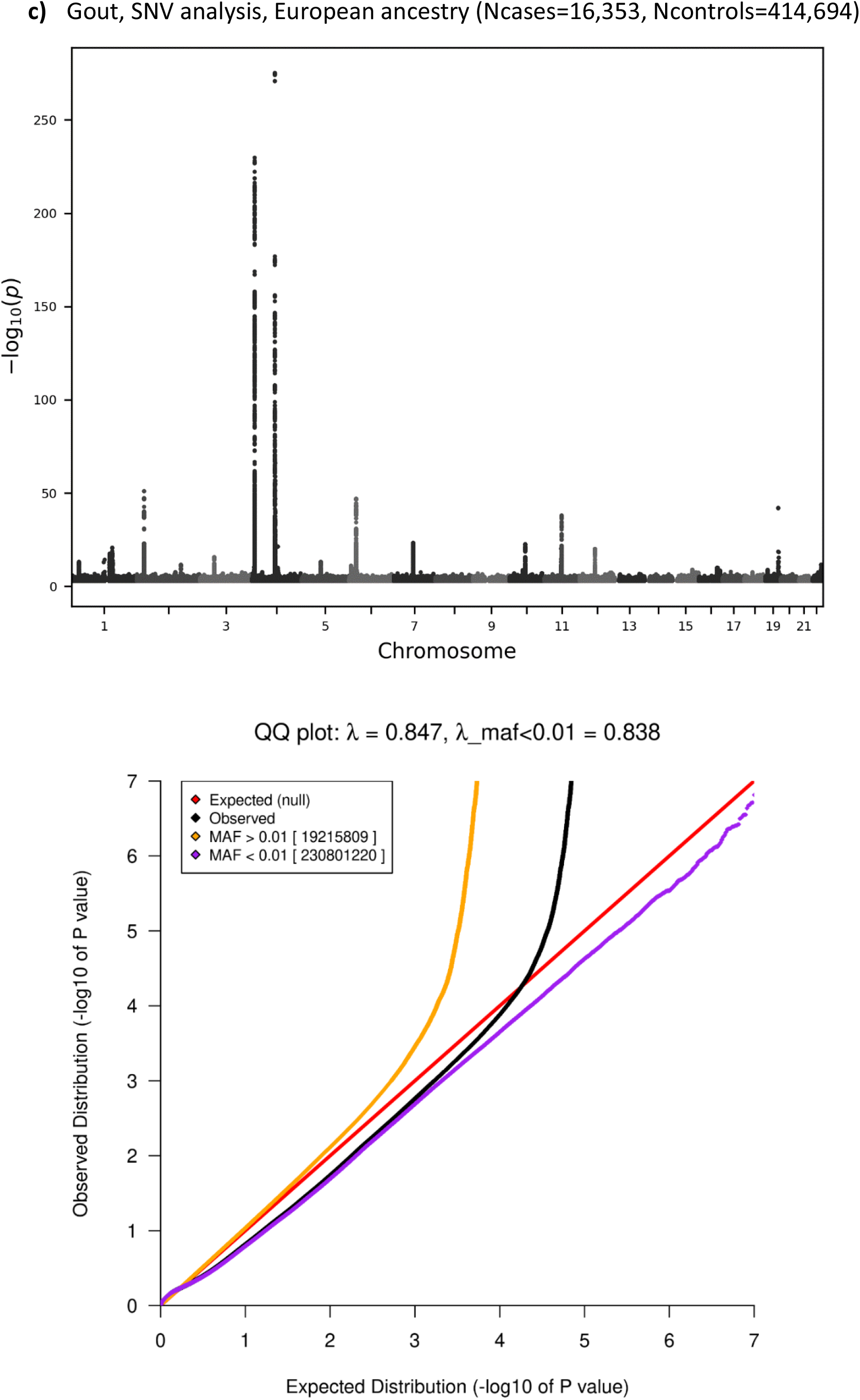
Manhattan plots and quantile-quantile (QQ) plots for case-control phenotypes with significant results reported in this manuscript. For Manhattan plots, the x-axis represents chromosome locations and the y-axis shows the –log10 significance levels of the associations. For QQ plots, the inflation (λ) is shown in the title of each graph, for all variants and for rare variants only (λ_maf<0.01)

**Fig. S27.**
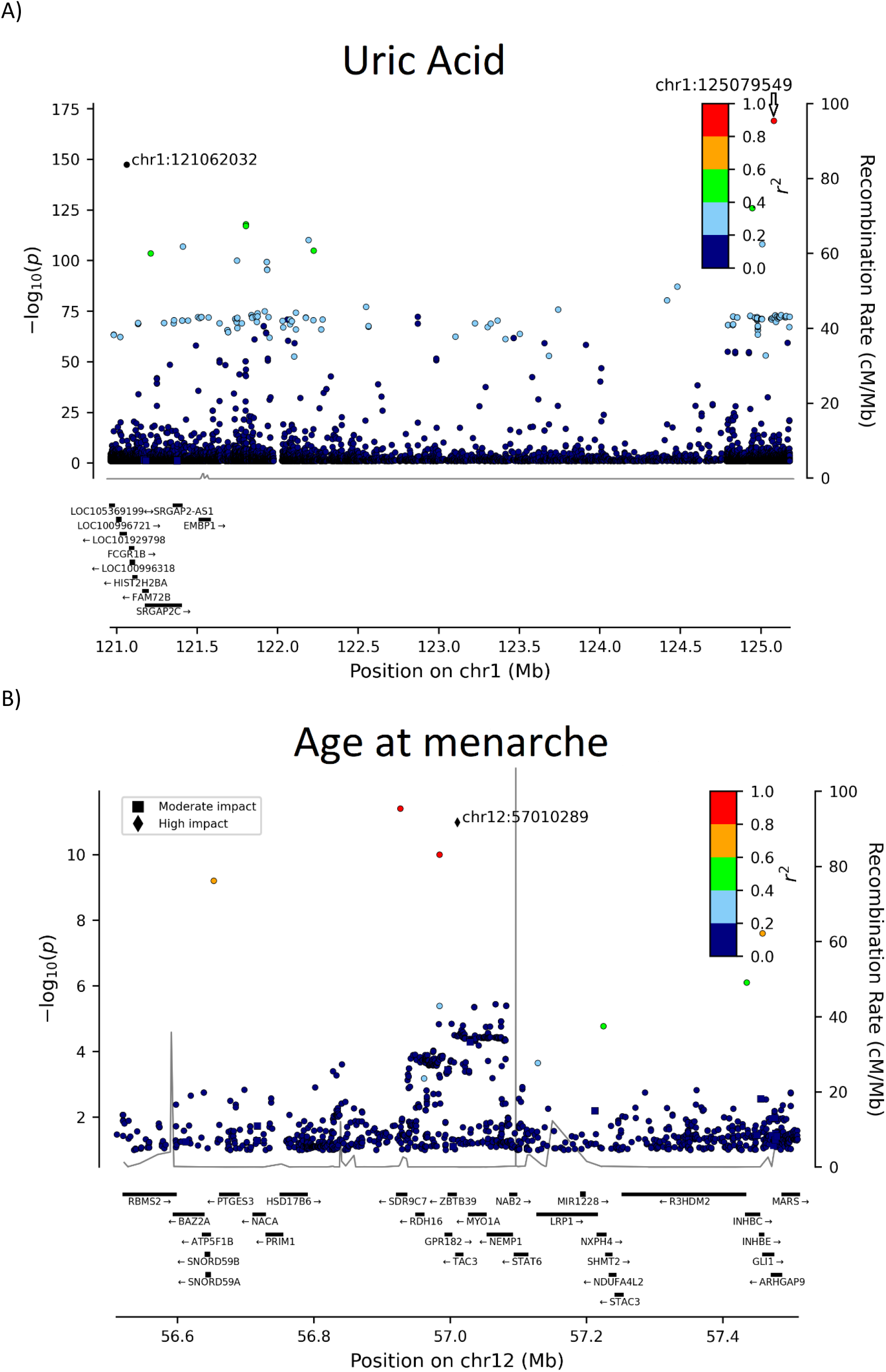
Locus plot for A) Uric acid and B) Age at menarche associations.

**Fig. S28.**
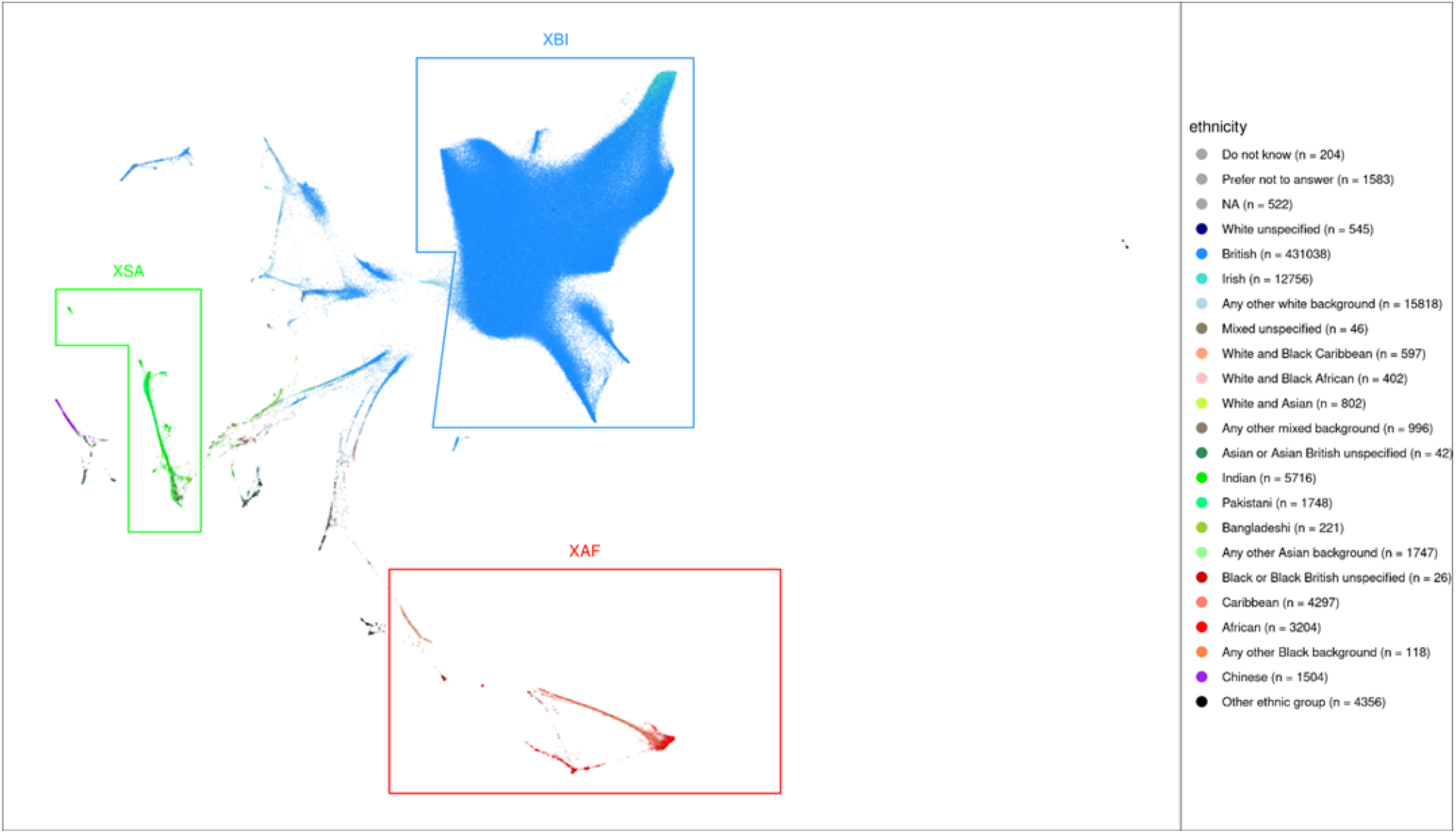
UMAP and ethnicity. 40 genetic principal components provided by UKB reduced to a latent space of 2 dimensions using UMAP (x and y axes). Individuals are colored according to self-identified ethnicity. The regions defined to delineate the three cohorts XAF, XBI, and XSA are indicated.

**Fig. S29.**
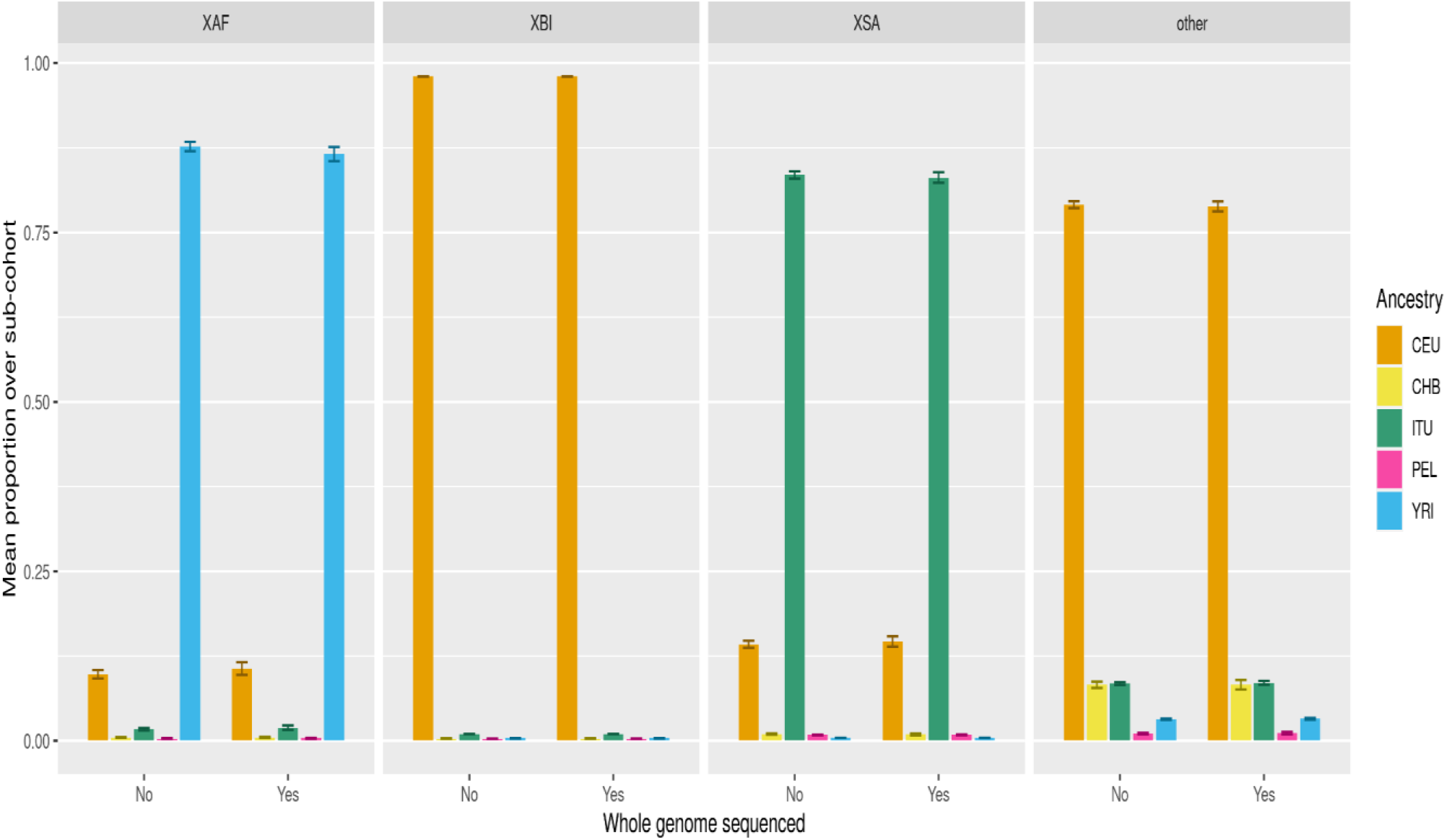
Cohort mean ADMIXTURE. Mean proportion of each of five 1000 Genome Project ancestry components assigned by ADMIXTURE (columns). Error bars represent 99.9% confidence intervals. CEU (Northern Europeans from Utah), CHB (Han Chinese in Beijing), ITU (Indian Telugu in the UK), PEL (Peruvians in Lima), and YRI (Yoruba in Ibadan, Nigeria).

**Fig. S30.**
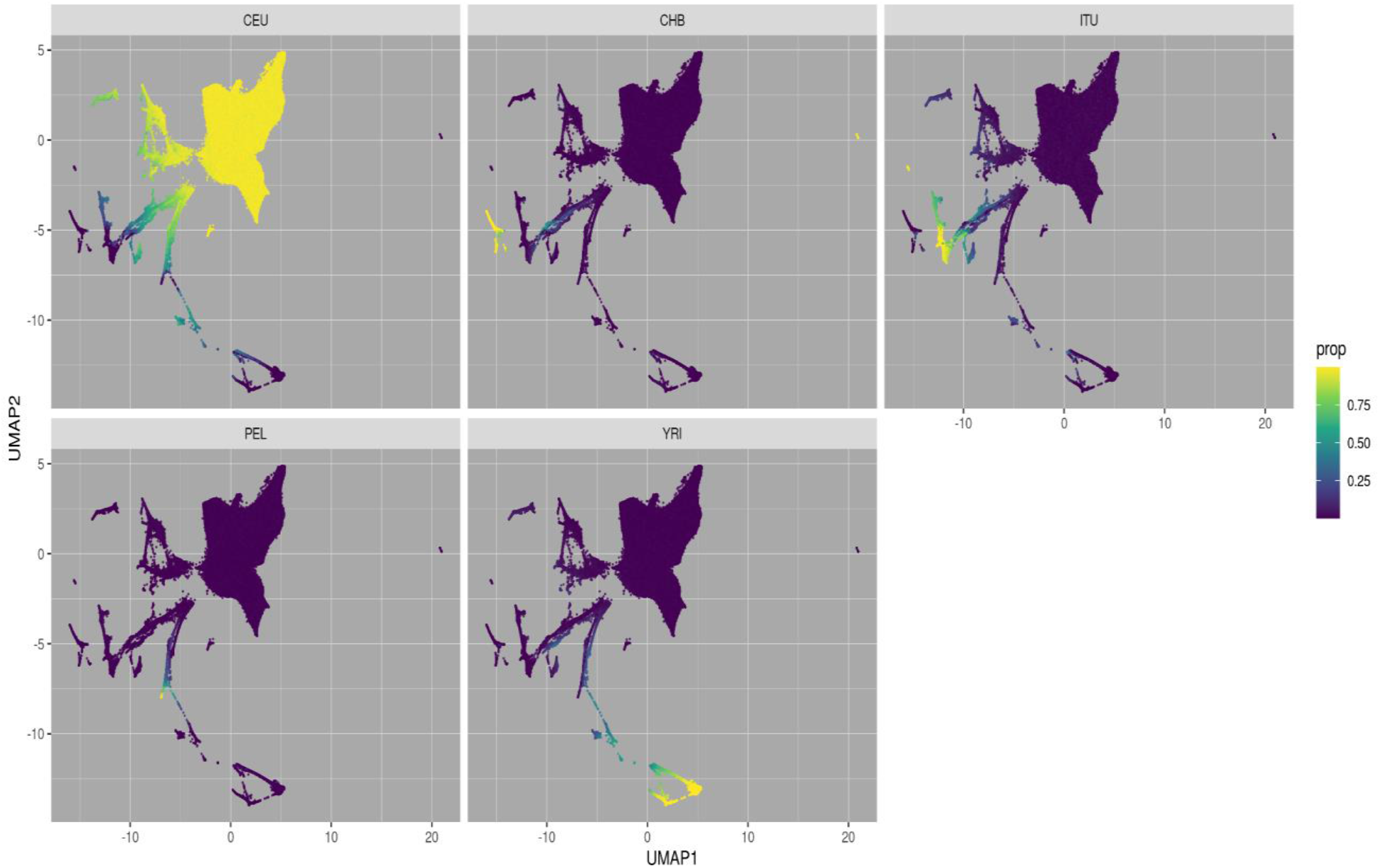
UMAP ADMIXTURE. 40 genetic principal components provided by UKB reduced to a latent space of 2 dimensions using UMAP (x and y axes). Individuals are colored according to proportion of ancestry assigned by supervised ADMIXTURE from five 1000GP training populations (facet headings): CEU (Northern Europeans from Utah), CHB (Han Chinese in Beijing), ITU (Indian Telugu in the UK), PEL (Peruvians in Lima), and YRI (Yoruba in Ibadan, Nigeria).

**Fig. S31.**
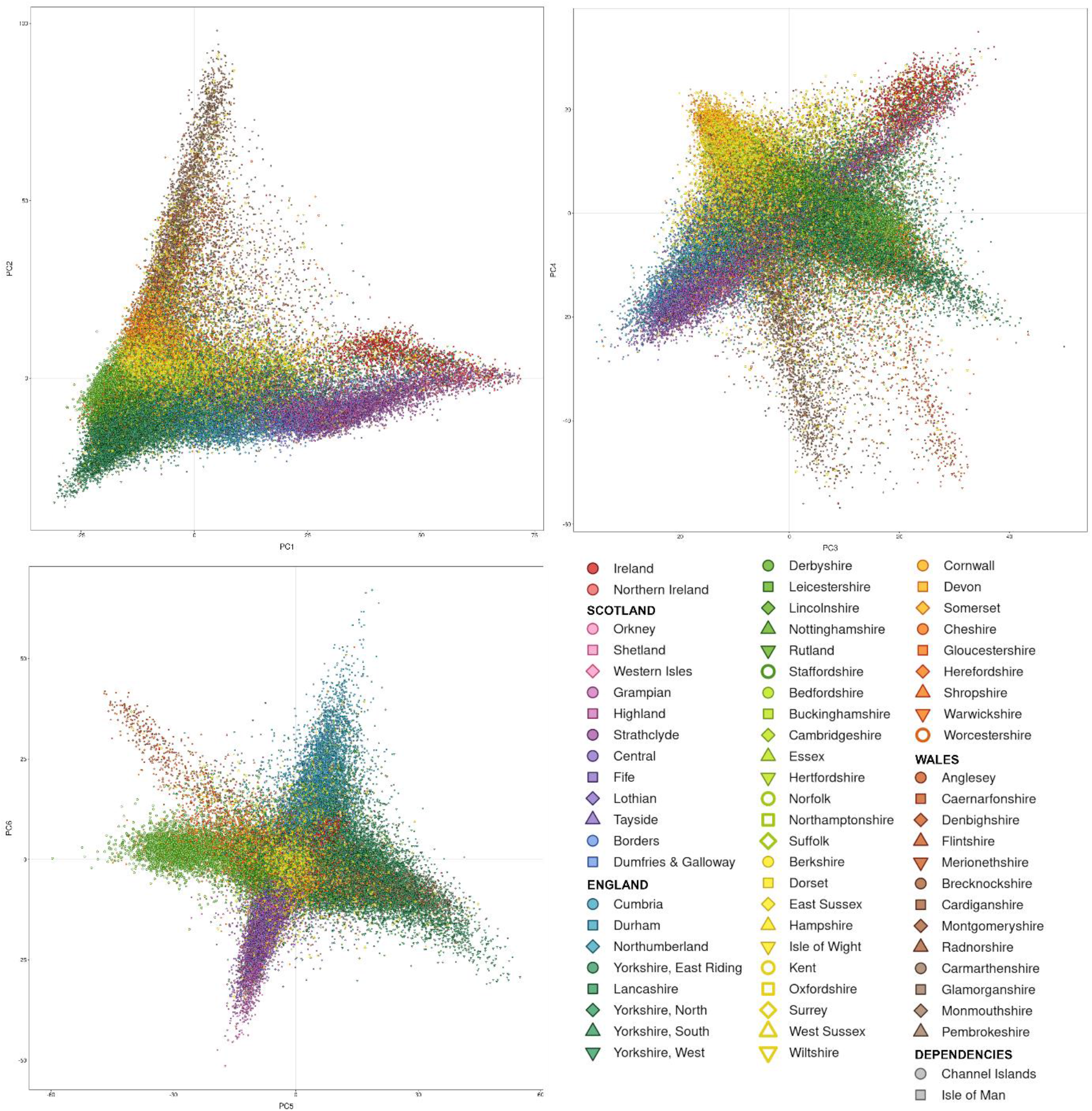
The first six principal components of the XBI cohort, plots show PC1 vs PC2, PC3 vs PC4 and PC5 vs PC6. Points represent individuals, colored by place of birth. To show geographic structure in the UK more clearly, we do not show individuals who report being born in urban areas with many internal migrants (Tyne & Wear, Merseyside, Greater Manchester, West Midlands, Bristol, London) or places outside the British-Irish Isles.

**Fig. S32.**
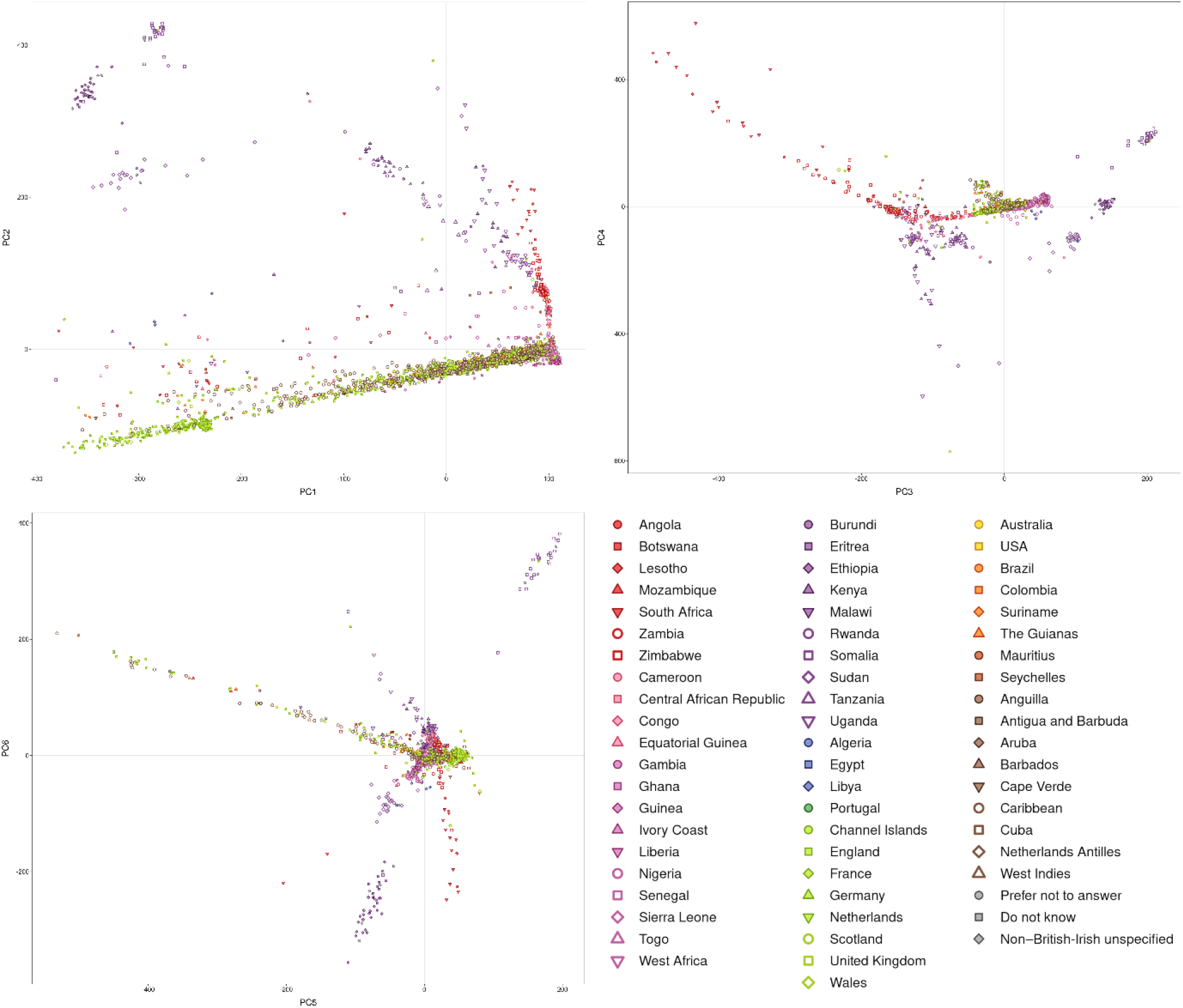
The first six principal components of the XAF cohort, plots show PC1 vs PC2, PC3 vs PC4 and PC5 vs PC6. Points represent individuals, colored by place of birth.

**Fig. S33.**
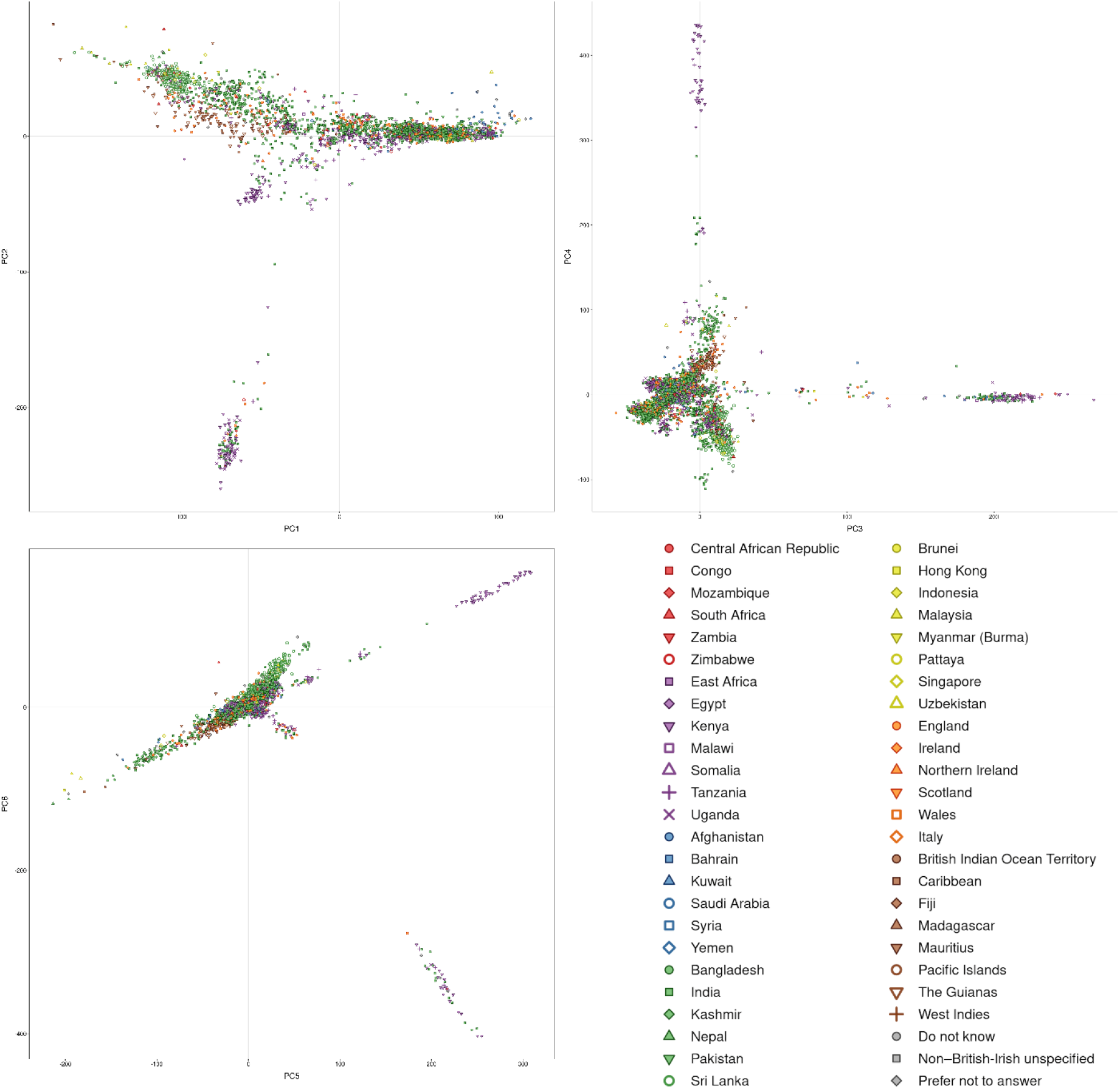
The first six principal components of the XSA cohort, plots show PC1 vs PC2, PC3 vs PC4 and PC5 vs PC6. Points represent individuals, colored by place of birth.

### Supplementary Tables

**Table S1.**
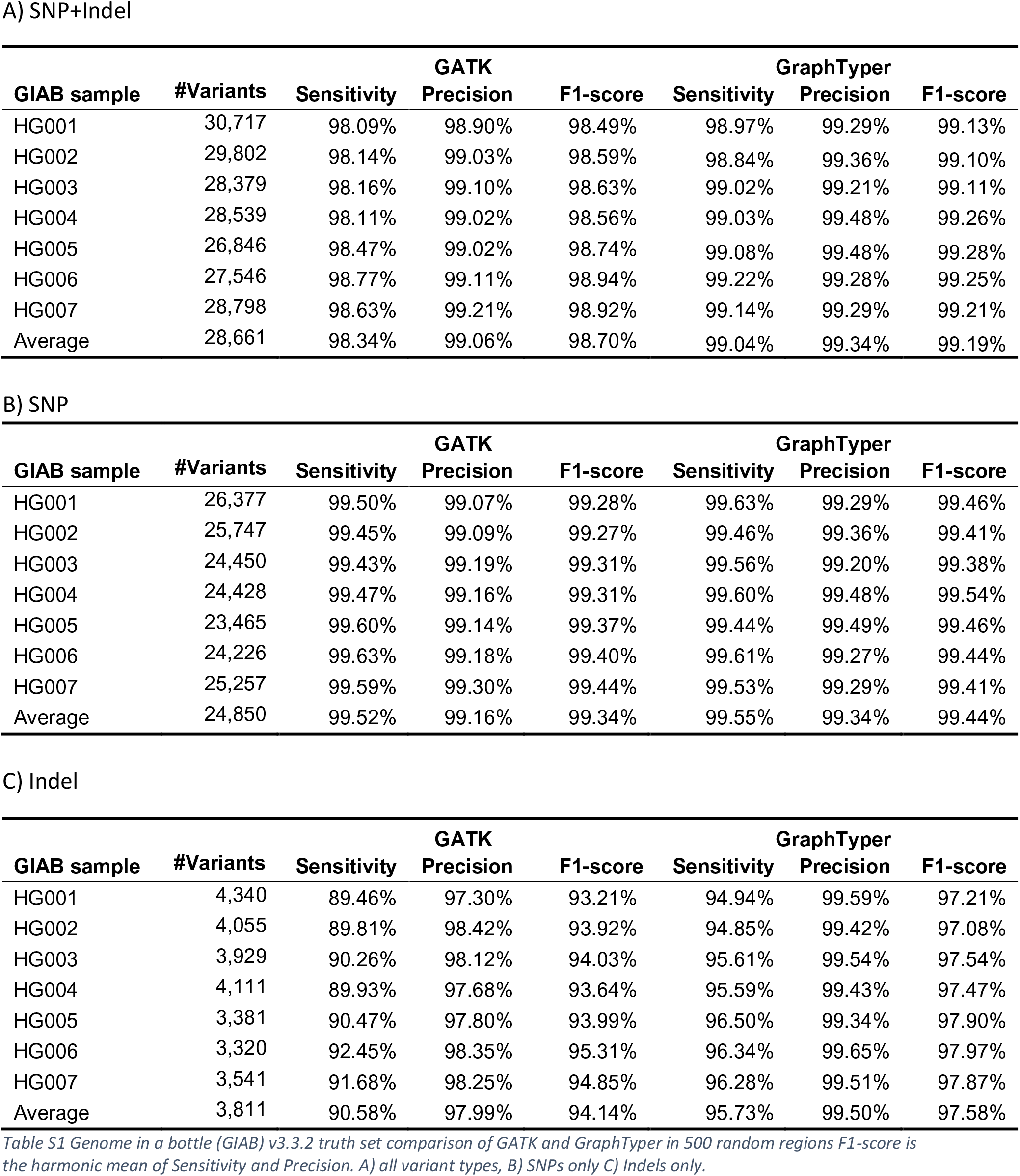
Genome in a bottle (GIAB) v3.3.2 truth set comparison of GATK and GraphTyper in 500 random regions F1-score is the harmonic mean of Sensitivity and Precision. A) all variant types, B) SNPs only C) Indels only.

**Table S2.**
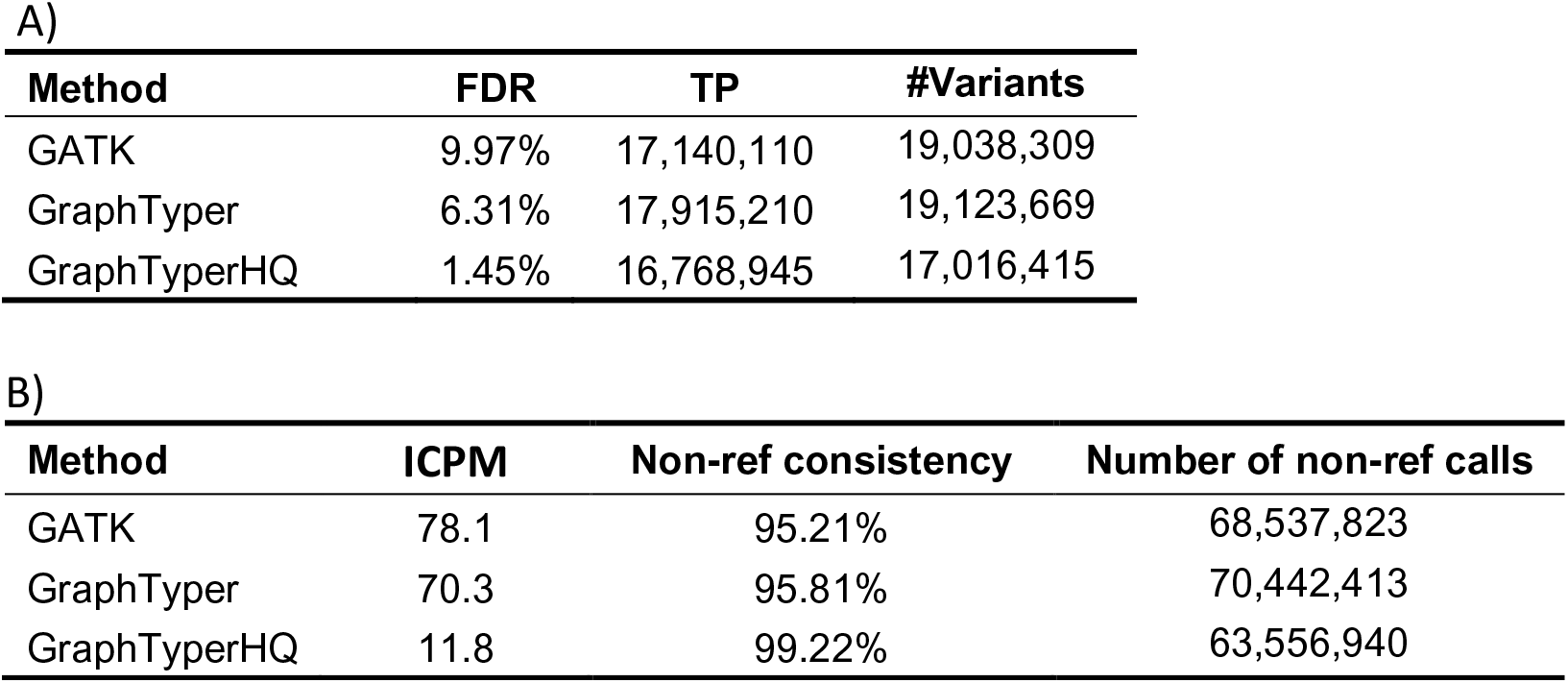
A) Estimate of false discovery rate (FDR) and number of true positive (TP) variants among the 28 parent-offspring trios. The estimates are determined from the allele transmission ratios from parent to offspring. B) Genotype consistency across among the 14 monozygotic twin pairs. ICPM = number of inconsistent genotypes per 1Mb.

**Table S3.**
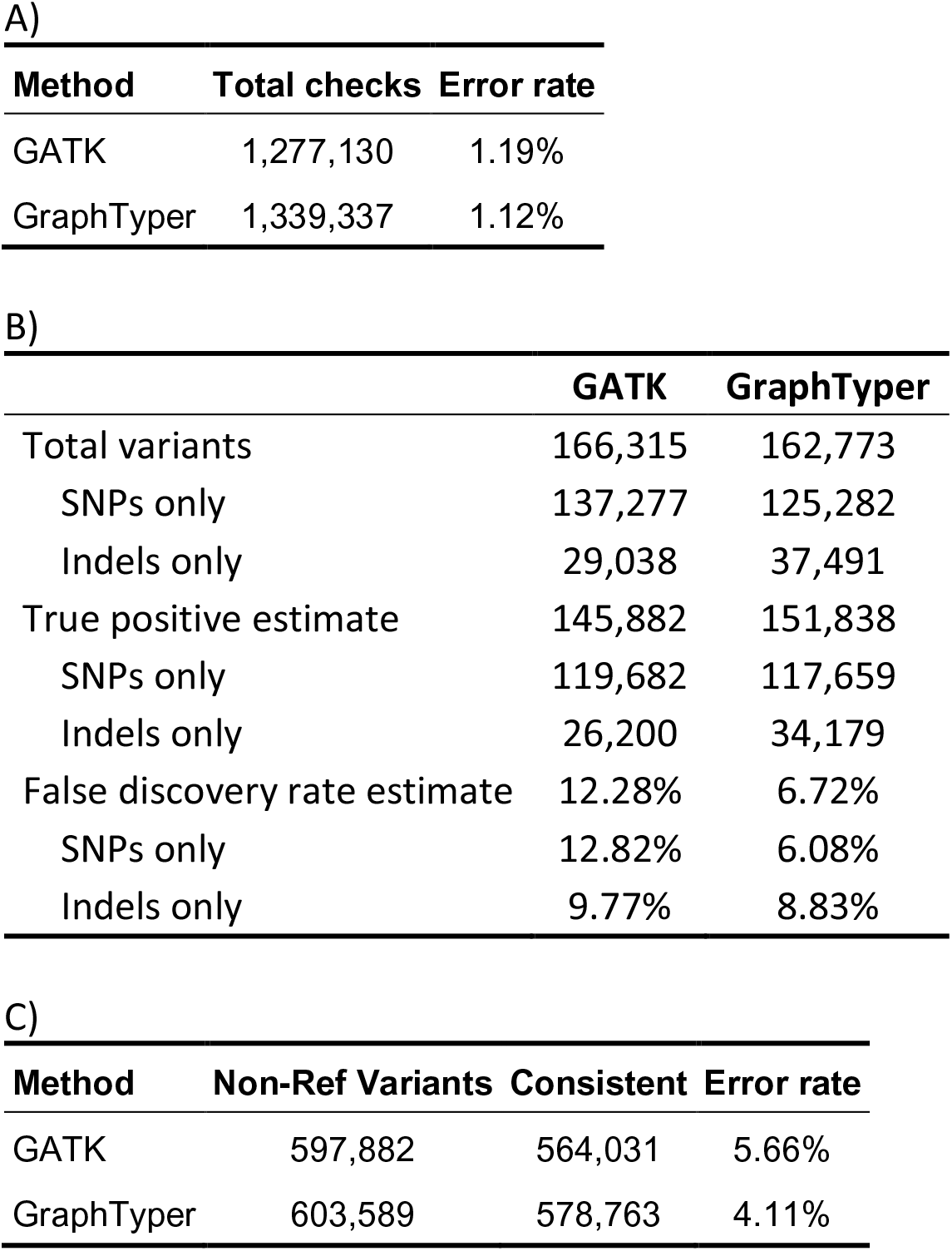
Analysis of variant transmission of related samples in the 500 randomly selected 50kb test regions. A) Number of inheritance errors among the 28 parent-offspring trios. B) Estimates of number True Positives and False discovery rate in GATK and GraphTyper datasets in the trios. The estimates are determined from the allele transmission ratios from parent to offspring. C) Genotype consistency among the 14 pairs of monozygote twins.

**Table S4.**
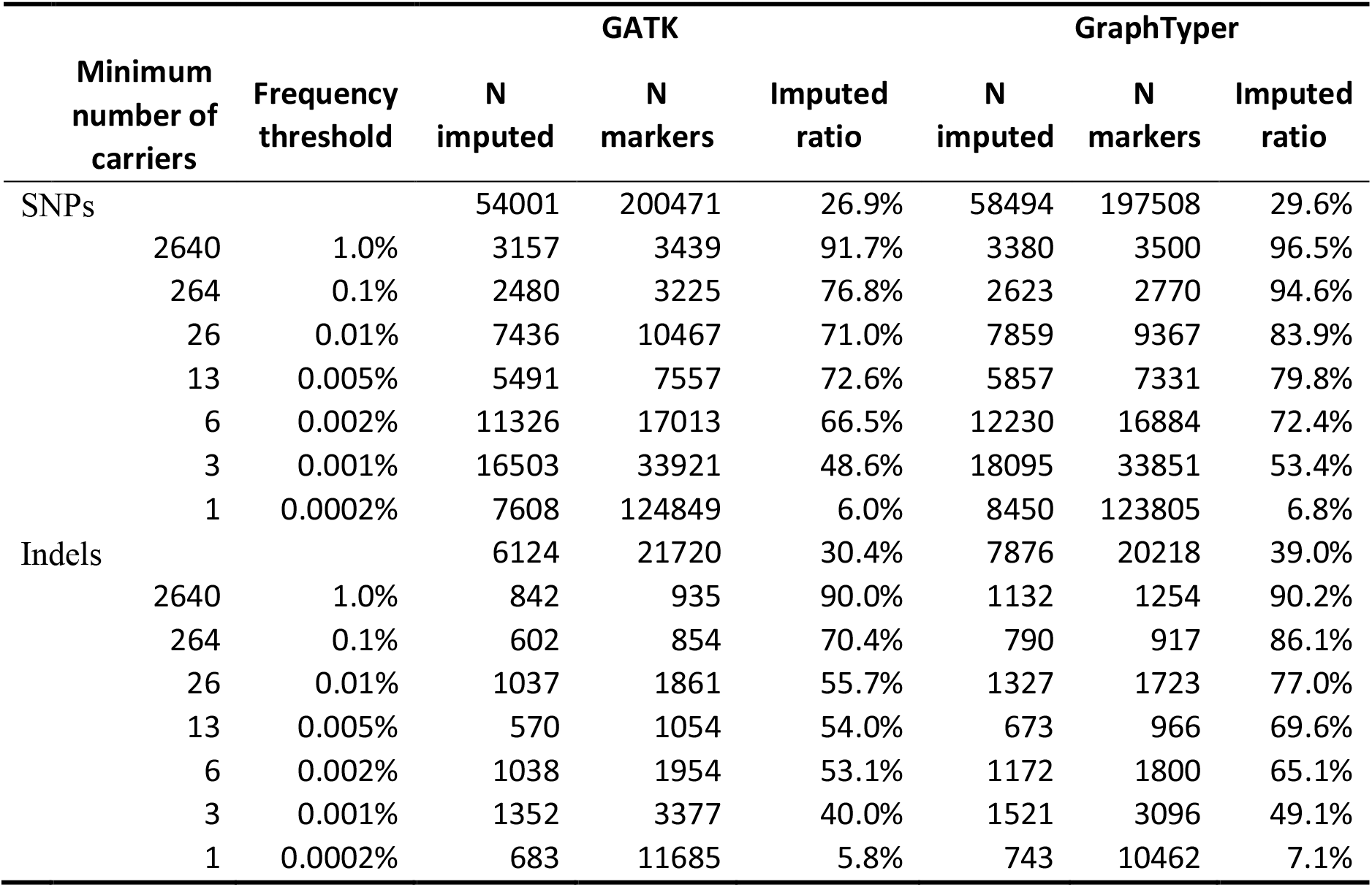
Comparison of imputation of variants from the GATK and GraphTyper call sets on chr22 10-11Mb in the XBI dataset. A variant is considered imputed if phasing Leave-on-out-r2 (L1or2) is greater than 0.5 and imputation info is greater than 0.8.

**Table S5.**
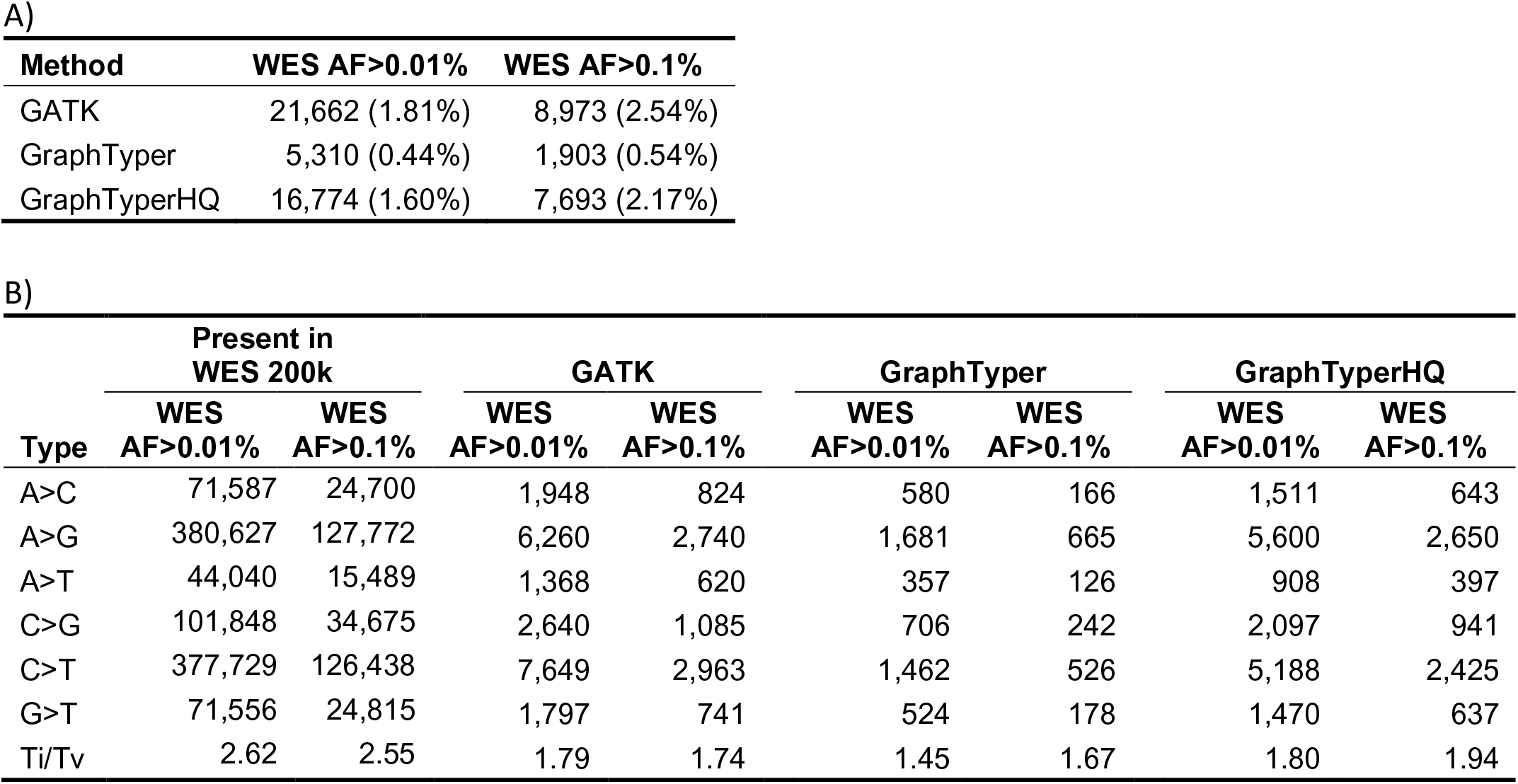
A) Number of variants in the WES 200k dataset that are missing from GATK, GraphTyper and GraphTyperHQ datasets, conditioned on the frequency in WES 200k. The fractions of missing variants are inside the parenthesis. B) Total number of SNP types present in WES 200K conditioned on frequency and how many of those are missing from our WGS datasets, stratified by variant type. Ti = number of transitions, Tv = number of transversions.

**Table S6.**
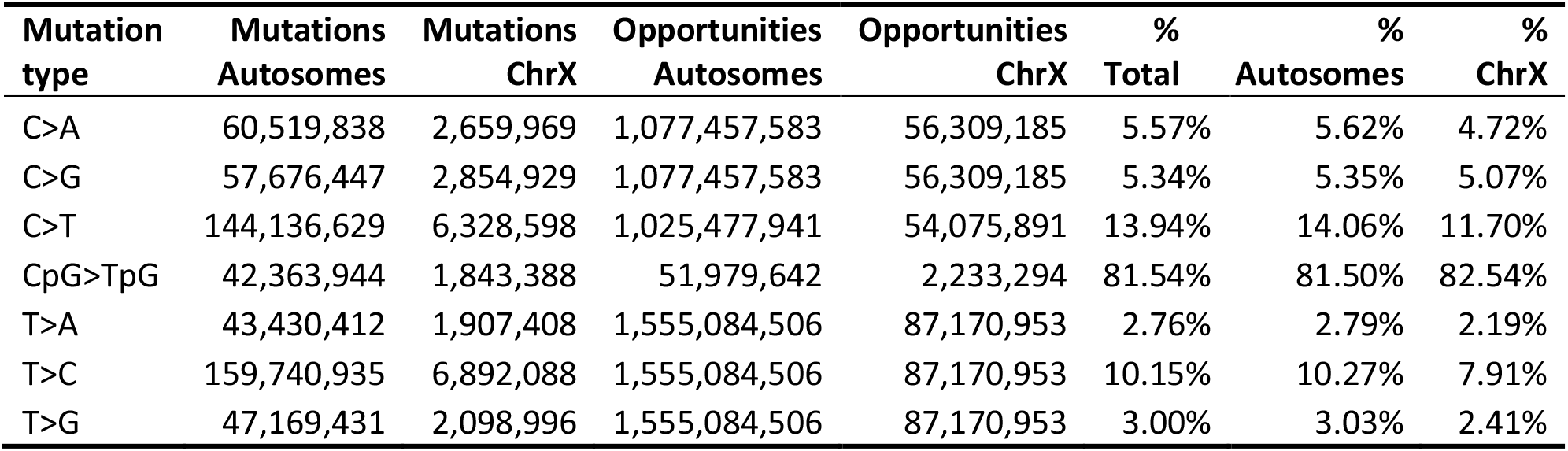
Mutation saturation, results presented for autosomes and chrX separately. Table shows the number of observed mutations in the GraphTyperHQ dataset and the number of possible mutation opportunities in regions of the genome amenable to short read sequence analysis.

**Table S7.**
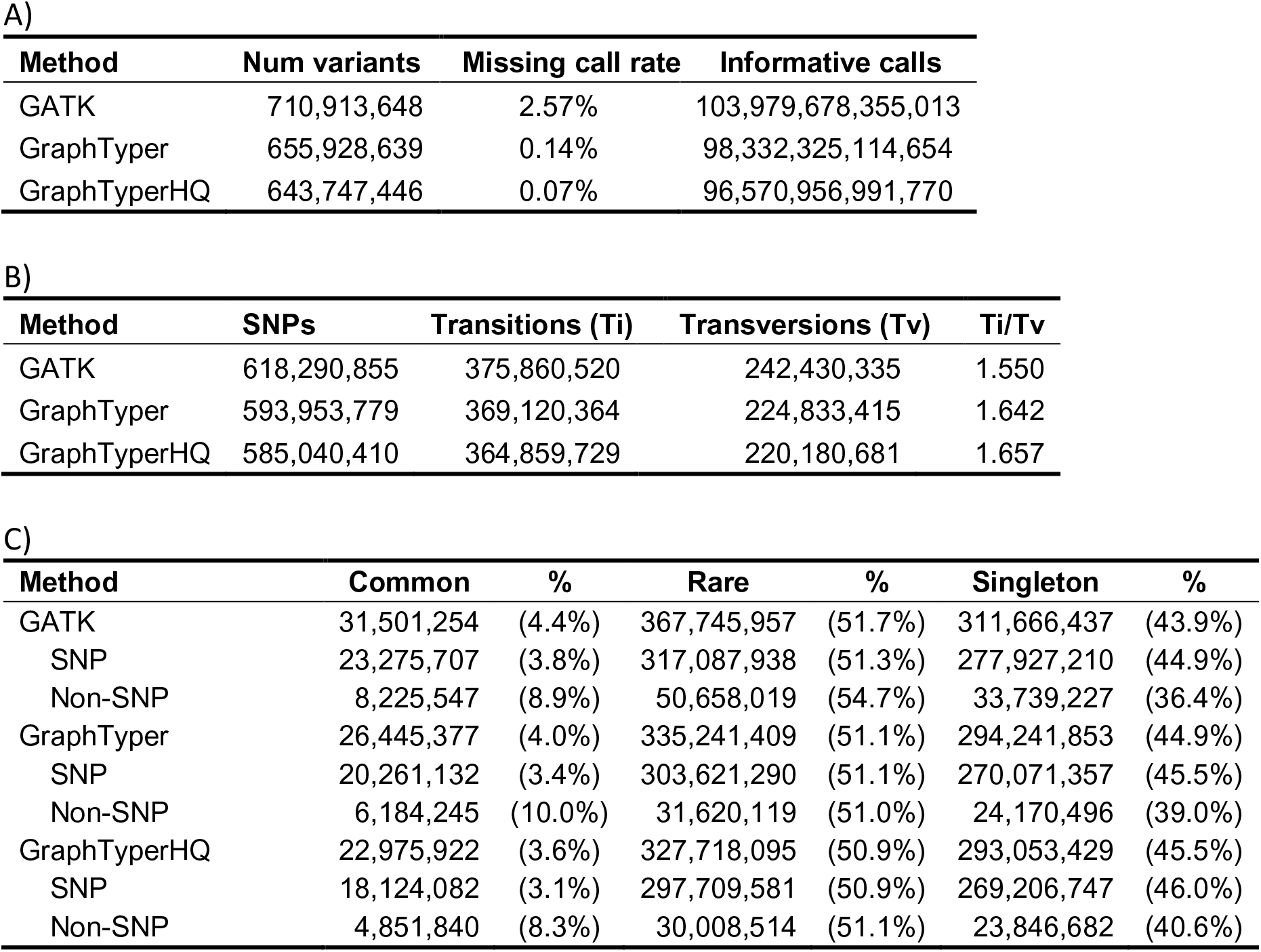
A) Number of variants in GATK, GraphTyper and GraphTyperHQ dataset. B) Variants split by transitions and transversions. C) Common = variants with frequency > 0.1%, rare = carried by more than one individual and frequency < 0.1%, singleton = carried by a single individual.

**Table S8.**
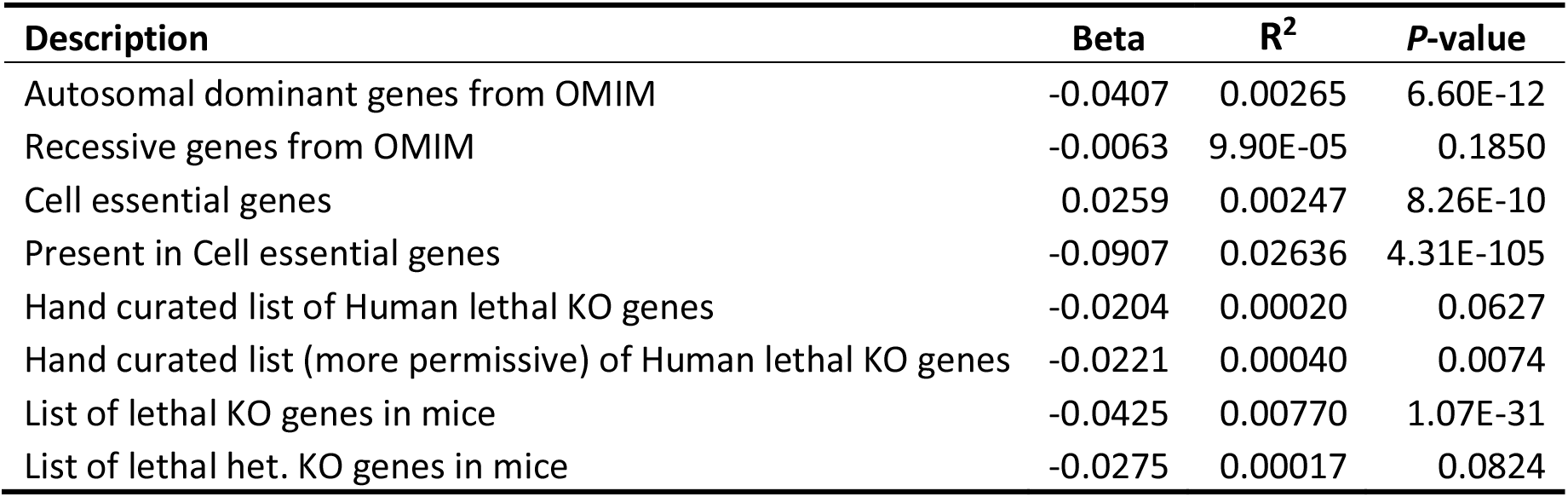
Regression of average DR overlapping gene exons on annotations from Gene discovery informatics toolkit^40^.

**Table S9.**
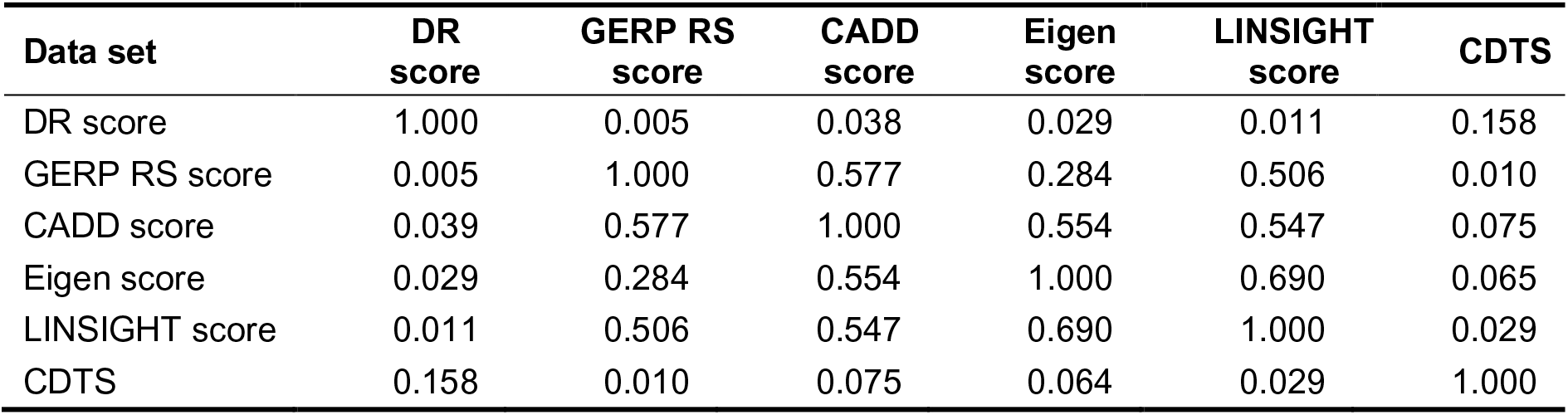
Pearson correlation coefficient between DR score and measures of sequence constraint and functional impact, computed over all autosomal chromosomes. For each one of the 500bp overlapping windows in which the DR score (*dr*) is defined we compute the average value of the published scores (*ps*) in that window and then conduct linear regression analysis (*ps* ∼ *dr*). The values shown in the table are the squared correlation coefficients of that regression. The correlation between the published datasets is computed from a set of 50bp non-overlapping windows using the average score within each window. A similar regression is conducted between each of the published datasets to obtain the squared correlation coefficient. Note, that the p-value for the linear regression fit is below computational threshold (2.2 × 10^−308^) for each pair of data sets in the table. CADD, Eigen and LINSIGHT all incorporate GERP into their annotation and are consequently not independent of each other or GERP. DR score and CDTS employ an analogous methodology, but scores are derived independently of each other and the other metrics.

**Table S10.**
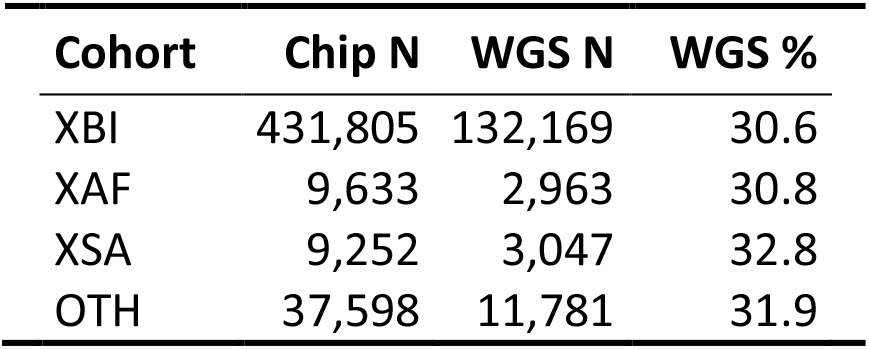
Number of individuals in the three cohorts described in this study.

**Table S11.**
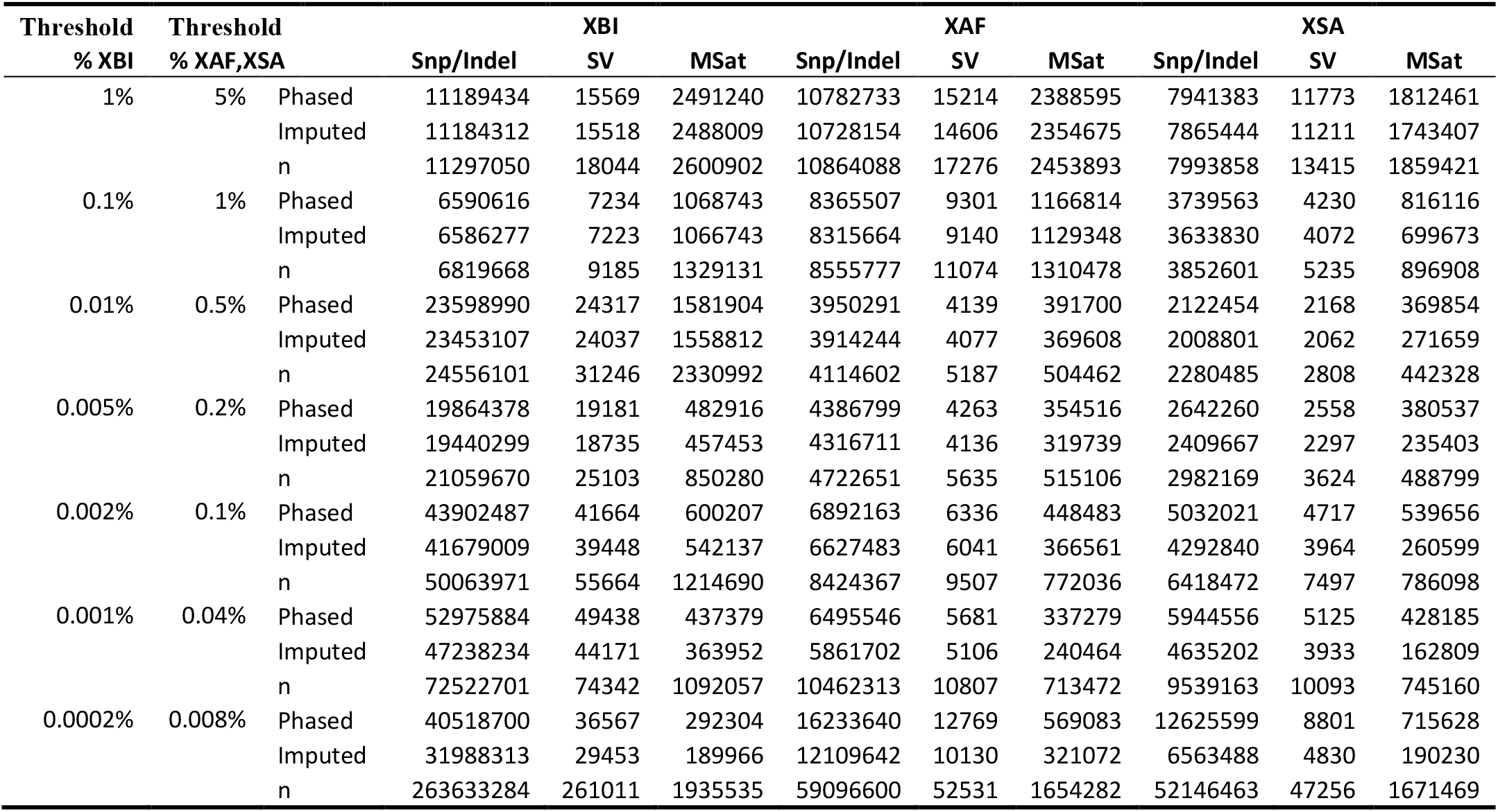
Imputation and phasing accuracy as a function of frequency within each cohort. Phased refers to number of variants with Leave-one-out-r2 value > 0.5 and imputed refers to phased variants that also have imputation info > 0.8. Numbers are for variants at frequency above the given threshold and not included in frequency thresholds in earlier lines, e.g., in the XBI population 72,522,701 variants have frequency between 0.001 and 0.002%, of which 52,975,884 could be phased and 47,238,234 could be imputed.

**Table S12.**
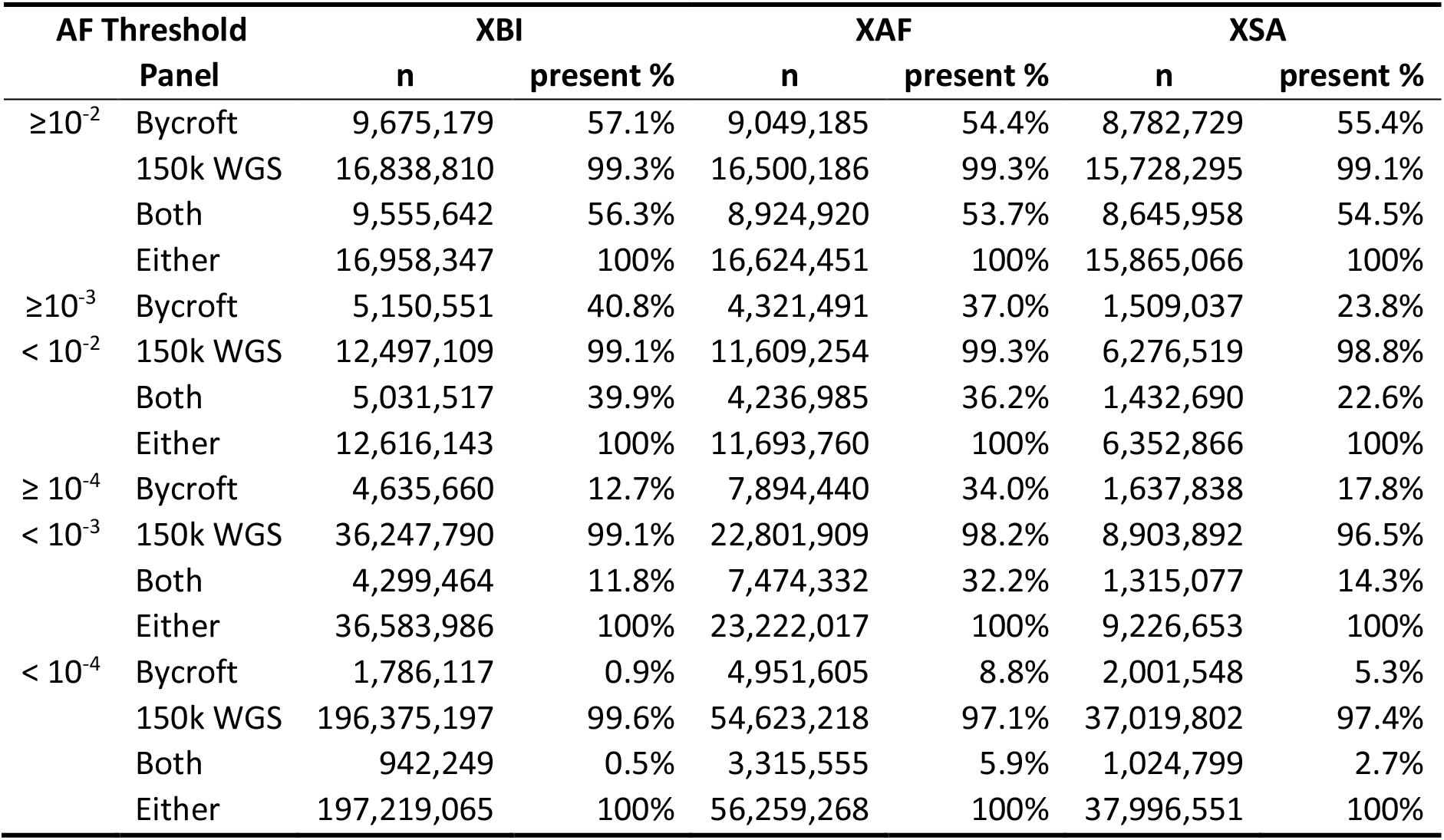
Number of markers that impute (Imp Info > .8) in 500k set of UKB using the imputation panel presented here (150k WGS) and an imputation by Bycroft et al.^5^. Both represents number of markers imputed by both panels, either the number of markers in either panel.

**Table S13.**
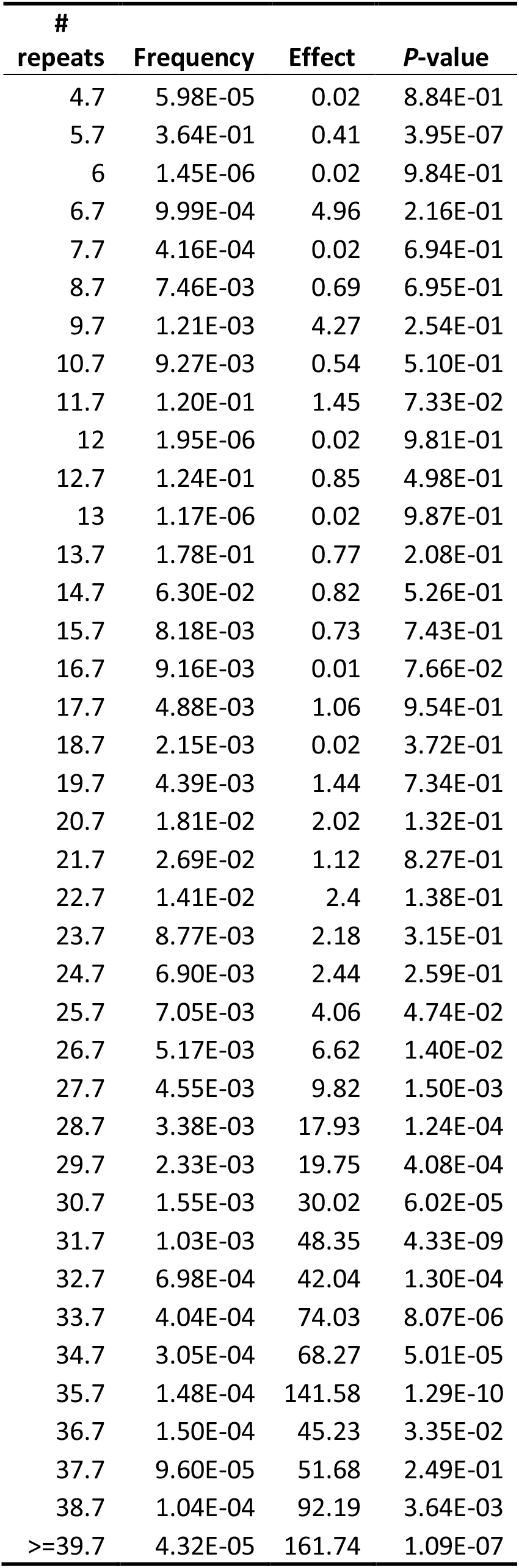
Association of number of repeat copies of microsatellite in 3’ UTR in DMPK with myotonic dystrophy. Individuals carrying 39.7 or more copies of the repeat are grouped together by popSTR^64^.

**Table S14.**
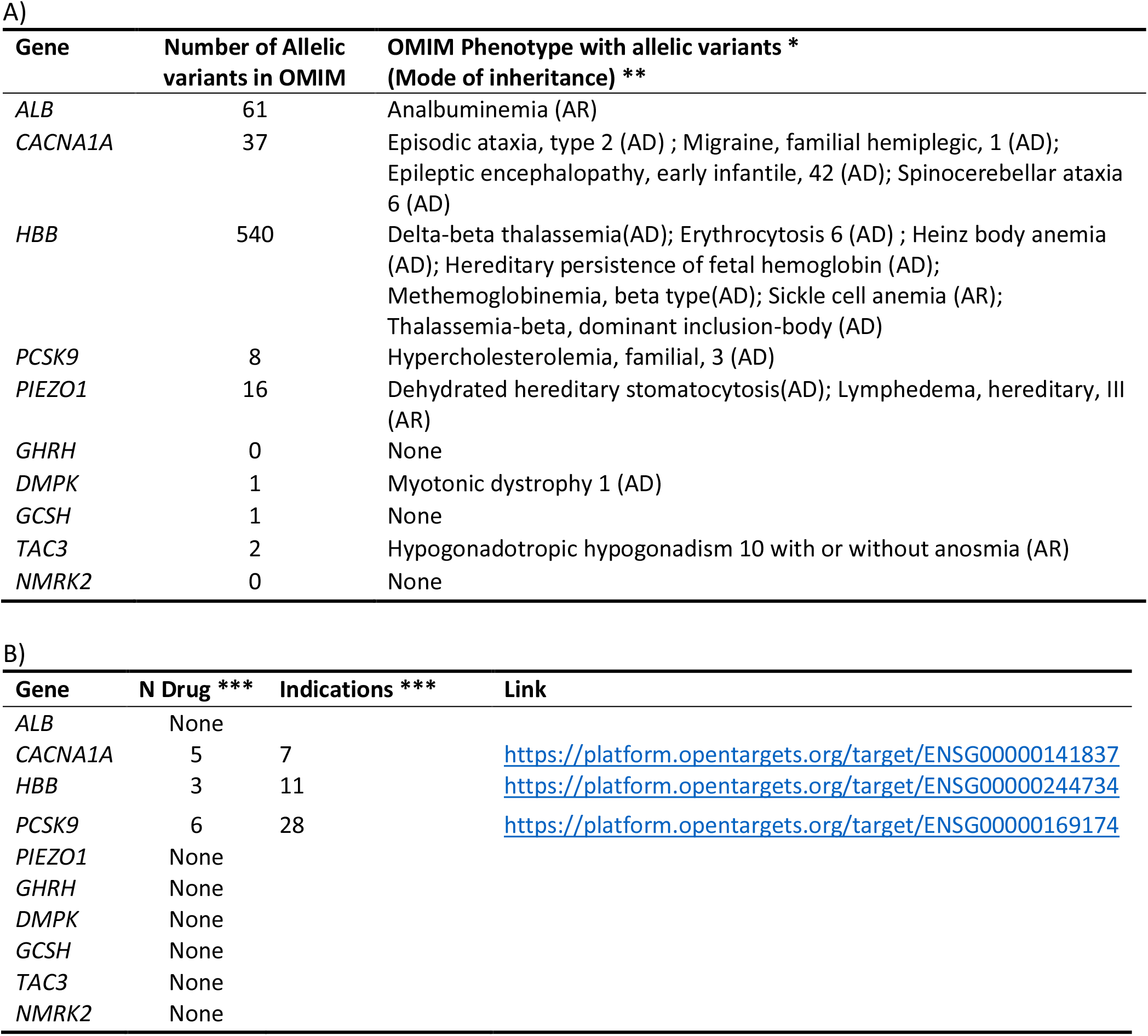
Information on genes presented. A) Phenotypes and allelic variants in OMIM for selected genes. B) Known drug data and in open targets for selected targets. *Excluding the ones with provisional phenotype gene relationship “?”; multifactorial diseases”{ }” and non diseases”[]” ** Mode of inheritance : AD Autosomal dominant; AR Autosomal recessive. ***Known drug data according to Open Targets^77^.

**Table S15.**
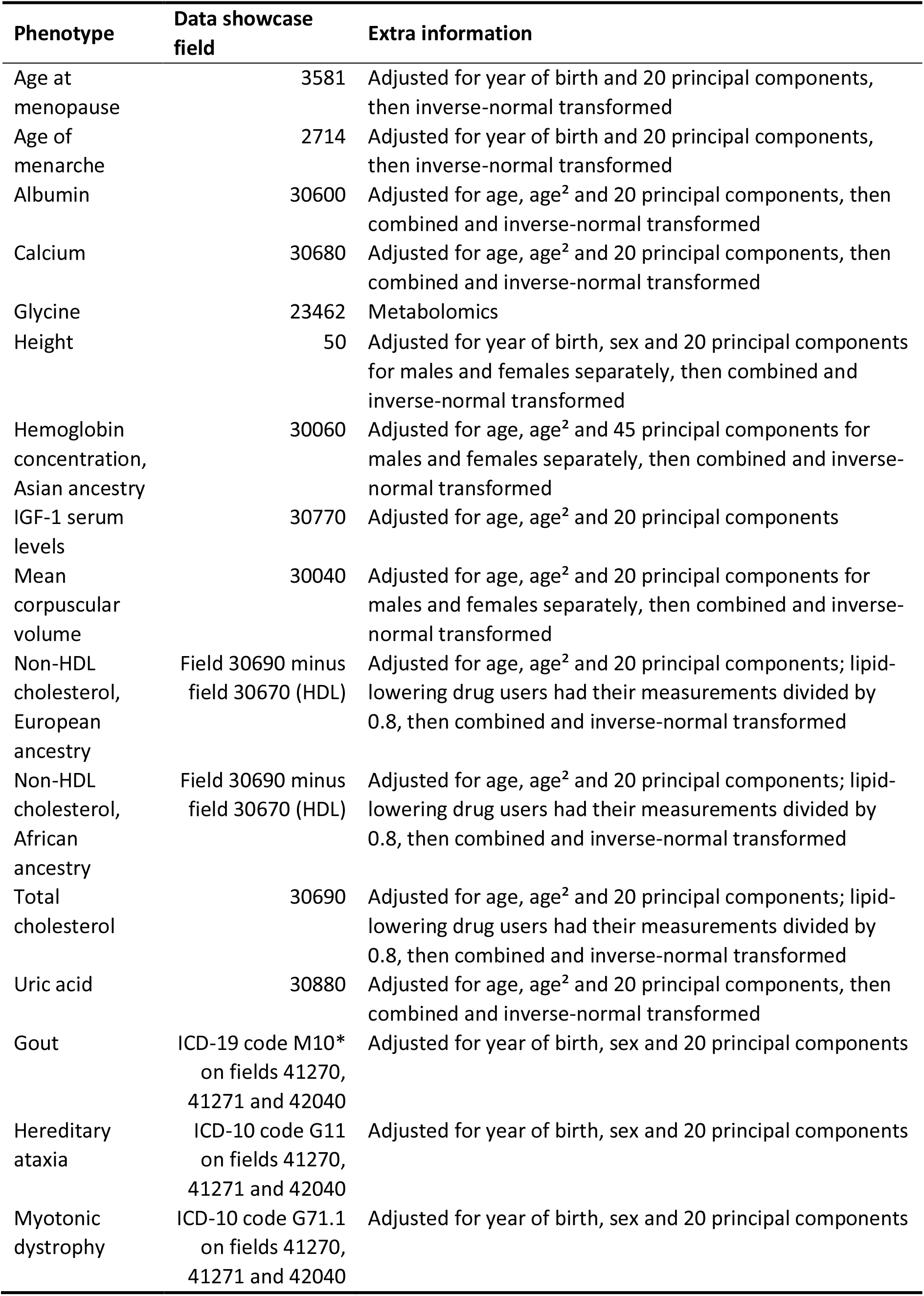
Phenotypes used in this study, their field in the UKB data showcase and adjustments performed prior to association analysis

**Table S16.**
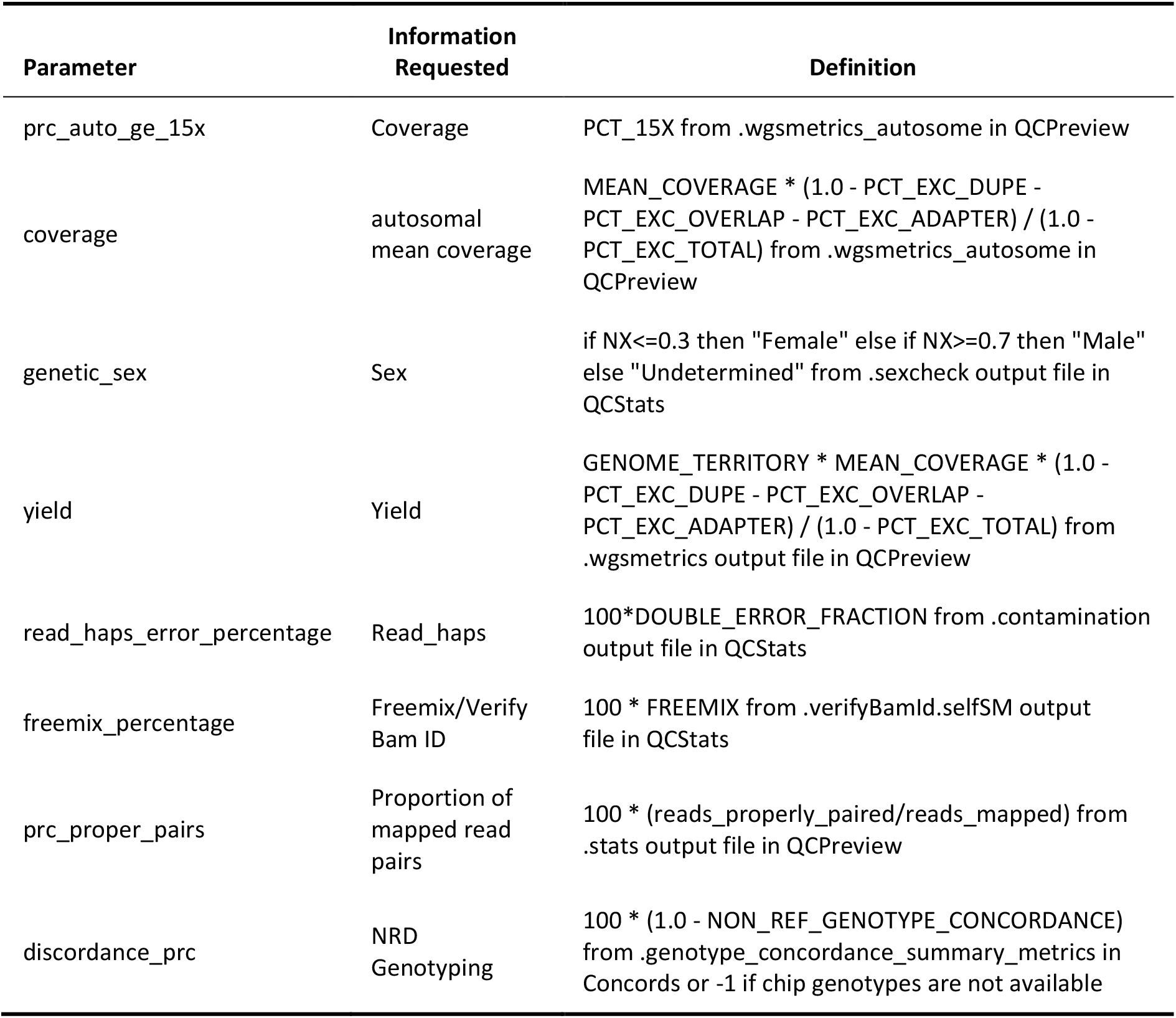
QA/QC metrics derived from the files delivered to the UKB. The result is written to a file, qaqc_metric.

**Table S17.**
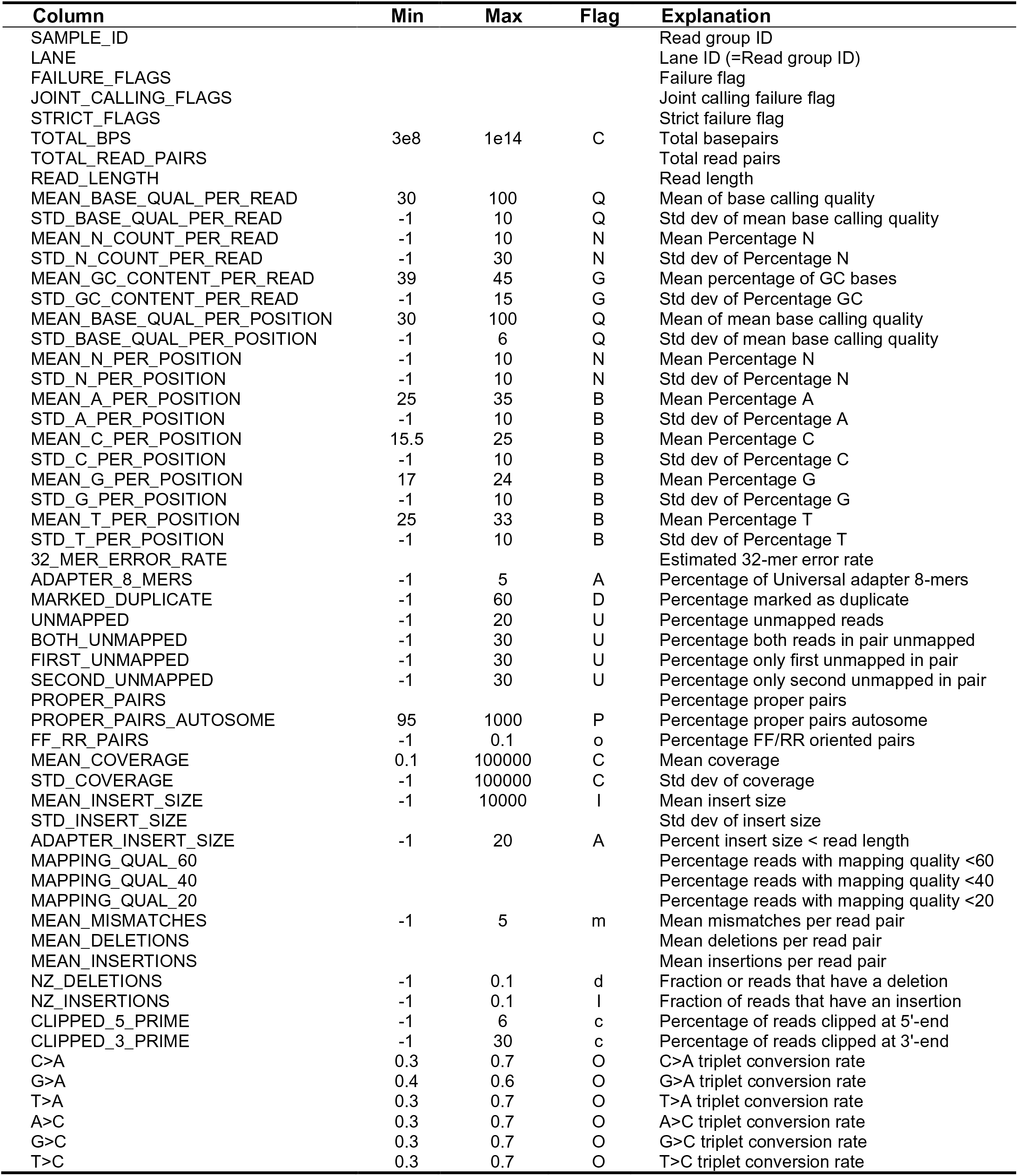
Metrics collected for each lane by bamqc_summary. If any flag is raised, the lane is excluded from the merge process. The values, per read group, are collected in the file .bamqc_summary.

**Table S18.**
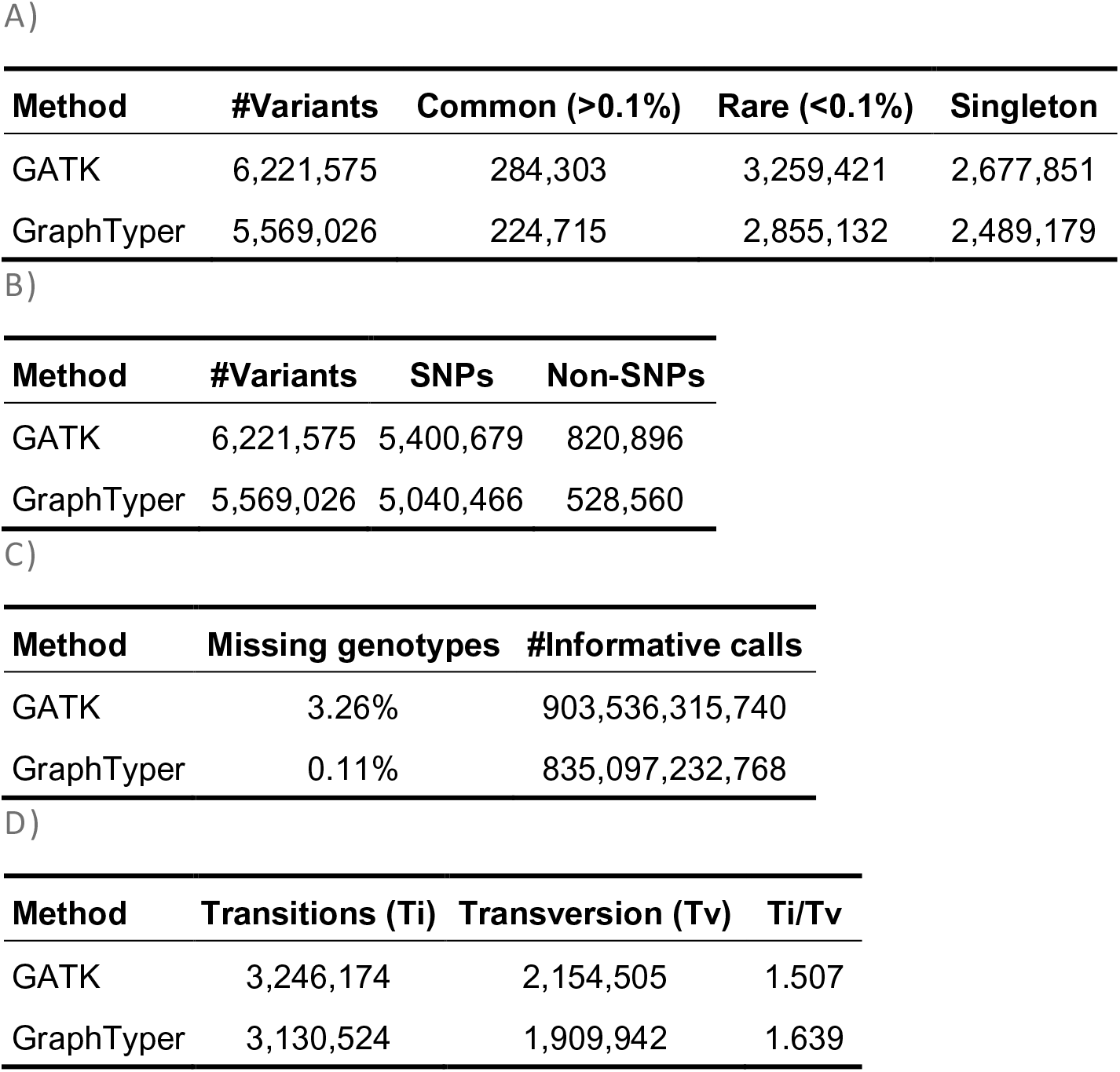
Results for 500 random test regions. A) Number of variants called by GATK and GraphTyper conditioned on frequency class. B) Number of variants conditioned on variant type. C) Fraction of missing variant calls. D) Number of transitions and transversions.

**Table S19.**
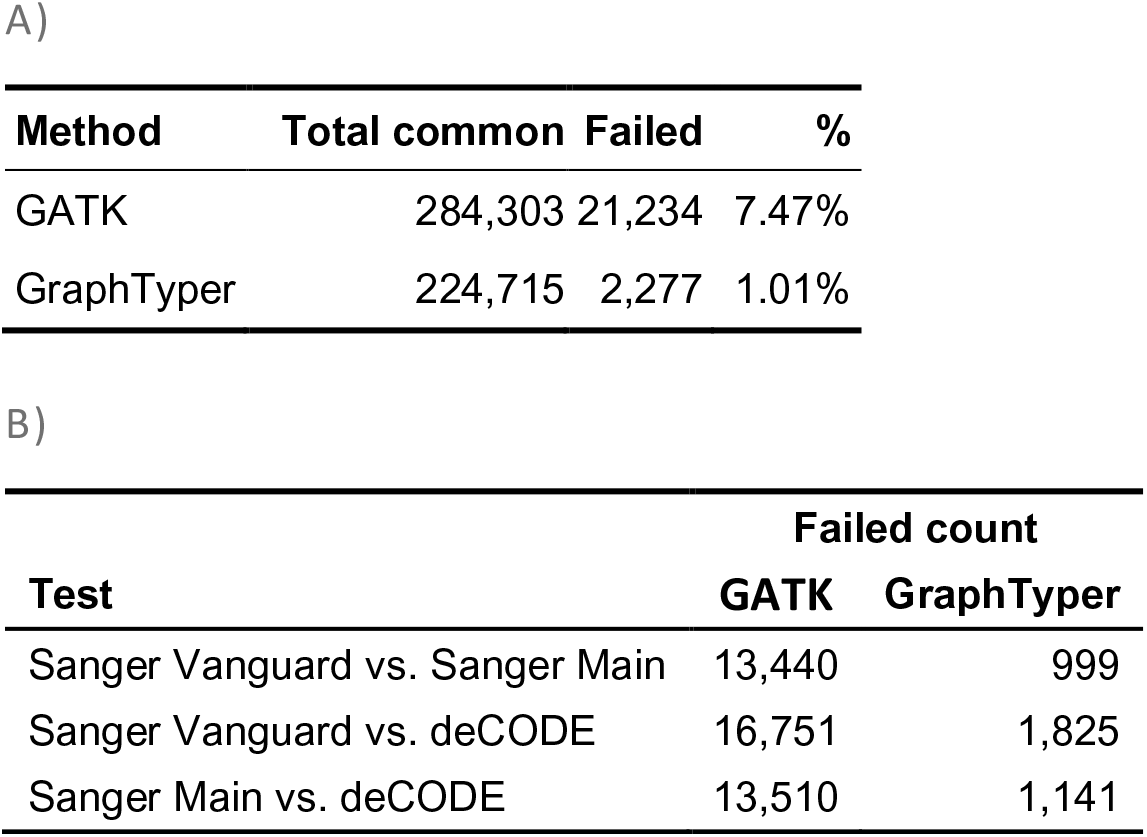
Number of common variants (frequency > .1%) that showed significant association with sequencing center in the 500 random regions test set., A) Total number of variants that failed in any test. B) Number of failed variants stratified by sequencing protocol. Variant is considered “Failed” if p-value < 1e-6, Fisher’s exact test.

**Table S20.**
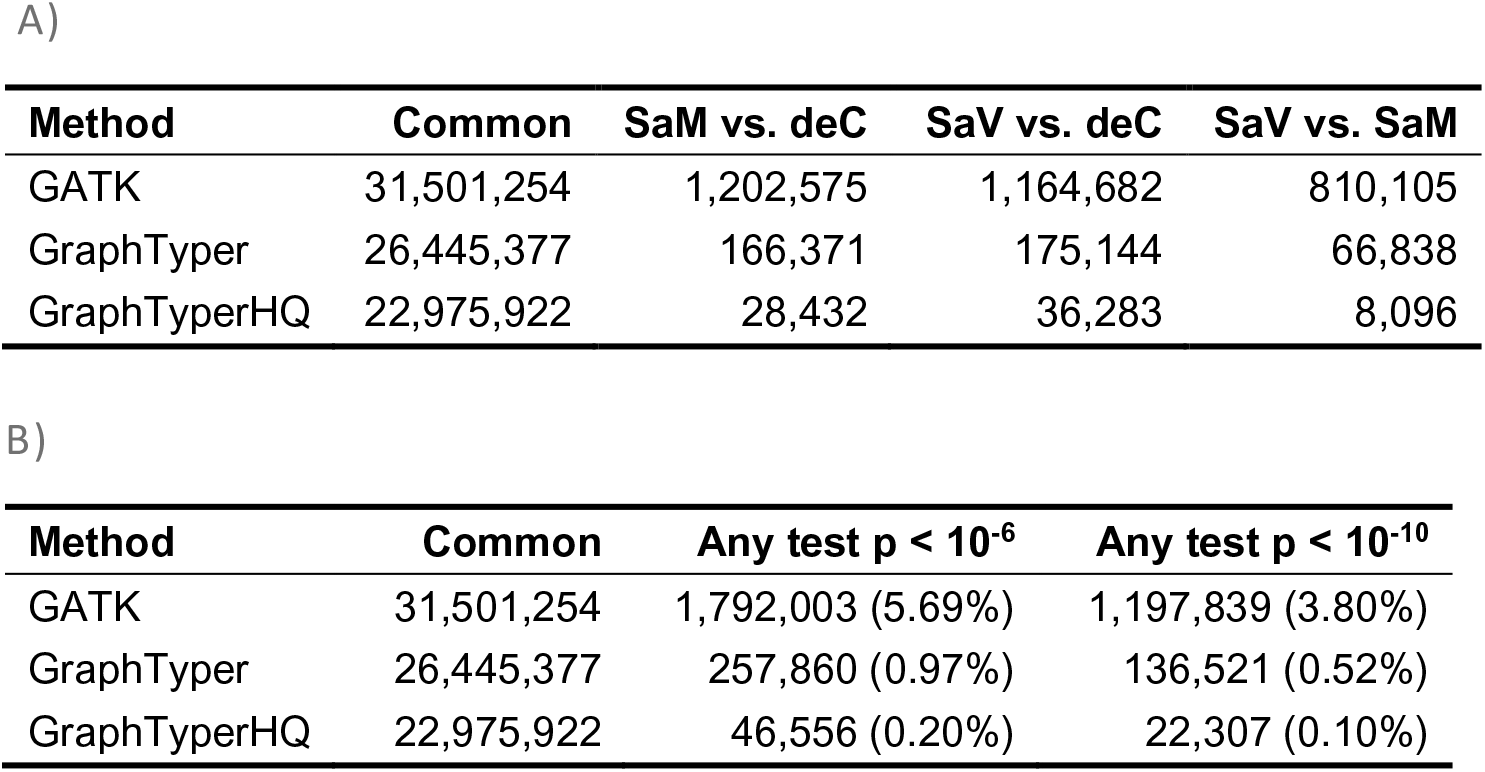
Number of common variants (frequency > 0.1%) that show significant association to sequencing center, indicating batch effects, using a Fisher’s exact test, for common (> 0.1% frequency) variants. A) Number of failed variants stratified by test using p < 10^-6^. deC = samples sequenced at deCODE genetics. SaV = samples sequenced using the Sanger Vanguard processing pipeline. SaM = samples sequenced using the Sanger main phase pipeline. B) Total number of variants that failed in any test, using both p < 10^-6^ and p < 10^-10^.

**Table S21.**
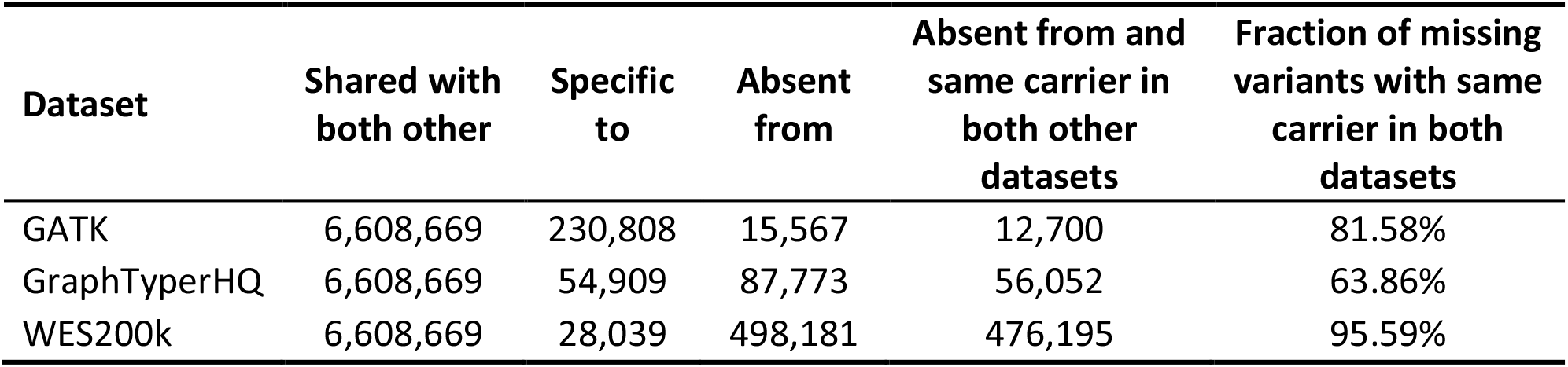
Three-way comparison between the GraphTyperHQ, GATK and WES200k^76^ call analyzed inside WES capture regions within the set of 109,618 individuals present in both the WES200k call set an our set of 150,119 individuals.

**Table S22.**
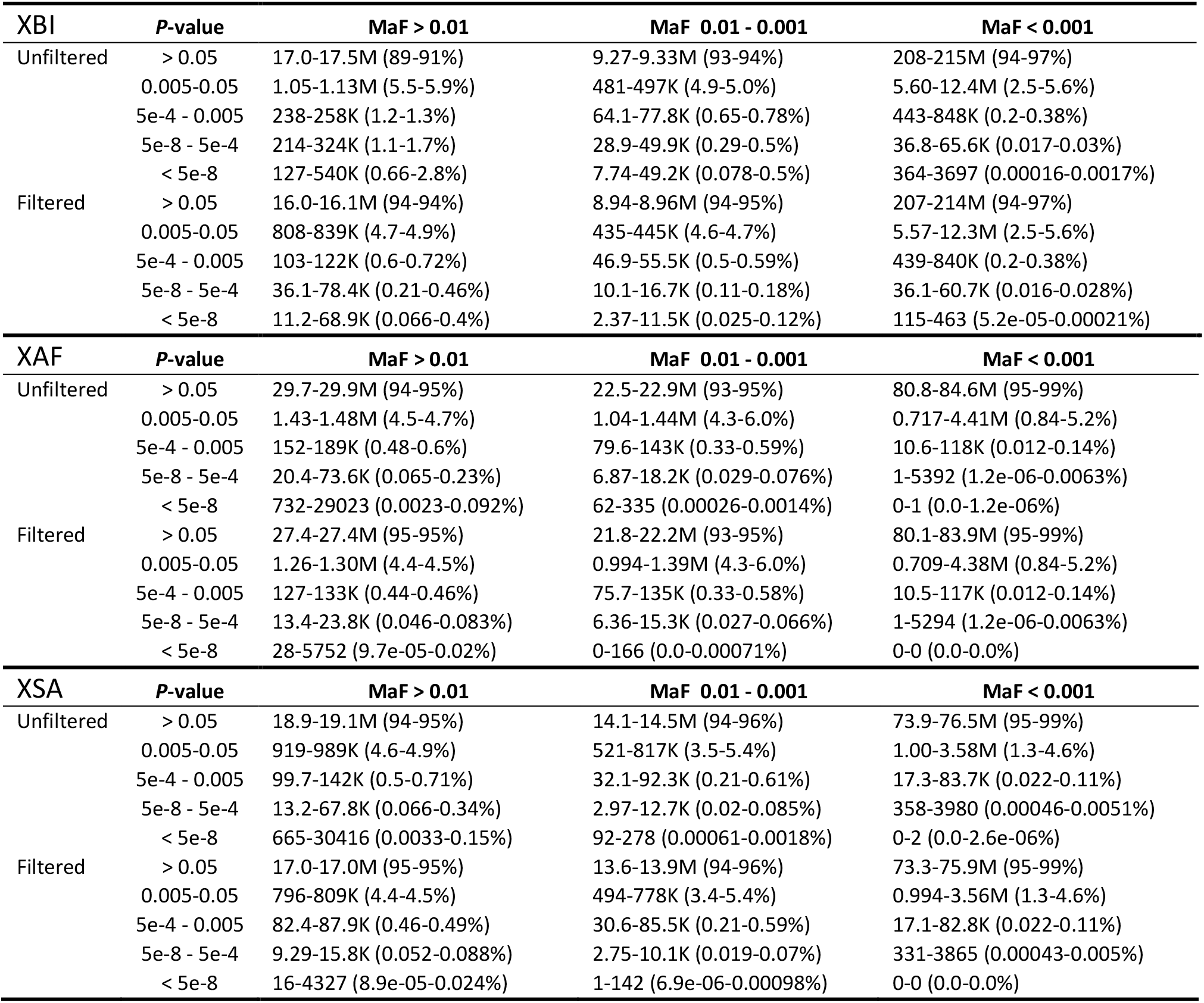
Batch effects for sequencing center in the raw genotype calls. Six phenotypes for batch effects are tested. Results are conditioned on marker minor allele frequency (MAF). Table shows the minimum and maximum number and fraction of markers, across the six phenotypes) with p-value in each p-value range. E.g., when considering the unfiltered dataset and the XSA cohort, MAF > 0.01, between 919 and 989k markers have p-value between 0.005 and 0.05, corresponding to 4.6-4.9% of markers with MAF > 0.01.

**Table S23.**
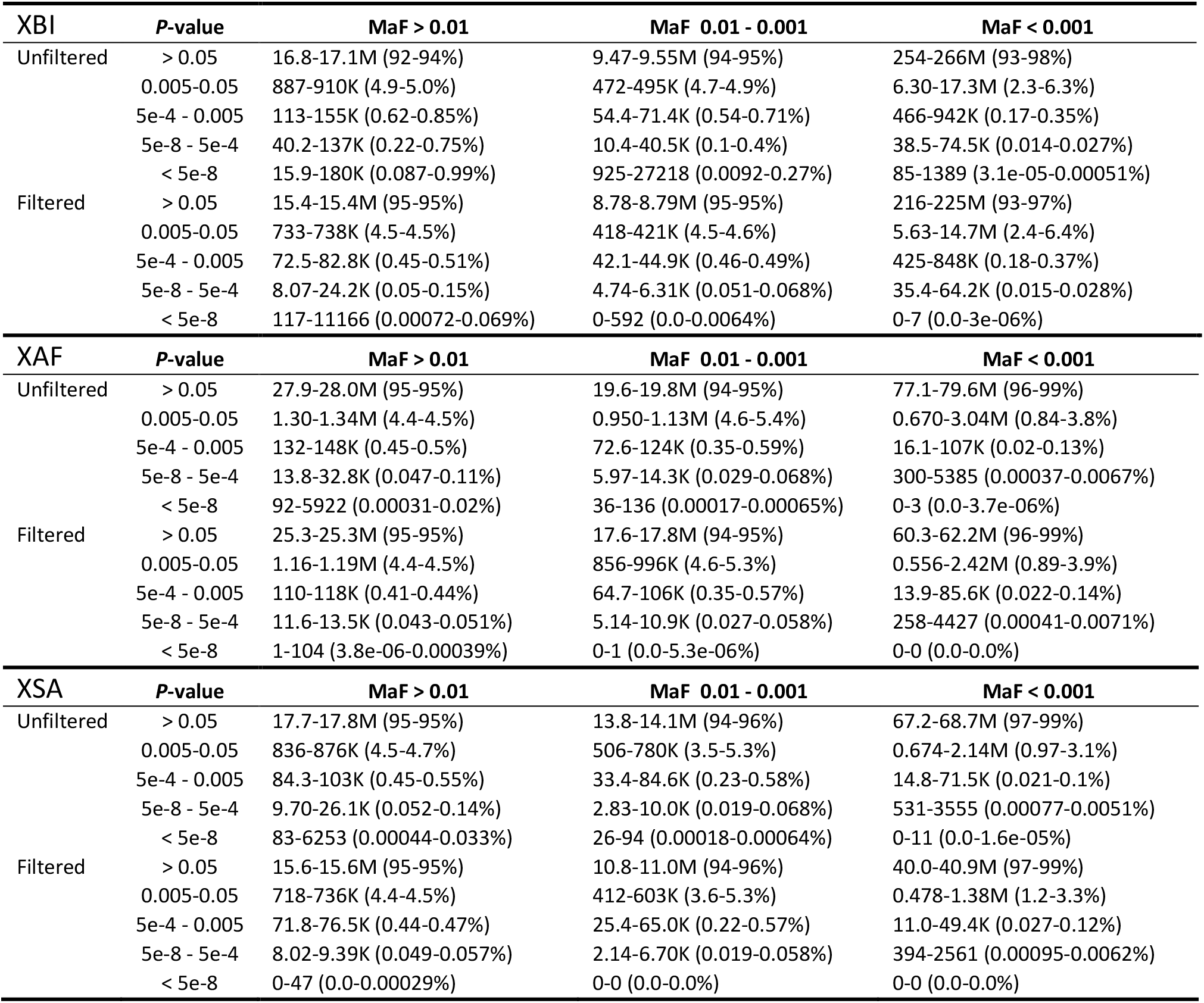
Batch effects for sequencing center in the imputed genotype calls. Six phenotypes for batch effects are tested.Results are conditioned on marker minor allele frequency (MAF). Table shows the minimum and maximum number and fraction of markers, across the six phenotypes) with p-value in each p-value range. E.g., when considering the unfiltered dataset and the XSA cohort, MAF > 0.01, between 836 and 876k markers have p-value between 0.005 and 0.05, corresponding to 4.5-4.7% of markers with MAF > 0.01.

**Table S24.**
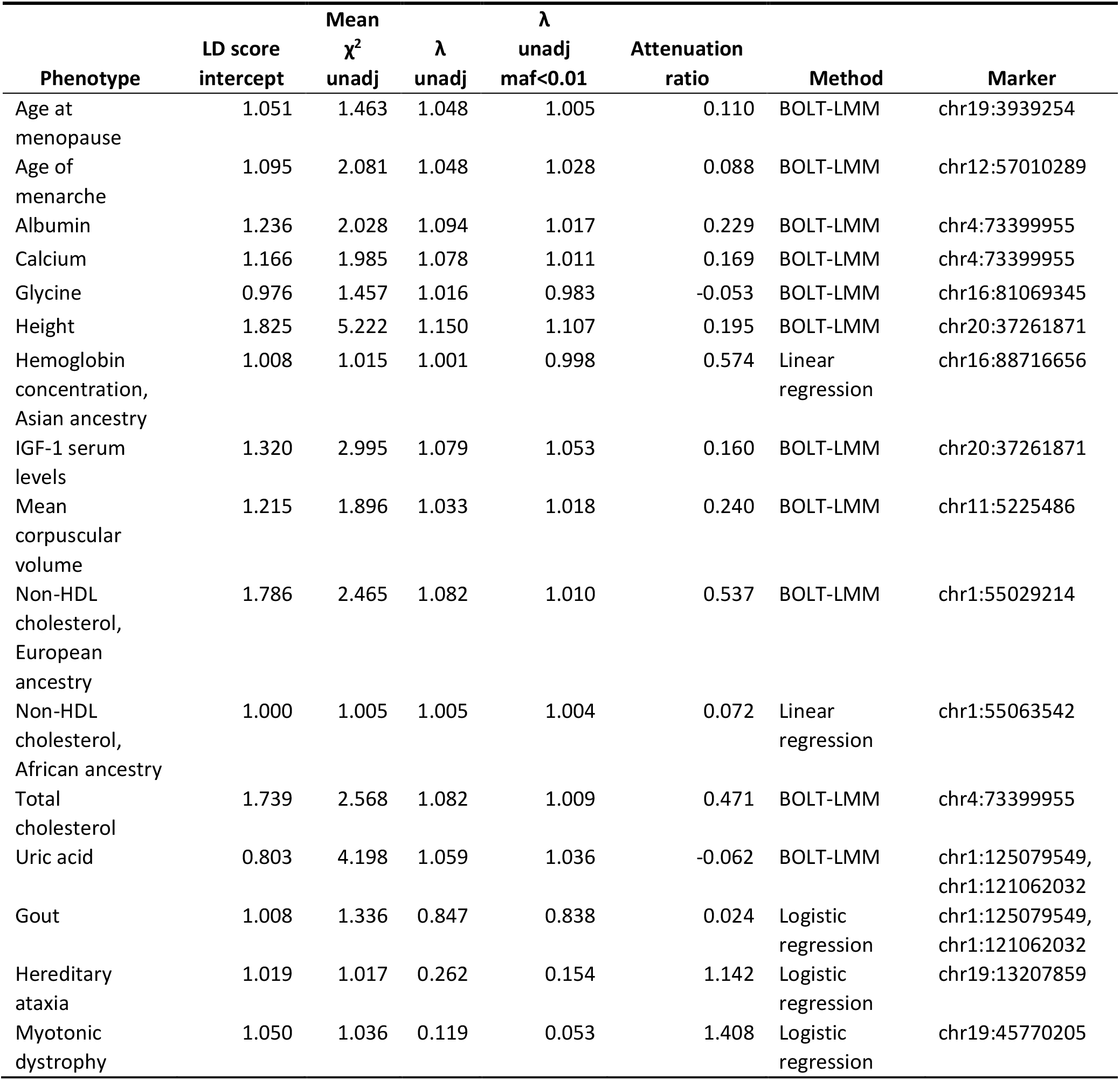
Correction factors and inflation metrics from phenotypes used in this study; LD score intercept, mean chi-squared unadjusted value, unadjusted lambda value, unadjusted lambda value for rare (< 1% MAF) markers and attenuation ratio. Marker represents the ID of the association reported.

**Table S25.**
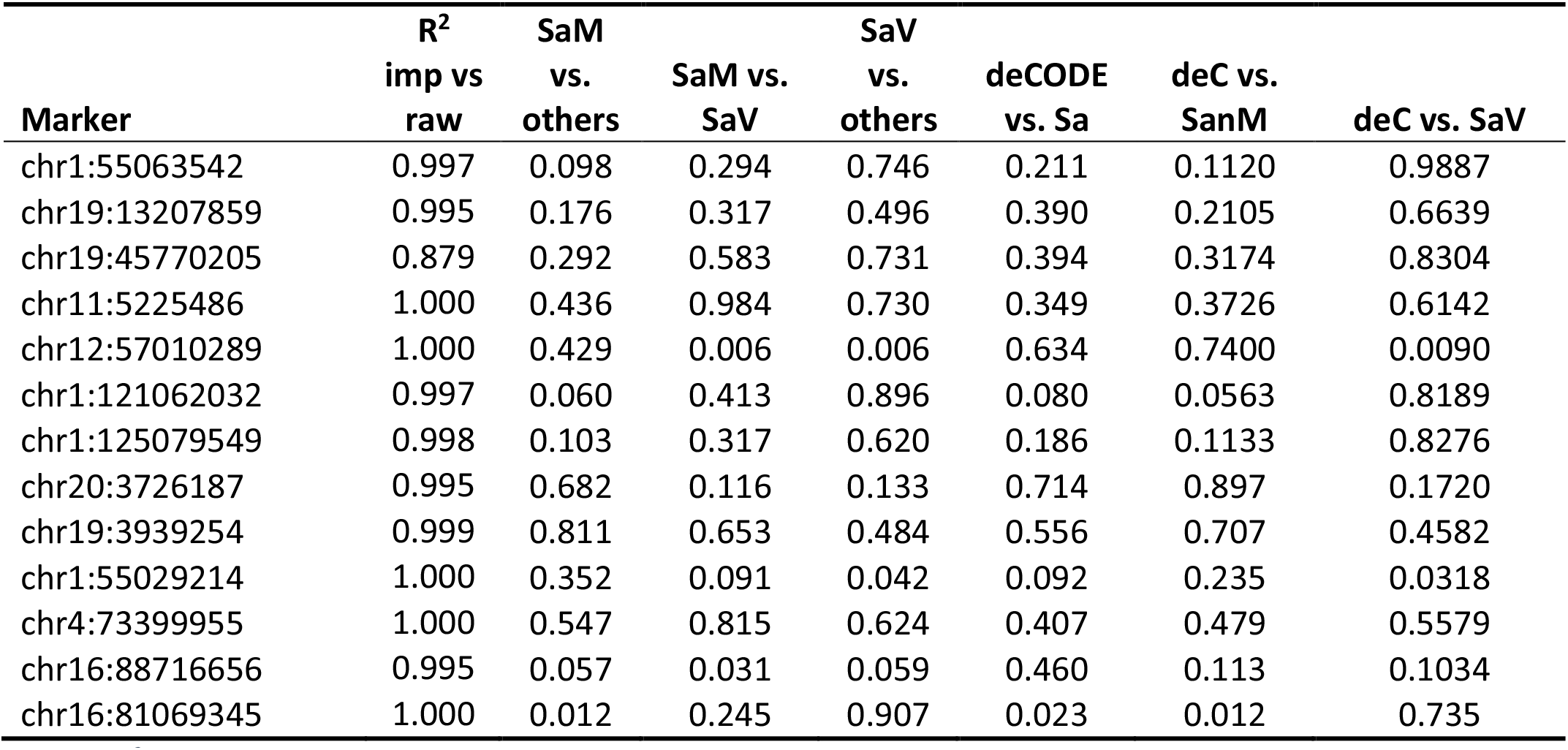
r^2^ between raw genotypes and imputed markers in the XBI cohort. p-value for batch effect in the XBI cohort for markers presented in this study. deC = samples sequenced at deCODE genetics. SaV = samples sequenced using the Sanger Vanguard processing pipeline. SaM = samples sequenced using the Sanger main phase pipeline. Sa = samples sequenced at Sanger. Relationship between marker IDs and phenotypes can be seen in Table S24.

## Methods

### Datasets

#### UKB data

The UKB phenotype and genotype data were collected following an informed consent obtained from all participants. The North West Research Ethics Committee reviewed and approved UKB’s scientific protocol and operational procedures (REC Reference Number: 06/MRE08/65). Data for this study were obtained and research conducted under the UKB applications license numbers 24898 and 68574.

Phenotypes were downloaded from the UKB, and we provide information corresponding to how we processed the resources and created phenotype lists with reference to the field identity available in the UKB data showcase (Table S15Table S15).

#### Icelandic data

The gout sample set^78^, a total of 1740 Icelanders, was recruited through multiple sources. A subset of these individuals were regular users of anti-gout medication corresponding to the Anatomical Therapeutic Chemical Classification System class M04 (ATC-M04). Individuals using ATC-M04 were identified through questionnaires at the time of entry into genetics projects at deCODE and provided by the Directorate of Heahth from entry in the Prescription Medicines Register (2005-2020) or the Register of RAI Assessments and Minimum Data Set (MDS) for residents and applicants of nursing homes (1993–2018). Furthermore, about half had received a clinical diagnosis of gout (International Classification of Disease: ICD-9 code 274 or ICD-10 code M10) between 1984 and 2019 at Landspitali, the National University Hospital of Iceland or at two rheumatology clinics, or such a diagnosis was determined by examining RAI and MDS medical records.

Serum uric acid levels in blood samples from 95,086 Icelanders were obtained from Landspitali, the National University Hospital of Iceland and the Icelandic Medical Center (Laeknasetrid) Laboratory in Mjodd (RAM) between 1990 and 2020. Serum uric acid levels were normalized to a standard normal distribution using quantile-quantile normalization and then adjusted for sex, year of birth and age at measurement. For individuals for whom more than one measurement was available, we used the average of the normalized value. Serum uric acid levels are determined from an enzymatic reaction in which uricase oxidizes urate to allantoin and hydrogen peroxide, which with the aid of peroxidase and a dye forms a colored complex that can be measured in a photometer at a wavelength of 670 nm.

All participating individuals who donated blood signed informed consent. The identities of participants were encrypted using a third-party system approved and monitored by the Icelandic Data Protection Authority. The study was approved by was approved by the National Bioethics Committee of Iceland (Approval no. VSN-15-023) following evaluation of the Icelandic Data Protection Authority. All data processing complies with the instructions of the Data Protection Authority (PV_2017060950ÞS).

RNA sequence data analysis was approved by the Icelandic Data Protection Authority and the National Bioethics Committee of Iceland (no. VSNb2015030021).

#### Danish data

Data was provided from the Danish Blood Donor Study (DBDS)^79^. The DBDS genetic study has been approved by the Danish National Committee on Health Research Ethics (NVK-1700407) and by the Danish Capital Region Data Protection Office (P-2019-99).

### WGS data quality specification

Sequencing was performed at the two sequencing providers, deCODE genetics and the Wellcome Trust Sanger Institute, according to the specifications set forth in the material transfer agreement for UKB Access application nr. 52293 – Summarized as follows:

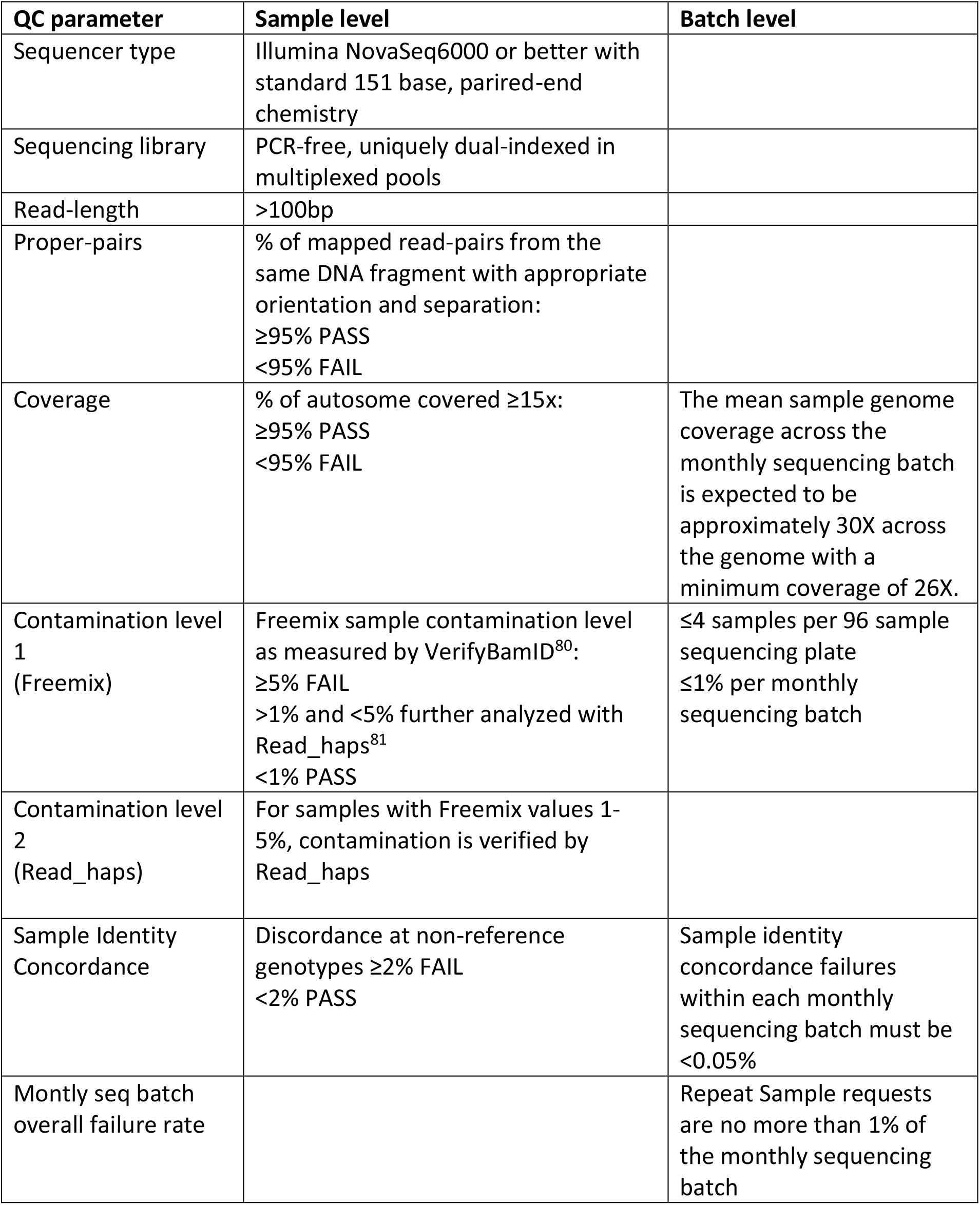

All calculations of data quantity (yield) and coverage must exclude duplicate reads, adaptors, overlapping bases from reads from the same fragment, soft-clipped bases

### Whole genome sequencing

DNA samples were selected by UK Biobank using its picking algorithm which ensures pseudo-randomisation of recruitment centres and collection times across batches, to avoid potential batch effects and shipped on dry-ice to the sequencing centers at Welcome Sanger Institute in Cambridgeshire, UK (WSI) and deCODE genetics in Reykjavik, Iceland (deCODE). The samples were in 70 µL aliquots in Fluid-X 0.3 mL, externally threaded 2D barcoded tubes in 96-well racks with linear barcodes (Brooks Life Sciences) at a normalized, target DNA concentration of 12 ng/µL in 1x TE buffer (10 mM Tris-HCl, 1.0mM EDTA, pH 8.0). Upon arrival, samples/plates were registered in the respective Laboratory Information Management System (LIMS) and stored until use at -20 °C. DNA concentration was confirmed by UV/VIS spectrophotometry (Trinean DropSense system or equivalent). Sequencing libraries were prepared using the NEBNext Ultra™ II PCR-free kit (New England Biolabs). In short, 500 ng of genomic DNA was fragmented to a mean target size of 450-500 bp using high frequency Adaptive Focused Acoustics Technology (AFA) from Covaris Inc (LE220plus instruments and 96-well TPX-AFA plates). End repair and A-tailing was performed in a single step followed by ligation of unique dual indexed sequencing adaptors (IDT for Illumina) and two rounds of SPRI-bead purification (0.6X) using an automatic 96/8-channel liquid handler (Hamilton Microlab STAR and Tecan Freedom EVO). Quality (concentration and insert size) of sequencing libraries was determined using the LabChip GX (96-samples) instrument (Perkin Elmer). Sequencing libraries were pooled appropriately using automatic 8-channel liquid handlers and sequenced using Illuminás NovaSeq6000 instruments. Paired-end sequencing on the S4 flowcell (v1.0 chemistry) was performed with a read length of 2×151 cycles of incorporation and imaging, in addition to 2*8 index cycles to a mean coverage of at least 26X per sample. Real-time analysis (RTA) involved conversion of image data to base-calling in real-time. All steps in the workflow were monitored using the in-LIMS with barcode tracking of all samples/plates and reagents.

### Sequence processing pipeline

The deCODE pipeline (Fig. S16, Fig. S17) for UKB consists of the following steps. An automated pipeline monitors the data coming off the sequencers and starts processing the data when the sequence run folder is ready. The steps taken are:

1. bcl2fastq is run on the sequencer run folder to demultiplex the data and convert each (lane,index) combination into fastq pairs. A checksum is generated for each fastq pair and stored for future reference. The reads in the fastq files are counted and compared against the expected counts coming from the sequencer. The Undetermined read files are inspected, looking for reads that haven’t been accounted for.
2. Each pair of fastq files is processed to create a CRAM file. The steps are

a. Align against GRCh38
b. Fix mate pair information
c. Mark duplicates.
d. Sort in genomic order
e. calculate checksum and compare with fastq checksum. Failure if they don’t match and process is rerun
3. CRAM file is compared with chip genotypes for same sample. Result reported back to the lab. Failure if mismatch rate >2% (potential sample error)
4. QC stats are collected and thresholds applied (Fig. S18). Results are reported back to the lab and CRAM is failed if it doesn’t pass all quality parameter thresholds. Failed lanes are archived and not used in further processing.
5. A merge process monitors the (lane,index) data and merges the data when it is likely that sufficient data have been collected for a sample. The merge process injects all the necessary header information into the file making it ready for export to UKB.
6. When the file has been created, a checksum is generated for each read group and compared with the corresponding checksums for the fastq files. Failure if the don’t match and the merge process is rerun.
7. The merged CRAM file is archived and the upstream data are marked for deletion.
8. Variant calling is performed on the CRAM file and the result is prepared for export to UKB. This includes the production of the BQSR25 table as well as a gVCF file.
9. QC stats for the merged file are collected and thresholds applied. Results are reported back to the lab.

a. If the file fails on quantity only, the file is held, the lab initiates a top-up run which is processed as described above and upon completion is merged with the held CRAM file into a new merged CRAM file. That new merged CRAM file is then processed again as described above
b. If the file fails on other quality parameters, the file is failed and the sample is flagged in the lab. The lab must decide the appropriate action (abandon sample, request a new library)
10. The merged CRAM file, along with variant calling and auxiliary data are sent to UK Biobank

### Pipeline details

#### Alignment

Each read group is aligned to GRCh38 reference (GRCh38 reference with alt contigs plus additional decoy contigs and HLA genes) with bwa mem (v0.7.17)^23^ using parameters ‘-K 100000000 -Y -t 24’. To add MC and MQ tags, samblaster^82^ (v0.1.24) is used with parameters ‘-a --addMateTags’. Duplicates are marked using Picard MarkDuplicates (v2.20.3) with parameters “ASSUME_SORT_ORDER=queryname READ_NAME_REGEX=’[a-zA- Z0-9-]+:[0-9]+:[a-zA-Z0-9]+:[0-9]:([0-9]+):([0-9]+):([0-9]+)’”, then the results are coordinate sorted using samtools^83^ (v1.9).

#### Merging

Internal thresholds are set for total sequence yield and read count, GC fraction (first and second read in pair) and bias compared to reference, flagging of base conversions in sample preparation, where certain trinucleotides are more commonly observed in sequencing than their reverse complement, flagging of base conversions in sample preparation, where certain trinucleotides are more commonly observed in sequencing than their reverse complement, percentage aligned library read pairs, library insert fragment size distribution, sequencing adapter contamination level, sequence run base call quality values, genotype concordance rate against supplied genome-wide genotype data supplied by UKB for each participant sample, sequence error rate, sequence contamination rate and genome coverage. Read group bam files are assessed for these parameters and those that pass all the thresholds are merged using samtools^83^ merge (v1.9) and converted to CRAM format.

#### Single sample variant calling

A base quality recalibration table is created using GATK BaseRecalibrator (v4.0.12) with known sites files dbSNP138, Mills and 1000G gold standard indels, and known indels from GATK resource bundle and parameters “--preserve-qscores-less-than 6 -L chr1 .. -L chr22”. For each chromosome in chr1 .. chr22, chrX, chrY, the resulting base recalibration table is applied using GATK ApplyBQSR (v4.0.12) with parameters “--preserve-qscores-less-than 6 --static-quantized-quals 10 --static-quantized-quals 20 --static-quantized-quals 30 -- create-output-bam-index” and then variants are called using GATK^25^ HaplotypeCaller (v4.0.12) with parameters “-ERC GVCF”. The resulting 24 chromosome g.vcf files are then combined using Picard^25^ MergeVcfs (v2.20.3).

#### Quality assessment reports

Reports (Table S16) to assess the data quality are created using the following programs (in the steps Lane QC, QCPreview and QCStats):

- BamQC (v1.0.0) run on each lane before merge (Table S17).
- samtools^83^ stats (v1.9) using parameters “-d -p”, i.e. excluding duplicates and overlapping basepairs
- Picard CollectWGSMetrics (v2.20.3) is run with parameters “USE_FAST_ALGORITHM=True MINIMUM_BASE_QUALITY=0 MINIMUM_MAPPING_QUALITY=0 COVERAGE_CAP=1000” once for whole genome, once for autosomes only
- Genotypes are called from .g.vcf files using GATK GenotypeGVCFs (v4.0.12)
- Sample contamination is assessed by running verifyBamId^80^ (v1.1.3) with parameters “--ignoreRG --chip-none --free-full --maxDepth 100 --precise” using 1000G phase 3 autosomal SNPs with European MAF > 0.01
- Sample contamination is accessed again using read_haps^81^ “-q 30 -mq 30 -c 1 -w 1000”
- Genetic sex is determined using a set of some 100 000 chrX SNPs from gnomad with Non-Finnish European MAF > 0.2. For each variant, the genotype is called using GATK GenotypeGVCFs. Then the ratio of observed to expected heterozygosity assuming diploidy is computed. If ratio > 0.7 the sample is called female, if ratio < 0.3 the sample is called male, otherwise undetermined. Implemented using in-house script gvcf_sexcheck.py
- Picard^25^ Genotypeconcordance (v2.20.3) is run with parameter “MIN_GQ=30” to determine concordance with genotypes for quality variants from a chip array.

### Sequence coverage

Our design was to have at least 95% of the genome covered to at least 15x coverage in each sample. Nearly half of the variants detected in this study are singletons, detected in only one sample and a large majority of the variants are rare. GraphTyper requires that at least 4 high quality reads be observed at position for a marker to be called. At 15x coverage the probability that a variant observed in a single individual would be misclassified due to random sampling is 3.5%. Sequence coverage across the genome computed over 1,000 randomly selected samples can be seen in Fig. S19.

### SNP and indel calling with GraphTyper

Prior to running GraphTyper we preprocessed all input CRAI indices by extracting a large single file containing all CRAI index entries with sample_id for a 50kb window (with 1 kb padding at each side of the region) for all samples. For each region, we then created a chopped CRAI for each sample by processing the large file for the corresponding region, substantially reducing the amount of CRAI index entries read.

Further, we created a sequence cache of the reference FASTA file using the ‘seq_cache_populate.pl’ script distributed with samtools 1.9. In each region we copied the corresponding sequence cache to the local disk and used it for reading the CRAM files by setting the ‘REF_CACHÈ environment variable.

We ran GraphTyper (v2.7.1) using the ‘genotypè subcommand. The full command we ran was in the format:

~~~
graphtyper genotype ${UKBIO_REFERENCE}
 --sams=${SAMS}
 --sams_index=${CRAI_TMP}/crai_filelist.txt
 --avg_cov_by_readlen=${COVERAGES}
 --region=${REGION}
 --threads=${THREADS}
 --verbose
~~~

Where `UKBIO_REFERENCE` is the `GRCh38_full_analysis_set_plus_decoy_hla` FASTA sequence file, `SAMS` is a list of all input BAM/CRAM files, `CRAI_TMP` is a path to the chopped CRAI files on the local disk, `COVERAGES` is the coverage divided by the read length for each input file, `REGION` is the genotyping region and `THREADS` is the number of threads to use.

#### Running time

All jobs were run using 12 cors with 60GB of reserved RAM. Approximately 1% of jobs were rerun using 24 cores with 120GB reserved RAM. A few jobs requiring more cores and memory, with a single job finishing with 48 cores and 1000GB of RAM. Total reserved CPU time on cluster was 5.8M CPU hours and total effective compute time 5.0M CPU hours. The difference in these numbers is explained by the fact that not all cores reserved for the program may not utilize all at the same time.

### SNP and indel calling with Calling with GATK

We used GATK versions 4.1.7.0 for all regions. Regions that failed were rerun with version 4.1.8.1.

The process starts by slicing the 50kb region (padded with 1kb) of every sample file with tabix (from htslib^83^ version 1.9) onto local disk and then builds a GenomicsDB with GATK GenomicsDBImport. The command we ran was the following:

~~~
gatk --java-options “-Xmx${JAVAMEM_TOTAL}G
 -Xms${JAVAMEM_TOTAL}G
 -DGATK_STACKTRACE_ON_USER_EXCEPTION=true”
    GenomicsDBImport
    --genomicsdb-workspace-path ${GDB}
    --intervals ${REGION_PADDED}
    --tmp-dir ${GDB_TMP}
    --sample-name-map ${SNMAP}
    --batch-size ${BATCH_SIZE}
    --reader-threads ${RTHREADS}
~~~

where `SNMAP` is the tab-delimited text file of sample names and paths to samples. The parameters `--batch-size` and `--reader-threads` are used to reduce memory usage. We then split the padded region into as many smaller regions as the number of threads, and pad those regions again with 1kb. The GenotypeGVCFs command was then ran wrapped in GNU parallel

~~~
parallel --halt=now,fail=1
 --jobs=${NTHREADS}
 --xapply
    ”${GATK_WITH_OPTS} GenotypeGVCFs
    --genomicsdb-use-vcf-codec
      -R ${REF}
      -V gendb://${GDB}
      --tmp-dir=${tmpdir}
      -L {1}
      -O {2} &&
    ${GATK_WITH_OPTS} SelectVariants -R ${REF}
    -V {2}
    -L {3}
    -O {4}”
    :::: ${REGIONS_PADDED} ${SPLITFILES_PADDED} ${REGIONS} ${SPLITFILES}
~~~

where `REF` is the reference, `REGIONS_PADDED` is a file containing the padded subregions, SPLITFILES_PADDED is a file containing the intermediate padded output file paths, `REGIONS` is a file containing the subregions and `SPLITFILES` is a file containing the intermediate output file paths after selecting the variants.

We then run the following command to combine the intermediate output files

~~~
gatk --java-options “-Djava.io.tmpdir=$tmpdir
 -Xmx${JAVAMEM_TOTAL}G
 -Xms${JAVAMEM_TOTAL}G
 GatherVcfs -R ${REF}
  -O ${OUT}
  --arguments_file ${VARARGS}
~~~

where `VARARGS` is a file containing arguments for all input intermediate vcfs.

It should be noted that running GATK out of the box will cause every job to read the entire gVCF index file (.tbi) for each of the 150,119 samples. The average size of the index files is 4.15MB, so each job would have to read 4.15*150,126 = 623GB of data on top of the actual gVCF slice data. For 60,000 jobs, this would amount to 623GB*60,000 = 37PB or 25.2GB/sec of additional read overhead if the jobs are run on 20,000 cores in 17 days. This read overhead will definitely prevent 20,000 cores from being used simultaneously. However, this problem was avoided by pre-processing the .tbi files and modifying the software reading the gVCF files from the central storage in a similar fashion as we did for GraphTyper and the CRAM index files (.crai).

All jobs were run initially with 6 cores and 100GB of RAM. Jobs that failed due to memory were rerun with more memory, up to a maximum of 1,458GB. Calling for 320 of the 50kb regions failed using GATK version 4.1.7.0, either due to 1,458GB of memory being insuffient or program failure. These regions were split into 3,066 5kb regions (regions at the end of chromosomes were smaller than 50kb) and rerun with GATK version 4.1.8.1. 320 regions, representing 1.6Mb, of the 3,066 regions again failed calling with GATK version 4.1.8.1. No further attempt was made to call these regions. Total reserved CPU time on cluster was 9.6M CPU hours and total effective compute time 4.0M CPU hours. The difference in these numbers is explained by the fact that while 6 cores reserved for the program it may not utilize all at the same time.

### Evaluation of SNP and indel callers across 500 random regions

Prior to running variant calling on the whole dataset, we evaluated joint variant callers for the UKB sequencing effort. We evaluated the quality of the genotype calls and feasibility of variant calling 150,000 or more WGS samples. There were some minor differences between this call set and the final set, for example we included seven Genome in a Bottle (GIAB) samples for evaluation purposes in the evaluation set. However, we believe these differences should have minimal effects on the results.

#### Input data

The evaluation was run on the set of 150,126 WGS samples including 7 WGS samples obtained from the GIAB Consortium (websites).

All of the GIAB BAM files were down sampled to approximately 30x coverage using `samtools view -s 42.FRAC` option with seed 42 and FRAC was the fraction of reads to keep such that 30x was obtained to represent more closely the target coverage of the other input files. Samtools version 1.9 was used.

We evaluated 500 regions (50kb each). We selected the regions at random by listing all such regions (only excluding regions which contained only Ns) and using the first 500 regions from the output of `sort -R`.

#### SNP and indel calling with GraphTyper

We ran GraphTyper as described for the whole dataset, with the additional option `-- normal_and_no_variant_overlapping`. This was done to simplify the comparison to the GIAB truth sets using the files which contained no variant overlaps as `rtg vcfeval` sometimes misinterprets overlapping variants. This option however should normally be omitted to generate only a set where variants may overlap. We used the non-overlapping set when comparing to the GIAB truth sets but in all other analysis of GraphTyper variants we used the “normal” variants set.

#### Resource Requirements

##### GraphTyper

The GraphTyper jobs were run on 12 cores and 60GB of memory reserved for each job (5GB/core). Average CPU time was 82 hours and average elapsed walltime was 7.8 hours, resulting in average reserved core time (walltime*12) of 93.6 hours. For 150k samples and the entire genome (60,000 50kb slices), this translates to overall compute time of 93.6*60,000 = 5.62M hours, or 12 days if the jobs are run in parallel on 20,000 cores. The input data to GraphTyper are CRAM files. The average size of an input CRAM file is 17.8GB, so the total size of data to be read is 17.8GB*150,126 = 2.7PB. Reading those data once over a period of 12 days was estimated to result in average sustained read rate of 2.6GB/sec, assuming no overhead.

##### GATK HaplotypeCaller

The GATK jobs were run on 6 cores and 80GB of memory reserved for each job (13.33GB/core). With these settings, 488 of the 500 jobs completed. The 12 remaining jobs finished when given more memory. The average cpu time was 53.4 hours and average elapsed walltime was 22.5 hours, resulting in average reserved core time (walltime*6) of 135.1 hours. For 150k samples and the entire genome (60,000 50kb slices), this translates to overall compute time of 135*60,000 = 8.1M hours, or 17 days if the jobs are run in parallel on 20,000 cores.

#### Output sizes

Both programs return a gzip compressed vcf file (.vcf.gz), one for each region. The average file size for GATK is 12.0GB while for GraphTyper it is 7.6GB. For 150k samples and the entire genome, this translates to a total estimated output size of 12GB*60,000 = 720TB for GATK, while the output for GraphTyper was 7.6GB*60,000 = 445TB. This difference in size may in part be explained by the fact that GATK reports more variants and in part by the fact that GATK does not cap genotype likelihoods at 255 like GraphTyper, thus resulting in worse compression ratio.

#### Comparison to the GIAB truth sets

In both sets we genotyped seven GIAB samples. We extracted the calls made in each of those sample in the 150k sample run and compared to their v3.3.2 truth set in high confidence regions. Variant callers do not generally have the same output when genotyping a single sample compared to extracting the sample from a multi-sample run. We ran the tool RTG-vcfeval^84^ to make the comparison to the truth set in the high confidence regions which overlapped the 500 regions. For all of the samples, GraphTyper had both higher sensitivity and precision than GATK on the full sets (Table S1). The difference between the two callers was small (99.44% vs. 99.34%,Table S1) for SNPs but more marked for indels (97.58% vs. 94.14%, Table S1), were both methods performed much worse on indels only compared to single sample calling, indicating that indel calling is particularly difficult when genotyping a large population.

#### Overview of genotyping results

We analyzed the evaluation set to further learn the differences between the two genotyping datasets. In this analysis, all of the variants from the VCF were analyzed on per alternative allele basis. Therefore the number of variants we report here is higher than the number of VCF records due to multi-allelic variants.

#### Variant counts

We counted the number of variants in each dataset (Table S18, Fig. S20). We saw that there were more variants in the GATK dataset. However, GATK also had greater number of missing calls (genotype quality = 0 in the VCF). It is expected that the ratio of SNP transitions to transversion is roughly 2.1-2.3 in humans genome-wide. We saw lower ratios in the call sets, but it was higher in the GraphTyper set (1.639) than in the GATK set (1.507). Indel sizes were limited to 100 bp in the GraphTyper dataset but had a larger range in the GATK set (Fig. S21).

#### Batch Effect by Sequence Center

Further, we investigated how many common variants had genotype calls which were highly correlated to the sequence center for which the sample was sequenced in. As the batches had a highly different amount of samples we randomly selected 10,000 samples from each batch and restricted our analysis to those sample. We tested whether there were more alternative calls (either ref/alt or alt/alt calls) compared to the number of reference calls in each set using Fisher’s exact test. Only common variants were tested, as we expect fewer rare markers to be rejected due to smaller sample size. We used a p-value threshold of 10^−6^, any variants with a lower p-value in any of three tests were considered as failed.

To our surprise, we saw that a large fraction of the common variants are highly correlated with the sequence center (Table S19), on average of 7.47% and 1.01% of variants for GATK and GraphTyper, respectively.

#### Singletons variants

Fig. S22a) shows the distribution of singletons by mutation classes between and the variant allele frequency (VAF) of singletons. A VAF of 50% is expected for singletons.

#### Parent-Offspring Trio Analysis

There were 28 parent-offspring trios in the dataset. We analyzed Mendelian errors in the trios as well as the rate of transmission of alternative alleles from parent to offspring. We assume that the alleles transmit from parent to child with equal likelihood and use the transmission rate to estimate false discovery rate and number of germline variants in the datasets. More info on the method is described^24^.

#### Mendelian Errors

We measured non-reference Mendelian errors by checking for Mendelian consistency when a parent had an alternative genotype (ref/alt or alt/alt) (Table S3).

#### Estimating FDR and number of TP in trios

Using transmission rate in trios we estimate both false discovery rate (FDR) and the number of true positive (TP) variants^24^. We also stratified the results by variant type. We estimated that GraphTyper finds slighlty more true positive variants across all variant types with a much lower false discovery rate than GATK (Table S3). GATK finds more true positive SNPs, but GraphTyper more true positive indels.

#### Monozygotic Twin Non-Ref Error Rate

There were 14 pairs of monozygotic twins in the dataset. We checked how many of the non-reference variants were consistent between a pair of monozygotic twins. We considered a variant to be non-ref if either twin had an alternative allele in their genotyped. GraphTyper had lower error rate between monozygotic twins (Table S3C).

#### Summary

Overall, we find that GraphTyper performs consistently slightly better than GATK in the variant quality experiments. Despite that GATK reports more variants than GraphTyper, we estimate that GraphTyper’s sensitivity is better in both the GIAB truth set comparison and family trio analysis. There appears to be larger gap between the methods in terms of noise, GATK performs worse in precision in the GIAB comparison, in the family trios we estimated that GATK’s false discovery rate is twice as much as GraphTyper’s, and 7-fold more common GATK variants failed the batch effect test compared to GraphTyper.

### Comparison of final GraphTyper and GATK call sets

In addition to the two callsets, we also define the set “GraphTyperHQ” as the set of GraphTyper alternative alleles with AAScore above 0.5.

#### Variant counts and frequency classes

We counted total number of variants in the sets (Table S7). When counting the number of “variants” in any context hereafter, we are referring to alternative alleles excluding the alleles that are denoted as ‘*’ in the VCF.

An informative call is one with non-zero quality (GQ > 0). We saw that GATK had more variants but also much more missing calls. We split the sets into three frequency classes: Common (Allele frequency (AF) > 0.1%), rare (AF <0.1%, excluding singletons) and singletons (one called carrier in the set). A vast majority of the datasets (95.6% - 96.0%) are have an allele frequency below 0.1%. Singletons account for nearly half of the variants (43.9-45.5%) (Table S7).

The transition transversion ratio was 1.550, 1.642 and 1.657 for the GATK, GraphTyper and GraphTyperHQ datasets, respectively (Table S7B, Fig. S23).

#### Batch effect by sequence center

We investigated how many common variants had genotype calls which were highly correlated to the sequence center, i.e. the location which the sample was sequenced at. We randomly selected 10,000 samples from each sequencing center analysis pipeline and restricted our analysis to those samples. We tested whether there were more alternative calls (either ref/alt or alt/alt calls) compared to the number of reference calls in each set using Fisher’s exact test. Only common variants were tested, as we expect rare variants are less likely to be rejected due to limited sample size. The same variant often fails multiple tests, 5.69%, 0.97% and 0.20% of common variants associate with sequencing center for the GATK, GraphTyper and GraphTyperHQ datasets, respectively (Table S20).

#### Variant transmission in parent-offspring trios and monozygotic twin pairs

There were 28 parent-offspring trios in the dataset. We analyzed the rate of transmission of alternative alleles from parent to offspring. We assume that the alleles transmit from parent to child with equal likelihood and use the transmission rate to estimate false discovery rate (FDR) and number of germline true positive (TP) variants in the datasets^24^. From the family trios we estimate that GraphTyper has more true positive variants while also having lower rate of false positive ones. GraphTyperHQ has considerably lower false discovery rate than the GATK call set (Table S2). There were 14 pairs of monozygotic twins in the dataset. We checked how many inconsistent genotypes in the twins were on average in a 1MB region (ICPM). We also calculate the total non-reference consistency rate among, by checking for consistency among all calls where either twin had a call with an alternative allele. The raw GATK and GraphTyper datasets have many inconsistent calls between monozygotic twins but the filtered GraphTyper dataset is much more consistent (Table S2).

### Batch effects in final dataset

Sequencing was performed in three batches; individuals sequenced at deCODE genetics (deCODE), sequenced at the Welcome Trust Sanger Institute processed using Vanguard phase pipeline (Sanger Vanguard), sequenced at the Welcome Trust Sanger Instititute using the main phase pipeline (Sanger Main). From the lists of individuals, we constructed six different phenotypes, comparing each sequencing batch both to the two other sequencing batches both jointly and separately. Association tests were performed per cohort and both for the raw genotypes and the imputed dataset, following the protocol describe in subsection “Association testing”. Association results are presented for both a filtered and an unfiltered dataset. For the raw genotypes the filtered set refers to markers with AAscore > 0.5, or the GraphTyper HQ set. For the imputed genotypes the filtered set refers to markers markers with AAscore > 0.5 and Imp info > 0.8.

Batch effects for sequencing center are shown in Table S22 for raw genotypes and in Table S23 for imputed genotypes, with results conditioned on frequency and association p-value. Considerable batch effects can be observed in all datasets. As expected, lower levels of batch effects were detected for the filtered dataset. More common variants show higher levels of batch effects. We note that marker batch effect is conflated with missing data in genotype calling.

For the purpose of the Table S22 and Table S23 frequency is computed from genotype likelihoods, where the likelihoods are transformed into probabilities that the individual is a carrier. In this way an individuals with no sequence reads is assigned frequency 50%, upweighing rare markers where a large fraction of markers have missing data. Alternatively frequencies can be computed from the carrier status of individuals without missing data.

### Overlap with UKBB WES SNPs

#### Comparison based on minor allele frequency

A recent UKB WES dataset has 200,000 individuals (WES200k^76^). In the dataset there are 1,047,397 SNPs with WES AF >0.01% and 353,889 with WES AF >0.1%. We checked how many of those were not found in the WGS datasets. 1.81, 0.44 and 1.60% of variants with frequency > 0.01% in the WES200k dataset were missing the in the GATK, GraphTyper and GraphTyperHQ datasets, respectively (Table S5).

#### Variant normalization

To reliably compare two datasets (the result of different samples, technologies or tools), the data needs to be in a standardized format. The commonly used VCF format is unfortunately very ambiguous:

a. Two variation events may be represented as a single multi-allelic VCF record in one set or as two VCF records in another.
b. A single variation event has many equivalent representations, i.e. variants are not required to be left-aligned and parsimonious^85^.
c. While records are required to be ordered by POS, two records with the same POS have no defined order. This makes line-wise comparisons and merges difficult. In particular, the order generated by bcftools norm is not alphabetical.
d. Different conventions exist for how to name chromosomes (”Chr1” vs “1”; “ChrX” vs “Chr23” vs “23”).
e. IDs are absent from some files, making it more difficult to return to the original entry after changes have happened.

Our normalization pipeline employs bcftools norm to split multi-allelic variants and to left-align and trim them. It enforces a naming convention for the chromosomes (”Chr1” … “ChrX”) and adds an ID-String if missing. Finally, the data is split into 50KB regions and sorted by “Chrom,Pos,Ref,Alt”. Since normalization may influence the POS field of a VCF record, it may fall into a different 50KB bin than before; these cases are handled.

Once all datasets are normalized, a merged dataset is created from them. This consists of one set of VCF files where all INFO fields from the original datasets are included with a set-specific prefix, e.g. “GATK_AF” instead of “AF”. The original datasets’ ID, QUAL and FILTER fields are also included in the merged files’ INFO fields as “GATK_ID”, “GATK_QUAL” etc. This representation of the data is sparse because missing entries do not take up space. For analysis purposes, a TSV or GOR[Z] file can be created for individual regions or full chromosomes. The transformation from .VCF.GZ files to .GORZ and further operations (e.g. JOINs) are efficiently possible, because our VCF records are already fully sorted.

#### Comparison of WES and WGS call sets on the same sets of samples

In an attempt to make a judicial comparison between WES and WGS as well as between the GraphTyperHQ and GATK call sets we analyzed seperately the calls made for a subset of 109,618 individuals included in our dataset as well as the 200k release of WES data from the UKB^76^.

Variants not present in any of the 109,618 indivdiuals were removed from analysis, resulting in 558,128,486 GraphTyperHQ variants and 13,815,704 WES variants. We then split the variants by functional annotation and tabulated the number of variants shared between the two call sets and the number of variants absent from the other call set (Table 1).

To further explore the accuracy of genotype callers we analyzed specifically variants inside regions that are purportedly captured by exome sequencing (websites,Table S21), 6,608,669 variants are found in all three call sets. Variants in one call set and not another may be either true or false positives. A priori, we would expect that variants found in two call sets to be a strong indication of the variant being a true positive. This analysis is complicated by the fact that although we have filtered the set of GraphTyper variants GATK variants have not been filtered for true positives.

A total of 87,773 variants are found by both GATK and WES but missed by GraphTyperHQ. 32,875 of these variants were present in the unfilterd GraphTyper dataset but filtered due to low AAscore. 56,909 out of the 87,773 variants have the same primary carrier in both datasets, while the remaining 30,864 are found by both callers but not in the same sample. These variants represent a shared tendancy of false positive calls at the same variant (but in different samples) across both datasets. Best practices use of GATK recommends filtering of variants based on a number of factors. While we have not computed all of these, we computed for these variants what we believe are some of the most common causes of failure; failing variants that have variant allele frequency (VAF) below 25%, failing variants that are not supported by reads from both strands and failing variant that are not supported by both a read that is first in pair and one that is second in pair. Applying these three filters removed 69.3% of the 56,909 variants, suggesting at most a small fraction of the variants found by both GATK and WES, but not GraphTyper, are in fact called reliably enough to be used in a recommended genetic analysis.

Cursory analysis of the variants found by both GraphTyper and WES, but not GATK suggested that these were similarly possibly problematic.

Analysis of variants found by both GATK and GraphTyper however suggested that these were in large part true positives. We considered the distribution of the 898,764 singletons shared between the callers and found their distribution (XAF 78,229 (8.70%), XBI 564,346 (62.79%), XSA 71,823 (8.00%), OTH 184,366 (20.51%)), to be similar to that of the distribution of singleton calls overall (XAF 746,289 (8.40%), XBI 5,731,044 (64.50%), XSA 707,379 (7.96%), OTH 1,701,318 (19.15%)). We would expect false positive calls due to sequencing artifacts would be similar to the fraction of individuals from each cohort in our intersected sequencing set (XAF 2.05%, XBI 87.89%, XSA 2.08%, OTH 7.99%).

### SV calling with Manta and GraphTyper

We ran a structural variant (SV) genotyping pipeline similar to the one we had previously applied to 49,962 Icelanders^60^. In summary, we ran Manta^58^ v1.6 to discover SVs on all 150,119 individuals in the genotyping set. We also created a set of highly confident common SVs (imputation info above 0.95 with frequency above 0.1%) from our previous studies using both Illumina short reads^60^ and Oxford Nanopore long-read data^59^. Finally, we inferred a set of SVs from six publicly available assembly datasets using dipcall^86^, as described previously^60^. We used svimmer^60^ to merge these different SV datasets and we called the resulting SVs using GraphTyper^60^ version 2.7.1. By incorporating data from long read data and high quality assemblies, we are able call more true SVs compared using short reads only, particularly for common SVs.

A total of 895,054 variants were called, of which 637,321 variants were annoted as “Pass”. Variant counts are presented for variants annoted by GraphTyper as “Pass”, unless otherwise noted.

The majority of the SVs are deletions (81.3%), however we observe only slightly more deletions than insertions and duplications on average per individual (Fig. 3a). This is because the source for many insertions are long reads and assembly data, and thus many rare insertions are missing. Deletions are typically easier to discover in short read data. Individuals that belong in the XAF cohort carry more SVs than in the other cohorts (Fig. 3b).

### Microsatellite calling with popSTR

We followed the protocol described above for Graphtyper before we ran PopSTR(v2.0) and created chopped CRAI indices for all samples as well as a reference sequence cache for each processed region.

We scanned all CRAM files in 50kb regions using the popSTR subcommand computeReadAttributes.

The format of the command was:

~~~
popSTR computeReadAttributes ${CRAI_TMP}/sampleList.txt ${RESULT_TMP} markerList flanking <(readLength-2*flanking) “.” longRepeats N
~~~

Results over a predetermined set of microsatellites from chr21(our kernel) were used to estimate a slippage rate for each individual using the popSTR subcommand computePnSlippageDefault.

The format of the command was:

~~~
popSTR computePnSlippageDefault
–PL $sample
–AD ${RESULT_TMP}/attributes/chr21/
-OF ${outDir}/pnSlippage
-FP $sampleIDx
-MS ${codeDir}/kernelSlippageRates
-MD ${codeDir}/kernel/kernelModels
~~~

Combining CRAM analysis results and sample slippage rates we performed genomewide genotyping using the popSTR subcommand msGenotyperDefault

The format of the command was:

~~~
popSTR msGenotyperDefault –ADCN ${RESULT_TMP}/attributes/${chrom}/ -PNS pnSlippage –MS ${RESULT_TMP}/markerSlipps/${chrom}/markerSlippage –VD
${RESULT_TMP} –VN vcfName –ML markerList –I $idx –FP 1
~~~

`CRAI_TMP` is a path to the chopped CRAI files on the local disk, `RESULT_TMP` is a folder on the local disk to store results, `flanking` is a parameter specifying the number of bps required to anchor a read to the microsatellite, `readLength` is the length of reads in the CRAM file, `markerList` is a list of all microsatellites in the 50kb region being analysed, `outDir` is a directory to store sample slippage results, `sampleIDx` is the index of the sample being analysed in the `sampleList.txt`, `codeDir` is the directory where popSTR and its dependencies are stored and `$idx` is the index of the region being analyzed.

#### Filtering of microsatellites

We recommend the following best practice filtering guidelines.

Filter marker where:

average coverage < 10 or average coverage > 75

command: `bcftools query -f`

~~~
‘%CHROM\t%POS\t%INFO/nReads\t%INFO/nPnsWithReads\n‘ $file |
awk ‘{print $1,$2,$3/$4}‘ | awk ‘{if ($3>10 && $3<75){print
$1\t$2}}‘ > pass; bcftools view -T pass -o filtered_${file} -O z $file; tabix filtered_${file}
~~~

average genotype quality < 20

command: bcftools query -f ‘%CHROM\t%POS[\t%GT\t%GQ]\n‘ $file | awk ‘{sum=0; miss=0; avail=0; for (i=4;i<=NF;i+=2){if ($(i- 1)==“./.”){miss+=1}else{sum+=$i; avail+=1}} if(avail>0){mean=sum/avail}else{mean=0} print $1,$2,mean}‘ | awk ‘{if ($3>20){print $1\t$2}}‘ > pass; bcftools view -T pass -o filtered_${file} -O z $file; tabix filtered_${file}

number of individuals with reads < 75,000 command: `bcftools query -f`

~~~
‘%CHROM\t%POS\t%INFO/nPnsWithReads\n‘ $file |awk ‘{if ($3>75000){print $1\t$2}}‘ > pass; bcftools view -T pass -o filtered_${file} -O z $file; tabix filtered_${file}
~~~

number of reads not supporting genotype/number of reads available > 0.3 command: `bcftools query –f`

~~~
‘%CHROM\t%POS\t%INFO/nNonSupportReads\t%INFO/nReads\n‘ $file | awk ‘{if ($3/$4<0.3){print $1\t$2}}‘ > pass; bcftools view -T pass -o filtered_${file} -O z $file; tabix filtered_${file}
~~~

A total of 2,393,292 variants pass these filters.

### Imputation and phasing

The UKB samples were SNP chip genotyped with a custom-made Affymetrix chip, UK BiLEVE Axiom in the first 50,000 individuals^87^, and the Affymetrix UKB Axiom array^88^ in the remaining participants. We used the existing long-range phasing of the SNP chip genotyped samples^5^.

We excluced SNP and indel sequence variants where at least 50% of the samples had no coverage (GQ score = 0), if the Hardy Weinberg p-value was less than 10^-30^ or if heterozygous excess was less than 0.5 or greater than 1.5.

We used the remaining sequence variants and the long-range phased chip data to create a haplotype reference panel using inhouse tools^1, 89^. We then imputed the haplotype reference panel variants into the chip genotyped samples using inhouse tools and methods described previously^1, 89^.

The imputation consists of estimating, for each haplotype, haplotype sharing with haplotypes in the haplotype reference panel, giving haplotype weights for each haplotype. These weights along with allele probabilities for each haplotype in the haplotype reference panel allow imputation with a Li and Stephens^90^ model similar to the one used in IMPUTE2^91^. Estimation of haplotype weights was based on long-range phased chip haplotypes.

Sequence variant phasing consists of iteratively imputing the phase in each sequenced sample based on the other sequenced samples and the estimated phase from last iteration. The imputed genotypes, along with the original genotypes are weighted together to estimate new allele probabilites for the haplotypes. Imputation is done as described above.

We compute a leave-one-out r-squared score (L1oR2) as the squared correlation (r^2^ value) of the original genotype calls with the genotypes imputed for each sample when excluding the original genotype of the sample from the imputation input.

#### Imputation results

We refer to a variant as being reliably imputed if its L1oR2 score is greater than 0.5 and imputation info^1^ was above 0.8.

Imputation and phasing accuracy of SNPs and indels for the GraphTyperHQ set is shown in (Fig. 2, Fig. S14, Table S11). GraphTyperHQ filters variants based on an AAscore of 0.5.

Requiring higher AAscore allows a higher fraction of variants to be imputed (Fig. S24). We found that variants located > 100kb from a chip genotyped variant and variants in regions that were placed on different chromosomes on GRCh38^22^ and CHM13^71^ imputed less accurately than others.

SVs and microsatellites are imputed less accurately than SNPs and indels (Fig. S14), in part due to difficulty in genotyping those variants. For microsatellites, this may in part be attributed to the high mutation rate of microsatellites and in part to the fact that the results are presented for the unfiltered microsatellite set, we expect that a higher fraction of microsatellites would impute after filtering.

#### Comparison of imputation from GATK and GraphTyper variants

We imputed all variants genotyped by GATK and GraphTyper across chr22, 10-11Mb. We define a variant to be imputed if the phasing leave-one-out r2^1^ (L1or2) was at least 0.5 and imputation info^1^ was at least 0.5. We present the number of variants that could be imputed as a function of frequency and variant type (Table S4). Although more variants are called by GATK, there are more variants called by GraphTyper that can be imputed, across all frequency classes and variant types.

### Genome annotation

We downloaded Refseq and Ensembl gene map annotations from Ensembl^92^, version 100 database. The gene maps were transformed to segments with each position in GRCh38 annotated as at least one of 3’utr, 5’utr, coding, downstream, intergenic, intronic, spliceregion, splicesite, upstream.

These regions were grouped and ordered by precedence:

1. coding – coding
2. splice – spliceregion, splicesite
3. 5‘UTR – 5‘UTR
4. 3‘UTR – 3‘UTR
5. proximal – upstream, downstream, intronic
6. – intergenic – intergenic

Each position was then given annotation according to its lowest precedence rank annotation, e.g. a position annotated as both spliceregion and 5‘UTR was given the annotation “splice”.

### Identification of functionally important regions

To identify functionally important regions, we start by estimating whether reliable basecalls can be expected to be made at each site in the genome. The sequence coverage at each bp in GRCh38 was computed for each of 1,000 randomly selected individuals. At each bp we then computed the mean and s.d. of coverage across the 1,000 individuals. Bps with mean coverage at least 20 and s.d. of coverage at most 12 were considered reliable bps. Only variants in GraphTyperHQ (AAscore > 0.5) were considered in the analysis.

#### Recurrent mutations, and spectra under saturation

Using the classification of SNP variants from above, we calculate the ratio of all SNP’s incGraphTyperHQ that falls into each category. Then we do the same restricting to singletons, i.e. calculate the proportion of singletons falling into each mutation class. For comparison, we calculate the fractions of each SNP class in all 181,258 SNP’s from a curated list of 194,687 de novo mutations in 2,976 Icelandic trios^29^. We use this distribution on mutation classes to calculate the transitons/tranversions ratio in each case.

To get a list of recurrent mutations, we join this list of de novo mutations with GraphTyperHQ. This overlap is almost certainly cases of the same alleles originating from separate mutation events.

#### Saturation for general mutation classes

We restrict our analysis to the reliable bps described above and group bps and their complement and consider each A or T base in the genome as a mutation opportunity for T>A, T>C or T>G mutations. Similarly, we consider each G or C base as potential C>A, C>G or C>T mutation, splitting C>T into two classes based on whether they occur in a CpG context or not. We then compute the saturation ratio as the number of observed mutations in GraphTyperHQ divided by the number of mutation opportunities at reliable bps. Computation is done separately for the autosomes and chromosome X. 95% CIs are computed using a normal approximation to the binomial distribution, treating each site as an independent observation.

#### Sites methylated in the germline

We determine sites on GRCh38 that are methylated in the germline using ENCODE Whole Genome Bisulfite Sequencing^10^ (WGBS) data from samples of human testes and ovaries. More precisely we use sample ENCFF946UQB and ENCFF157ZPP for testes and ENCFF561KYJ, ENCFF545XYI and ENCFF515OOQ for ovaries.

We assume that methylation is strand symmetric and compute methylation ratio for each CpG dinucleotide in a given tissue type by tabulating the number of reads supporting methylation or non-methylation in each dinucleotide, summing over all samples of a given tissue type and then compute the fraction of reads that support methylation.

We consider a site in a CpG dinucleotide on the reference genome methylated in the germline if its methylation ratio is at least 0.7 in both testes and ovaries, and the combined depth is at least 20 for testes and 30 for ovaries, or 10 times the number of samples in each tissue type. This resulted in a list of 17,902,255 CpG dinucleotides, harboring 35,804,510 CpG>TpG mutation opportunities.

#### Saturation at methylated CpG sites

For each potential CpG>TpG at a methylated site we assessed its most significant potential consequence with Variant Effect Predictor^93^ v. 100. In case of multiple such consequences we chose the alphabetically last one. We also classified them based on the functional classifications described above. For each class we estimated the saturation as the ratio of variants of that functional class in GraphTyperHQ divided by the number of mutation opportunities. 95% CIs are computed using a normal approximation to the binomial distribution, treating each site as an independent observation.

#### Depletion rank (DR)

We followed a methodology akin to^35^. A variant depletion score is computed for an overlapping set of 500 bp windows in the genome with 50bp step size. A total of 49,104,026 500 bp windows where at least 450 bp were considered reliable bps were considered for further analysis. We tallied the number of occurrences of each possible heptamer (H) and the number of times the central bp in the heptamer was observed as a SNP (S), across the first set of non-overlapping windows. To account for regional mutational patterns in the genome^94^, we dichotomized the genome into two mutually exclusive subsets, inside and outside of C>G enriched regions (Supplementary Table 12 in^94^). The ratio S/H was then interpreted as the expected mutation rate of the heptamer, separately for each of the two subsets. For each window we then computed the observed number of variants (O) and then subtracted its expected number of variants (E), given its heptamers. This difference was divided by the square root of the expected value ((O-E)/ √E). We exclued windows from the analysis where the average AAscore was lower than 0.85 for variants within the window. These ((O-E)/ √E) numbers were then sorted and the window with the i-th lowest depletion score was assigned a Depletion Rank of 100(i-0.5)/n, where n is the total number of windows.

To compute DR restricted to the cohorts, we applied the same approach restricting to sequence variants that are present in each of the XBI, XSA and XAF cohorts.

### WGS individuals carrying actionable genotypes meeting ACMG criteria

The American College of Medical Genetics and Genomics (ACMG) recommends reporting secondary findings in a list of actionable genes associated with diseases that are highly penetrant and for which a well-established intervention is available^27^. The initial version (ACMG SF v1.0) was published in 2013 and included 56 actionable genes but has since been updated twice to ACMG SF v2.0 and v3.0 listing 59 and 73 actionable genes, respectively. 2.0% of the 49,960 WES individuals from the UKB were reported^28^ to carry an actionable variant in at least one gene from the ACMG v2.0 list of 59 genes. Using their criteria, we detected actionable genotypes in 2.6% of 150.119 WGS individuals. When applying the same criteria to the ACMG v3.0 gene list (73 genes), the fraction of individuals carrying an actionable genotype increases to 3.5%. In the ACMG v3.0 list of actionable genes, HFE p.Cys282Tyr homozygotes are recommended to be reported, but does not fullfill the previously described criteria^28^. In the set of 150,119 WGS individuals, we observe 929 HFE p.Cys282Tyr homozygotes (0.62%), thereby increasing the fraction of individuals carrying an actionable genotype in one of the ACMG v3.0 genes to 4.1%.

### Genotype count of rare LoF variants

We counted the number of autosomal heterozygous and homozygous genotypes per individual for rare LoF variants (minor allele frequency (MAF)<1% in all 3 groups, XBI, XAF and XSA). LoF variants are those annotated by the Variant Effect predictor as having consequence as one of: stop gained, frameshift, splice acceptor, splice donor og start loss. Heterozygous counts were based on WGS data, and homozygous counts were based on phased genotypes.

### GWAS enrichment analysis

We have previously described a likelihood-based inference model for estimating the enrichment of trait-associating sequence variants on the basis of their annotations^39^. Similar to our earlier work^39^ we defined a set of 22.8M high-quality sequence variants identified as mono-allelic SNPs or Indels in a set 28,075 whole genome sequenced individuals from the Icelandic population.

The high-quality SNP-indels (22.8M) were then tested for association to a selected set of 614 human diseases and other traits. For each trait, we split the genome into 10Mb windows and selected the strongest sequence variant association from each window where p < 1·10^-9^. Then, for each chromosome, we sorted the selected sequence variants according to P-value to then determine whether the second best variant still associates at p <1·10^-9^ after adjusting the trait for the strongest variant on that same chromosome. If so, this second best sequence variant was incorporated into a final set of “independently associated” variants for that trait, and the process continued for all other sequence variants down the list –each time adjusting for “stronger” variants on the same chromosome. This yielded a set of 3,431 independently associated sequence variants in 322 traits. For each of the 3,431 trait-associated variants, we searched for correlated sequence variants (r2>0.80) in the same Icelandic population. In this way, a given trait association variant along with its correlated variants (found in linkage disequilibrium; LD) defines an association signal. P-values were estimated by determining how often the enrichment estimate (E) is above or below E=1 by bootstrapping (N=5000) of the GWAS association signals. We then annotated sequence variants according to whether or not they are found within regions that show low and high DR scores (1st percentile versus 99th percentile; i.e. most and least conserved regions, respectively); refered to as DR-1% and DR-99%, respectively. In this model, we specified eleven other annotations of sequence variants: loss of function, missense, splice-donor/acceptor, splice region, synonymous, 5kb gene-upstream, 5kb gene-downstream, 3’UTR, 5’UTR, intronic and the remaining sequence variants as “other” (not found in any of the specified annotation categories). Similarly, we specified another model wherein we estimated enrichment for DR-5% and DR-95%.

### Overlap with ENCODE regions

We used annotations from ENCODE^10^ and compute the odds ratios these annotations in regions of different DR scores. We label each bp in the genome with a_11_,a_12_,a_21_ or a_22_, where the first number represent that the bp was annoted with the given ENCODE annotation (1) or not (2) and the second number represents that the DR score was above (1) or below (2) a given threshold.

The odds ratio for the ENCODE annotation given the DR score threshold is then: OR=a_11_/a_21_ ×a_22_/a_12_.

The marker label parameters are computed for each one of the annotations on a set of 1Mb windows across the regions annoted with a DR score. The mean odds ratio is computed by summing up the individual parameters for the complete set of windows. We use bootstrapping to estimate the confidence limits for the odds ratio we, for each bootstrap sample we sample with replacement from the complete set of 1Mb windows, sum up individually the resulting set a_ij_‘s and compute the odds ratio for the bootstrap sample. The odds ratio is computed for a total of 1000 bootstrap samples and the confidence intervals defined between the 2.5% and 97.5% quantile of the resulting dataset.

### Association testing

We tested for association with quantitative traits based on the linear mixed model implemented in BOLT-LMM^95^. We used BOLT-LMM to calculate leave-one-chromosome out (LOCO) residuals which we then tested for association using simple linear regression. We used logistic regression to test for the association between sequence variants and binary traits. We tested variants for association under the additive model using the expected allele counts as a covariate for quantitative traits and integrating over the possible genotypes for binary traits. Sequencing status (whether the individual is one of the WGS individuals), other available individual characteristics that correlate with the trait were additionally included in the model; sex, age, and principal components (20 for XBI and XAF, 45 for XSA) in order to adjust for population stratification. Association analyses with XAF and XSA ethnicities have sample sizes <10,000 and therefore were done with linear regression directly instead of BOLT-LMM. The correction factor employed was the intercept of each regression analysis.

We used LD score regression to account for distribution inflation in the dataset due to cryptic relatedness and population stratification^13^. Using 1.1 million variants, we regressed the χ2 statistics from our GWASs against LD score and used the intercepts as a correction factor. Effect sizes based on the LOCO residuals are shrunk and we rescaled them based on the shrinkage of the 1.1 million variants used in the LD score regression. Table S24 lists statistics for the GWAS analysis of each of the association signals presented here.

Manhattan plots, quantile-quantile (QQ) plots and histograms of inverse-normal transformed values after adjustment for covariates age, sex and 40 principal components can be found in Fig. S25 and Fig. S26 for quantitative and binary phenotypes, respectively. Locus plots for Uric Acid and Menarche association can be found in Fig. S27.

All associations reported are for imputed genotypes. For comparison purposes associations were also performed on the genotypes directly. For the association testing perfomed on the directly genotyped markers the same set of covariates were used, apart from sequencing status (as all individuals are sequenced) and additionaly the sequencing center (deCODE, Sanger main, Sanger Vanguard) was used as a covariate. Table S25 shows correlation between the raw and the imputed genotypes and batch effects for sequencing center in the XBI cohort.

An individual was deemed to be a carrier of an allele if the probability that the individual carried the allele was at least 0.9. The association analysis was limited to markers were at least one (XAF, XSA), two (XBI, imputed dataset) or three (XBI, raw genotypes) individuals carried the minor allele. As association tests are frequently limited to a subset of the individuals in the datset the association analysis was further limited to those markers were there was at least one carrier among the individuals in the association test. In the imputed dataset association tests were further limited to those markers with imp info > 0.5 and in the raw genotype set to those markers with sequencing information^1^ > 0.8.

### RNA sequence data

RNA sequencing was performed on samples from cardiac right atrium of 169 Icelanders. The data and subsequent sequence alignment to GRCh38 has been described^96^. To estimate the effect of deletion of exon 6 in transcript ENST00000168977.6 of *NMRK2* we counted fragments aligning from the donor site of exon 5 to either acceptor site of exon 6 or exon 7 (Fig. S12, Fig. S13).

### Defining cohorts

Most studies of UKB data to date have been conducted on a list of around 410,000 “Caucasian” individuals created by UKB on the basis of “White British” self-identification and clustering on genetic principal components derived from microarray genotypes^5^. Like some recent studies^54, 97, 98^, we wished to capitalize on the diversity in the UKB. To achieve this, we defined three cohorts based on the most common ancestries identified among the participants, using a combination of 1) UMAP dimension reduction of 40 genetic principal components provided by UKB, 2) ADMIXTURE analysis supervised on five reference populations and self-reported ethnicity information.

In order to define the three cohorts, we followed previous work^99^ and applied UMAP to the 40 genetic principal components provided by UKB. UMAP was performed in R using umap::umap() using default parameters in v0.2.3, notably n_neighbours 15 and min_dist 0.1 UMAP placed the individuals in a two-dimensional latent space featuring several clusters and filaments. These structures showed a correspondence with self-described ethnicity (Fig. S28).

To provide a separate measure of ancestry that we could use to inform our interpretation of the UMAP clusters, we superimposed results from a supervised ADMIXTURE^100^ analysis of the UKB microarray genotypes (Supp Section ADMIXTURE), using five training populations from the 1000 Genomes Project^8^ (1000GP): CEU (Northern Europeans from Utah), CHB (Han Chinese in Beijing), ITU (Indian Telugu in the UK), PEL (Peruvians in Lima), and YRI (Yoruba in Ibadan, Nigeria). We observed a clear correspondence between UMAP coordinates and ancestry proportions assigned by ADMIXTURE (Fig. S29, Fig. S30). Using this correspondence and guided by self-reported ethnicity information, we defined the cohorts by manually delineating regions in the UMAP latent space that were limited to individuals with British– Irish ancestry (XBI, N=431,805), South Asian ancestry (XSA, N=9,633), and African ancestry (XAF, N=9,252). This left 37,598 individuals with genotype data, who were assigned to an arbitrary cohort we refer to as OTH (short for other). The distribution of ancestry estimated using the ADMIXTURE in each of the four cohorts (Fig. S29). Fig. S6, Fig. S7 and Fig. S8 show the geographical distribution of birthplaces for the XBI, XAF and XSA cohorts, respectively.

The most systematic difference between the XBI cohort and the prevailing UKB-defined “Caucasian” set is our inclusion in XBI of around 12,500 individuals identifying as White Irish. This is clearly justified, given the known geographical and cultural proximity of the populations of the Britain and island of Ireland. More importantly, both our analyses (and those of previous publications) clearly reveal evidence for extensive gene flow between them. Thus, the main Irish genetic cluster appears in PCA as an integrated component of continuous variation in the UK (Fig. S5), and is not clearly separated from others. Another major difference of the XBI cohort relative to the much-used Caucasian set, is the addition of around 10,900 individuals who did not identify as White-British, but we infer to have ancestry indistinguishable from British-Irish individuals. We note that the greater size of the XBI cohort should provide more statistical power to detect genotype-phenotype associations.

### Computing principal components within cohorts

#### Microarray data

For all cohorts, we first removed variants with missingness >3% and 135 individuals with genomewide missingness >5%. We then removed a canonical set of long-range high-LD regions and all indels.

For the XAF and XSA cohorts, the following procedure was followed. We first excluded both individuals from each pair of relatives with kinship coefficient 0.0625 or greater; these excluded individuals were later projected onto the principal components. We then pruned for variants in complete linkage disequilibrium (r2 = 1) using plink --indep-pairwise 200 25 0.999999, and then removed all variants with MAF <1%. PCA for these two cohorts was performed using smartpca^101^ with parameters numoutevec: 45, numoutlieriter: 0, ldregress: 200, and ldposlimit: 100000. We then projected all relatives using the OADP method implemented in bigsnpr’s^102^ function bed_projectSelfPCA().

A slightly different approach was used for the XBI set, due to the very large number of individuals. We first excluded: individuals from each pair of relatives at a kinship coefficient threshold of 0.0442 or greater; individuals with inbreeding of 0.1 or greater; individuals with genomewide missingness 1% or greater; and all remaining individuals defined as “HetMiss” (heterozygosity/missingness) outliers by UKB. We next removed variants with < 0.05% MAF and a Hardy-Weinberg disequilibrium p-value (calculated with plink --hwe midp) of <1e-100. Then LD clumping was performed using bigsnpr’s bed_clumping() function using thr.r2 = 0.2 and [window] size = 500 [kb]. We calculated 30 PCs on the remaining variants and individuals using bigsnpr’s bed_randomSVD(), and the previously excluded individuals were projected onto these PCs using OADP.

#### WGS data

To prepare each WGS cohort for PCA, we first removed all variants with missingness >3%. We then excluded individuals with genomic inbreeding over 0.1 and both individuals in any pair of 3rd degree or closer relatives. The excluded individuals were later projected onto the principal components. After excluding these individuals, we removed all singleton variants. For XBI in particular, we also removed all variants with minor allele count <10, in order to make computation more tractable and to minimise the influence of very recent genealogical structure.

bigsnpr^101^ was used to remove a canonical list of long-range, high-LD regions [long-range LD ref] and then perform LD clumping using bed_clumping() with an r2 threshold of 0.1 and a window size of 5 megabases. We then used bed_randomSVD() in bigsnpr to calculate 50 PCs on each of the cohorts.

The first six principal components in each cohort are shown in Fig. S31, Fig. S32 and Fig. S33.

#### Inbreeding

Genomic inbreeding in the form of F_ROH_ (proportion of the genome in runs of homozygosity) was calculated on microarray genotypes using PLINK^103^ v1.9 and the same parameters specified in ROHgen2^104^: homozyg-window-snp 50; homozyg-snp 50; homozyg-kb 1500; homozyg-gap 1000; homozyg-density 50; homozyg-window-missing 5; homozyg-window-het 1. Genotype data had been filtered to remove variants: that were not in the “in_HetMiss” set defined by UKB; that had >2% cohortwide missingness; or that were found to have highly discordant allele frequencies compared to other British–Irish datasets or to be in apparent inter-chromosomal LD^105^.

#### IBD segment computation

We called IBD segments between UKB individuals’ microarray genotypes using KING v2.2.4 - -ibdseg^106^. Genotype data was split into 90 batches and run using --projection mode to calculate IBD between batches. Kinship coefficients quoted throughout the supplementary refer to the PropIBD values reported by KING divided by 2. Genotype data had been filtered to remove variants with cohortwide missingness >3%.

### ADMIXTURE

We assigned proportions of continental-scale ancestry to all UKB microarray genotypes using ADMIXTURE^100^. ADMIXTURE was run on --supervised mode using the 1000G populations CEU (Northern Europeans from Utah), CHB (Han Chinese in Beijing), ITU (Indian Telugu in the UK), PEL (Peruvians in Lima), and YRI (Yoruba in Ibadan, Nigeria) as training data. The 1000G training data had previously been filtered to remove close (at least 2nd degree) relatives using KING^106^ --kinship, to remove some apparent genomic ancestry outliers using PCA and leave-one-out unsupervised ADMIXTURE (especially PEL individuals with high European ancestry), and also pruned for LD using PLINK^103^ v1.9 --indep-pairwise 50 5 0.2. The ADMIXTURE program was run for batches of 30 UKB individuals at a time and the results subsequently merged.

#### Birthplace data

All location analyses were performed in R using the sf package^107^, the sp package^108^, and the gstat package^109^. Spatial interpolation of birthplaces was performed using linear variogram models (gstat∷vgm(), range 60,000) and ordinary kriging (gstat∷krige(), nmax = 300).

For some analysis, we binned the birthplaces into the following administrative divisions: the ceremonial counties of England; the historic counties of Wales; the 1975 local government areas of Scotland; the Isle of Man, Northern Ireland, and the [Republic of] Ireland each as their own divisions; and Jersey and Guernsey grouped together into a division we labelled the Channel Islands.

### Websites

GraphTyper https://github.com/DecodeGenetics/graphtyper

GATK resource bundle

gs://genomics-public-data/resources/broad/hg38/v0

Svimmer

https://github.com/DecodeGenetics/svimmer

popSTR

https://github.com/DecodeGenetics/popSTR

Dipcall

https://github.com/lh3/dipcall

RTG Tools

https://github.com/RealTimeGenomics/rtg-tools

bcl2fastq

https://support.illumina.com/sequencing/sequencing_software/bcl2fastq-conversion-software.html

Samtools

http://www.htslib.org/

samblaster

https://github.com/GregoryFaust/samblaster

BamQC

https://github.com/DecodeGenetics/BamQC

GIAB WGS samples

• HG001 https://ftp-trace.ncbi.nlm.nih.gov/ReferenceSamples/giab/data/NA12878/NIST_NA12878_HG001_HiSeq_300x/NHGRI_Illumina300X_novoalign_bams/HG001.GRCh38_full_plus_hs38d1_analysis_set_minus_alts.300x.bam

• HG002 https://ftp-trace.ncbi.nlm.nih.gov/ReferenceSamples/giab/data/AshkenazimTrio/HG002_NA24385_son/NIST_HiSeq_HG002_Homogeneity-10953946/NHGRI_Illumina300X_AJtrio_novoalign_bams/HG002.GRCh38.60x.1.bam

• HG003 https://ftp-trace.ncbi.nlm.nih.gov/ReferenceSamples/giab/data/AshkenazimTrio/HG003_NA24149_father/NIST_HiSeq_HG003_Homogeneity-12389378/NHGRI_Illumina300X_AJtrio_novoalign_bams/HG003.GRCh38.60x.1.bam

• HG004 https://ftp-trace.ncbi.nlm.nih.gov/ReferenceSamples/giab/data/AshkenazimTrio/HG004_NA24143_mother/NIST_HiSeq_HG004_Homogeneity-14572558/NHGRI_Illumina300X_AJtrio_novoalign_bams/HG004.GRCh38.60x.1.bam

• HG005 https://ftp-trace.ncbi.nlm.nih.gov/ReferenceSamples/giab/data/ChineseTrio/HG005_NA24631_son/HG005_NA24631_son_HiSeq_300x/NHGRI_Illumina300X_Chinesetrio_novoalign_bams/HG005.GRCh38_full_plus_hs38d1_analysis_set_minus_alts.300x.bam

• HG006 https://ftp-trace.ncbi.nlm.nih.gov/ReferenceSamples/giab/data/ChineseTrio/HG006_NA24694-huCA017E_father/NA24694_Father_HiSeq100x/NHGRI_Illumina100X_Chinesetrio_novoalign_bams/HG006.GRCh38_full_plus_hs38d1_analysis_set_minus_alts.100x.bam

• HG007 https://ftp-trace.ncbi.nlm.nih.gov/ReferenceSamples/giab/data/ChineseTrio/HG007_NA24695-hu38168_mother/NA24695_Mother_HiSeq100x/NHGRI_Illumina100X_Chinesetrio_novoalign_bams/HG007.GRCh38_full_plus_hs38d1_analysis_set_minus_alts.100x.bam

ENSEMBL

https://m.ensembl.org/info/data/mysql.html

Shapefiles for UK

http://discover.ukdataservice.ac.uk/catalogue/?sn=5819&tyep=Data%20catalogue

http://census.ukdataservice.ac.uk/get-data/boundary-data.aspx

https://gadm.org/

Exon capture regions

http://biobank.ndph.ox.ac.uk/ukb/ukb/auxdata/xgen_plus_spikein.b38.bed

ClinVar

https://www.ncbi.nlm.nih.gov/clinvar/

UKB data showcase

https://biobank.ndph.ox.ac.uk/showcase/search.cgi

GERP

http://mendel.stanford.edu/SidowLab/downloads/gerp/hg19.GERP_scores.tar.gz

Eigen

http://www.funlda.com/toolkit

LINSIGHT

http://compgen.cshl.edu/LINSIGHT/

CADD

https://cadd.gs.washington.edu/download

Open Targets

https://genetics.opentargets.org/

AffiXcan

https://rdrr.io/bioc/AffiXcan/man/trainingCovariates.html

umap

https://github.com/tkonopka/umap

