## Supplement for "The sequences of 150,119 genomes in the UK biobank"

### Supplementary Figures

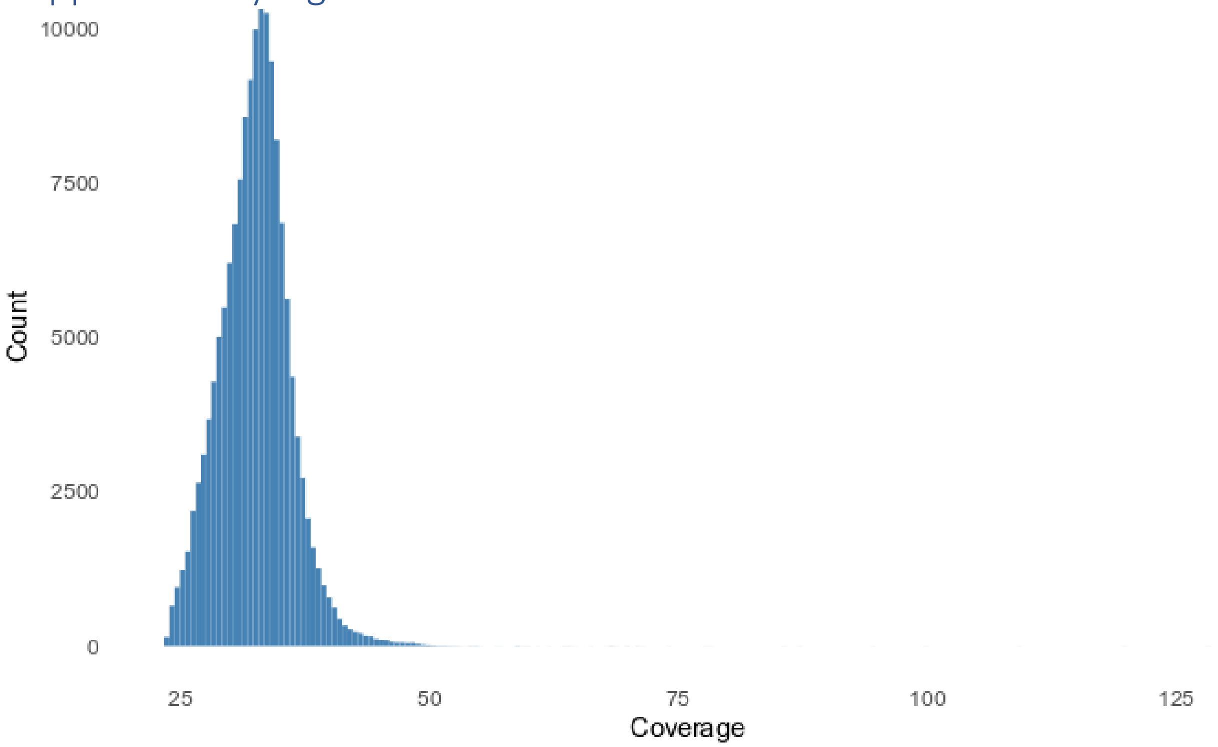


*Fig. S1 Histogram of average sequence coverage per sample in the 150,119 WGS samples.*

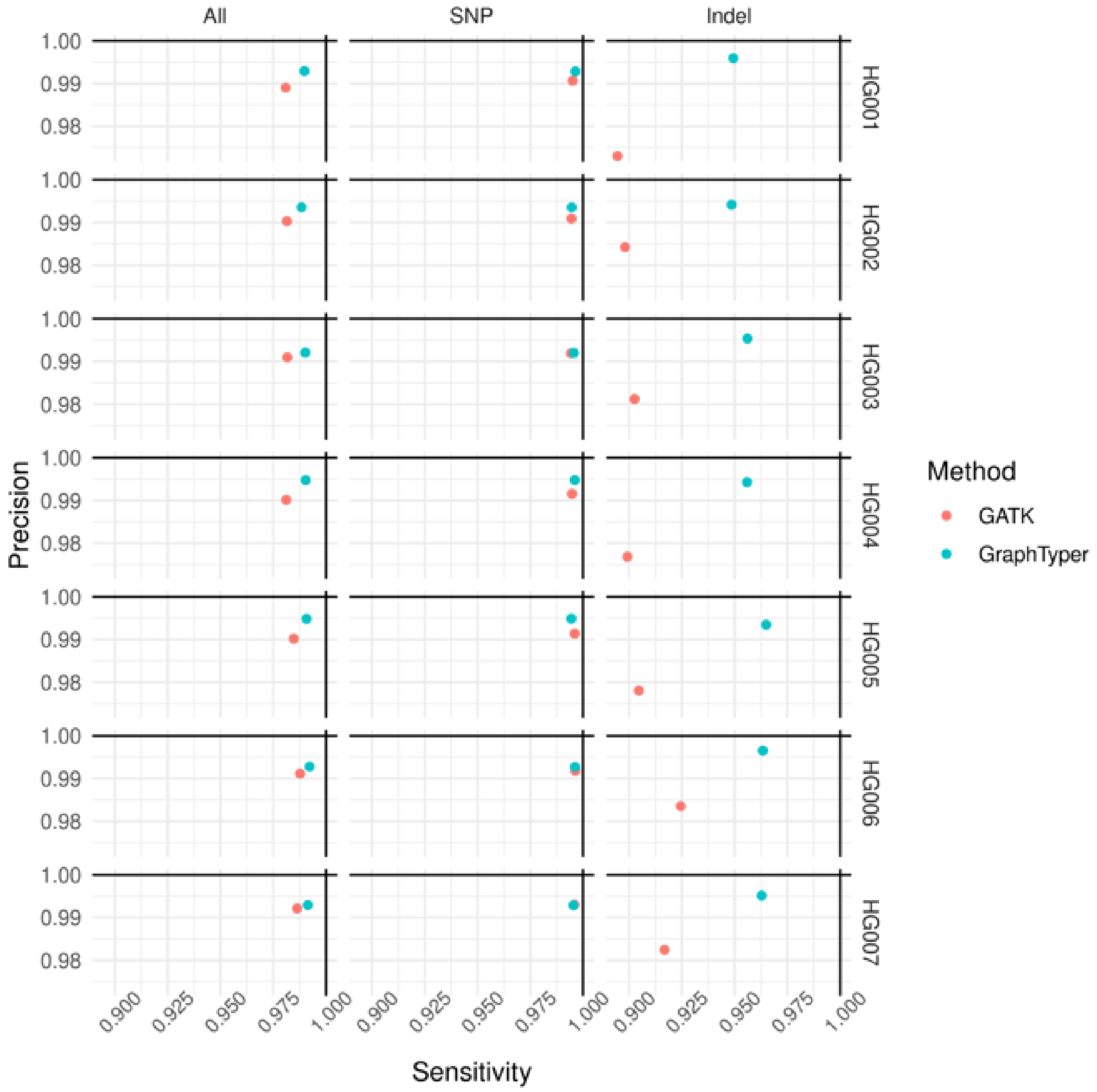


*Fig. S2 Sensitivity and precision for GATK and GraphTyper callsets in 500 regions benchmarking dataset across the seven Genome in a bottle (GIAB) v3.3.2 truth sets.*

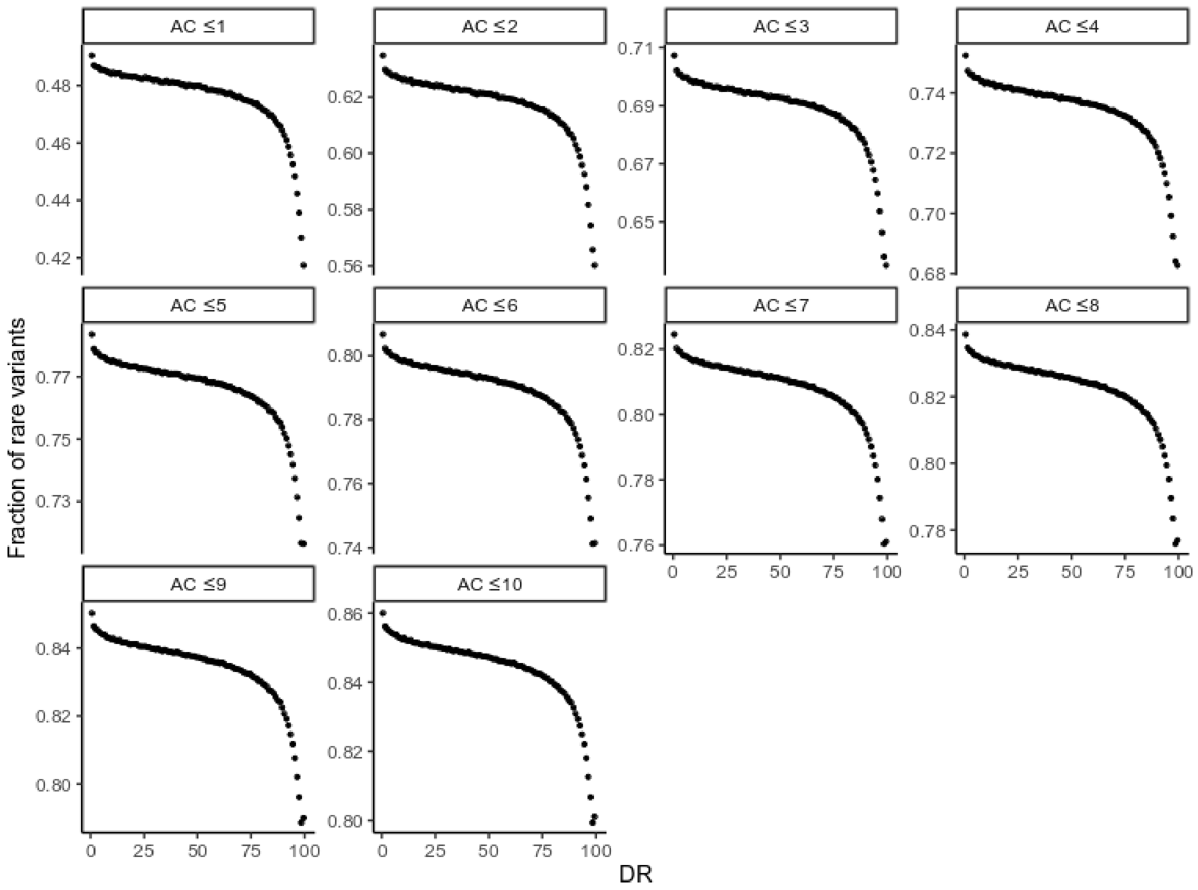


*Fig. S3 Fraction of rare variants (FRV) as a function of the definition of "rare", varying the allele count cutoff from at most 1 to at most 10 carriers. Note that homozygous carriers have an allele count of 2.*

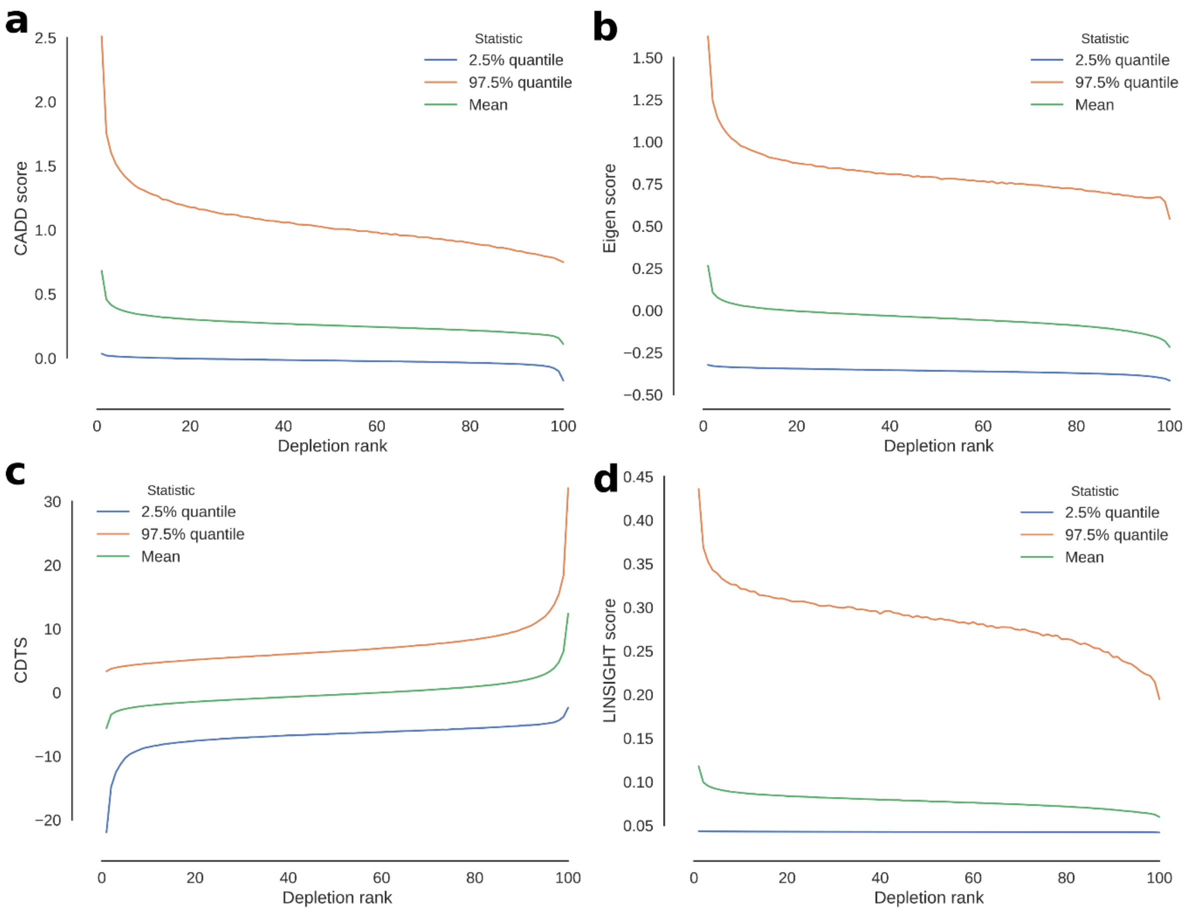


*Fig. S4 Average score in 500bp windows as a function of Depletion Rank for a) CADD b) Eigen c) CDTS and d) LINSIGHT. Green line represents average score, blue and red line 95-th percentile*

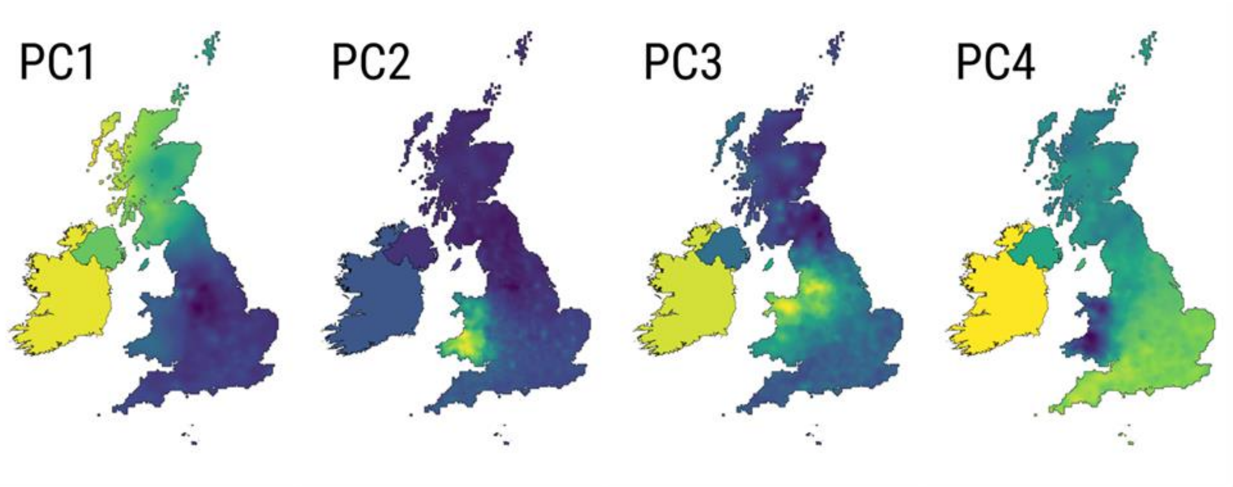

*Fig. S5 Geographic distribution of the loadings of the first four principal components of a PCA of the XBI population.*

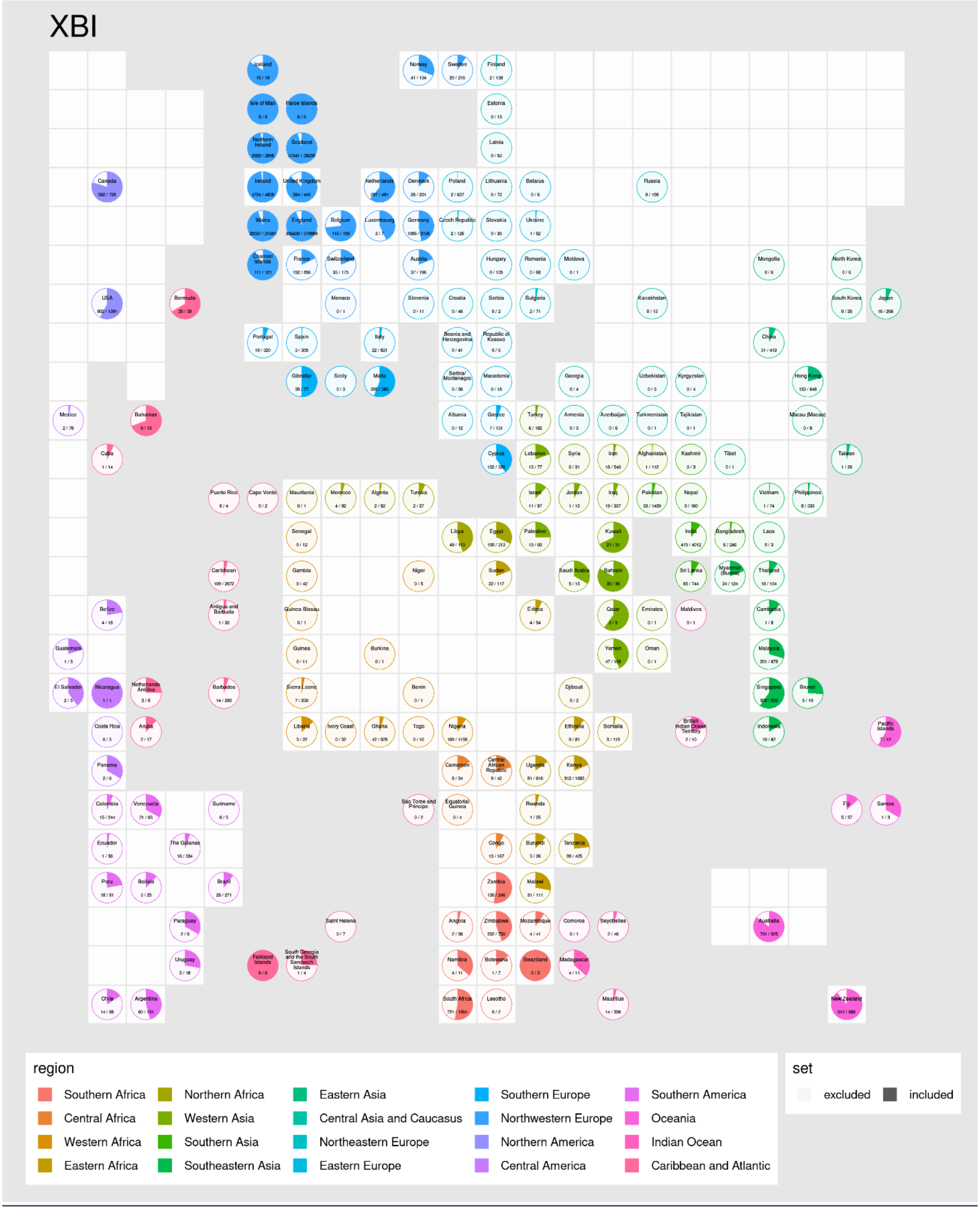


*Fig. S6 Cartogram-pies indicating the proportion of individuals born in each country (name shown on top of pies) in the XBI cohort. Pies are placed roughly according to their country's position on a world map. Grey and white squares represent sea and land respectively.*

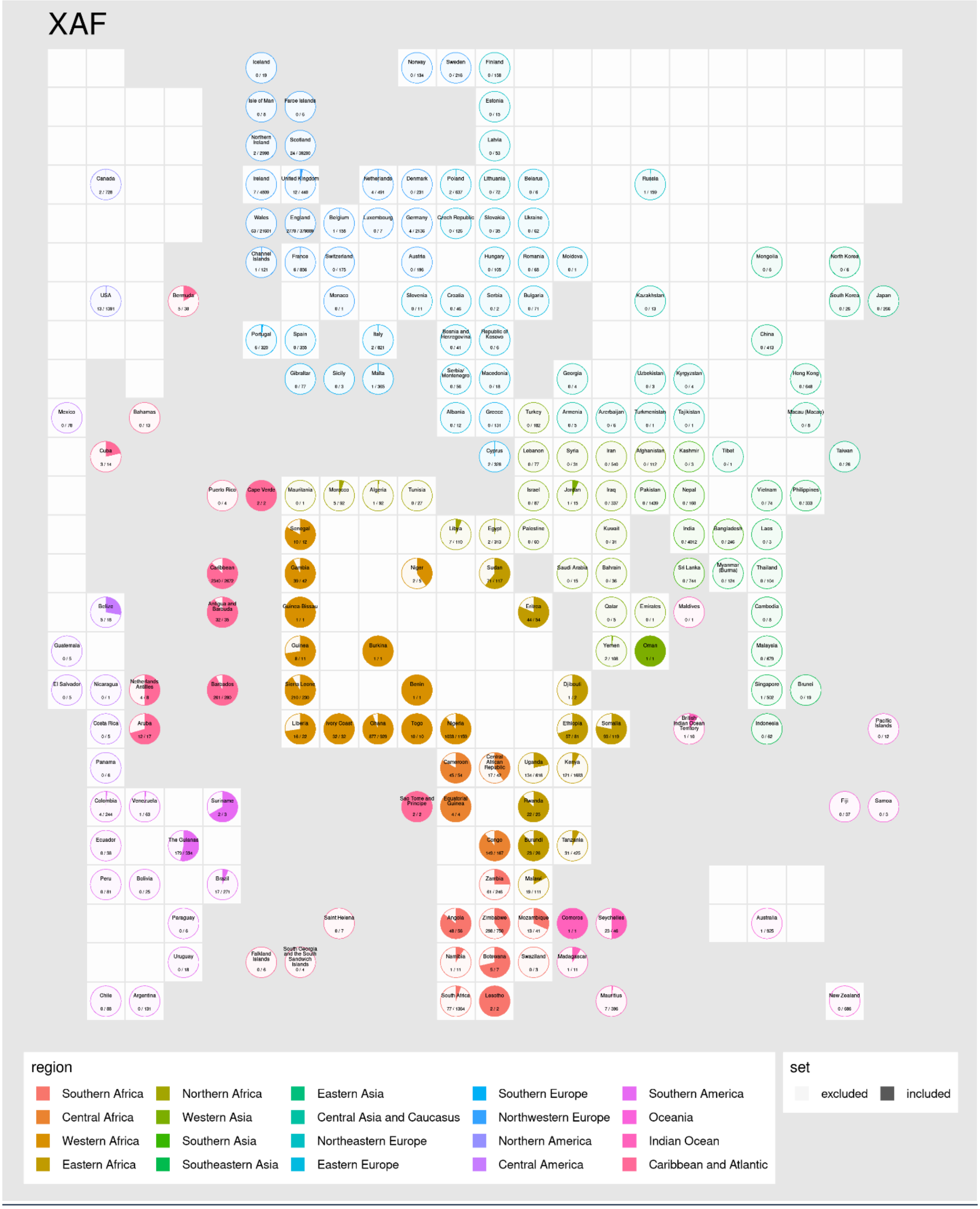


*Fig. S7 Cartogram-pies indicating the proportion of individuals born in each country (name shown on top of pies) in the XAF cohort. Pies are placed roughly according to their country's position on a world map. Grey and white squares represent sea and land respectively.*

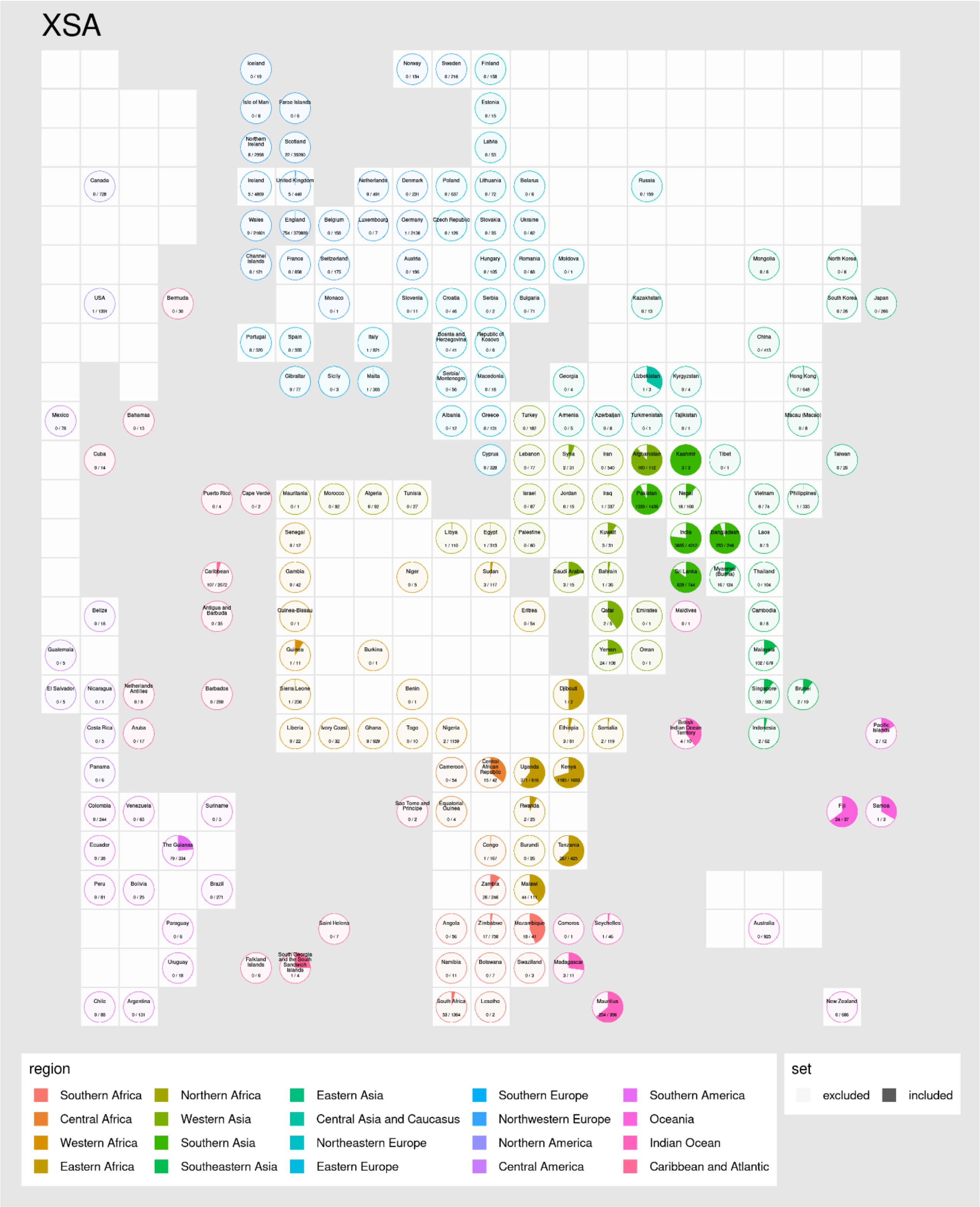


*Fig. S8 Cartogram-pies indicating the proportion of individuals born in each country (name shown on top of pies) in the XSA cohort. Pies are placed roughly according to their country's position on a world map. Grey and white squares represent sea and land respectively.*

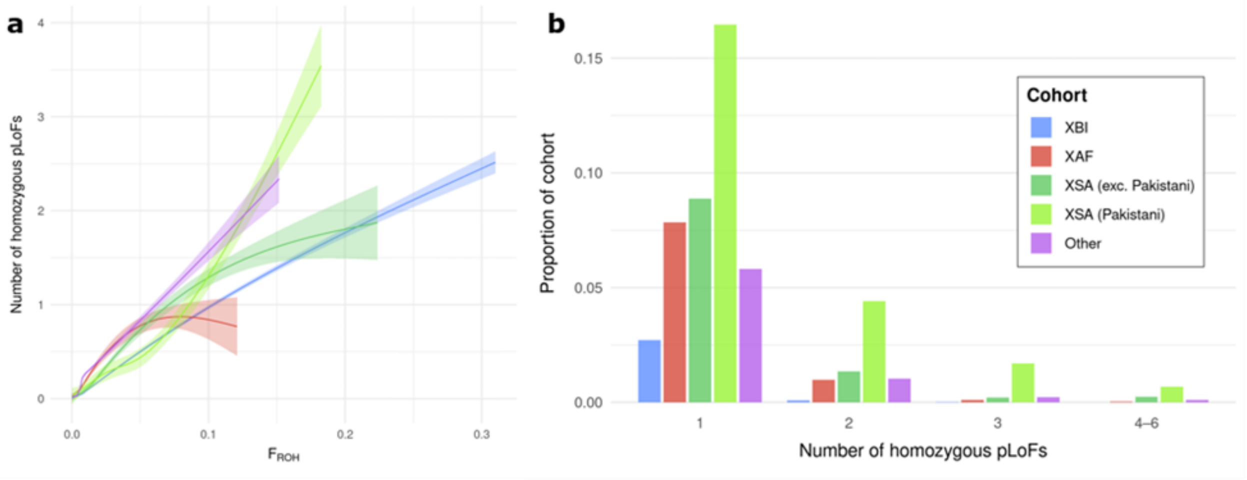


*Fig. S9 Loss-of-function a) Correlation between the number of LoF genes per sample and fraction of genome with runs of homozygosity. b) Number of homozygous loss-of-function (LoF) genes per sample. Count of homozygous genes annotated as high impact with frequency <1%. Results are presented for XBI, XAF, XSA excluding individuals self-identified as Pakistani, individuals self-identified as Pakistani from the XSA cohort and Others.*

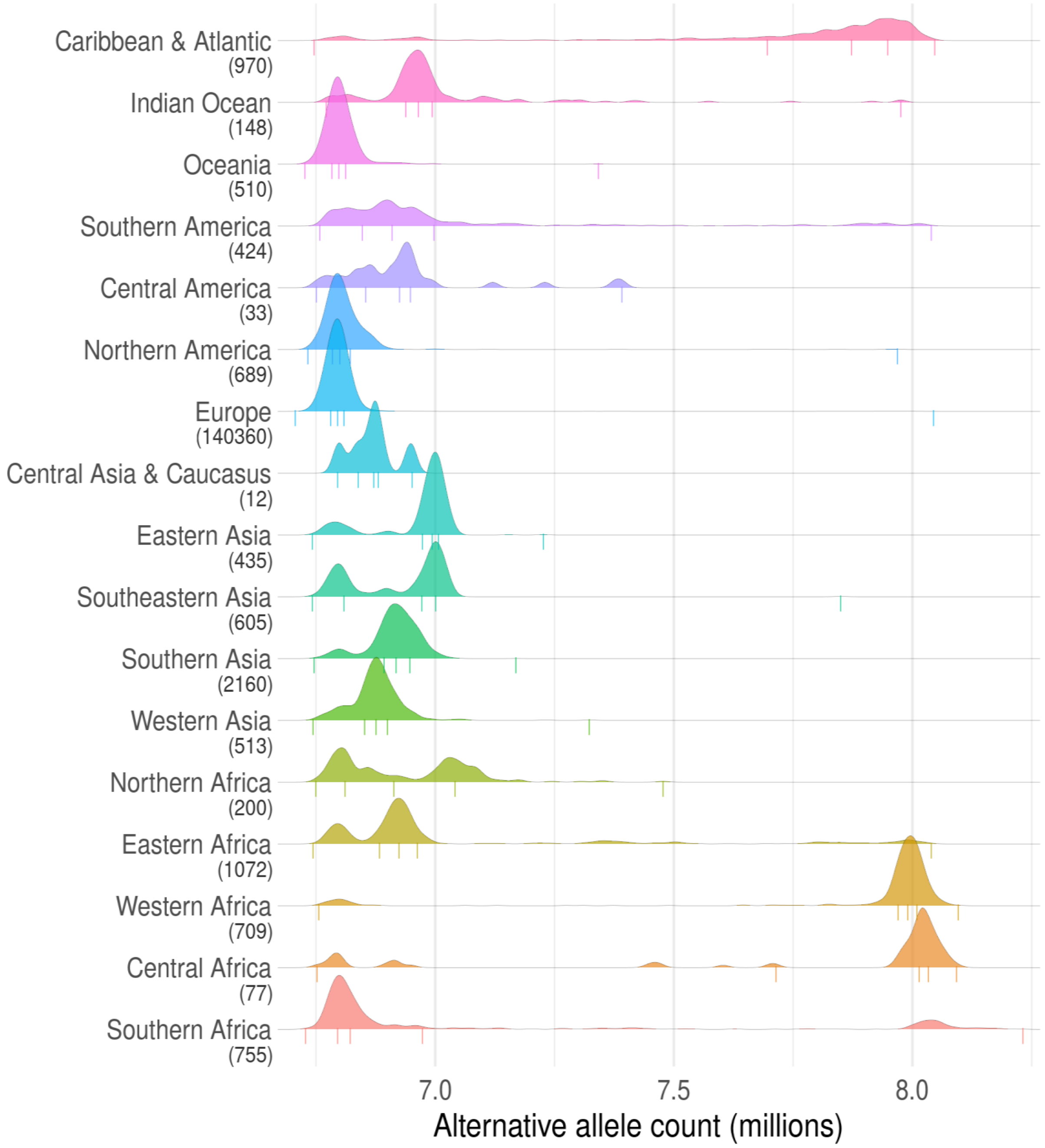


*Fig. S10: Alternative alleles by region. Numbers in brackets beneath region names indicate count of whole genome sequenced individuals with birthplaces in that region. Assignment of countries to regions is almost identical to the categorization displayed in the cohort cartogram pie figures, with the exception that all European regions are combined into one region in this figure. Vertical lines underneath density curves represent 0th, 25th, 50th, 75th, and 100th percentiles.*

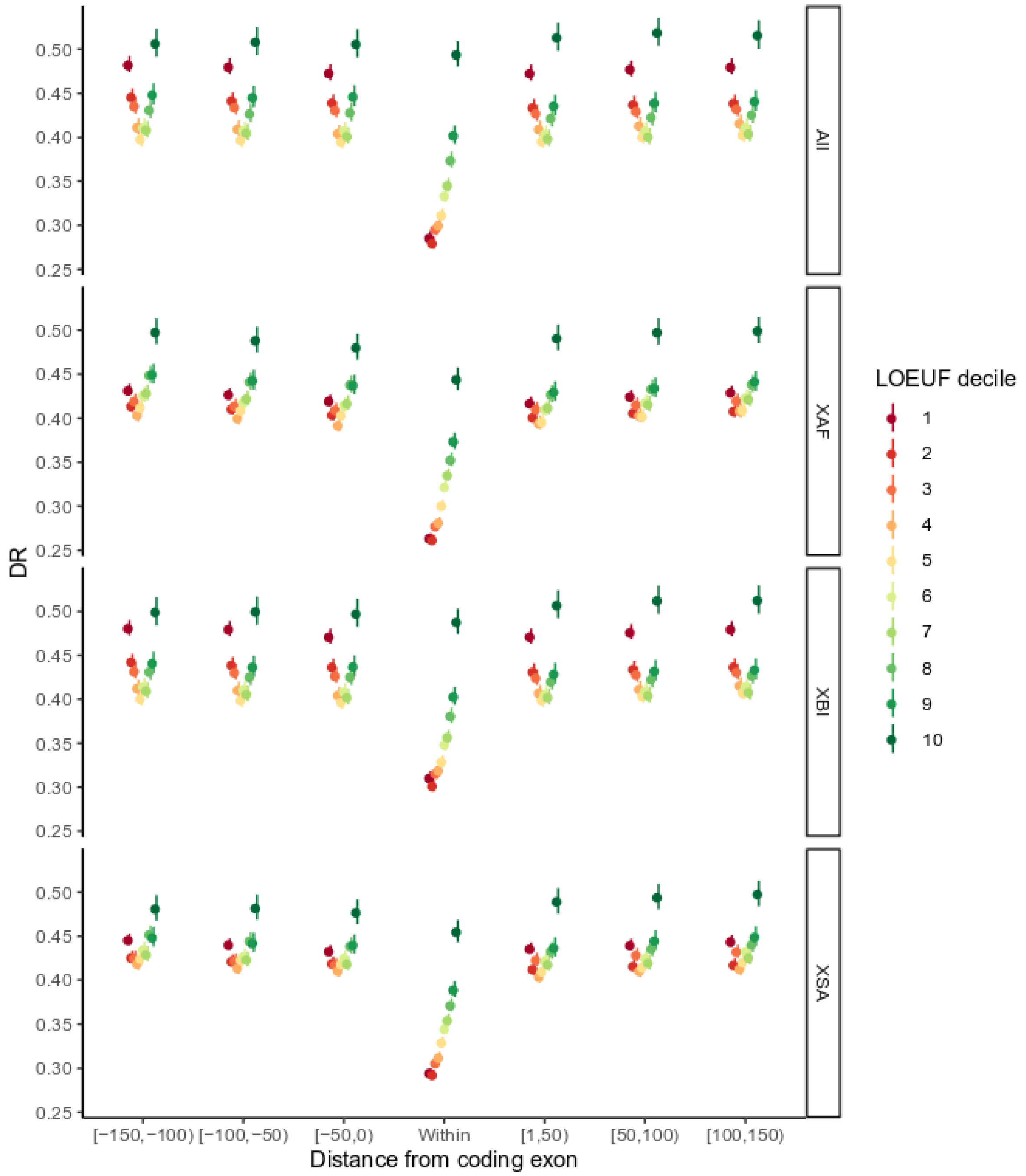


*Fig. S11 DR as a function of distance from coding exon partitioned by LOEUF[11] deciles. Results are shown separately for the overall dataset (All) and the individual cohorts, XBI, XAF and XSA.*

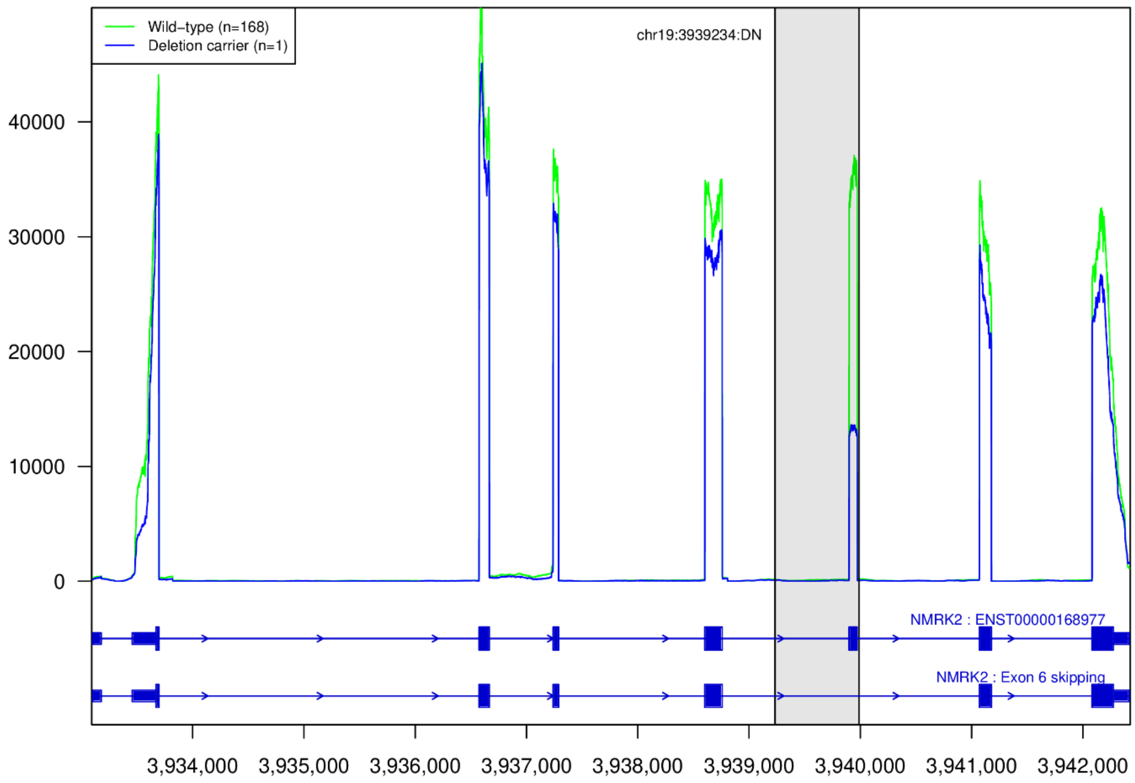


*Fig. S12 Coverage plot of RNA-sequenced reads from heart tissue from 169 heart tissue samples over the gene NMRK2. One individual is a carrier of a 754bp deletion depicted with gray rectangle that includes exon 6 of NMRK2. The RNA-coverage of the carrier (blue) is lower over exon 6 compared to median coverage of non-carriers (green). Shading marks the deleted region.*

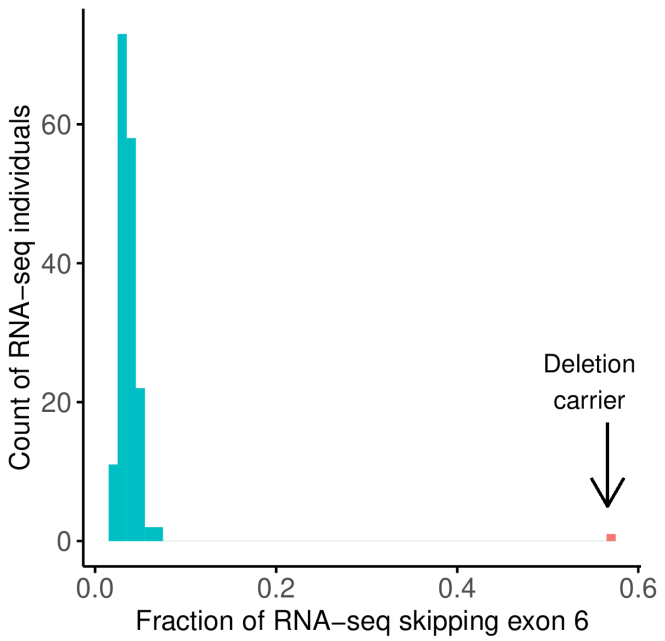


*Fig. S13 Histogram of fraction of RNA-sequenced fragments skipping exon 6 in NMRK2 out of all fragments aligning from the donor site of exon 5 to either acceptor site of exon 6 or exon 7. The median fraction fragments skipping for wild-type individuals is 0.035 and 0.57 for the carrier of the 754bp deletion.*

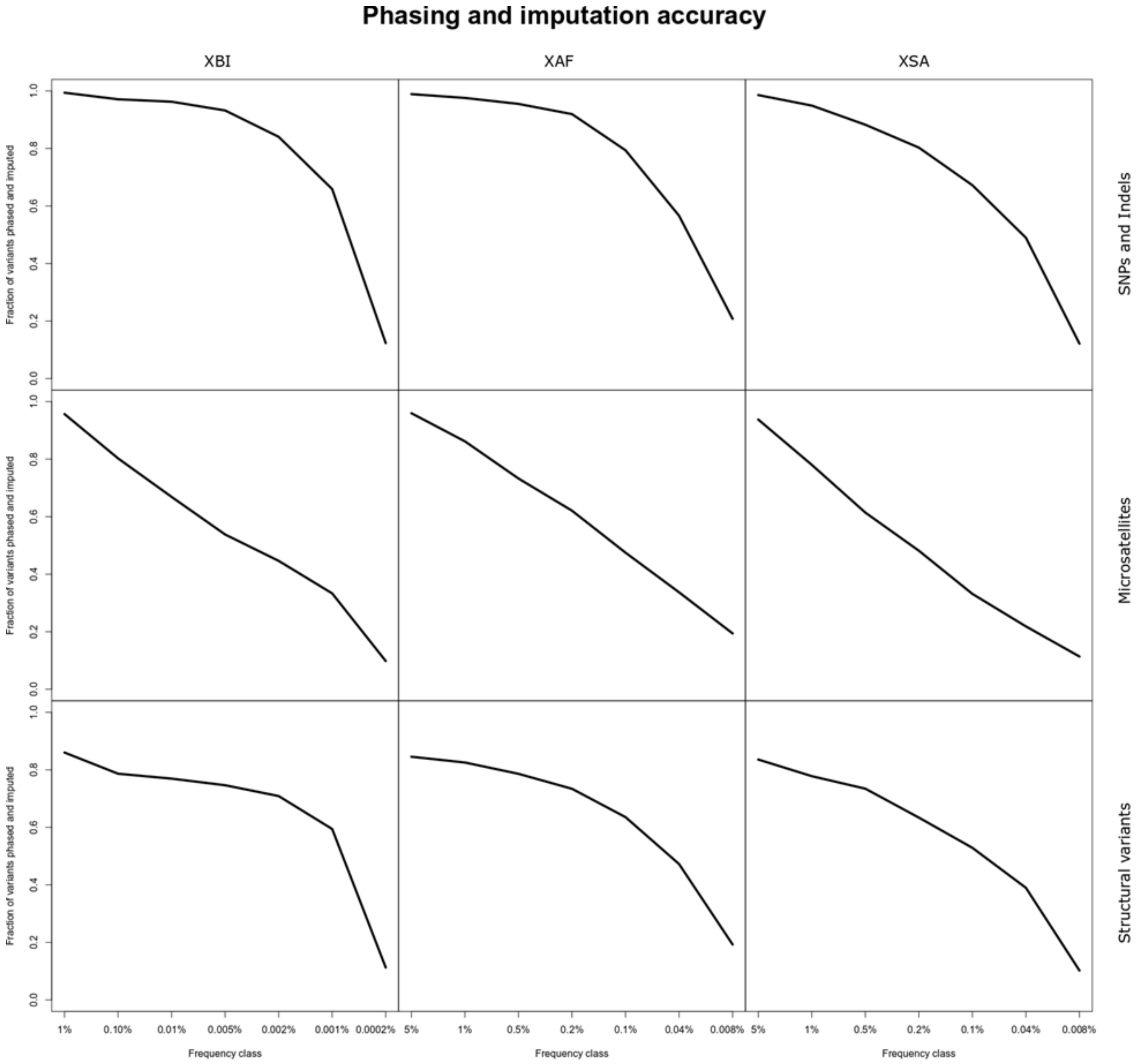

*Fig. S14 Imputation and phasing accuracy across variant datasets in the three populations. A variant is considered imputed if Leave one out r2 (L1or2) of phasing was greater than 0.5 and imputation information was greater than 0.8. x-axis splits variants into frequency classes based on the frequency in each cohort.*

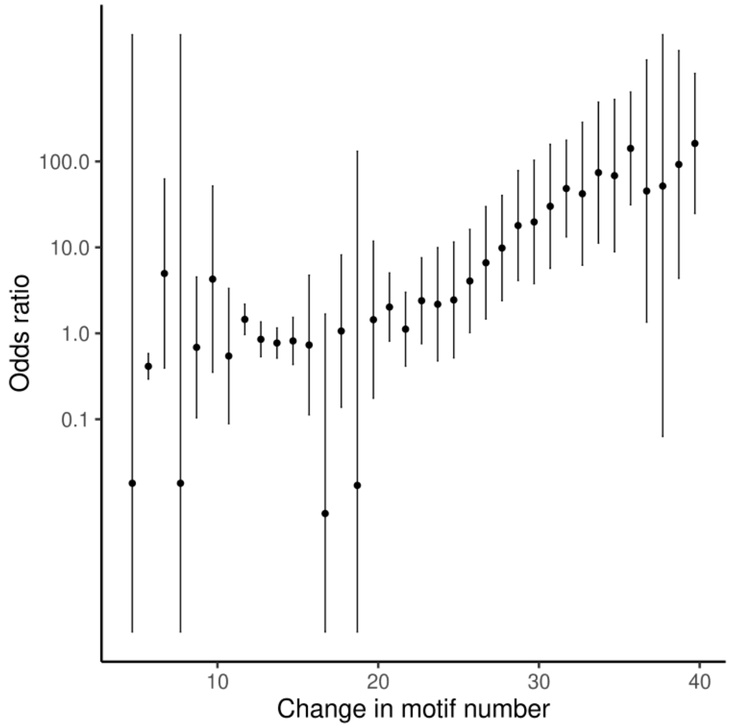


*Fig. S15 Odds ratio for risk of myotonic dystrophy as a function of repeat length in microsatellite at the 3' untranslated region of DMPK. Carriers of at least 39.7 copies of the microsatellite repeat motif have a 162-fold increased risk of myotonic dystrophy.*

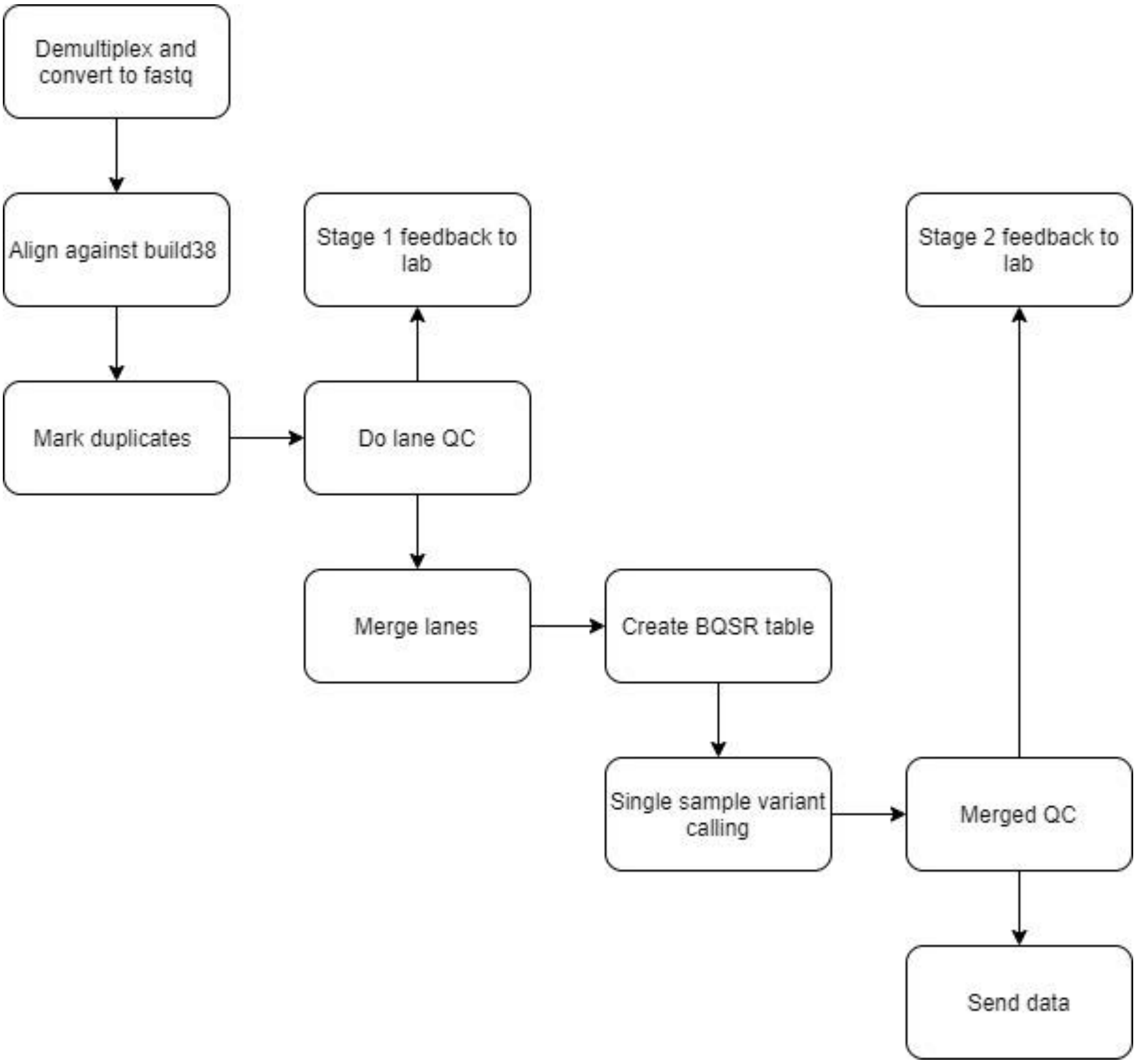


*Fig. S16 Process outline for UKB sequencing pipeline at deCODE genetics.*

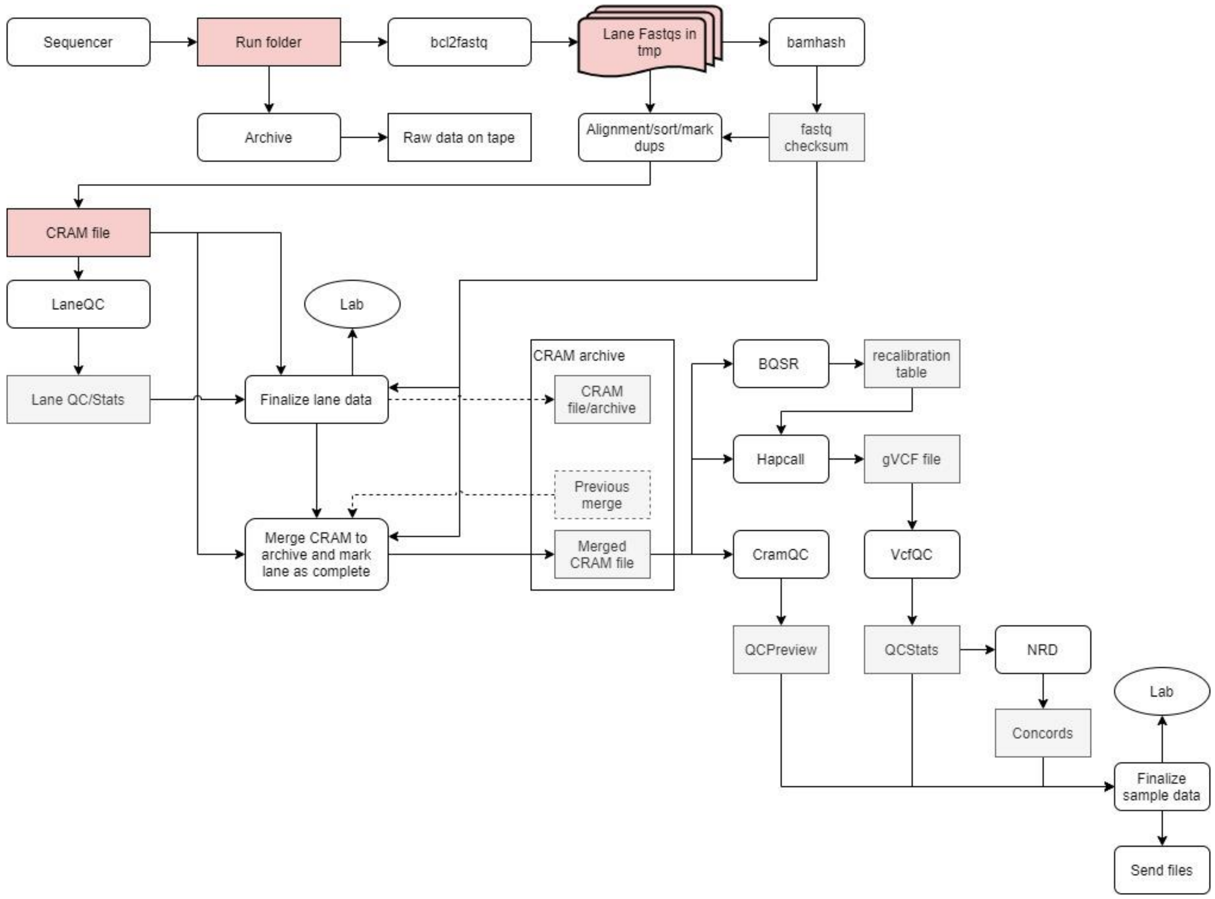


*Fig. S17 Pipeline for processing of sequence data at deCODE genetics.*

```
QC_VERDICT = 'PASS'

    if freemix_percentage >= 1.0:
        QC_VERDICT = 'REVIEW'

    if coverage < 26:
        QC_VERDICT = 'REVIEW'

    if freemix_percentage >= 5.0:
        QC_VERDICT = 'FAIL'

    if prc_proper_pairs < 95.0:
        QC_VERDICT = 'FAIL'

    if prc_auto_ge_15x < 95.0:
        QC_VERDICT = 'FAIL'

    if discordance_prc is not -1 and discordance_prc >= 2.0:
         QC_VERDICT = 'FAIL'
```

*Fig. S18 Logic used to compute PASS/FAIL for a WGS cram file.*

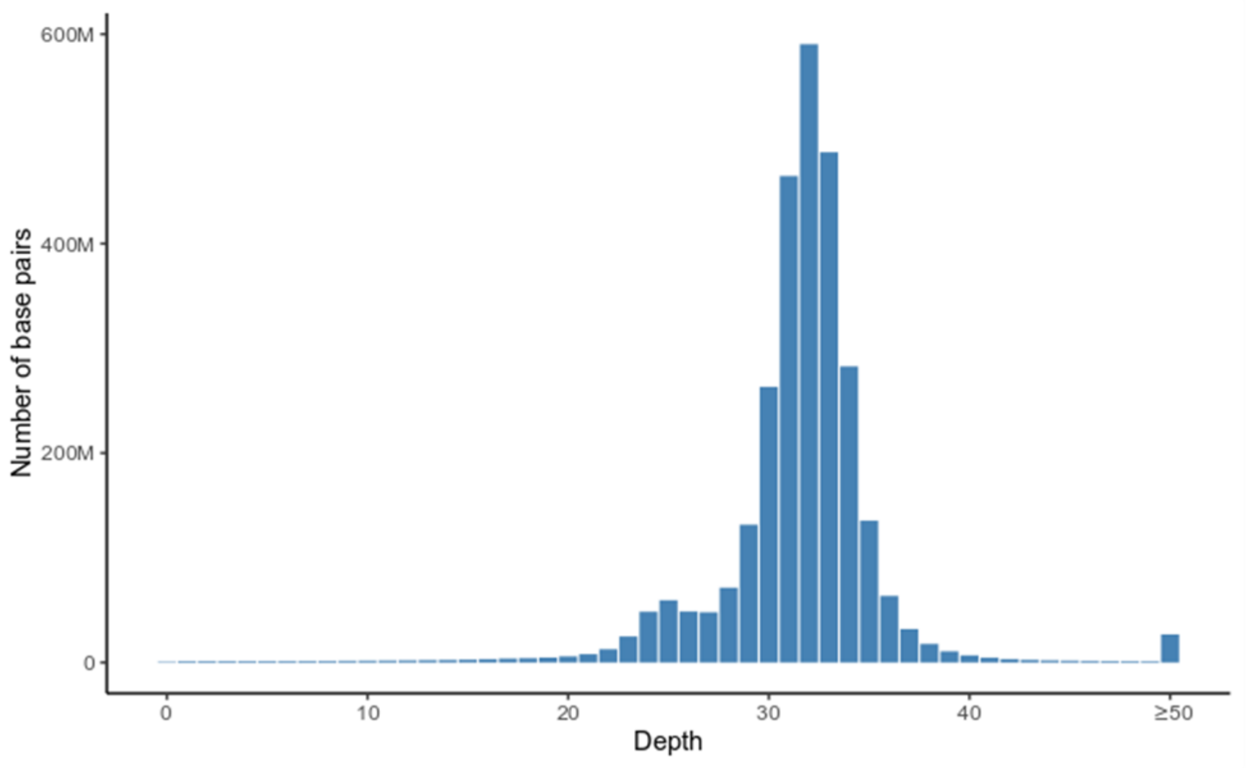


*Fig. S19 Average sequence coverage per base pair across the genome. The average coverage is computed from 1,000 randomly selected samples.*

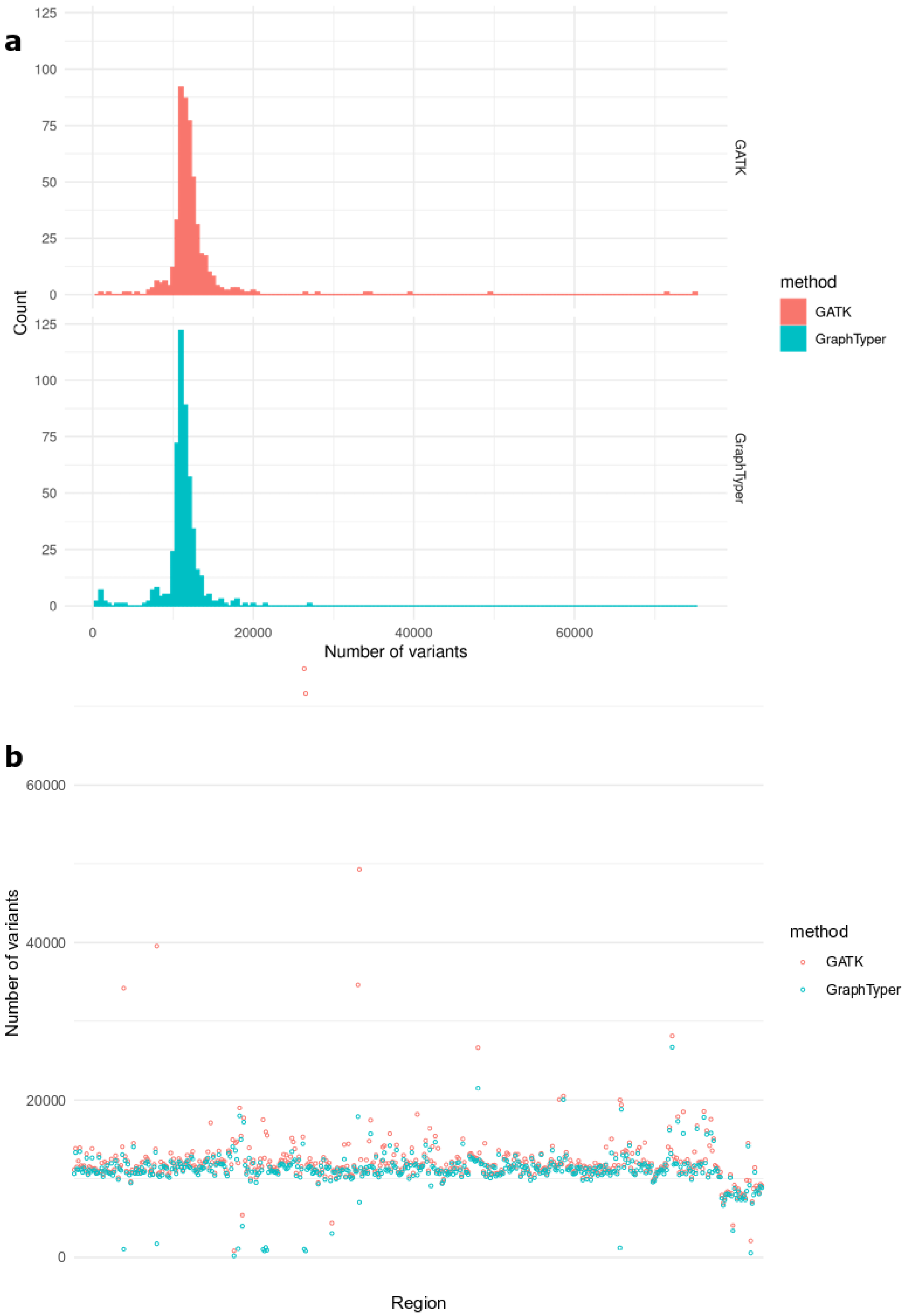

*Fig. S20 Number of variants per region in the 500 regions test set for the GATK and GraphTyper callsets, presented as a histogram a) and ordered by region b).*

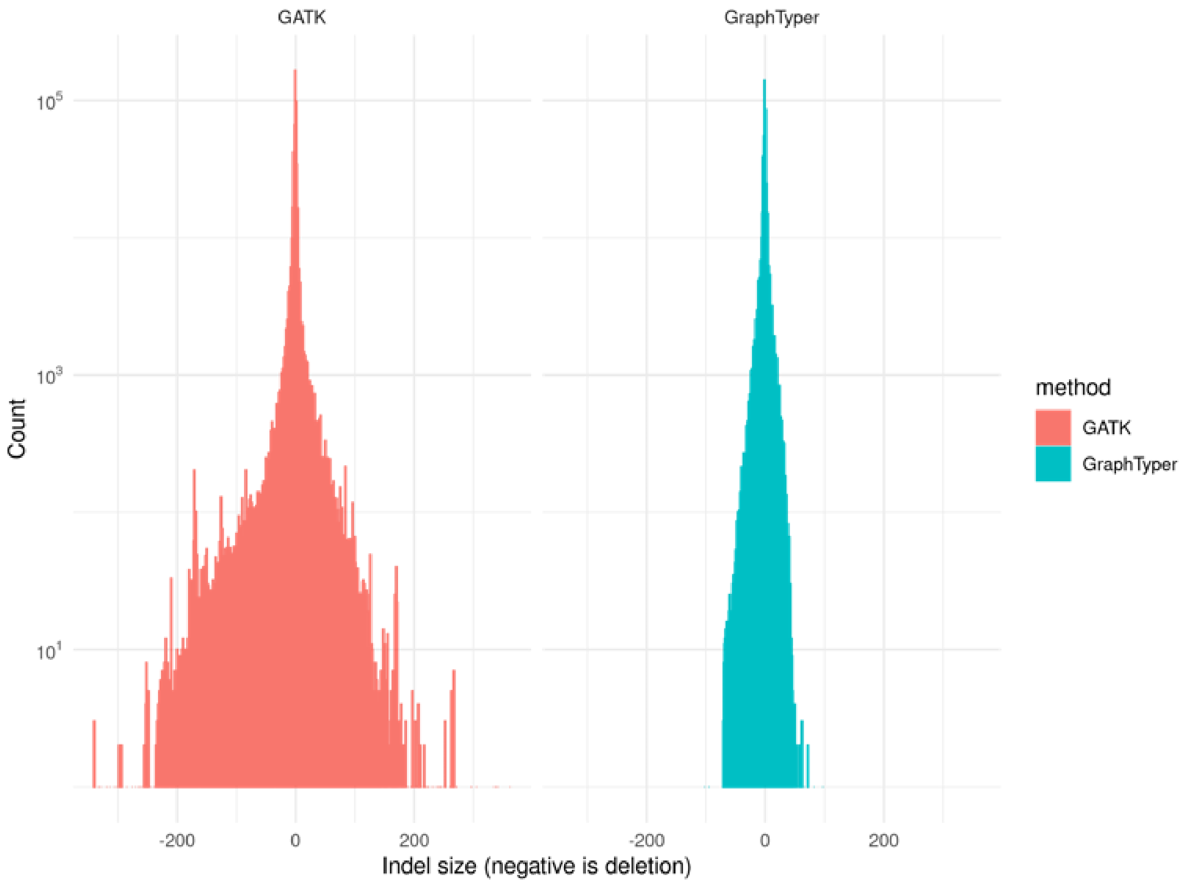

*Fig. S21 Distribution of indel sizes in GATK and GraphTyper callsets. Negative size indicates a deletion.*

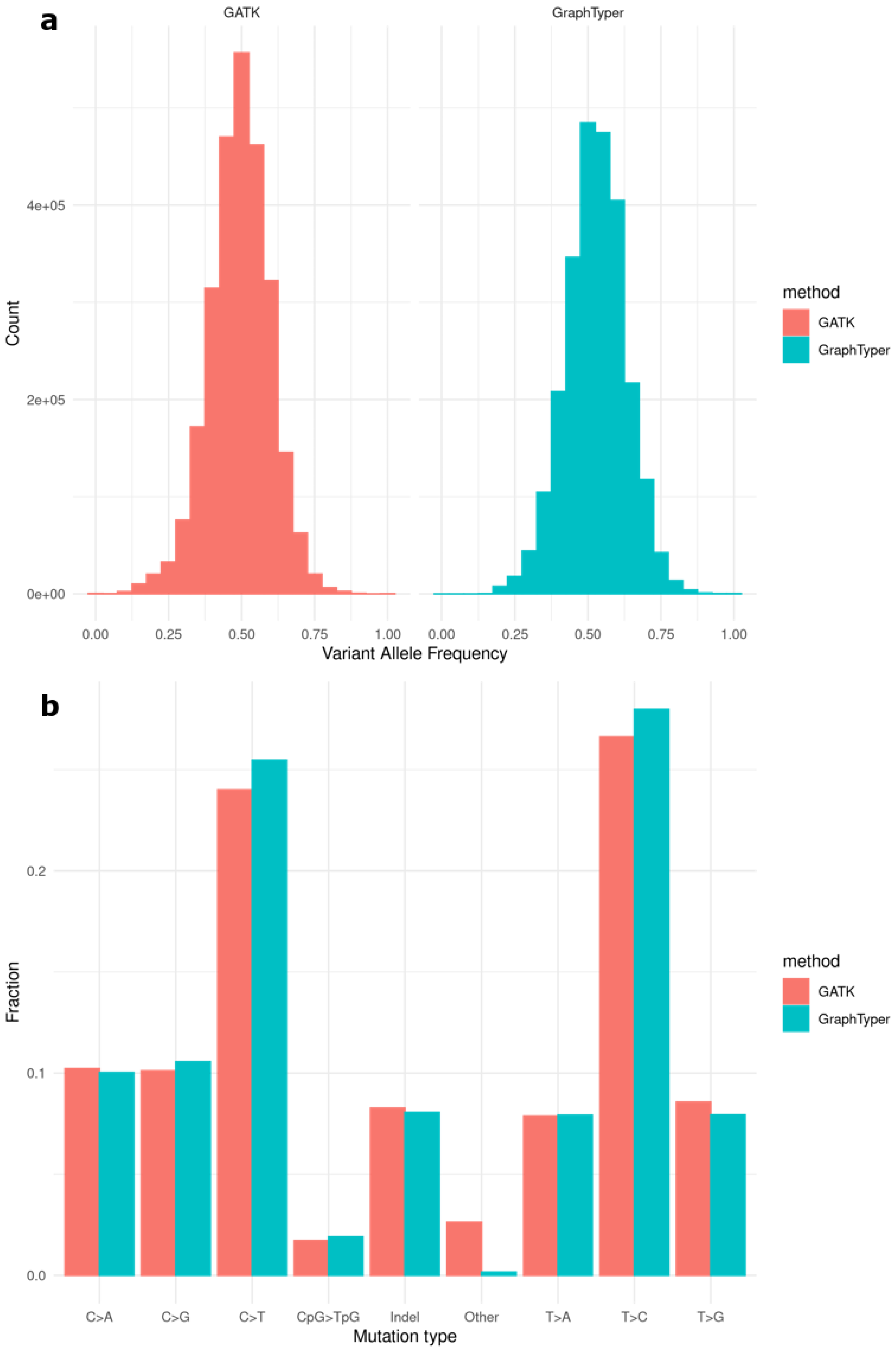


*Fig. S22 a) Variant allele frequencies (VAF) of singletons. b) Mutation classes of singletons. Results are for the GATK and GraphTyper callsets on 500 randomly selected regions.*

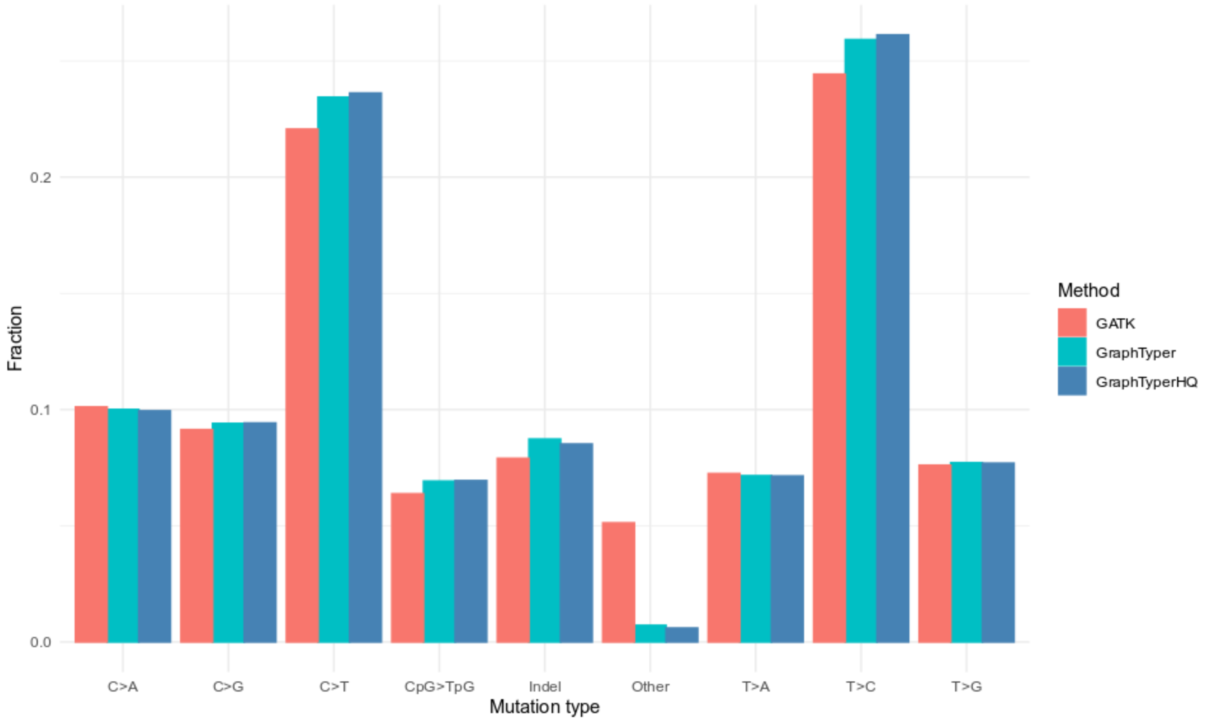


*Fig. S23 Fraction of variants by mutation type in the GATK, GraphTyper and GraphTyper HQ sets.*

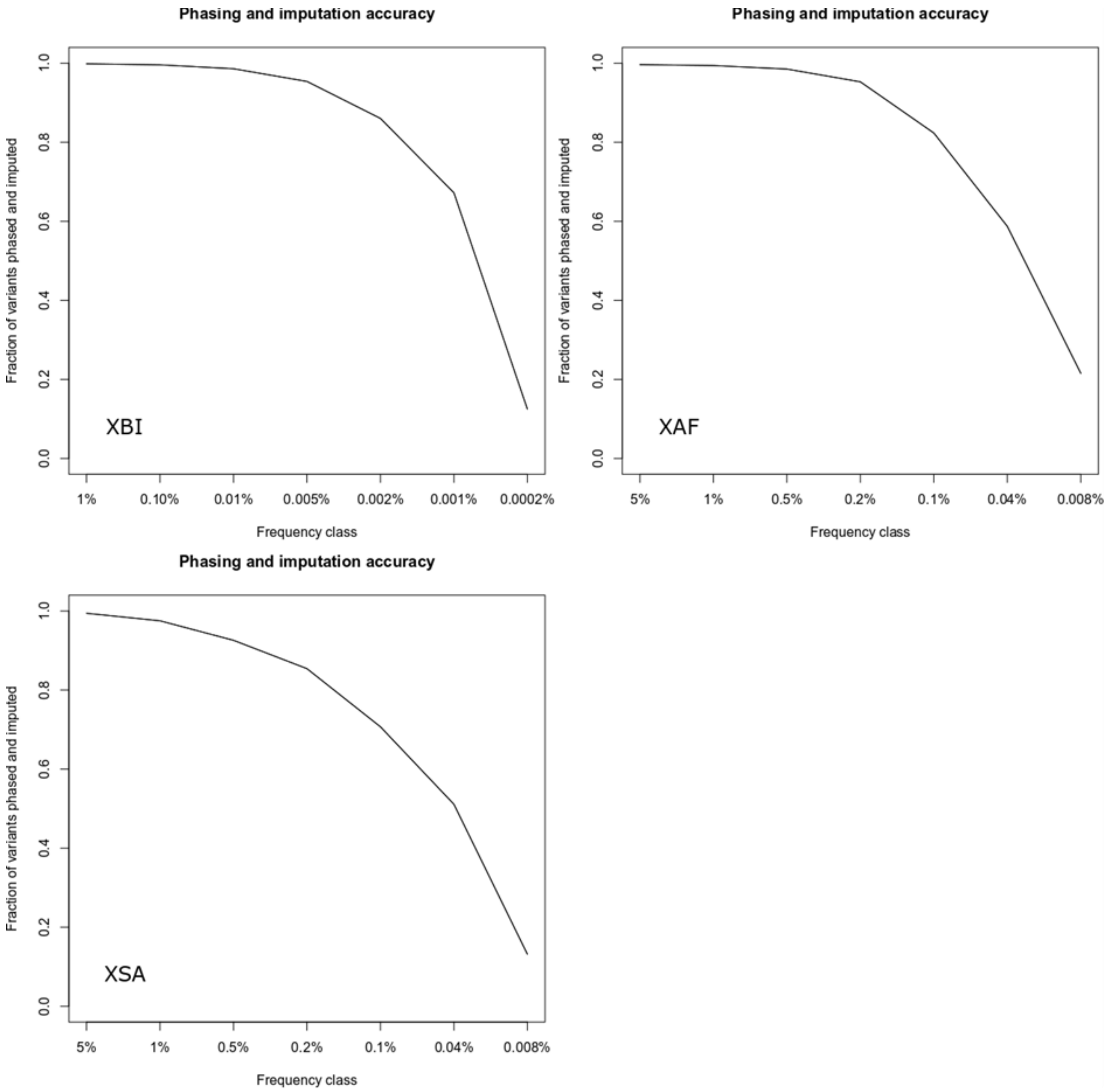


*Fig. S24 Imputation accuracy for variants with AAscore > 0.9 in the three populations, Top left: XBI, Top Right: XAF, Bottom: XSA. A variant was considered imputed if Leave one out r2 of phasing was greater than 0.5 and imputation information was greater than 0.8. x-axis splits variants into frequency classes based on the number of carriers in the sequence dataset, with the number representing the minimum number of carriers in the frequency class. Variants are split by variant type.*

*Fig. S25 Manhattan plots, quantile-quantile (QQ) plots and histograms of inverse-normal transformed values after adjustment for covariates age, sex and 40 principal components, when applicable, for quantitative traits with significant results reported in this manuscript. For Manhattan plots, the x-axis represents chromosome locations and the y-axis shows the –log10 significance levels of the associations. For QQ plots, the inflation (λ) is shown in the title of each graph, for all variants and for rare variants only (λ_maf<0.01). For the histograms, the x-axis shows the value range of the inverse-normal transformed points and the y-axis shows the count of individuals within value ranges.*

a) Total cholesterol, structural variant analysis, European ancestry (N=412,119)

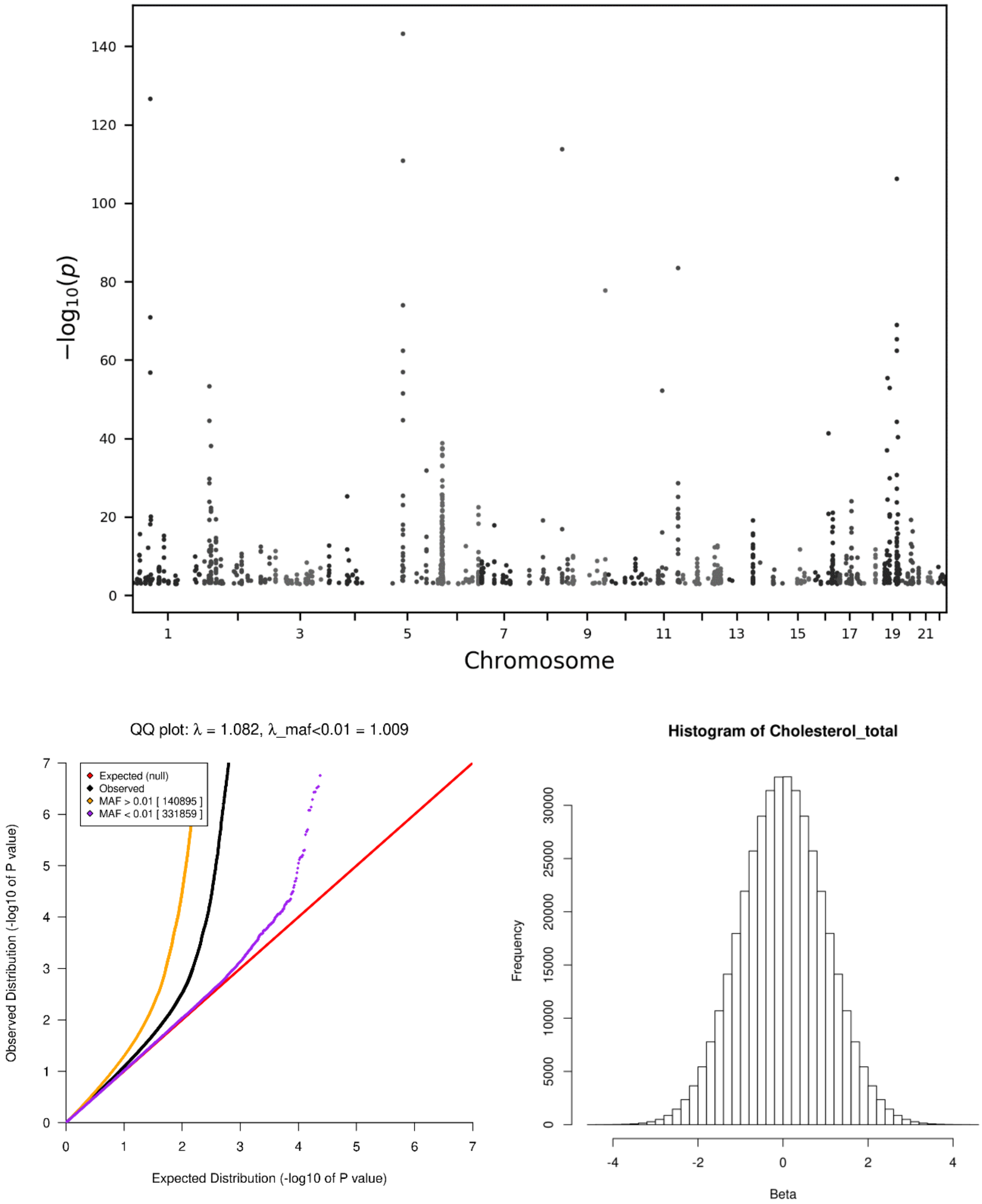

b) Calcium levels, structural variant analysis, European ancestry (N=378,246)

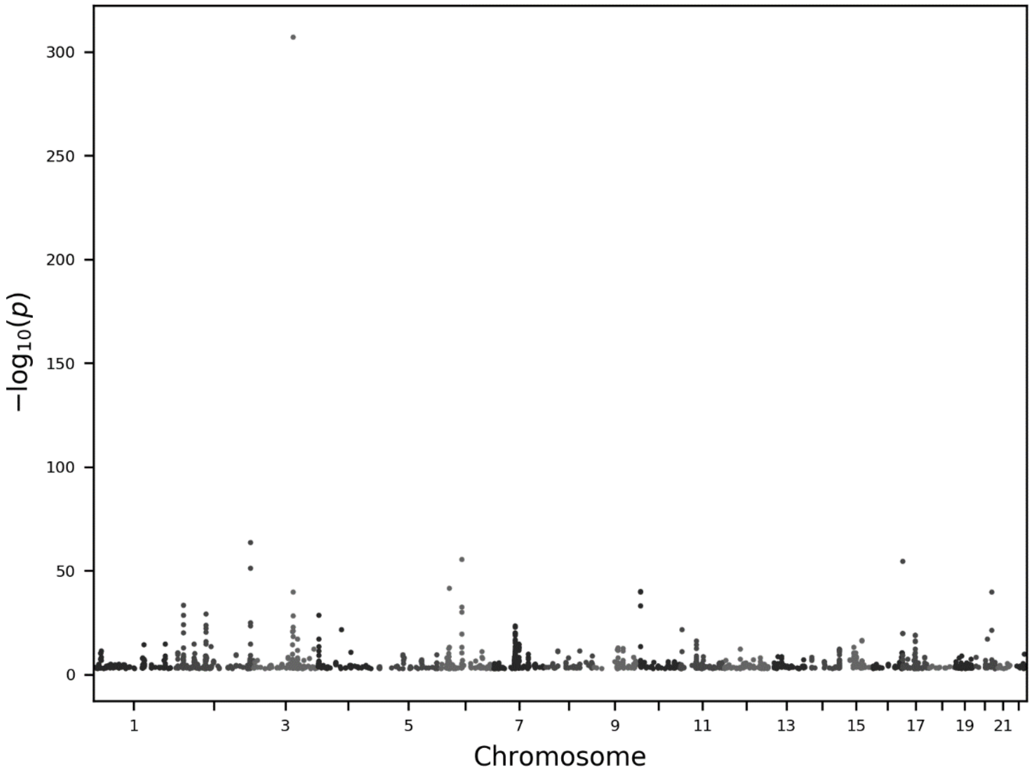


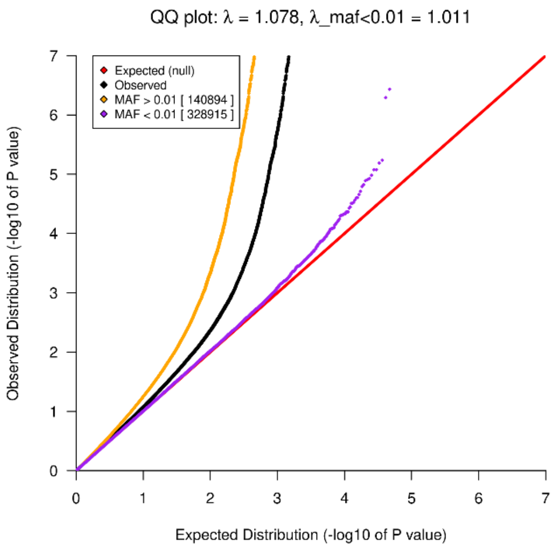


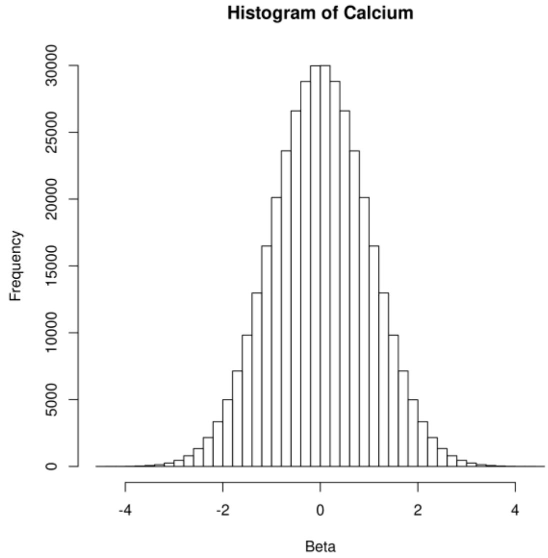

c) Albumin levels, structural variant analysis, European ancestry (N=378,395)

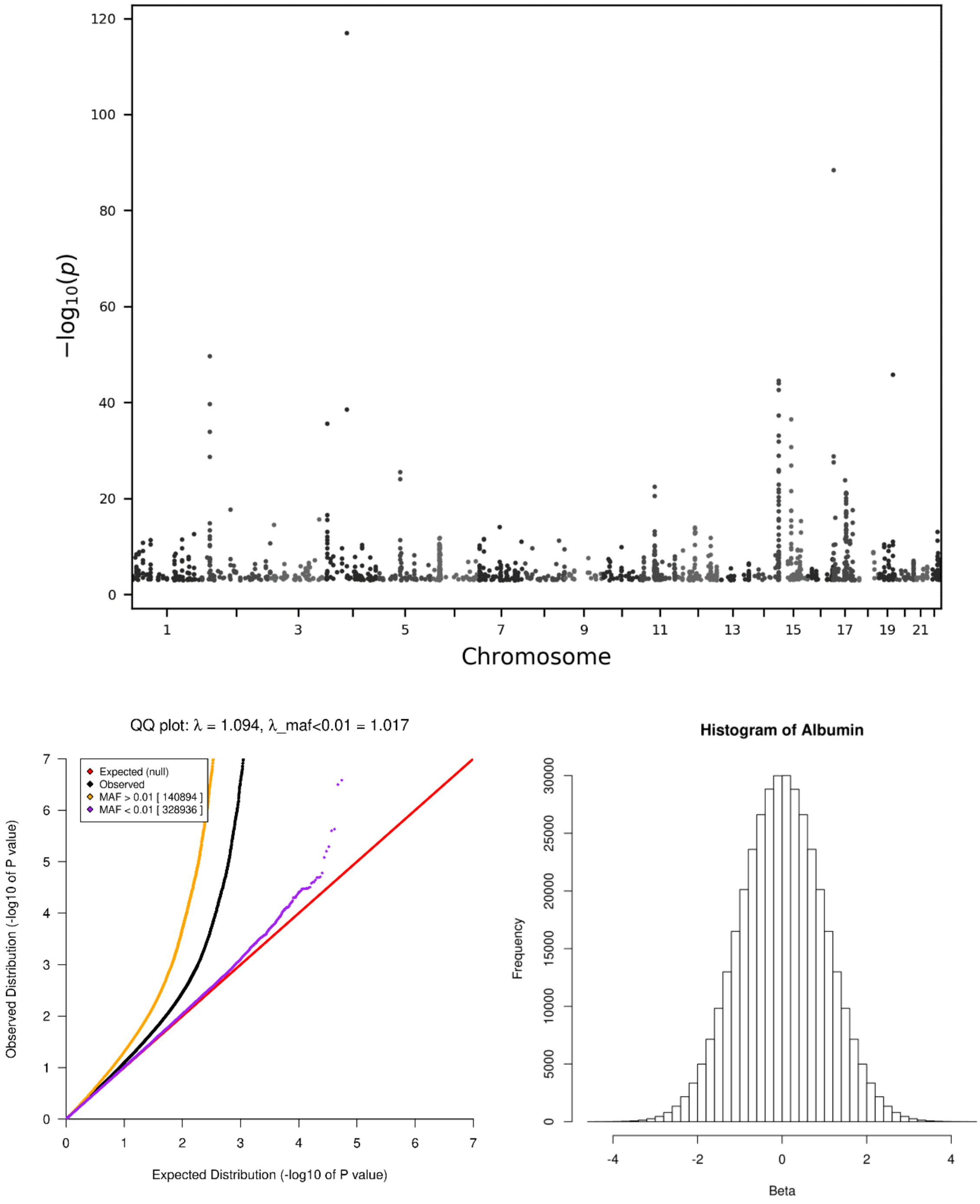

d) Standing height, SNV analysis, European ancestry (N=430,136)

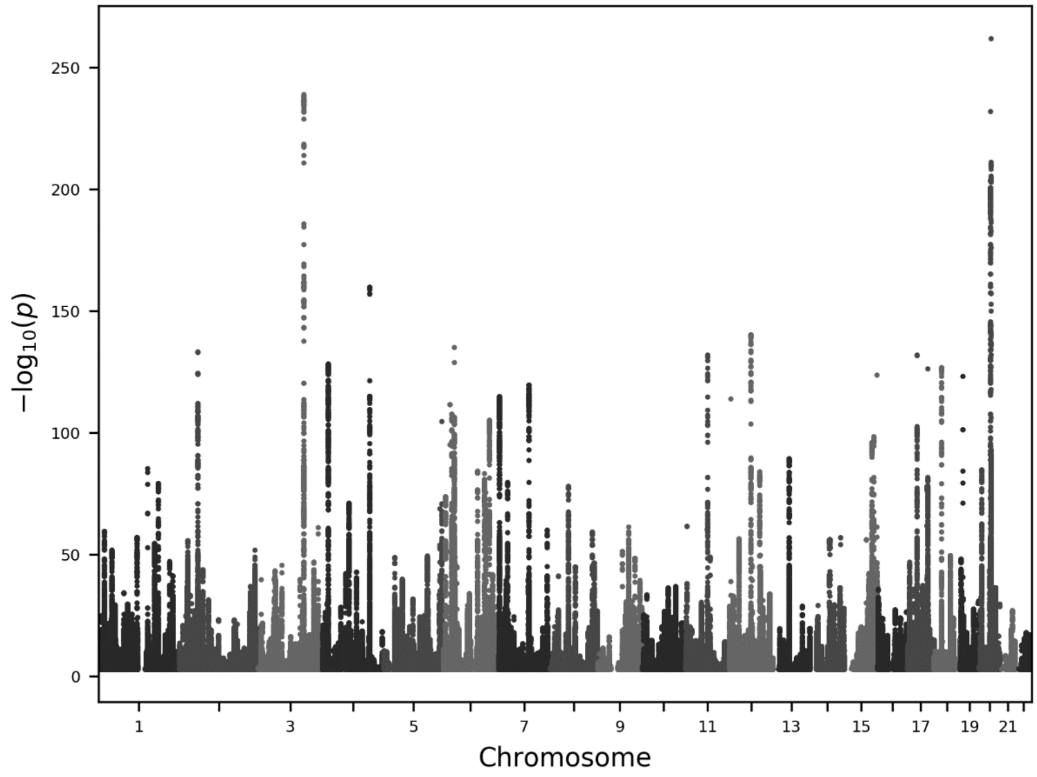


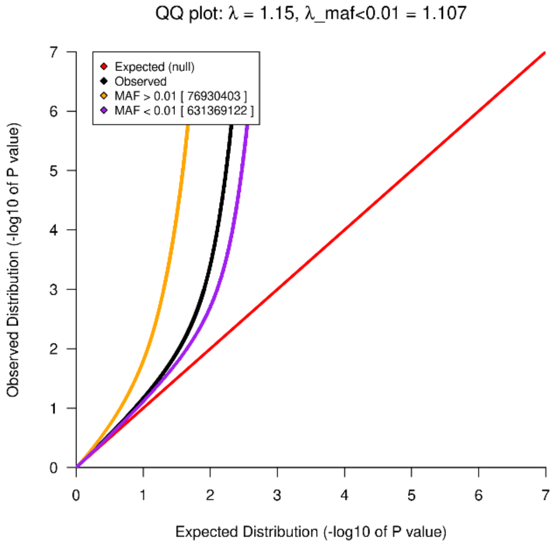


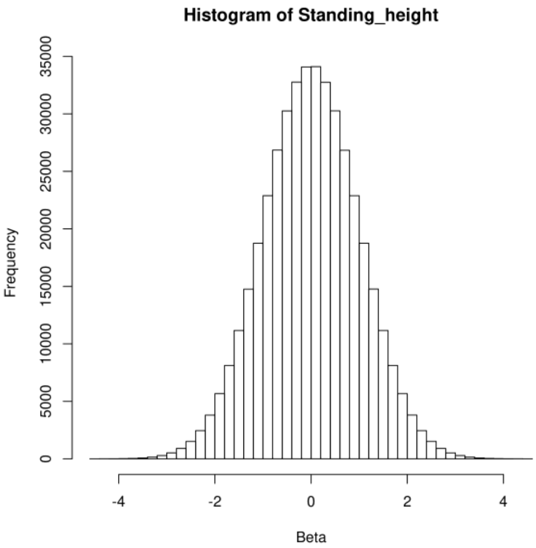

e) IGF-1 levels, SNV analysis, European ancestry (N=409,982)

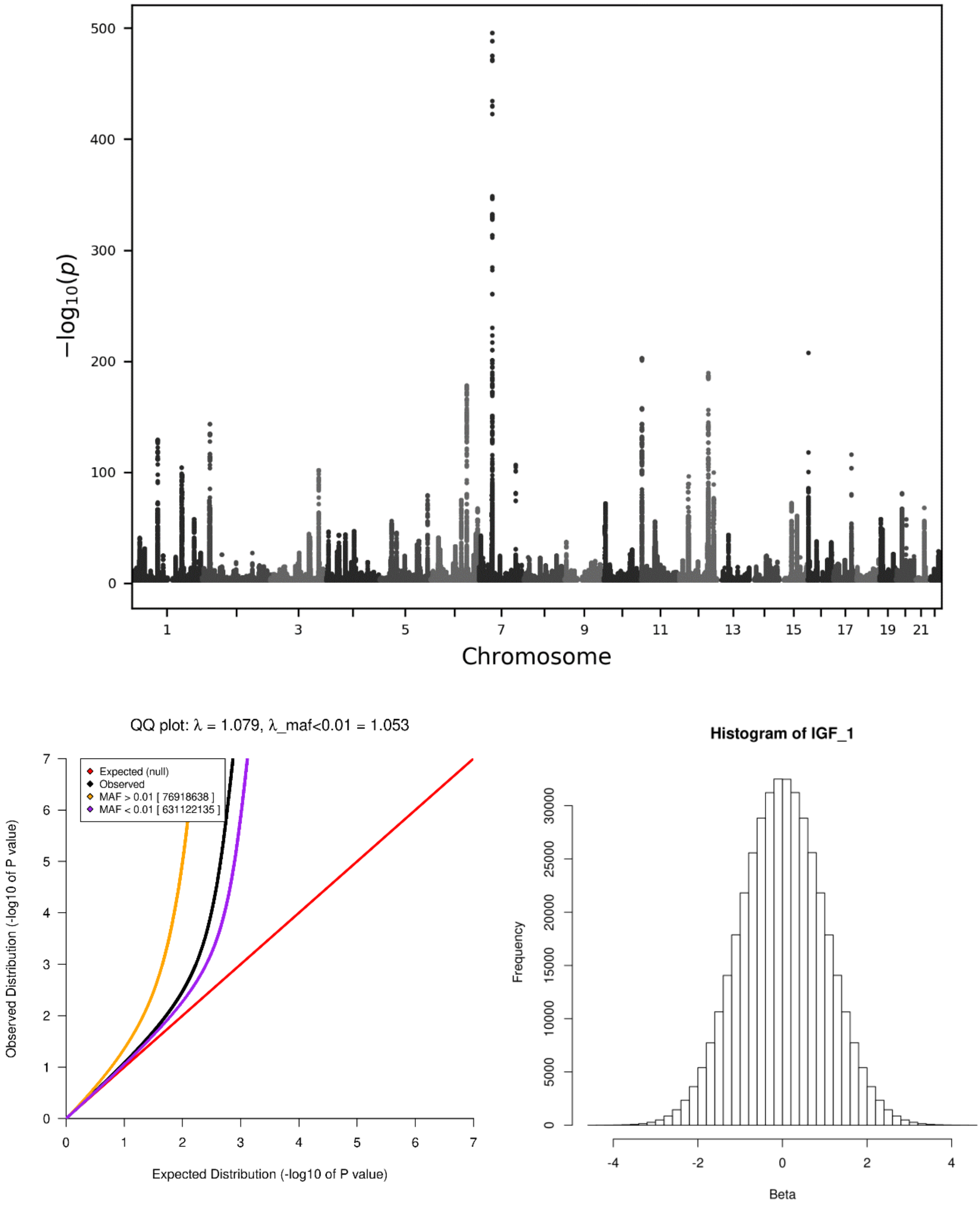

f) Mean corpuscular volume, SNV analysis, European ancestry, male sex (N=182,270)

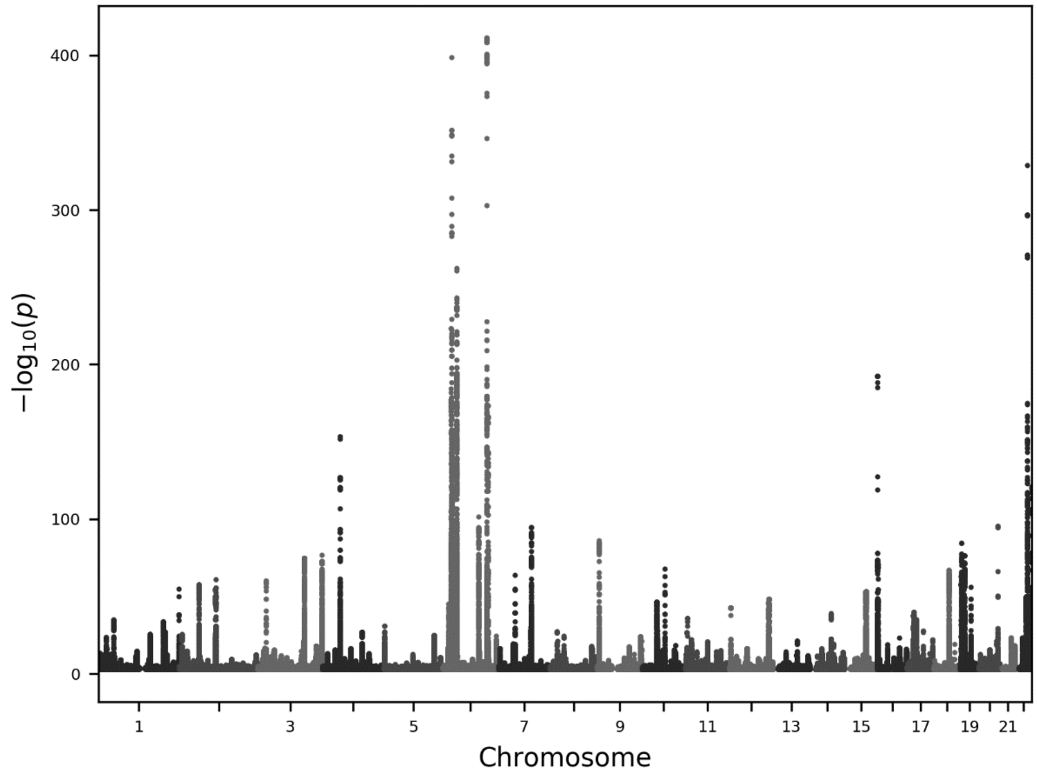


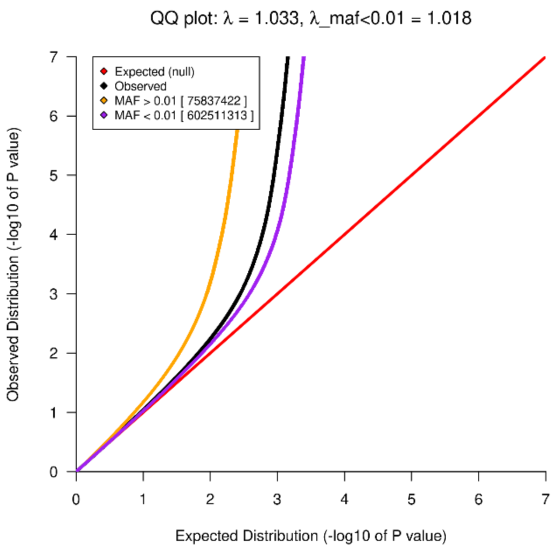


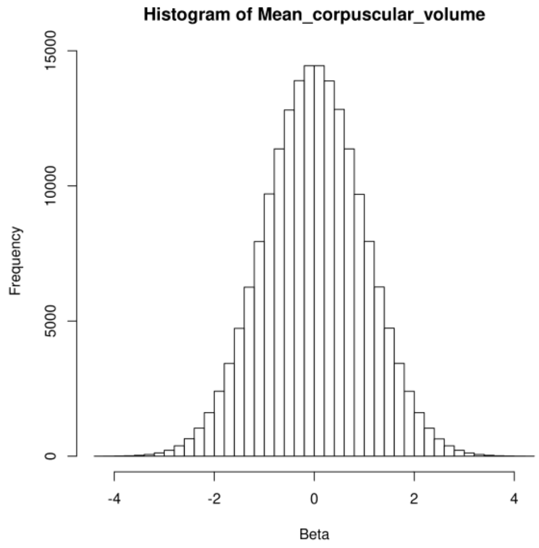

g) Age at menopause, structural variant analysis, European ancestry, female sex (N=141,129)

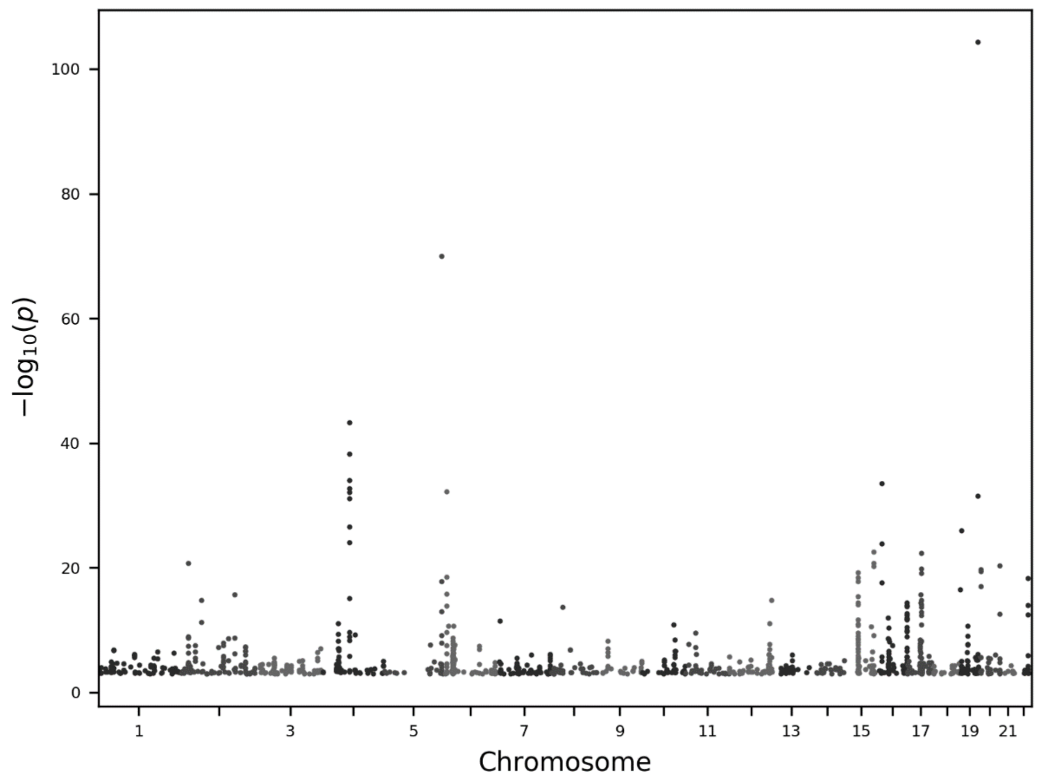


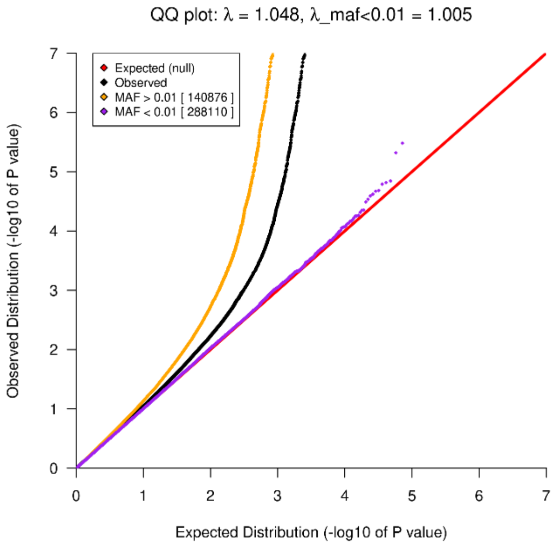


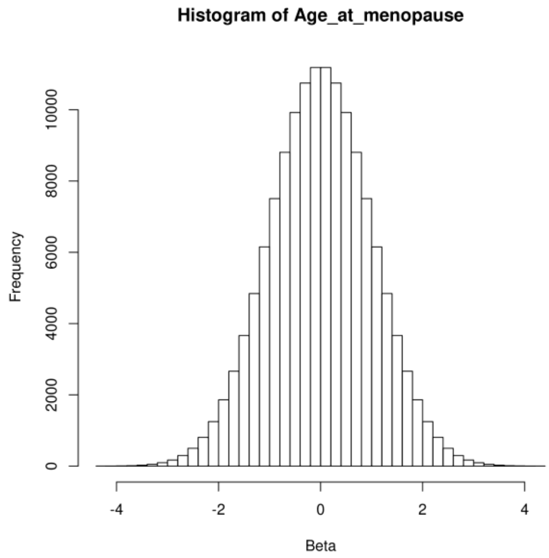

h) Non-high density lipoprotein, structural variant analysis, European ancestry (N=378,146)

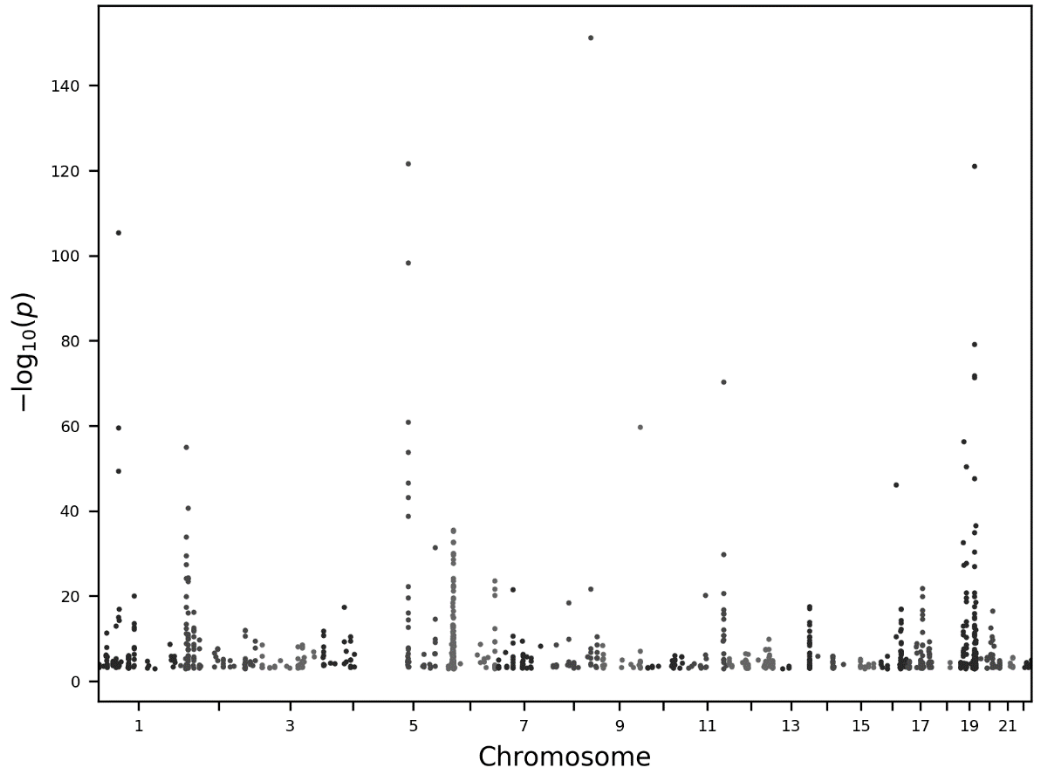


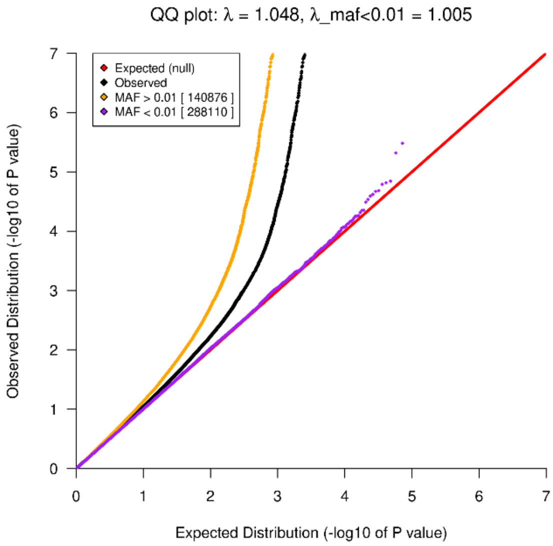


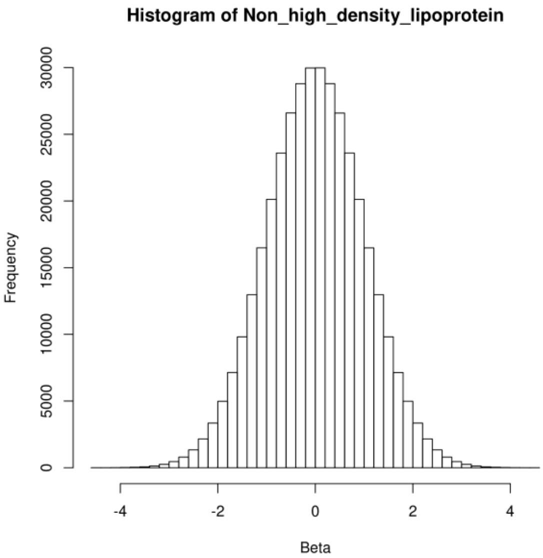

i) Non-high density lipoprotein, SNV analysis, African ancestry (N=8,359)

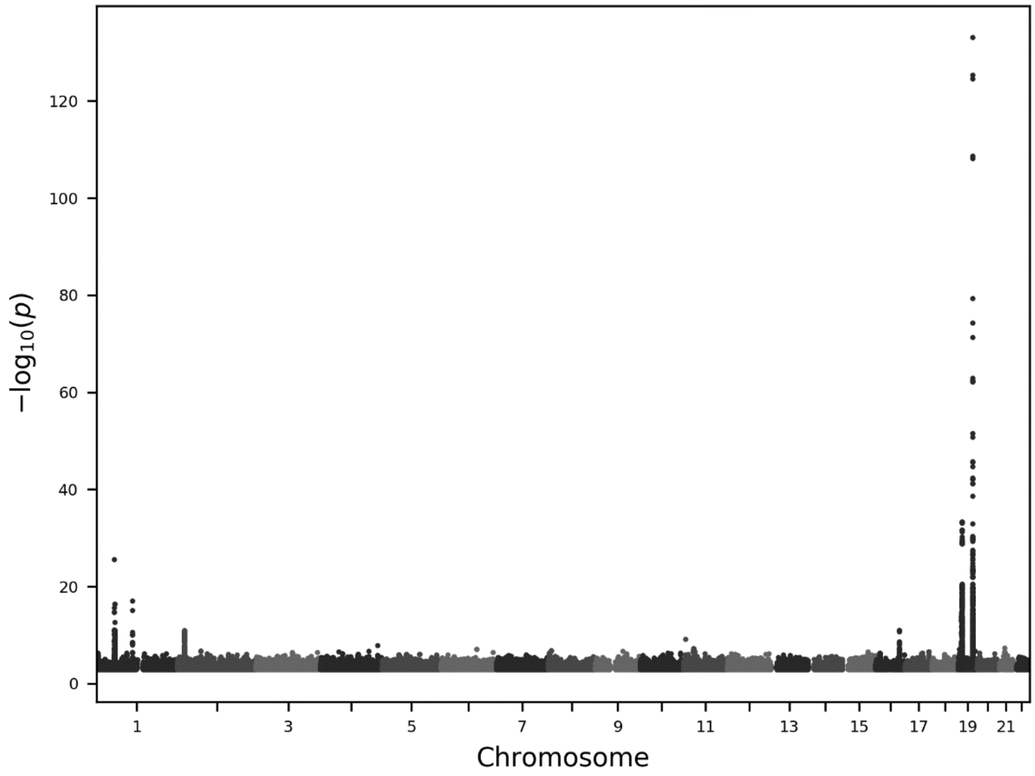


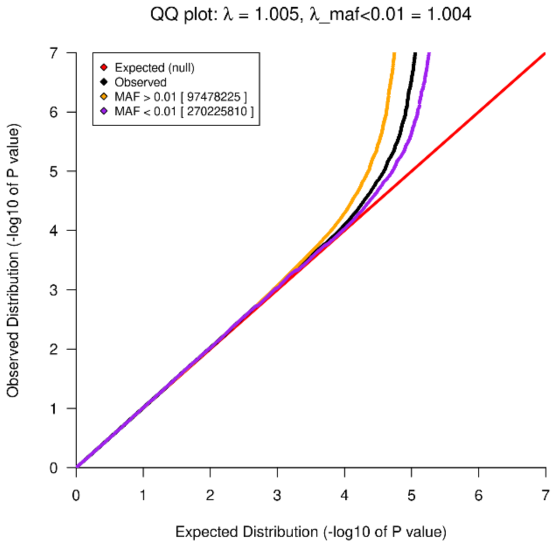


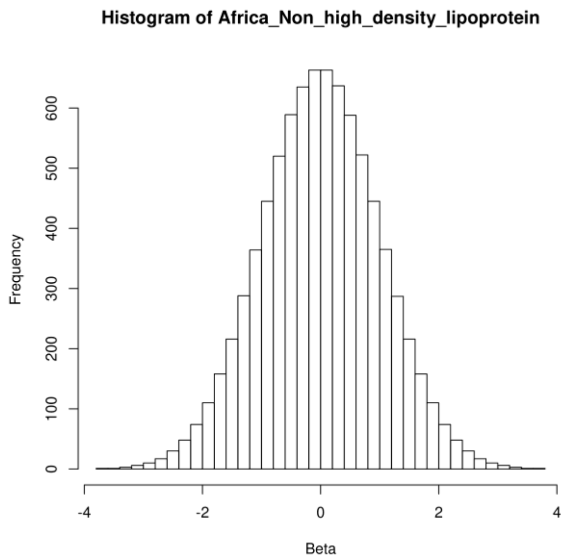

j) Hemoglobin concentration, SNV analysis, Asian ancestry (N=8,842)

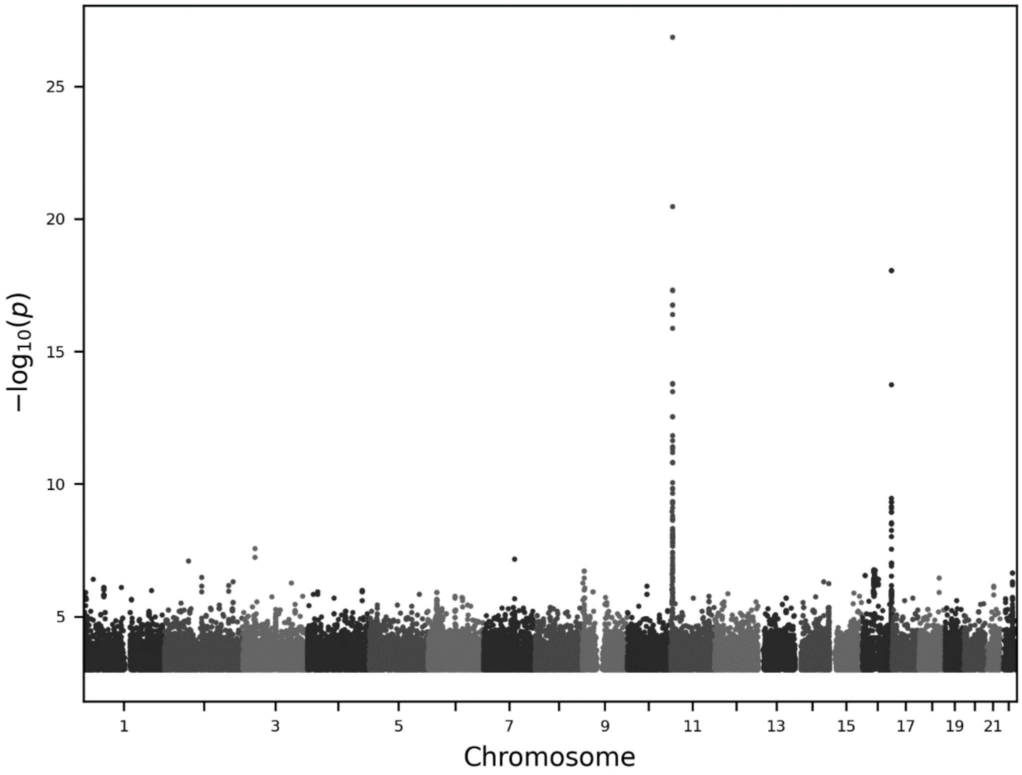


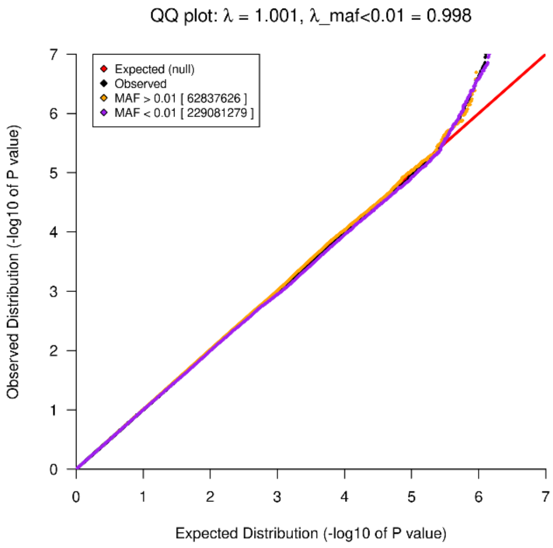


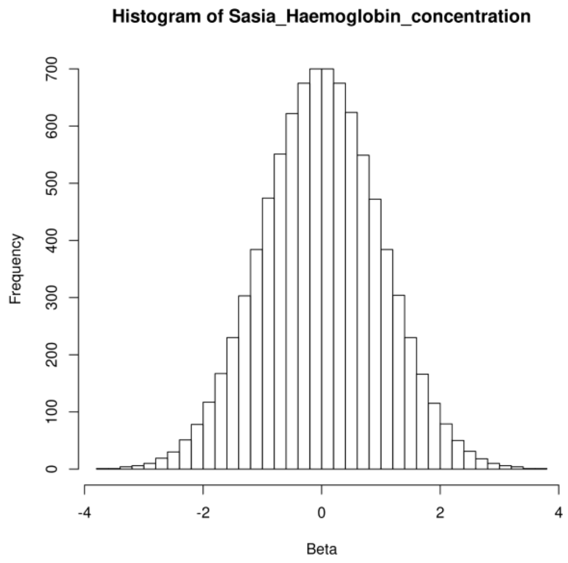

k) Age at menarche, SNV analysis, European ancestry (N=226,436)

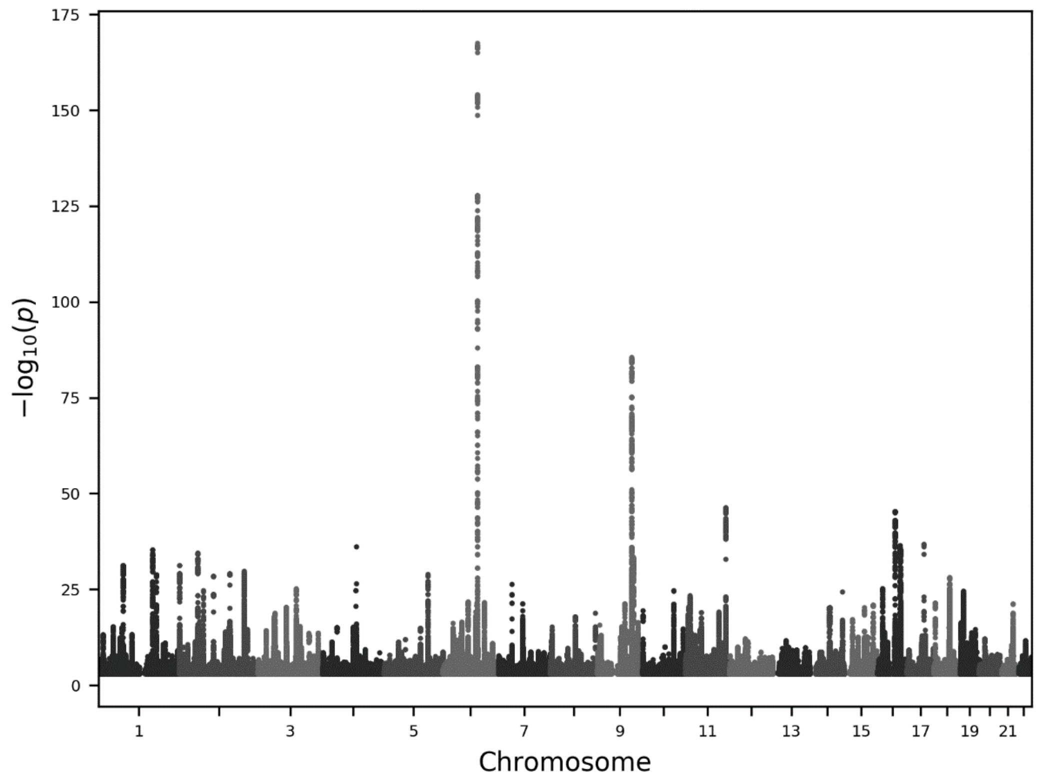


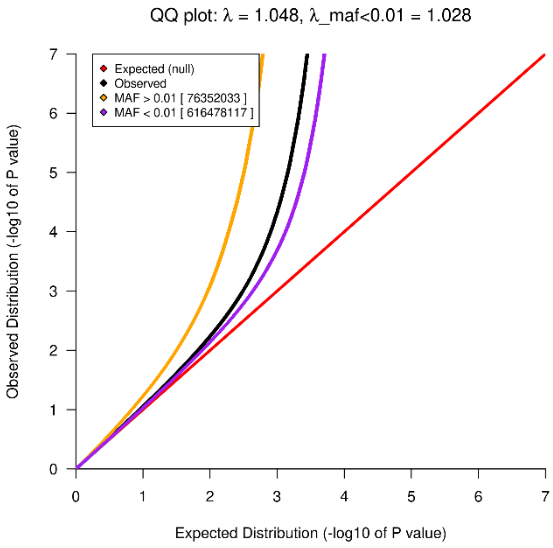


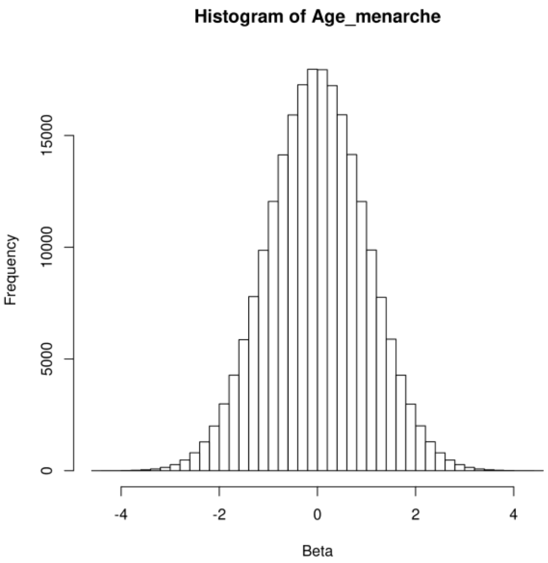

l) Urate levels, SNV analysis, European ancestry (N=411,640)

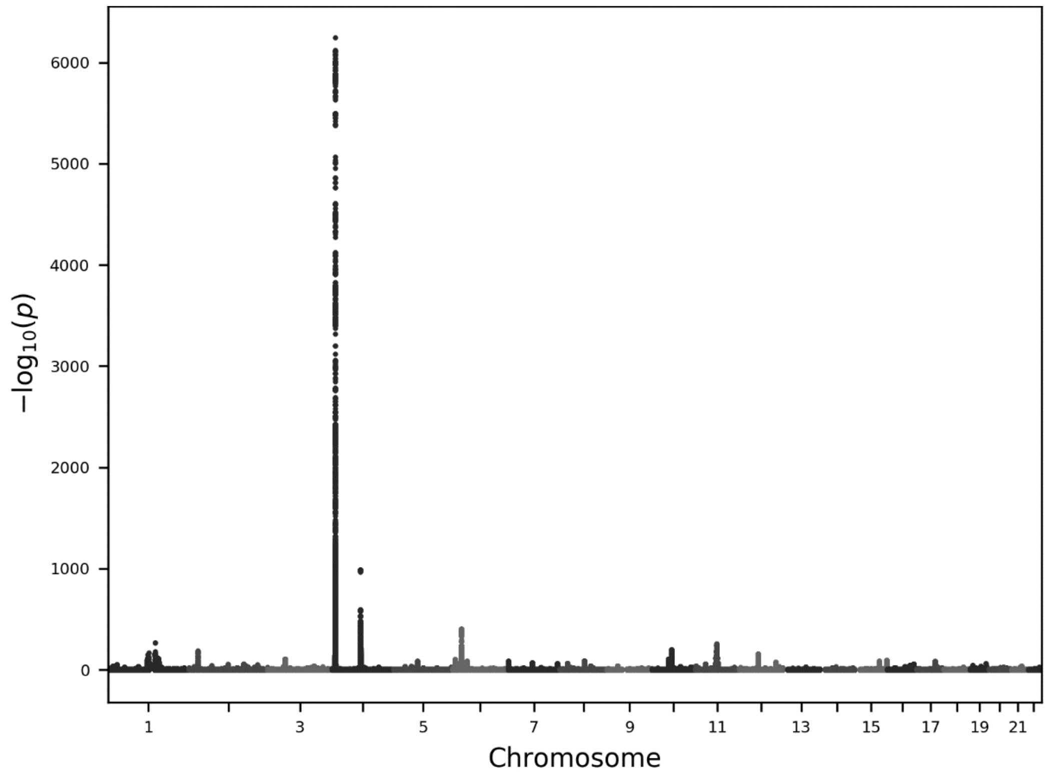


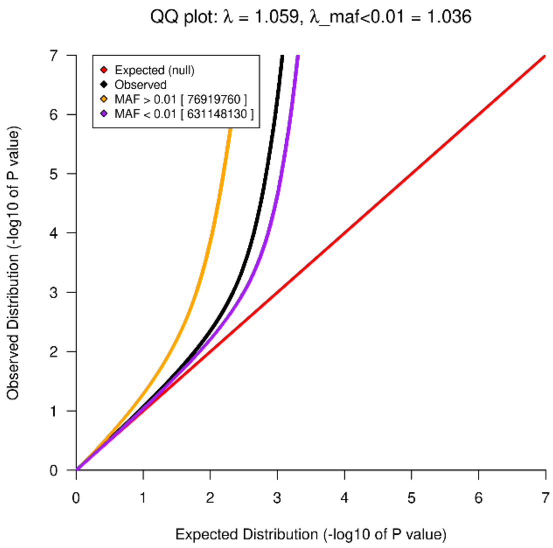


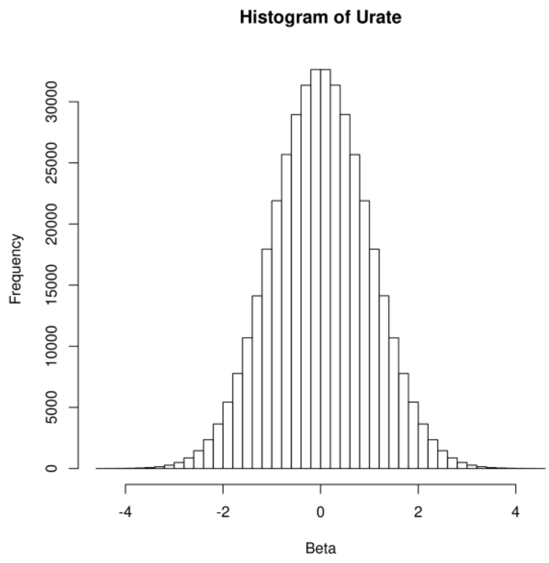

m) Glycine, metabolomics analysis, European ancestry (N=411,640)

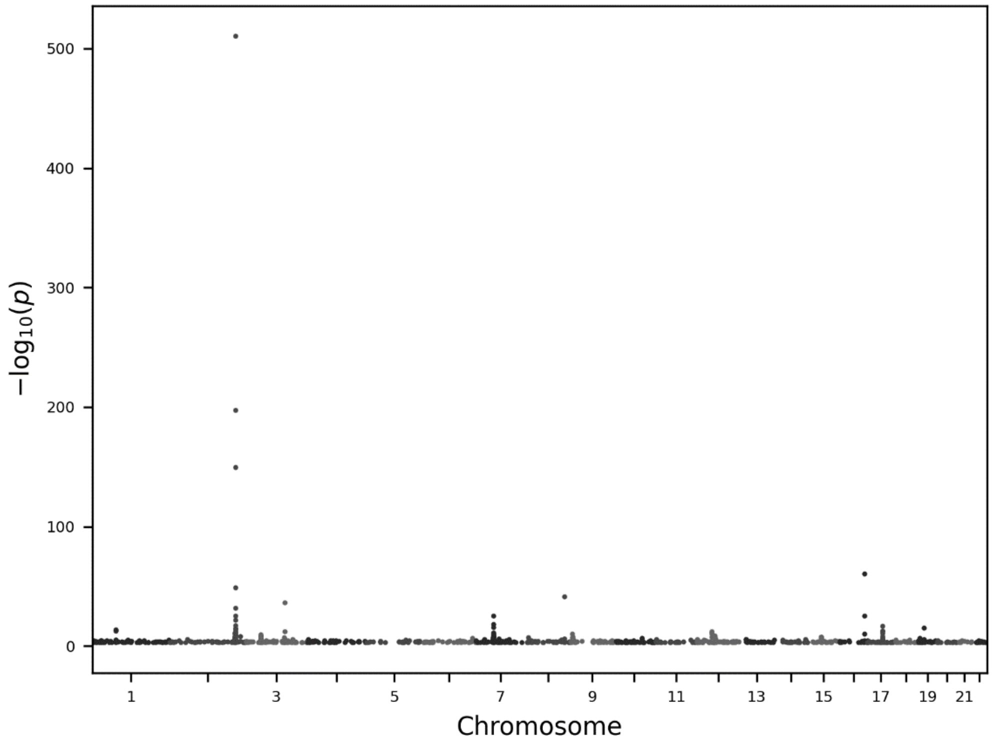


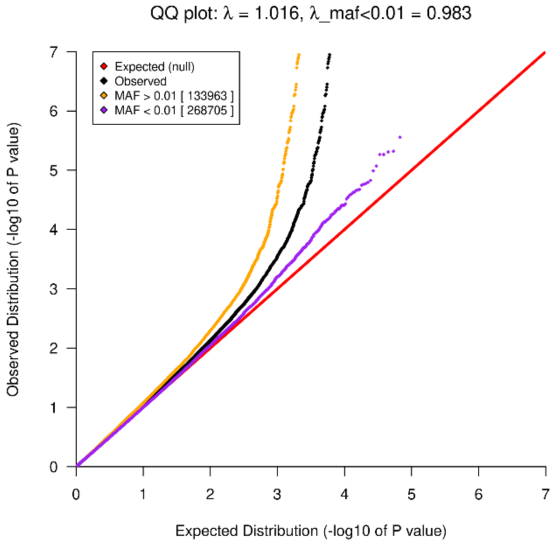


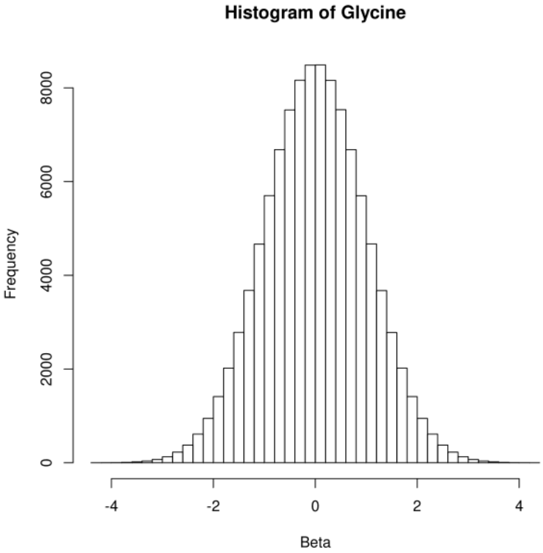

*Fig. S26 Manhattan plots and quantile-quantile (QQ) plots for case-control phenotypes with significant results reported in this manuscript. For Manhattan plots, the x-axis represents chromosome locations and the y-axis shows the –log10 significance levels of the associations. For QQ plots, the inflation (λ) is shown in the title of each graph, for all variants and for rare variants only (λ_maf<0.01)*

**a)** Hereditary ataxia, microsatellite analysis, European ancestry (Ncases=335, Ncontrols=430,603)

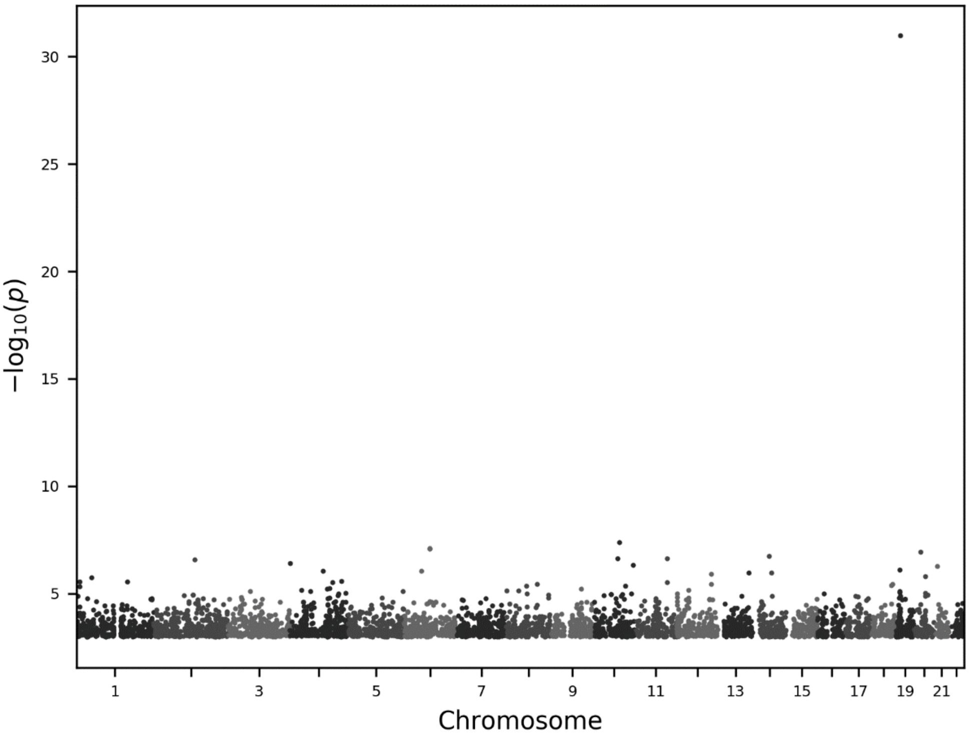


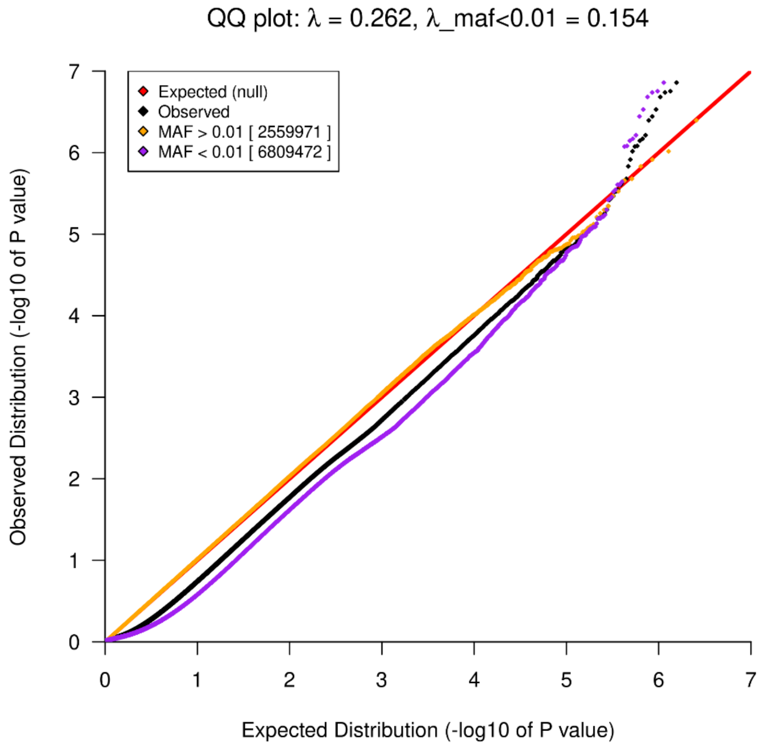

**b)** Myotonic disorders, microsatellite analysis, European ancestry (Ncases=99, Ncontrols=430,839)

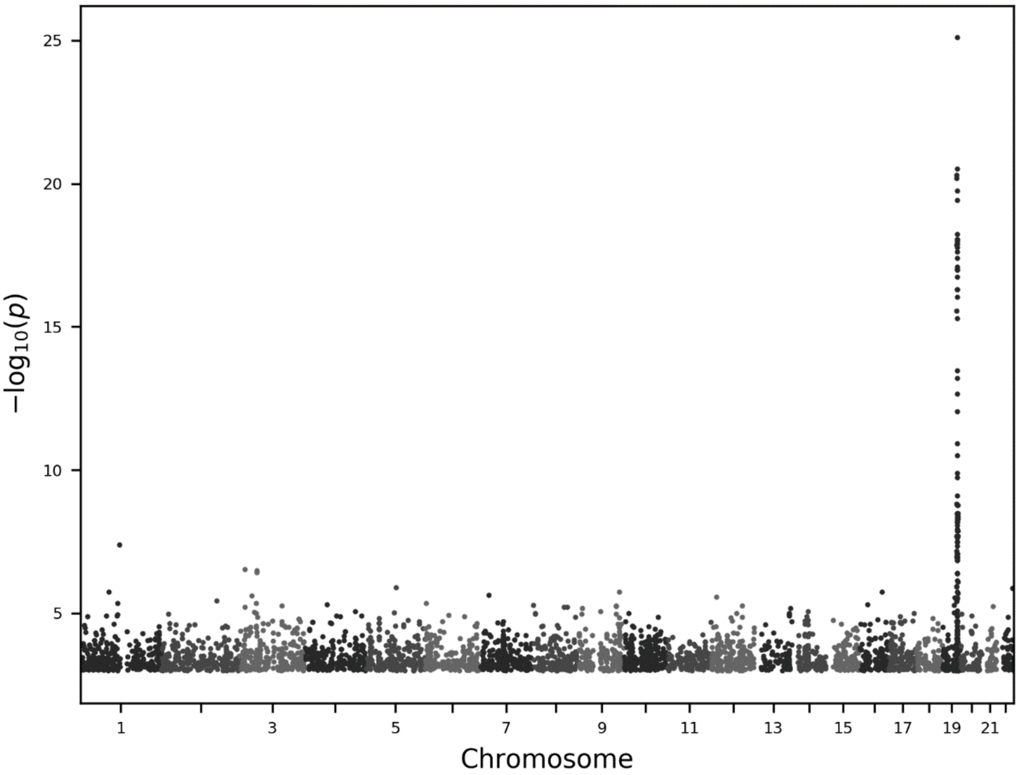


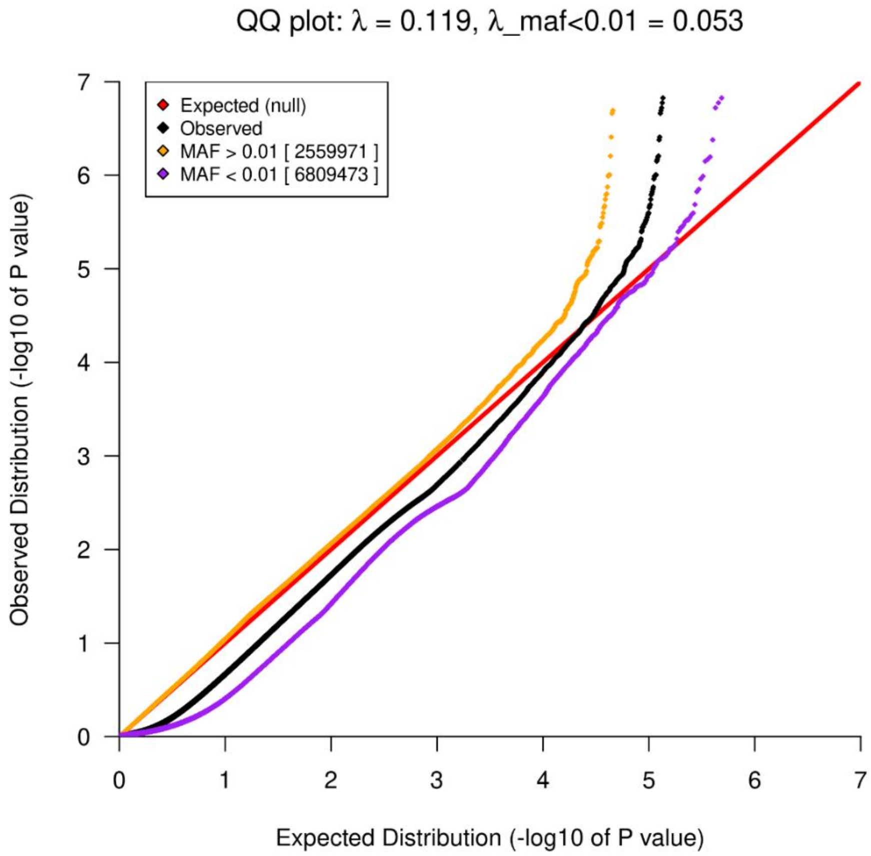

**c)** Gout, SNV analysis, European ancestry (Ncases=16,353, Ncontrols=414,694)

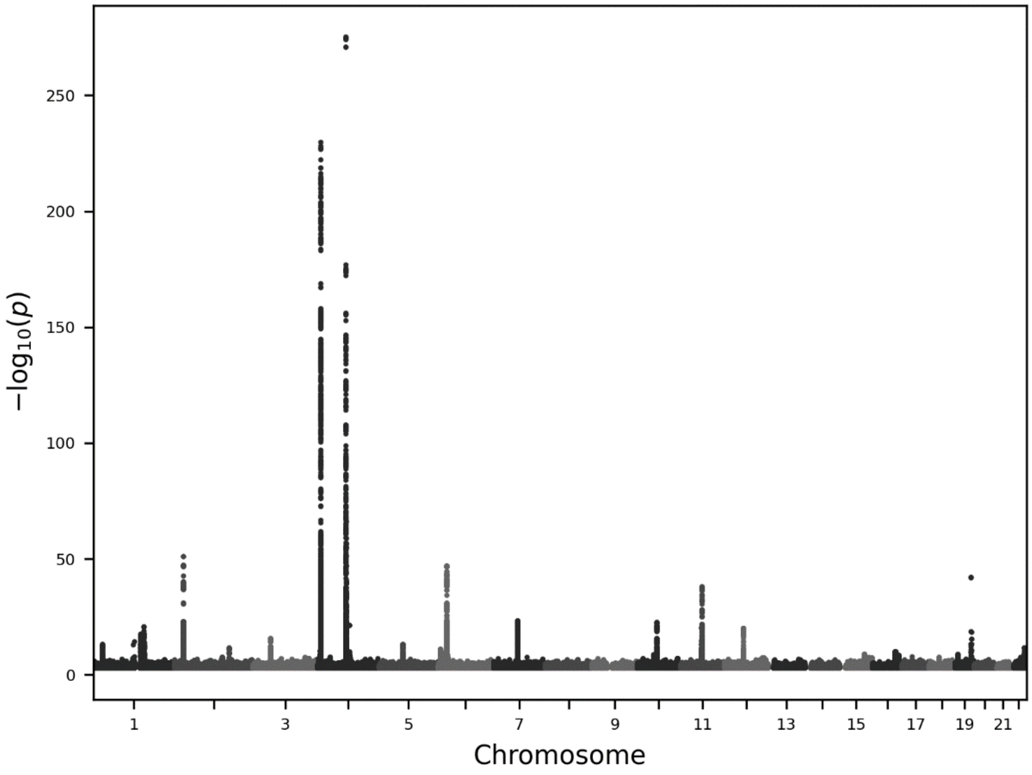


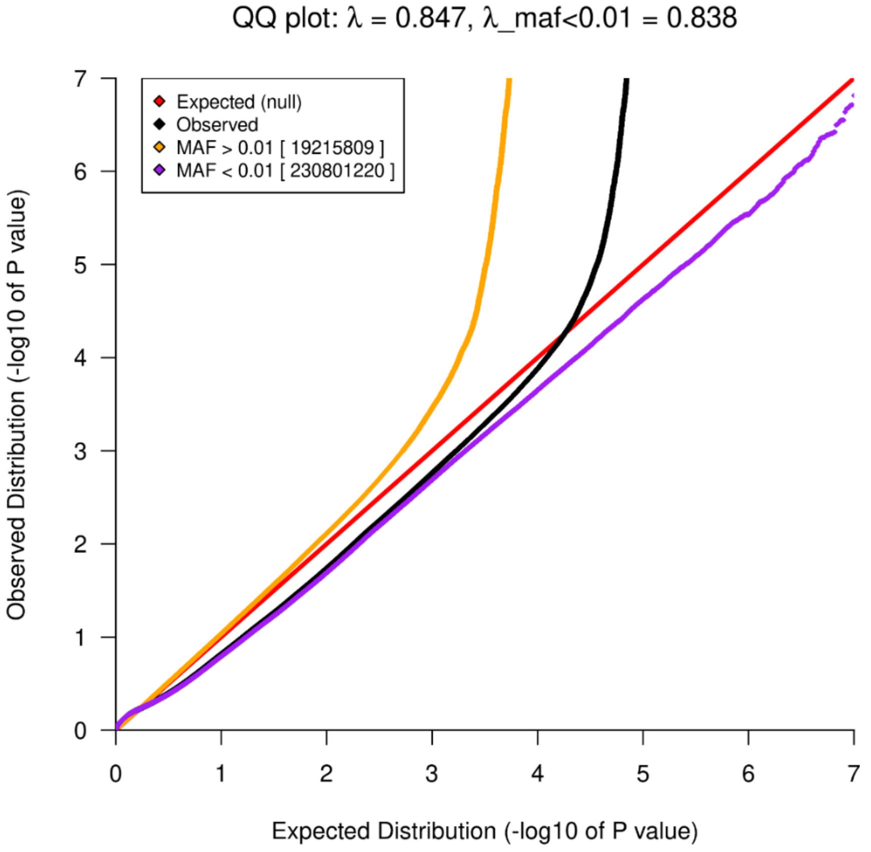

A)

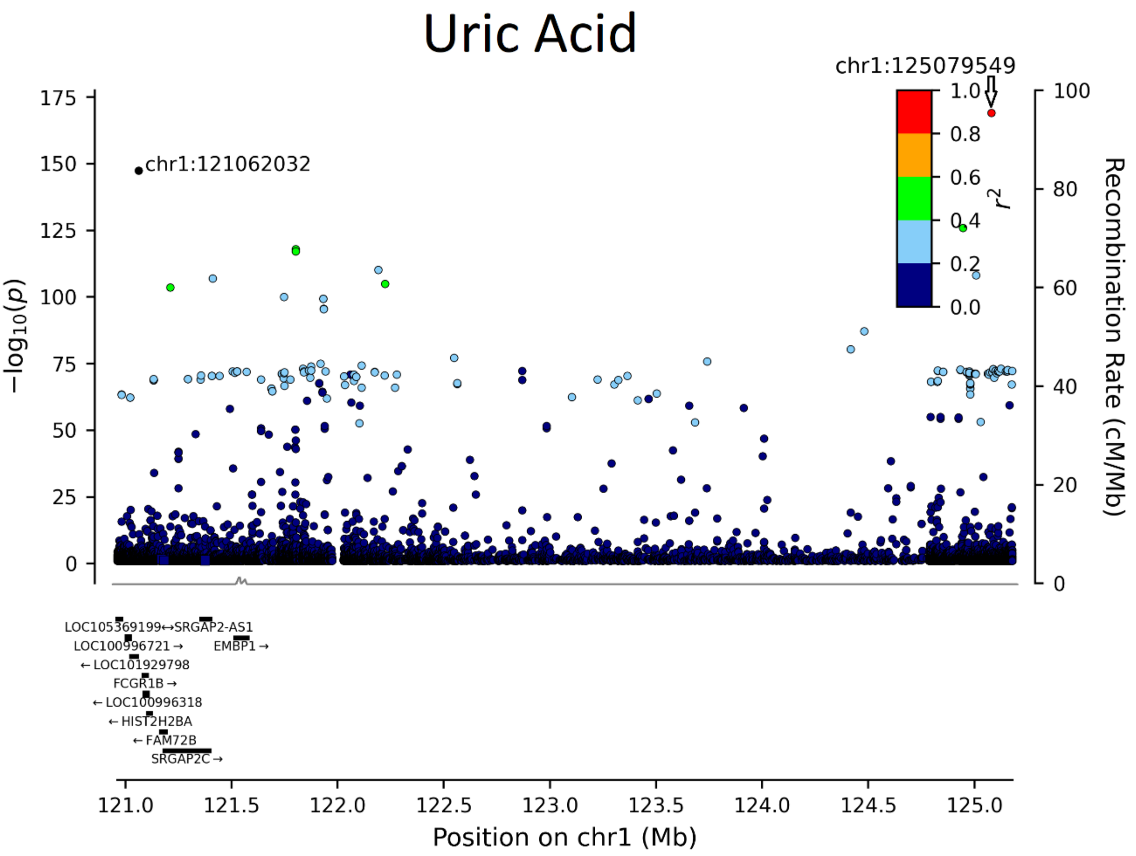


B)

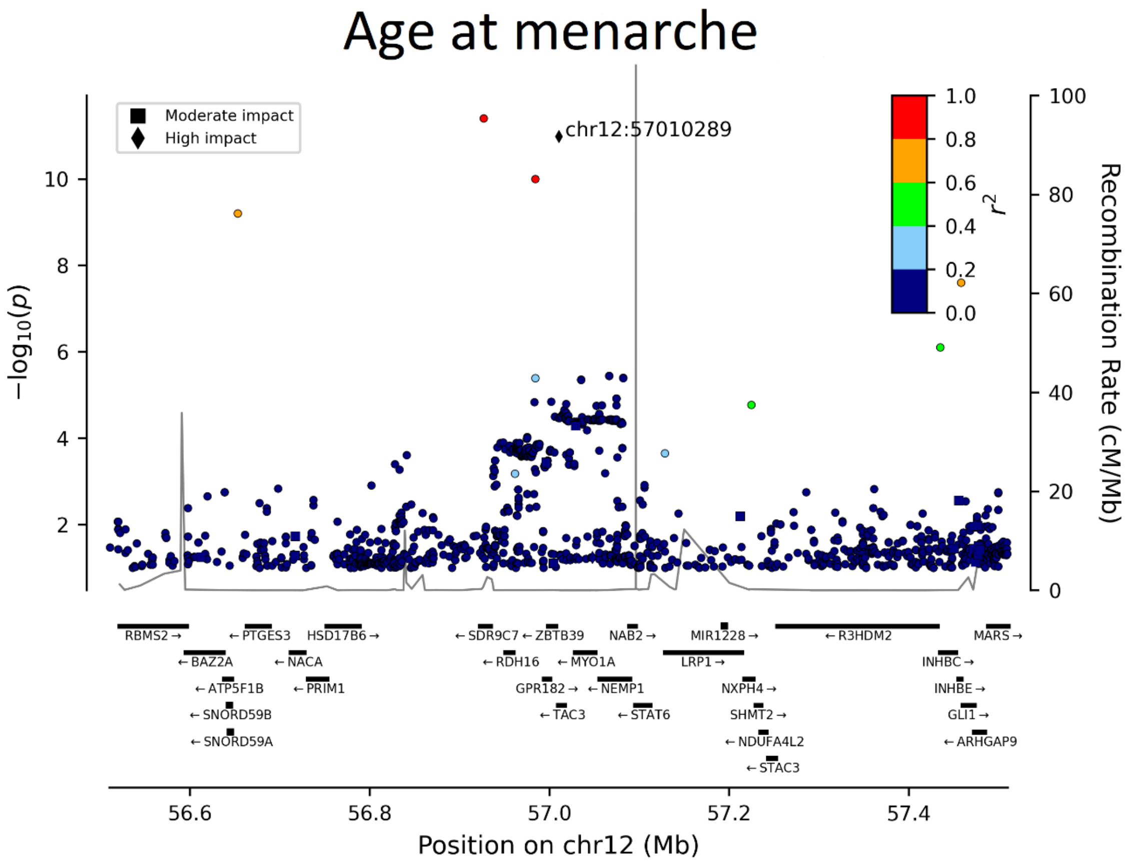


*Fig. S27 Locus plot for A) Uric acid and B) Age at menarche associations.*

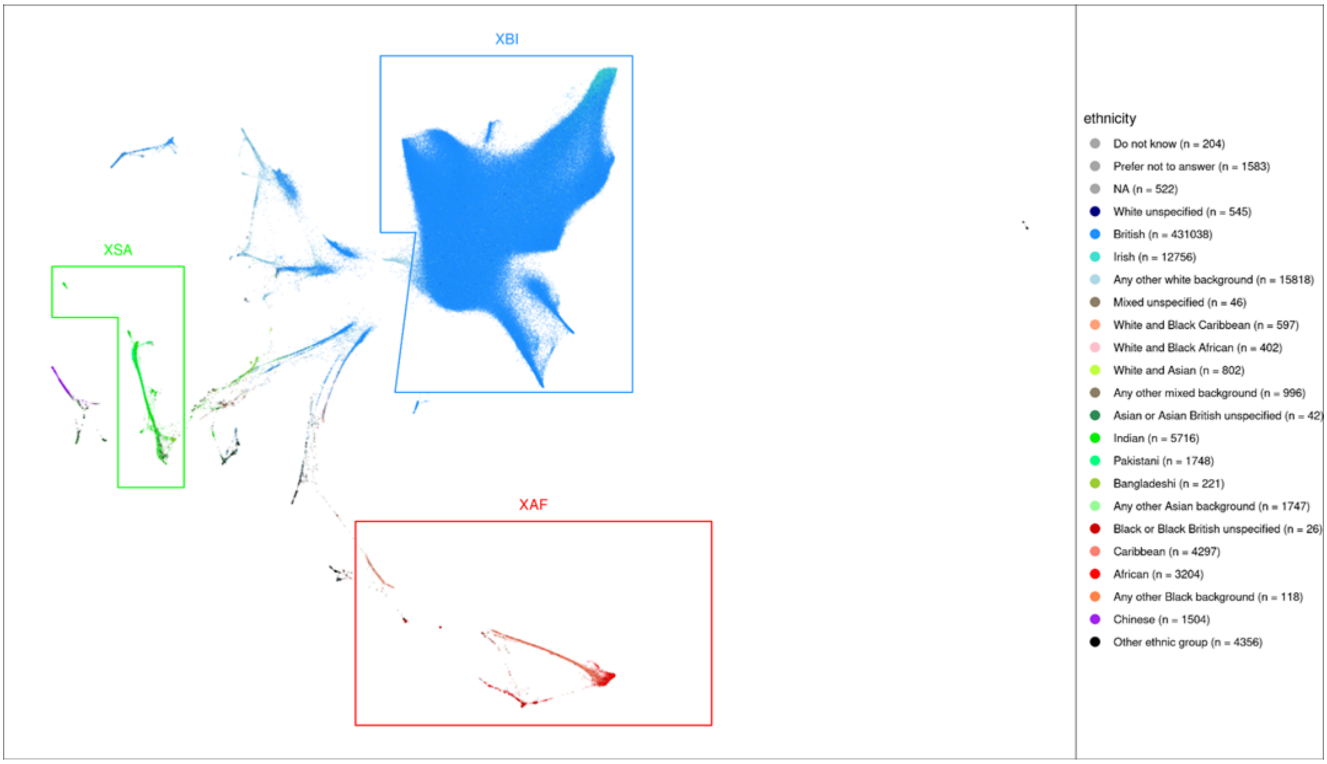


*Fig. S28 UMAP and ethnicity. 40 genetic principal components provided by UKB reduced to a latent space of 2 dimensions using UMAP (x and y axes). Individuals are colored according to self-identified ethnicity. The regions defined to delineate the three cohorts XAF, XBI, and XSA are indicated.*

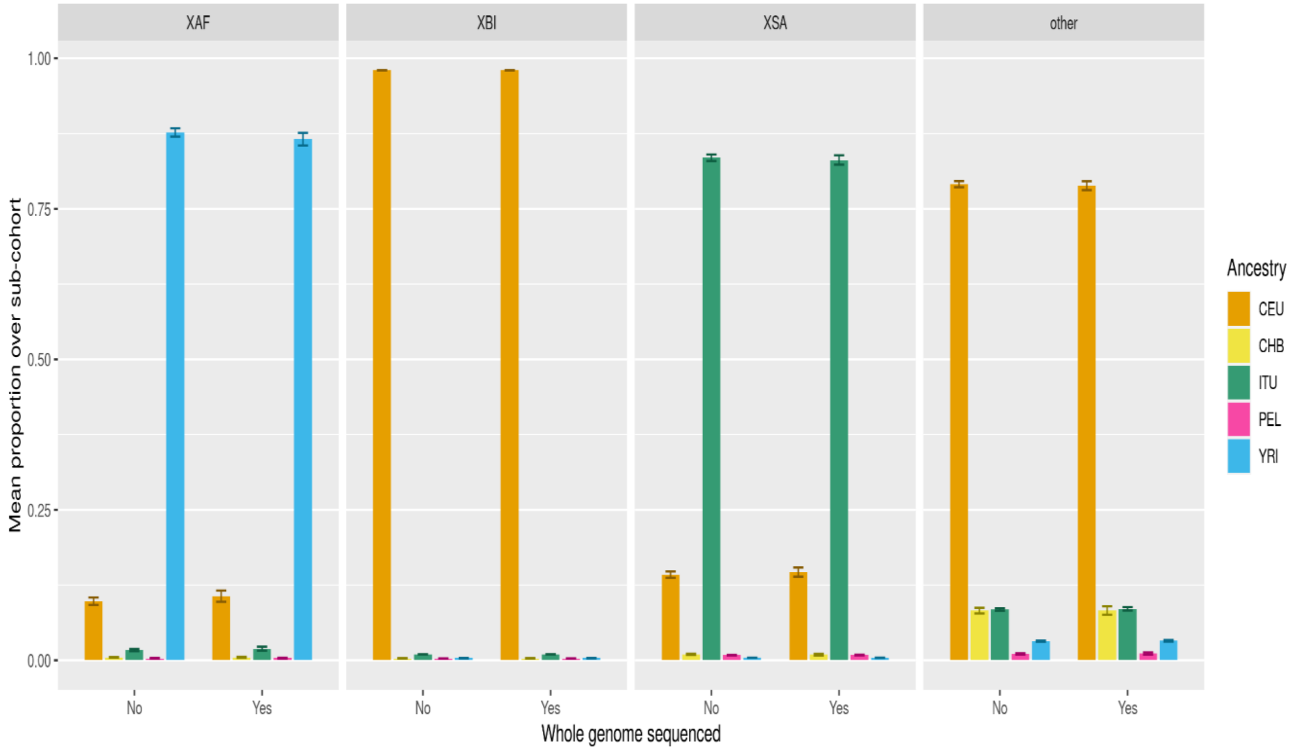


*Fig. S29 Cohort mean ADMIXTURE. Mean proportion of each of five 1000 Genome Project ancestry components assigned by ADMIXTURE (columns). Error bars represent 99.9% confidence intervals. CEU (Northern Europeans from Utah), CHB (Han Chinese in Beijing), ITU (Indian Telugu in the UK), PEL (Peruvians in Lima), and YRI (Yoruba in Ibadan, Nigeria).*

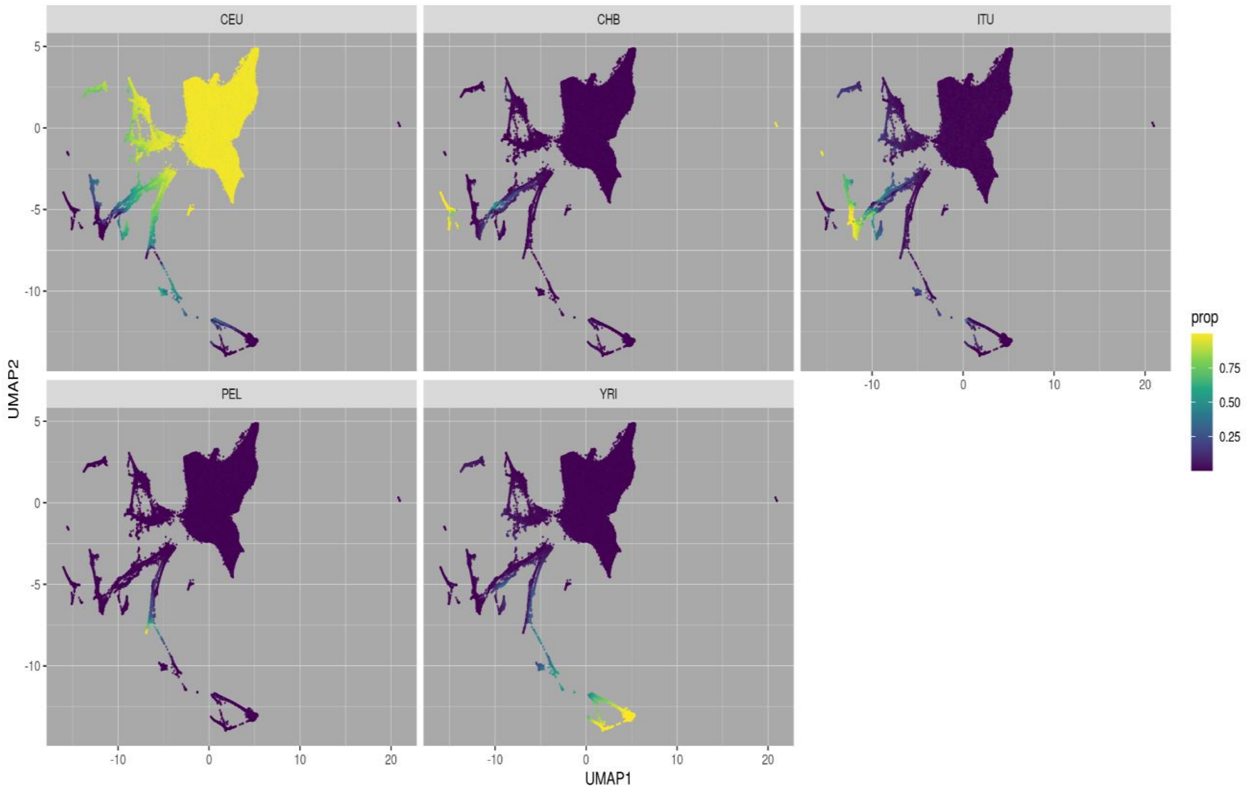


*Fig. S30 UMAP ADMIXTURE. 40 genetic principal components provided by UKB reduced to a latent space of 2 dimensions using UMAP (x and y axes). Individuals are colored according to proportion of ancestry assigned by supervised ADMIXTURE from five 1000GP training populations (facet headings): CEU (Northern Europeans from Utah), CHB (Han Chinese in Beijing), ITU (Indian Telugu in the UK), PEL (Peruvians in Lima), and YRI (Yoruba in Ibadan, Nigeria).*

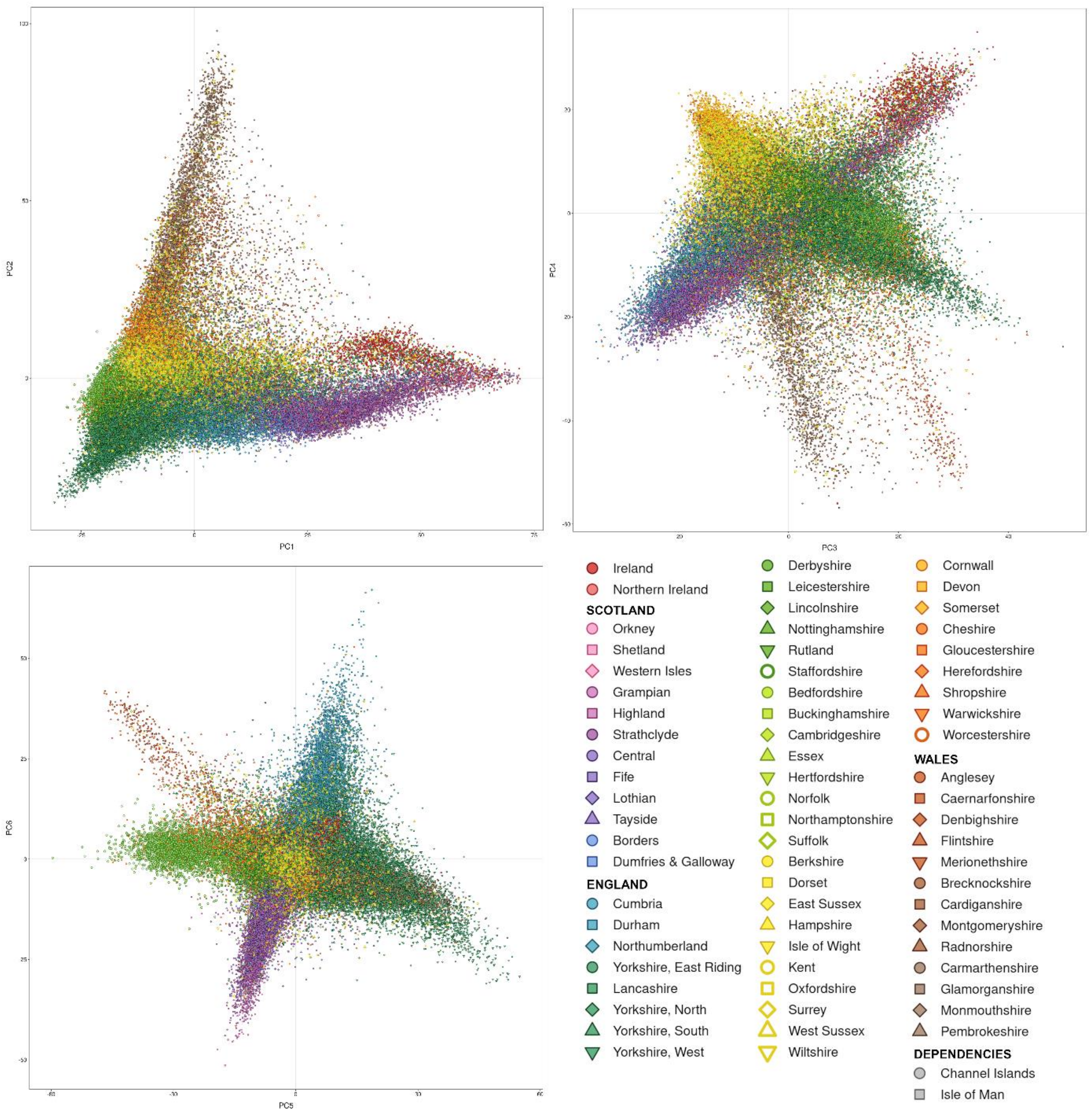


*Fig. S31 The first six principal components of the XBI cohort, plots show PC1 vs PC2, PC3 vs PC4 and PC5 vs PC6. Points represent individuals, colored by place of birth. To show geographic structure in the UK more clearly, we do not show individuals who report being born in urban areas with many internal migrants (Tyne & Wear, Merseyside, Greater Manchester, West Midlands, Bristol, London) or places outside the British-Irish Isles.*

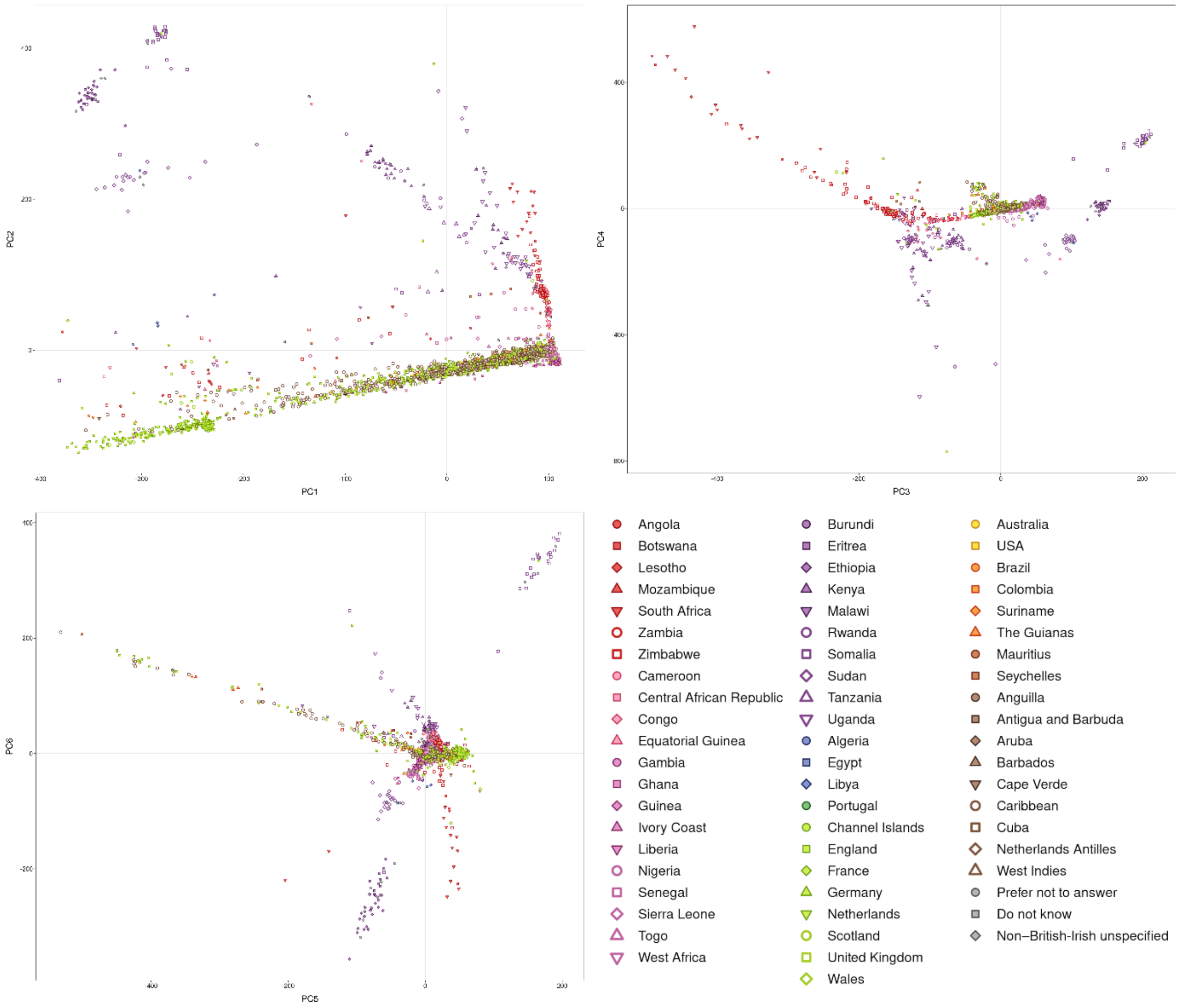


*Fig. S32 The first six principal components of the XAF cohort, plots show PC1 vs PC2, PC3 vs PC4 and PC5 vs PC6. Points represent individuals, colored by place of birth.*

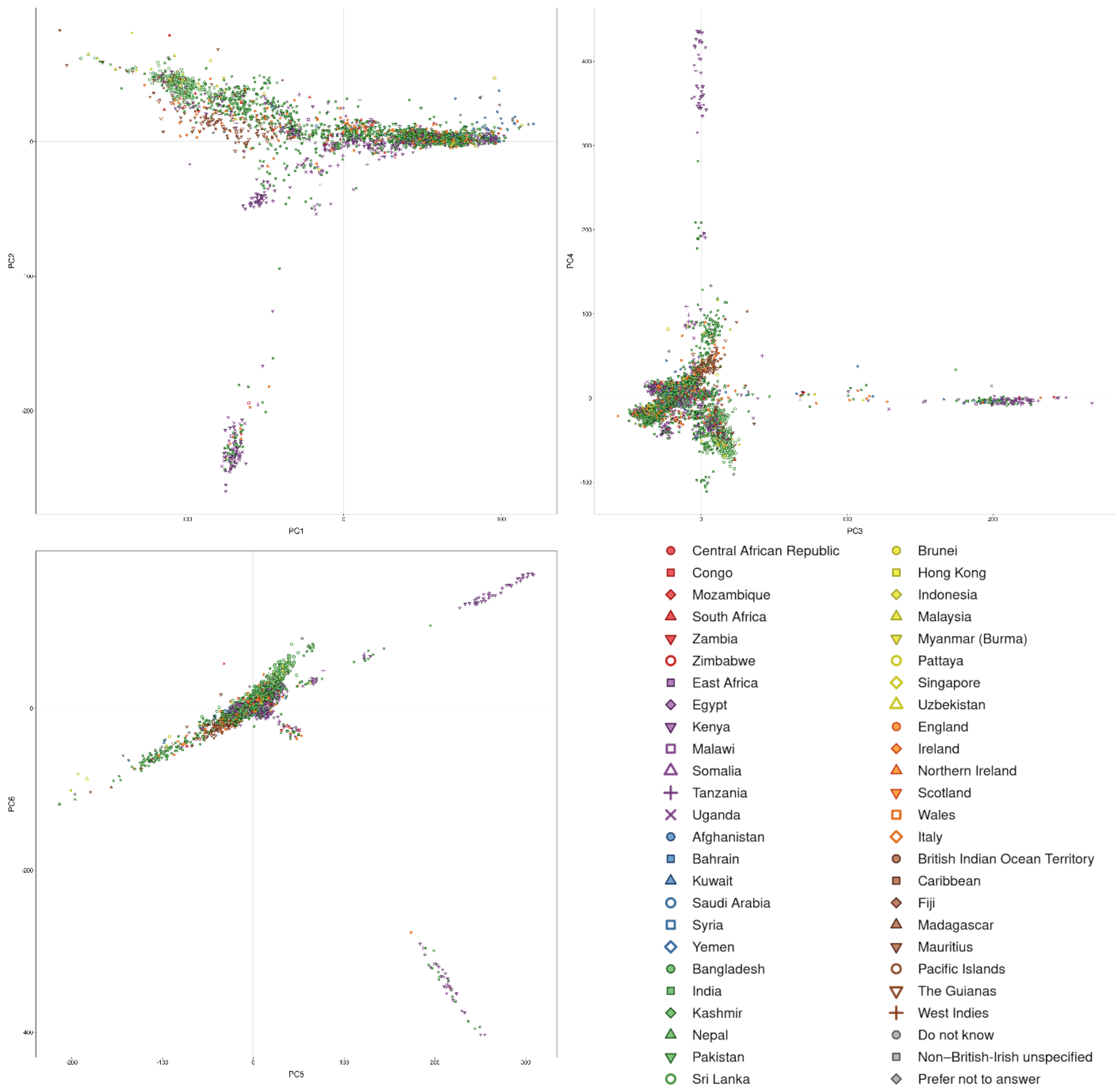


*Fig. S33 The first six principal components of the XSA cohort, plots show PC1 vs PC2, PC3 vs PC4 and PC5 vs PC6. Points represent individuals, colored by place of birth.*

#### Supplementary Tables

A) SNP+Indel

| GIAB sample | #Variants | GATK Sensitivity | GATK Precision | GATK F1-score | GraphTyper Sensitivity | GraphTyper Precision | GraphTyper F1-score |
|---|---|---|---|---|---|---|---|
| HG001 | 30,717 | 98.09% | 98.90% | 98.49% | 98.97% | 99.29% | 99.13% |
| HG002 | 29,802 | 98.14% | 99.03% | 98.59% | 98.84% | 99.36% | 99.10% |
| HG003 | 28,379 | 98.16% | 99.10% | 98.63% | 99.02% | 99.21% | 99.11% |
| HG004 | 28,539 | 98.11% | 99.02% | 98.56% | 99.03% | 99.48% | 99.26% |
| HG005 | 26,846 | 98.47% | 99.02% | 98.74% | 99.08% | 99.48% | 99.28% |
| HG006 | 27,546 | 98.77% | 99.11% | 98.94% | 99.22% | 99.28% | 99.25% |
| HG007 | 28,798 | 98.63% | 99.21% | 98.92% | 99.14% | 99.29% | 99.21% |
| Average | 28,661 | 98.34% | 99.06% | 98.70% | 99.04% | 99.34% | 99.19% |

B) SNP

| GIAB sample | #Variants | GATK Sensitivity | GATK Precision | GATK F1-score | GraphTyper Sensitivity | GraphTyper Precision | GraphTyper F1-score |
|---|---|---|---|---|---|---|---|
| HG001 | 26,377 | 99.50% | 99.07% | 99.28% | 99.63% | 99.29% | 99.46% |
| HG002 | 25,747 | 99.45% | 99.09% | 99.27% | 99.46% | 99.36% | 99.41% |
| HG003 | 24,450 | 99.43% | 99.19% | 99.31% | 99.56% | 99.20% | 99.38% |
| HG004 | 24,428 | 99.47% | 99.16% | 99.31% | 99.60% | 99.48% | 99.54% |
| HG005 | 23,465 | 99.60% | 99.14% | 99.37% | 99.44% | 99.49% | 99.46% |
| HG006 | 24,226 | 99.63% | 99.18% | 99.40% | 99.61% | 99.27% | 99.44% |
| HG007 | 25,257 | 99.59% | 99.30% | 99.44% | 99.53% | 99.29% | 99.41% |
| Average | 24,850 | 99.52% | 99.16% | 99.34% | 99.55% | 99.34% | 99.44% |

C) Indel

| GIAB sample | #Variants | GATK Sensitivity | GATK Precision | GATK F1-score | GraphTyper Sensitivity | GraphTyper Precision | GraphTyper F1-score |
|---|---|---|---|---|---|---|---|
| HG001 | 4,340 | 89.46% | 97.30% | 93.21% | 94.94% | 99.59% | 97.21% |
| HG002 | 4,055 | 89.81% | 98.42% | 93.92% | 94.85% | 99.42% | 97.08% |
| HG003 | 3,929 | 90.26% | 98.12% | 94.03% | 95.61% | 99.54% | 97.54% |
| HG004 | 4,111 | 89.93% | 97.68% | 93.64% | 95.59% | 99.43% | 97.47% |
| HG005 | 3,381 | 90.47% | 97.80% | 93.99% | 96.50% | 99.34% | 97.90% |
| HG006 | 3,320 | 92.45% | 98.35% | 95.31% | 96.34% | 99.65% | 97.97% |
| HG007 | 3,541 | 91.68% | 98.25% | 94.85% | 96.28% | 99.51% | 97.87% |
| Average | 3,811 | 90.58% | 97.99% | 94.14% | 95.73% | 99.50% | 97.58% |

*Table S1 Genome in a bottle (GIAB) v3.3.2 truth set comparison of GATK and GraphTyper in 500 random regions F1-score is the harmonic mean of Sensitivity and Precision. A) all variant types, B) SNPs only C) Indels only.*

A)

| Method | FDR | TP | #Variants |
|---|---|---|---|
| GATK | 9.97% | 17,140,110 | 19,038,309 |
| GraphTyper | 6.31% | 17,915,210 | 19,123,669 |
| GraphTyperHQ | 1.45% | 16,768,945 | 17,016,415 |

B)

| Method | ICPM | Non-ref consistency | Number of non-ref calls |
|---|---|---|---|
| GATK | 78.1 | 95.21% | 68,537,823 |
| GraphTyper | 70.3 | 95.81% | 70,442,413 |
| GraphTyperHQ | 11.8 | 99.22% | 63,556,940 |

*Table S2 A) Estimate of false discovery rate (FDR) and number of true positive (TP) variants among the 28 parent-offspring trios. The estimates are determined from the allele transmission ratios from parent to offspring. B) Genotype consistency across among the 14 monozygotic twin pairs. ICPM = number of inconsistent genotypes per 1Mb.*

A)

| Method | Total checks | Error rate |
|---|---|---|
| GATK | 1,277,130 | 1.19% |
| GraphTyper | 1,339,337 | 1.12% |

B)

| | GATK | GraphTyper |
|---|---|---|
| Total variants | 166,315 | 162,773 |
| SNPs only | 137,277 | 125,282 |
| Indels only | 29,038 | 37,491 |
| True positive estimate | 145,882 | 151,838 |
| SNPs only | 119,682 | 117,659 |
| Indels only | 26,200 | 34,179 |
| False discovery rate estimate | 12.28% | 6.72% |
| SNPs only | 12.82% | 6.08% |
| Indels only | 9.77% | 8.83% |

C)

| Method | Non-Ref Variants | Consistent | Error rate |
|---|---|---|---|
| GATK | 597,882 | 564,031 | 5.66% |
| GraphTyper | 603,589 | 578,763 | 4.11% |

*Table S3 Analysis of variant transmission of related samples in the 500 randomly selected 50kb test regions. A) Number of inheritance errors among the 28 parent-offspring trios. B) Estimates of number True Positives and False discovery rate in GATK and GraphTyper datasets in the trios. The estimates are determined from the allele transmission ratios from parent to offspring. C) Genotype consistency among the 14 pairs of monozygote twins.*

| Minimum number of carriers | Frequency threshold | GATK N imputed | GATK N markers | GATK Imputed ratio | GraphTyper N imputed | GraphTyper N markers | GraphTyper Imputed ratio |
|---|---|---|---|---|---|---|---|
| SNPs | | 54001 | 200471 | 26.9% | 58494 | 197508 | 29.6% |
| 2640 | 1.0% | 3157 | 3439 | 91.7% | 3380 | 3500 | 96.5% |
| 264 | 0.1% | 2480 | 3225 | 76.8% | 2623 | 2770 | 94.6% |
| 26 | 0.01% | 7436 | 10467 | 71.0% | 7859 | 9367 | 83.9% |
| 13 | 0.005% | 5491 | 7557 | 72.6% | 5857 | 7331 | 79.8% |
| 6 | 0.002% | 11326 | 17013 | 66.5% | 12230 | 16884 | 72.4% |
| 3 | 0.001% | 16503 | 33921 | 48.6% | 18095 | 33851 | 53.4% |
| 1 | 0.0002% | 7608 | 124849 | 6.0% | 8450 | 123805 | 6.8% |
| Indels | | 6124 | 21720 | 30.4% | 7876 | 20218 | 39.0% |
| 2640 | 1.0% | 842 | 935 | 90.0% | 1132 | 1254 | 90.2% |
| 264 | 0.1% | 602 | 854 | 70.4% | 790 | 917 | 86.1% |
| 26 | 0.01% | 1037 | 1861 | 55.7% | 1327 | 1723 | 77.0% |
| 13 | 0.005% | 570 | 1054 | 54.0% | 673 | 966 | 69.6% |
| 6 | 0.002% | 1038 | 1954 | 53.1% | 1172 | 1800 | 65.1% |
| 3 | 0.001% | 1352 | 3377 | 40.0% | 1521 | 3096 | 49.1% |
| 1 | 0.0002% | 683 | 11685 | 5.8% | 743 | 10462 | 7.1% |

*Table S4 Comparison of imputation of variants from the GATK and GraphTyper call sets on chr22 10-11Mb in the XBI dataset. A variant is considered imputed if phasing Leave-on-out-r2 (L1or2) is greater than 0.5 and imputation info is greater than 0.8.*

A)

| Method | WES AF>0.01% | WES AF>0.1% |
|---|---|---|
| GATK | 21,662 (1.81%) | 8,973 (2.54%) |
| GraphTyper | 5,310 (0.44%) | 1,903 (0.54%) |
| GraphTyperHQ | 16,774 (1.60%) | 7,693 (2.17%) |

B)

| | Present in WES 200k | | GATK | | GraphTyper | | GraphTyperHQ | |
|---|---|---|---|---|---|---|---|---|
| **Type** | **WES AF>0.01%** | **WES AF>0.1%** | **WES AF>0.01%** | **WES AF>0.1%** | **WES AF>0.01%** | **WES AF>0.1%** | **WES AF>0.01%** | **WES AF>0.1%** |
| A>C | 71,587 | 24,700 | 1,948 | 824 | 580 | 166 | 1,511 | 643 |
| A>G | 380,627 | 127,772 | 6,260 | 2,740 | 1,681 | 665 | 5,600 | 2,650 |
| A>T | 44,040 | 15,489 | 1,368 | 620 | 357 | 126 | 908 | 397 |
| C>G | 101,848 | 34,675 | 2,640 | 1,085 | 706 | 242 | 2,097 | 941 |
| C>T | 377,729 | 126,438 | 7,649 | 2,963 | 1,462 | 526 | 5,188 | 2,425 |
| G>T | 71,556 | 24,815 | 1,797 | 741 | 524 | 178 | 1,470 | 637 |
| Ti/Tv | 2.62 | 2.55 | 1.79 | 1.74 | 1.45 | 1.67 | 1.80 | 1.94 |

*Table S5 A) Number of variants in the WES 200k dataset that are missing from GATK, GraphTyper and GraphTyperHQ datasets, conditioned on the frequency in WES 200k. The fractions of missing variants are inside the parenthesis. B) Total number of SNP types present in WES 200K conditioned on frequency and how many of those are missing from our WGS datasets, stratified by variant type. Ti = number of transitions, Tv = number of transversions.*

| Mutation type | Mutations Autosomes | Mutations ChrX | Opportunities Autosomes | Opportunities ChrX | % Total | % Autosomes | % ChrX |
|---|---|---|---|---|---|---|---|
| C>A | 60,519,838 | 2,659,969 | 1,077,457,583 | 56,309,185 | 5.57% | 5.62% | 4.72% |
| C>G | 57,676,447 | 2,854,929 | 1,077,457,583 | 56,309,185 | 5.34% | 5.35% | 5.07% |
| C>T | 144,136,629 | 6,328,598 | 1,025,477,941 | 54,075,891 | 13.94% | 14.06% | 11.70% |
| CpG>TpG | 42,363,944 | 1,843,388 | 51,979,642 | 2,233,294 | 81.54% | 81.50% | 82.54% |
| T>A | 43,430,412 | 1,907,408 | 1,555,084,506 | 87,170,953 | 2.76% | 2.79% | 2.19% |
| T>C | 159,740,935 | 6,892,088 | 1,555,084,506 | 87,170,953 | 10.15% | 10.27% | 7.91% |
| T>G | 47,169,431 | 2,098,996 | 1,555,084,506 | 87,170,953 | 3.00% | 3.03% | 2.41% |

*Table S6 Mutation saturation, results presented for autosomes and chrX separately. Table shows the number of observed mutations in the GraphTyperHQ dataset and the number of possible mutation opportunities in regions of the genome amenable to short read sequence analysis.*

A)

| Method | Num variants | Missing call rate | Informative calls |
|---|---|---|---|
| GATK | 710,913,648 | 2.57% | 103,979,678,355,013 |
| GraphTyper | 655,928,639 | 0.14% | 98,332,325,114,654 |
| GraphTyperHQ | 643,747,446 | 0.07% | 96,570,956,991,770 |

B)

| Method | SNPs | Transitions (Ti) | Transversions (Tv) | Ti/Tv |
|---|---|---|---|---|
| GATK | 618,290,855 | 375,860,520 | 242,430,335 | 1.550 |
| GraphTyper | 593,953,779 | 369,120,364 | 224,833,415 | 1.642 |
| GraphTyperHQ | 585,040,410 | 364,859,729 | 220,180,681 | 1.657 |

C)

| Method | Common | % | Rare | % | Singleton | % |
|---|---|---|---|---|---|---|
| GATK | 31,501,254 | (4.4%) | 367,745,957 | (51.7%) | 311,666,437 | (43.9%) |
| SNP | 23,275,707 | (3.8%) | 317,087,938 | (51.3%) | 277,927,210 | (44.9%) |
| Non-SNP | 8,225,547 | (8.9%) | 50,658,019 | (54.7%) | 33,739,227 | (36.4%) |
| GraphTyper | 26,445,377 | (4.0%) | 335,241,409 | (51.1%) | 294,241,853 | (44.9%) |
| SNP | 20,261,132 | (3.4%) | 303,621,290 | (51.1%) | 270,071,357 | (45.5%) |
| Non-SNP | 6,184,245 | (10.0%) | 31,620,119 | (51.0%) | 24,170,496 | (39.0%) |
| GraphTyperHQ | 22,975,922 | (3.6%) | 327,718,095 | (50.9%) | 293,053,429 | (45.5%) |
| SNP | 18,124,082 | (3.1%) | 297,709,581 | (50.9%) | 269,206,747 | (46.0%) |
| Non-SNP | 4,851,840 | (8.3%) | 30,008,514 | (51.1%) | 23,846,682 | (40.6%) |

*Table S7 A) Number of variants in GATK, GraphTyper and GraphTyperHQ dataset. B) Variants split by transitions and transversions. C) Common = variants with frequency > 0.1%, rare = carried by more than one individual and frequency < 0.1%, singleton = carried by a single individual.*

| **Description** | **Beta** | **$R^2$** | ***P*-value** |
|---|---|---|---|
| Autosomal dominant genes from OMIM | -0.0407 | 0.00265 | 6.60E-12 |
| Recessive genes from OMIM | -0.0063 | 9.90E-05 | 0.1850 |
| Cell essential genes | 0.0259 | 0.00247 | 8.26E-10 |
| Present in Cell essential genes | -0.0907 | 0.02636 | 4.31E-105 |
| Hand curated list of Human lethal KO genes | -0.0204 | 0.00020 | 0.0627 |
| Hand curated list (more permissive) of Human lethal KO genes | -0.0221 | 0.00040 | 0.0074 |
| List of lethal KO genes in mice | -0.0425 | 0.00770 | 1.07E-31 |
| List of lethal het. KO genes in mice | -0.0275 | 0.00017 | 0.0824 |

*Table S8 Regression of average DR overlapping gene exons on annotations from Gene discovery informatics toolkit[40].*

| Data set | DR score | GERP RS score | CADD score | Eigen score | LINSIGHT score | CDTS |
|---|---|---|---|---|---|---|
| DR score | 1.000 | 0.005 | 0.038 | 0.029 | 0.011 | 0.158 |
| GERP RS score | 0.005 | 1.000 | 0.577 | 0.284 | 0.506 | 0.010 |
| CADD score | 0.039 | 0.577 | 1.000 | 0.554 | 0.547 | 0.075 |
| Eigen score | 0.029 | 0.284 | 0.554 | 1.000 | 0.690 | 0.065 |
| LINSIGHT score | 0.011 | 0.506 | 0.547 | 0.690 | 1.000 | 0.029 |
| CDTS | 0.158 | 0.010 | 0.075 | 0.064 | 0.029 | 1.000 |

*Table S9 Pearson correlation coefficient between DR score and measures of sequence constraint and functional impact, computed over all autosomal chromosomes. For each one of the 500bp overlapping windows in which the DR score ($dr$) is defined we compute the average value of the published scores ($ps$) in that window and then conduct linear regression analysis ($ps \sim dr$). The values shown in the table are the squared correlation coefficients of that regression. The correlation between the published datasets is computed from a set of 50bp non-overlapping windows using the average score within each window. A similar regression is conducted between each of the published datasets to obtain the squared correlation coefficient. Note, that the p-value for the linear regression fit is below computational threshold ($2.2 \times 10^{-308}$) for each pair of data sets in the table. CADD, Eigen and LINSIGHT all incorporate GERP into their annotation and are consequently not independent of each other or GERP. DR score and CDTS employ an analogous methodology, but scores are derived independently of each other and the other metrics.*

| Cohort | Chip N | WGS N | WGS % |
|---|---|---|---|
| XBI | 431,805 | 132,169 | 30.6 |
| XAF | 9,633 | 2,963 | 30.8 |
| XSA | 9,252 | 3,047 | 32.8 |
| OTH | 37,598 | 11,781 | 31.9 |

*Table S10 Number of individuals in the three cohorts described in this study.*

| Threshold % XBI | Threshold % XAF,XSA | | XBI Snp/Indel | XBI SV | XBI MSat | XAF Snp/Indel | XAF SV | XAF MSat | XSA Snp/Indel | XSA SV | XSA MSat |
|---|---|---|---|---|---|---|---|---|---|---|---|
| 1% | 5% | Phased | 11189434 | 15569 | 2491240 | 10782733 | 15214 | 2388595 | 7941383 | 11773 | 1812461 |
| | | Imputed | 11184312 | 15518 | 2488009 | 10728154 | 14606 | 2354675 | 7865444 | 11211 | 1743407 |
| | | n | 11297050 | 18044 | 2600902 | 10864088 | 17276 | 2453893 | 7993858 | 13415 | 1859421 |
| 0.1% | 1% | Phased | 6590616 | 7234 | 1068743 | 8365507 | 9301 | 1166814 | 3739563 | 4230 | 816116 |
| | | Imputed | 6586277 | 7223 | 1066743 | 8315664 | 9140 | 1129348 | 3633830 | 4072 | 699673 |
| | | n | 6819668 | 9185 | 1329131 | 8555777 | 11074 | 1310478 | 3852601 | 5235 | 896908 |
| 0.01% | 0.5% | Phased | 23598990 | 24317 | 1581904 | 3950291 | 4139 | 391700 | 2122454 | 2168 | 369854 |
| | | Imputed | 23453107 | 24037 | 1558812 | 3914244 | 4077 | 369608 | 2008801 | 2062 | 271659 |
| | | n | 24556101 | 31246 | 2330992 | 4114602 | 5187 | 504462 | 2280485 | 2808 | 442328 |
| 0.005% | 0.2% | Phased | 19864378 | 19181 | 482916 | 4386799 | 4263 | 354516 | 2642260 | 2558 | 380537 |
| | | Imputed | 19440299 | 18735 | 457453 | 4316711 | 4136 | 319739 | 2409667 | 2297 | 235403 |
| | | n | 21059670 | 25103 | 850280 | 4722651 | 5635 | 515106 | 2982169 | 3624 | 488799 |
| 0.002% | 0.1% | Phased | 43902487 | 41664 | 600207 | 6892163 | 6336 | 448483 | 5032021 | 4717 | 539656 |
| | | Imputed | 41679009 | 39448 | 542137 | 6627483 | 6041 | 366561 | 4292840 | 3964 | 260599 |
| | | n | 50063971 | 55664 | 1214690 | 8424367 | 9507 | 772036 | 6418472 | 7497 | 786098 |
| 0.001% | 0.04% | Phased | 52975884 | 49438 | 437379 | 6495546 | 5681 | 337279 | 5944556 | 5125 | 428185 |
| | | Imputed | 47238234 | 44171 | 363952 | 5861702 | 5106 | 240464 | 4635202 | 3933 | 162809 |
| | | n | 72522701 | 74342 | 1092057 | 10462313 | 10807 | 713472 | 9539163 | 10093 | 745160 |
| 0.0002% | 0.008% | Phased | 40518700 | 36567 | 292304 | 16233640 | 12769 | 569083 | 12625599 | 8801 | 715628 |
| | | Imputed | 31988313 | 29453 | 189966 | 12109642 | 10130 | 321072 | 6563488 | 4830 | 190230 |
| | | n | 263633284 | 261011 | 1935535 | 59096600 | 52531 | 1654282 | 52146463 | 47256 | 1671469 |

*Table S11 Imputation and phasing accuracy as a function of frequency within each cohort. Phased refers to number of variants with Leave-one-out-r2 value > 0.5 and imputed refers to phased variants that also have imputation info > 0.8. Numbers are for variants at frequency above the given threshold and not included in frequency thresholds in earlier lines, e.g., in the XBI population 72,522,701 variants have frequency between 0.001 and 0.002%, of which 52,975,884 could be phased and 47,238,234 could be imputed.*

| AF Threshold | Panel | XBI n | XBI present % | XAF n | XAF present % | XSA n | XSA present % |
|---|---|---|---|---|---|---|---|
| $\geq 10^{-2}$ | Bycroft | 9,675,179 | 57.1% | 9,049,185 | 54.4% | 8,782,729 | 55.4% |
| | 150k WGS | 16,838,810 | 99.3% | 16,500,186 | 99.3% | 15,728,295 | 99.1% |
| | Both | 9,555,642 | 56.3% | 8,924,920 | 53.7% | 8,645,958 | 54.5% |
| | Either | 16,958,347 | 100% | 16,624,451 | 100% | 15,865,066 | 100% |
| $\geq 10^{-3}$ | Bycroft | 5,150,551 | 40.8% | 4,321,491 | 37.0% | 1,509,037 | 23.8% |
| $< 10^{-2}$ | 150k WGS | 12,497,109 | 99.1% | 11,609,254 | 99.3% | 6,276,519 | 98.8% |
| | Both | 5,031,517 | 39.9% | 4,236,985 | 36.2% | 1,432,690 | 22.6% |
| | Either | 12,616,143 | 100% | 11,693,760 | 100% | 6,352,866 | 100% |
| $\geq 10^{-4}$ | Bycroft | 4,635,660 | 12.7% | 7,894,440 | 34.0% | 1,637,838 | 17.8% |
| $< 10^{-3}$ | 150k WGS | 36,247,790 | 99.1% | 22,801,909 | 98.2% | 8,903,892 | 96.5% |
| | Both | 4,299,464 | 11.8% | 7,474,332 | 32.2% | 1,315,077 | 14.3% |
| | Either | 36,583,986 | 100% | 23,222,017 | 100% | 9,226,653 | 100% |
| $< 10^{-4}$ | Bycroft | 1,786,117 | 0.9% | 4,951,605 | 8.8% | 2,001,548 | 5.3% |
| | 150k WGS | 196,375,197 | 99.6% | 54,623,218 | 97.1% | 37,019,802 | 97.4% |
| | Both | 942,249 | 0.5% | 3,315,555 | 5.9% | 1,024,799 | 2.7% |
| | Either | 197,219,065 | 100% | 56,259,268 | 100% | 37,996,551 | 100% |

*Table S12 Number of markers that impute (Imp Info > .8) in 500k set of UKB using the imputation panel presented here (150k WGS) and an imputation by Bycroft et al.[5]. Both represents number of markers imputed by both panels, either the number of markers in either panel.*

| # repeats | Frequency | Effect | *P*-value |
|---|---|---|---|
| 4.7 | 5.98E-05 | 0.02 | 8.84E-01 |
| 5.7 | 3.64E-01 | 0.41 | 3.95E-07 |
| 6 | 1.45E-06 | 0.02 | 9.84E-01 |
| 6.7 | 9.99E-04 | 4.96 | 2.16E-01 |
| 7.7 | 4.16E-04 | 0.02 | 6.94E-01 |
| 8.7 | 7.46E-03 | 0.69 | 6.95E-01 |
| 9.7 | 1.21E-03 | 4.27 | 2.54E-01 |
| 10.7 | 9.27E-03 | 0.54 | 5.10E-01 |
| 11.7 | 1.20E-01 | 1.45 | 7.33E-02 |
| 12 | 1.95E-06 | 0.02 | 9.81E-01 |
| 12.7 | 1.24E-01 | 0.85 | 4.98E-01 |
| 13 | 1.17E-06 | 0.02 | 9.87E-01 |
| 13.7 | 1.78E-01 | 0.77 | 2.08E-01 |
| 14.7 | 6.30E-02 | 0.82 | 5.26E-01 |
| 15.7 | 8.18E-03 | 0.73 | 7.43E-01 |
| 16.7 | 9.16E-03 | 0.01 | 7.66E-02 |
| 17.7 | 4.88E-03 | 1.06 | 9.54E-01 |
| 18.7 | 2.15E-03 | 0.02 | 3.72E-01 |
| 19.7 | 4.39E-03 | 1.44 | 7.34E-01 |
| 20.7 | 1.81E-02 | 2.02 | 1.32E-01 |
| 21.7 | 2.69E-02 | 1.12 | 8.27E-01 |
| 22.7 | 1.41E-02 | 2.4 | 1.38E-01 |
| 23.7 | 8.77E-03 | 2.18 | 3.15E-01 |
| 24.7 | 6.90E-03 | 2.44 | 2.59E-01 |
| 25.7 | 7.05E-03 | 4.06 | 4.74E-02 |
| 26.7 | 5.17E-03 | 6.62 | 1.40E-02 |
| 27.7 | 4.55E-03 | 9.82 | 1.50E-03 |
| 28.7 | 3.38E-03 | 17.93 | 1.24E-04 |
| 29.7 | 2.33E-03 | 19.75 | 4.08E-04 |
| 30.7 | 1.55E-03 | 30.02 | 6.02E-05 |
| 31.7 | 1.03E-03 | 48.35 | 4.33E-09 |
| 32.7 | 6.98E-04 | 42.04 | 1.30E-04 |
| 33.7 | 4.04E-04 | 74.03 | 8.07E-06 |
| 34.7 | 3.05E-04 | 68.27 | 5.01E-05 |
| 35.7 | 1.48E-04 | 141.58 | 1.29E-10 |
| 36.7 | 1.50E-04 | 45.23 | 3.35E-02 |
| 37.7 | 9.60E-05 | 51.68 | 2.49E-01 |
| 38.7 | 1.04E-04 | 92.19 | 3.64E-03 |
| >=39.7 | 4.32E-05 | 161.74 | 1.09E-07 |

*Table S13 Association of number of repeat copies of microsatellite in 3' UTR in DMPK with myotonic dystrophy. Individuals carrying 39.7 or more copies of the repeat are grouped together by popSTR[64].*

A)

| Gene | Number of Allelic variants in OMIM | OMIM Phenotype with allelic variants * (Mode of inheritance) ** |
|---|---|---|
| *ALB* | 61 | Analbuminemia (AR) |
| *CACNA1A* | 37 | Episodic ataxia, type 2 (AD) ; Migraine, familial hemiplegic, 1 (AD); Epileptic encephalopathy, early infantile, 42 (AD); Spinocerebellar ataxia 6 (AD) |
| *HBB* | 540 | Delta-beta thalassemia(AD); Erythrocytosis 6 (AD) ; Heinz body anemia (AD); Hereditary persistence of fetal hemoglobin (AD); Methemoglobinemia, beta type(AD); Sickle cell anemia (AR); Thalassemia-beta, dominant inclusion-body (AD) |
| *PCSK9* | 8 | Hypercholesterolemia, familial, 3 (AD) |
| *PIEZO1* | 16 | Dehydrated hereditary stomatocytosis(AD); Lymphedema, hereditary, III (AR) |
| *GHRH* | 0 | None |
| *DMPK* | 1 | Myotonic dystrophy 1 (AD) |
| *GCSH* | 1 | None |
| *TAC3* | 2 | Hypogonadotropic hypogonadism 10 with or without anosmia (AR) |
| *NMRK2* | 0 | None |

B)

| Gene | N Drug *** | Indications *** | Link |
|---|---|---|---|
| *ALB* | None | | |
| *CACNA1A* | 5 | 7 | https://platform.opentargets.org/target/ENSG00000141837 |
| *HBB* | 3 | 11 | https://platform.opentargets.org/target/ENSG00000244734 |
| *PCSK9* | 6 | 28 | https://platform.opentargets.org/target/ENSG00000169174 |
| *PIEZO1* | None | | |
| *GHRH* | None | | |
| *DMPK* | None | | |
| *GCSH* | None | | |
| *TAC3* | None | | |
| *NMRK2* | None | | |

*Table S14 Information on genes presented. A) Phenotypes and allelic variants in OMIM for selected genes. B) Known drug data and in open targets for selected targets. *Excluding the ones with provisional phenotype gene relationship "?"; multifactorial diseases"{ }" and non diseases"[ ]" ** Mode of inheritance : AD Autosomal dominant; AR Autosomal recessive. ***Known drug data according to Open Targets[77].*

| Phenotype | Data showcase field | Extra information |
|---|---|---|
| Age at menopause | 3581 | Adjusted for year of birth and 20 principal components, then inverse-normal transformed |
| Age of menarche | 2714 | Adjusted for year of birth and 20 principal components, then inverse-normal transformed |
| Albumin | 30600 | Adjusted for age, age² and 20 principal components, then combined and inverse-normal transformed |
| Calcium | 30680 | Adjusted for age, age² and 20 principal components, then combined and inverse-normal transformed |
| Glycine | 23462 | Metabolomics |
| Height | 50 | Adjusted for year of birth, sex and 20 principal components for males and females separately, then combined and inverse-normal transformed |
| Hemoglobin concentration, Asian ancestry | 30060 | Adjusted for age, age² and 45 principal components for males and females separately, then combined and inverse-normal transformed |
| IGF-1 serum levels | 30770 | Adjusted for age, age² and 20 principal components |
| Mean corpuscular volume | 30040 | Adjusted for age, age² and 20 principal components for males and females separately, then combined and inverse-normal transformed |
| Non-HDL cholesterol, European ancestry | Field 30690 minus field 30670 (HDL) | Adjusted for age, age² and 20 principal components; lipid-lowering drug users had their measurements divided by 0.8, then combined and inverse-normal transformed |
| Non-HDL cholesterol, African ancestry | Field 30690 minus field 30670 (HDL) | Adjusted for age, age² and 20 principal components; lipid-lowering drug users had their measurements divided by 0.8, then combined and inverse-normal transformed |
| Total cholesterol | 30690 | Adjusted for age, age² and 20 principal components; lipid-lowering drug users had their measurements divided by 0.8, then combined and inverse-normal transformed |
| Uric acid | 30880 | Adjusted for age, age² and 20 principal components, then combined and inverse-normal transformed |
| Gout | ICD-19 code M10* on fields 41270, 41271 and 42040 | Adjusted for year of birth, sex and 20 principal components |
| Hereditary ataxia | ICD-10 code G11 on fields 41270, 41271 and 42040 | Adjusted for year of birth, sex and 20 principal components |
| Myotonic dystrophy | ICD-10 code G71.1 on fields 41270, 41271 and 42040 | Adjusted for year of birth, sex and 20 principal components |

*Table S15 Phenotypes used in this study, their field in the UKB data showcase and adjustments performed prior to association analysis*

| Parameter | Information Requested | Definition |
|---|---|---|
| prc_auto_ge_15x | Coverage | PCT_15X from .wgsmetrics_autosome in QCPreview |
| coverage | autosomal mean coverage | MEAN_COVERAGE * (1.0 - PCT_EXC_DUPE - PCT_EXC_OVERLAP - PCT_EXC_ADAPTER) / (1.0 - PCT_EXC_TOTAL) from .wgsmetrics_autosome in QCPreview |
| genetic_sex | Sex | if NX<=0.3 then "Female" else if NX>=0.7 then "Male" else "Undetermined" from .sexcheck output file in QCStats |
| yield | Yield | GENOME_TERRITORY * MEAN_COVERAGE * (1.0 - PCT_EXC_DUPE - PCT_EXC_OVERLAP - PCT_EXC_ADAPTER) / (1.0 - PCT_EXC_TOTAL) from .wgsmetrics output file in QCPreview |
| read_haps_error_percentage | Read_haps | 100*DOUBLE_ERROR_FRACTION from .contamination output file in QCStats |
| freemix_percentage | Freemix/Verify Bam ID | 100 * FREEMIX from .verifyBamId.selfSM output file in QCStats |
| prc_proper_pairs | Proportion of mapped read pairs | 100 * (reads_properly_paired/reads_mapped) from .stats output file in QCPreview |
| discordance_prc | NRD Genotyping | 100 * (1.0 - NON_REF_GENOTYPE_CONCORDANCE) from .genotype_concordance_summary_metrics in Concords or -1 if chip genotypes are not available |

*Table S16 QA/QC metrics derived from the files delivered to the UKB. The result is written to a file, qaqc_metric.*

| Column | Min | Max | Flag | Explanation |
|---|---|---|---|---|
| SAMPLE_ID | | | | Read group ID |
| LANE | | | | Lane ID (=Read group ID) |
| FAILURE_FLAGS | | | | Failure flag |
| JOINT_CALLING_FLAGS | | | | Joint calling failure flag |
| STRICT_FLAGS | | | | Strict failure flag |
| TOTAL_BPS | 3e8 | 1e14 | C | Total basepairs |
| TOTAL_READ_PAIRS | | | | Total read pairs |
| READ_LENGTH | | | | Read length |
| MEAN_BASE_QUAL_PER_READ | 30 | 100 | Q | Mean of base calling quality |
| STD_BASE_QUAL_PER_READ | -1 | 10 | Q | Std dev of mean base calling quality |
| MEAN_N_COUNT_PER_READ | -1 | 10 | N | Mean Percentage N |
| STD_N_COUNT_PER_READ | -1 | 30 | N | Std dev of Percentage N |
| MEAN_GC_CONTENT_PER_READ | 39 | 45 | G | Mean percentage of GC bases |
| STD_GC_CONTENT_PER_READ | -1 | 15 | G | Std dev of Percentage GC |
| MEAN_BASE_QUAL_PER_POSITION | 30 | 100 | Q | Mean of mean base calling quality |
| STD_BASE_QUAL_PER_POSITION | -1 | 6 | Q | Std dev of mean base calling quality |
| MEAN_N_PER_POSITION | -1 | 10 | N | Mean Percentage N |
| STD_N_PER_POSITION | -1 | 10 | N | Std dev of Percentage N |
| MEAN_A_PER_POSITION | 25 | 35 | B | Mean Percentage A |
| STD_A_PER_POSITION | -1 | 10 | B | Std dev of Percentage A |
| MEAN_C_PER_POSITION | 15.5 | 25 | B | Mean Percentage C |
| STD_C_PER_POSITION | -1 | 10 | B | Std dev of Percentage C |
| MEAN_G_PER_POSITION | 17 | 24 | B | Mean Percentage G |
| STD_G_PER_POSITION | -1 | 10 | B | Std dev of Percentage G |
| MEAN_T_PER_POSITION | 25 | 33 | B | Mean Percentage T |
| STD_T_PER_POSITION | -1 | 10 | B | Std dev of Percentage T |
| 32_MER_ERROR_RATE | | | | Estimated 32-mer error rate |
| ADAPTER_8_MERS | -1 | 5 | A | Percentage of Universal adapter 8-mers |
| MARKED_DUPLICATE | -1 | 60 | D | Percentage marked as duplicate |
| UNMAPPED | -1 | 20 | U | Percentage unmapped reads |
| BOTH_UNMAPPED | -1 | 30 | U | Percentage both reads in pair unmapped |
| FIRST_UNMAPPED | -1 | 30 | U | Percentage only first unmapped in pair |
| SECOND_UNMAPPED | -1 | 30 | U | Percentage only second unmapped in pair |
| PROPER_PAIRS | | | | Percentage proper pairs |
| PROPER_PAIRS_AUTOSOME | 95 | 1000 | P | Percentage proper pairs autosome |
| FF_RR_PAIRS | -1 | 0.1 | o | Percentage FF/RR oriented pairs |
| MEAN_COVERAGE | 0.1 | 100000 | C | Mean coverage |
| STD_COVERAGE | -1 | 100000 | C | Std dev of coverage |
| MEAN_INSERT_SIZE | -1 | 10000 | I | Mean insert size |
| STD_INSERT_SIZE | | | | Std dev of insert size |
| ADAPTER_INSERT_SIZE | -1 | 20 | A | Percent insert size < read length |
| MAPPING_QUAL_60 | | | | Percentage reads with mapping quality <60 |
| MAPPING_QUAL_40 | | | | Percentage reads with mapping quality <40 |
| MAPPING_QUAL_20 | | | | Percentage reads with mapping quality <20 |
| MEAN_MISMATCHES | -1 | 5 | m | Mean mismatches per read pair |
| MEAN_DELETIONS | | | | Mean deletions per read pair |
| MEAN_INSERTIONS | | | | Mean insertions per read pair |
| NZ_DELETIONS | -1 | 0.1 | d | Fraction or reads that have a deletion |
| NZ_INSERTIONS | -1 | 0.1 | I | Fraction of reads that have an insertion |
| CLIPPED_5_PRIME | -1 | 6 | c | Percentage of reads clipped at 5'-end |
| CLIPPED_3_PRIME | -1 | 30 | c | Percentage of reads clipped at 3'-end |
| C>A | 0.3 | 0.7 | O | C>A triplet conversion rate |
| G>A | 0.4 | 0.6 | O | G>A triplet conversion rate |
| T>A | 0.3 | 0.7 | O | T>A triplet conversion rate |
| A>C | 0.3 | 0.7 | O | A>C triplet conversion rate |
| G>C | 0.3 | 0.7 | O | G>C triplet conversion rate |
| T>C | 0.3 | 0.7 | O | T>C triplet conversion rate |

*Table S17 Metrics collected for each lane by bamqc_summary. If any flag is raised, the lane is excluded from the merge process. The values, per read group, are collected in the file .bamqc_summary.*

A)

| Method | #Variants | Common (>0.1%) | Rare (<0.1%) | Singleton |
|---|---|---|---|---|
| GATK | 6,221,575 | 284,303 | 3,259,421 | 2,677,851 |
| GraphTyper | 5,569,026 | 224,715 | 2,855,132 | 2,489,179 |

B)

| Method | #Variants | SNPs | Non-SNPs |
|---|---|---|---|
| GATK | 6,221,575 | 5,400,679 | 820,896 |
| GraphTyper | 5,569,026 | 5,040,466 | 528,560 |

C)

| Method | Missing genotypes | #Informative calls |
|---|---|---|
| GATK | 3.26% | 903,536,315,740 |
| GraphTyper | 0.11% | 835,097,232,768 |

D)

| Method | Transitions (Ti) | Transversion (Tv) | Ti/Tv |
|---|---|---|---|
| GATK | 3,246,174 | 2,154,505 | 1.507 |
| GraphTyper | 3,130,524 | 1,909,942 | 1.639 |

*Table S18 Results for 500 random test regions. A) Number of variants called by GATK and GraphTyper conditioned on frequency class. B) Number of variants conditioned on variant type. C) Fraction of missing variant calls. D) Number of transitions and transversions.*

A)

| Method | Total common | Failed | % |
|---|---|---|---|
| GATK | 284,303 | 21,234 | 7.47% |
| GraphTyper | 224,715 | 2,277 | 1.01% |

B)

| | Failed count | |
|---|---|---|
| **Test** | **GATK** | **GraphTyper** |
| Sanger Vanguard vs. Sanger Main | 13,440 | 999 |
| Sanger Vanguard vs. deCODE | 16,751 | 1,825 |
| Sanger Main vs. deCODE | 13,510 | 1,141 |

*Table S19 Number of common variants (frequency > .1%) that showed significant association with sequencing center in the 500 random regions test set., A) Total number of variants that failed in any test. B) Number of failed variants stratified by sequencing protocol. Variant is considered “Failed” if p-value < 1e-6, Fisher’s exact test.*

A)

| Method | Common | SaM vs. deC | SaV vs. deC | SaV vs. SaM |
|---|---|---|---|---|
| GATK | 31,501,254 | 1,202,575 | 1,164,682 | 810,105 |
| GraphTyper | 26,445,377 | 166,371 | 175,144 | 66,838 |
| GraphTyperHQ | 22,975,922 | 28,432 | 36,283 | 8,096 |

B)

| Method | Common | Any test $p < 10^{-6}$ | Any test $p < 10^{-10}$ |
|---|---|---|---|
| GATK | 31,501,254 | 1,792,003 (5.69%) | 1,197,839 (3.80%) |
| GraphTyper | 26,445,377 | 257,860 (0.97%) | 136,521 (0.52%) |
| GraphTyperHQ | 22,975,922 | 46,556 (0.20%) | 22,307 (0.10%) |

*Table S20 Number of common variants (frequency > 0.1%) that show significant association to sequencing center, indicating batch effects, using a Fisher's exact test, for common (> 0.1% frequency) variants. A) Number of failed variants stratified by test using $p < 10^{-6}$. deC = samples sequenced at deCODE genetics. SaV = samples sequenced using the Sanger Vanguard processing pipeline. SaM = samples sequenced using the Sanger main phase pipeline. B) Total number of variants that failed in any test, using both $p < 10^{-6}$ and $p < 10^{-10}$.*

| Dataset | Shared with both other | Specific to | Absent from | Absent from and same carrier in both other datasets | Fraction of missing variants with same carrier in both datasets |
|---|---|---|---|---|---|
| GATK | 6,608,669 | 230,808 | 15,567 | 12,700 | 81.58% |
| GraphTyperHQ | 6,608,669 | 54,909 | 87,773 | 56,052 | 63.86% |
| WES200k | 6,608,669 | 28,039 | 498,181 | 476,195 | 95.59% |

*Table S21 Three-way comparison between the GraphTyperHQ, GATK and WES200k[76] call analyzed inside WES capture regions within the set of 109,618 individuals present in both the WES200k call set an our set of 150,119 individuals.*

| XBI | *P*-value | MaF > 0.01 | MaF 0.01 - 0.001 | MaF < 0.001 |
|---|---|---|---|---|
| Unfiltered | > 0.05 | 17.0-17.5M (89-91%) | 9.27-9.33M (93-94%) | 208-215M (94-97%) |
| | 0.005-0.05 | 1.05-1.13M (5.5-5.9%) | 481-497K (4.9-5.0%) | 5.60-12.4M (2.5-5.6%) |
| | 5e-4 - 0.005 | 238-258K (1.2-1.3%) | 64.1-77.8K (0.65-0.78%) | 443-848K (0.2-0.38%) |
| | 5e-8 - 5e-4 | 214-324K (1.1-1.7%) | 28.9-49.9K (0.29-0.5%) | 36.8-65.6K (0.017-0.03%) |
| | < 5e-8 | 127-540K (0.66-2.8%) | 7.74-49.2K (0.078-0.5%) | 364-3697 (0.00016-0.0017%) |
| Filtered | > 0.05 | 16.0-16.1M (94-94%) | 8.94-8.96M (94-95%) | 207-214M (94-97%) |
| | 0.005-0.05 | 808-839K (4.7-4.9%) | 435-445K (4.6-4.7%) | 5.57-12.3M (2.5-5.6%) |
| | 5e-4 - 0.005 | 103-122K (0.6-0.72%) | 46.9-55.5K (0.5-0.59%) | 439-840K (0.2-0.38%) |
| | 5e-8 - 5e-4 | 36.1-78.4K (0.21-0.46%) | 10.1-16.7K (0.11-0.18%) | 36.1-60.7K (0.016-0.028%) |
| | < 5e-8 | 11.2-68.9K (0.066-0.4%) | 2.37-11.5K (0.025-0.12%) | 115-463 (5.2e-05-0.00021%) |
| **XAF** | ***P*-value** | **MaF > 0.01** | **MaF 0.01 - 0.001** | **MaF < 0.001** |
| Unfiltered | > 0.05 | 29.7-29.9M (94-95%) | 22.5-22.9M (93-95%) | 80.8-84.6M (95-99%) |
| | 0.005-0.05 | 1.43-1.48M (4.5-4.7%) | 1.04-1.44M (4.3-6.0%) | 0.717-4.41M (0.84-5.2%) |
| | 5e-4 - 0.005 | 152-189K (0.48-0.6%) | 79.6-143K (0.33-0.59%) | 10.6-118K (0.012-0.14%) |
| | 5e-8 - 5e-4 | 20.4-73.6K (0.065-0.23%) | 6.87-18.2K (0.029-0.076%) | 1-5392 (1.2e-06-0.0063%) |
| | < 5e-8 | 732-29023 (0.0023-0.092%) | 62-335 (0.00026-0.0014%) | 0-1 (0.0-1.2e-06%) |
| Filtered | > 0.05 | 27.4-27.4M (95-95%) | 21.8-22.2M (93-95%) | 80.1-83.9M (95-99%) |
| | 0.005-0.05 | 1.26-1.30M (4.4-4.5%) | 0.994-1.39M (4.3-6.0%) | 0.709-4.38M (0.84-5.2%) |
| | 5e-4 - 0.005 | 127-133K (0.44-0.46%) | 75.7-135K (0.33-0.58%) | 10.5-117K (0.012-0.14%) |
| | 5e-8 - 5e-4 | 13.4-23.8K (0.046-0.083%) | 6.36-15.3K (0.027-0.066%) | 1-5294 (1.2e-06-0.0063%) |
| | < 5e-8 | 28-5752 (9.7e-05-0.02%) | 0-166 (0.0-0.00071%) | 0-0 (0.0-0.0%) |
| **XSA** | ***P*-value** | **MaF > 0.01** | **MaF 0.01 - 0.001** | **MaF < 0.001** |
| Unfiltered | > 0.05 | 18.9-19.1M (94-95%) | 14.1-14.5M (94-96%) | 73.9-76.5M (95-99%) |
| | 0.005-0.05 | 919-989K (4.6-4.9%) | 521-817K (3.5-5.4%) | 1.00-3.58M (1.3-4.6%) |
| | 5e-4 - 0.005 | 99.7-142K (0.5-0.71%) | 32.1-92.3K (0.21-0.61%) | 17.3-83.7K (0.022-0.11%) |
| | 5e-8 - 5e-4 | 13.2-67.8K (0.066-0.34%) | 2.97-12.7K (0.02-0.085%) | 358-3980 (0.00046-0.0051%) |
| | < 5e-8 | 665-30416 (0.0033-0.15%) | 92-278 (0.00061-0.0018%) | 0-2 (0.0-2.6e-06%) |
| Filtered | > 0.05 | 17.0-17.0M (95-95%) | 13.6-13.9M (94-96%) | 73.3-75.9M (95-99%) |
| | 0.005-0.05 | 796-809K (4.4-4.5%) | 494-778K (3.4-5.4%) | 0.994-3.56M (1.3-4.6%) |
| | 5e-4 - 0.005 | 82.4-87.9K (0.46-0.49%) | 30.6-85.5K (0.21-0.59%) | 17.1-82.8K (0.022-0.11%) |
| | 5e-8 - 5e-4 | 9.29-15.8K (0.052-0.088%) | 2.75-10.1K (0.019-0.07%) | 331-3865 (0.00043-0.005%) |
| | < 5e-8 | 16-4327 (8.9e-05-0.024%) | 1-142 (6.9e-06-0.00098%) | 0-0 (0.0-0.0%) |

*Table S22 Batch effects for sequencing center in the raw genotype calls. Six phenotypes for batch effects are tested. Results are conditioned on marker minor allele frequency (MAF). Table shows the minimum and maximum number and fraction of markers, across the six phenotypes) with p-value in each p-value range. E.g., when considering the unfiltered dataset and the XSA cohort, MAF > 0.01, between 919 and 989k markers have p-value between 0.005 and 0.05, corresponding to 4.6-4.9% of markers with MAF > 0.01.*

| XBI | *P*-value | MaF > 0.01 | MaF 0.01 - 0.001 | MaF < 0.001 |
|---|---|---|---|---|
| Unfiltered | > 0.05 | 16.8-17.1M (92-94%) | 9.47-9.55M (94-95%) | 254-266M (93-98%) |
| | 0.005-0.05 | 887-910K (4.9-5.0%) | 472-495K (4.7-4.9%) | 6.30-17.3M (2.3-6.3%) |
| | 5e-4 - 0.005 | 113-155K (0.62-0.85%) | 54.4-71.4K (0.54-0.71%) | 466-942K (0.17-0.35%) |
| | 5e-8 - 5e-4 | 40.2-137K (0.22-0.75%) | 10.4-40.5K (0.1-0.4%) | 38.5-74.5K (0.014-0.027%) |
| | < 5e-8 | 15.9-180K (0.087-0.99%) | 925-27218 (0.0092-0.27%) | 85-1389 (3.1e-05-0.00051%) |
| Filtered | > 0.05 | 15.4-15.4M (95-95%) | 8.78-8.79M (95-95%) | 216-225M (93-97%) |
| | 0.005-0.05 | 733-738K (4.5-4.5%) | 418-421K (4.5-4.6%) | 5.63-14.7M (2.4-6.4%) |
| | 5e-4 - 0.005 | 72.5-82.8K (0.45-0.51%) | 42.1-44.9K (0.46-0.49%) | 425-848K (0.18-0.37%) |
| | 5e-8 - 5e-4 | 8.07-24.2K (0.05-0.15%) | 4.74-6.31K (0.051-0.068%) | 35.4-64.2K (0.015-0.028%) |
| | < 5e-8 | 117-11166 (0.00072-0.069%) | 0-592 (0.0-0.0064%) | 0-7 (0.0-3e-06%) |
| **XAF** | ***P*-value** | **MaF > 0.01** | **MaF 0.01 - 0.001** | **MaF < 0.001** |
| Unfiltered | > 0.05 | 27.9-28.0M (95-95%) | 19.6-19.8M (94-95%) | 77.1-79.6M (96-99%) |
| | 0.005-0.05 | 1.30-1.34M (4.4-4.5%) | 0.950-1.13M (4.6-5.4%) | 0.670-3.04M (0.84-3.8%) |
| | 5e-4 - 0.005 | 132-148K (0.45-0.5%) | 72.6-124K (0.35-0.59%) | 16.1-107K (0.02-0.13%) |
| | 5e-8 - 5e-4 | 13.8-32.8K (0.047-0.11%) | 5.97-14.3K (0.029-0.068%) | 300-5385 (0.00037-0.0067%) |
| | < 5e-8 | 92-5922 (0.00031-0.02%) | 36-136 (0.00017-0.00065%) | 0-3 (0.0-3.7e-06%) |
| Filtered | > 0.05 | 25.3-25.3M (95-95%) | 17.6-17.8M (94-95%) | 60.3-62.2M (96-99%) |
| | 0.005-0.05 | 1.16-1.19M (4.4-4.5%) | 856-996K (4.6-5.3%) | 0.556-2.42M (0.89-3.9%) |
| | 5e-4 - 0.005 | 110-118K (0.41-0.44%) | 64.7-106K (0.35-0.57%) | 13.9-85.6K (0.022-0.14%) |
| | 5e-8 - 5e-4 | 11.6-13.5K (0.043-0.051%) | 5.14-10.9K (0.027-0.058%) | 258-4427 (0.00041-0.0071%) |
| | < 5e-8 | 1-104 (3.8e-06-0.00039%) | 0-1 (0.0-5.3e-06%) | 0-0 (0.0-0.0%) |
| **XSA** | ***P*-value** | **MaF > 0.01** | **MaF 0.01 - 0.001** | **MaF < 0.001** |
| Unfiltered | > 0.05 | 17.7-17.8M (95-95%) | 13.8-14.1M (94-96%) | 67.2-68.7M (97-99%) |
| | 0.005-0.05 | 836-876K (4.5-4.7%) | 506-780K (3.5-5.3%) | 0.674-2.14M (0.97-3.1%) |
| | 5e-4 - 0.005 | 84.3-103K (0.45-0.55%) | 33.4-84.6K (0.23-0.58%) | 14.8-71.5K (0.021-0.1%) |
| | 5e-8 - 5e-4 | 9.70-26.1K (0.052-0.14%) | 2.83-10.0K (0.019-0.068%) | 531-3555 (0.00077-0.0051%) |
| | < 5e-8 | 83-6253 (0.00044-0.033%) | 26-94 (0.00018-0.00064%) | 0-11 (0.0-1.6e-05%) |
| Filtered | > 0.05 | 15.6-15.6M (95-95%) | 10.8-11.0M (94-96%) | 40.0-40.9M (97-99%) |
| | 0.005-0.05 | 718-736K (4.4-4.5%) | 412-603K (3.6-5.3%) | 0.478-1.38M (1.2-3.3%) |
| | 5e-4 - 0.005 | 71.8-76.5K (0.44-0.47%) | 25.4-65.0K (0.22-0.57%) | 11.0-49.4K (0.027-0.12%) |
| | 5e-8 - 5e-4 | 8.02-9.39K (0.049-0.057%) | 2.14-6.70K (0.019-0.058%) | 394-2561 (0.00095-0.0062%) |
| | < 5e-8 | 0-47 (0.0-0.00029%) | 0-0 (0.0-0.0%) | 0-0 (0.0-0.0%) |

*Table S23 Batch effects for sequencing center in the imputed genotype calls. Six phenotypes for batch effects are tested. Results are conditioned on marker minor allele frequency (MAF). Table shows the minimum and maximum number and fraction of markers, across the six phenotypes) with p-value in each p-value range. E.g., when considering the unfiltered dataset and the XSA cohort, MAF > 0.01, between 836 and 876k markers have p-value between 0.005 and 0.05, corresponding to 4.5-4.7% of markers with MAF > 0.01.*

| Phenotype | LD score intercept | Mean $\chi^2$ unadj | $\lambda$ unadj | $\lambda$ unadj maf<0.01 | Attenuation ratio | Method | Marker |
|---|---|---|---|---|---|---|---|
| Age at menopause | 1.051 | 1.463 | 1.048 | 1.005 | 0.110 | BOLT-LMM | chr19:3939254 |
| Age of menarche | 1.095 | 2.081 | 1.048 | 1.028 | 0.088 | BOLT-LMM | chr12:57010289 |
| Albumin | 1.236 | 2.028 | 1.094 | 1.017 | 0.229 | BOLT-LMM | chr4:73399955 |
| Calcium | 1.166 | 1.985 | 1.078 | 1.011 | 0.169 | BOLT-LMM | chr4:73399955 |
| Glycine | 0.976 | 1.457 | 1.016 | 0.983 | -0.053 | BOLT-LMM | chr16:81069345 |
| Height | 1.825 | 5.222 | 1.150 | 1.107 | 0.195 | BOLT-LMM | chr20:37261871 |
| Hemoglobin concentration, Asian ancestry | 1.008 | 1.015 | 1.001 | 0.998 | 0.574 | Linear regression | chr16:88716656 |
| IGF-1 serum levels | 1.320 | 2.995 | 1.079 | 1.053 | 0.160 | BOLT-LMM | chr20:37261871 |
| Mean corpuscular volume | 1.215 | 1.896 | 1.033 | 1.018 | 0.240 | BOLT-LMM | chr11:5225486 |
| Non-HDL cholesterol, European ancestry | 1.786 | 2.465 | 1.082 | 1.010 | 0.537 | BOLT-LMM | chr1:55029214 |
| Non-HDL cholesterol, African ancestry | 1.000 | 1.005 | 1.005 | 1.004 | 0.072 | Linear regression | chr1:55063542 |
| Total cholesterol | 1.739 | 2.568 | 1.082 | 1.009 | 0.471 | BOLT-LMM | chr4:73399955 |
| Uric acid | 0.803 | 4.198 | 1.059 | 1.036 | -0.062 | BOLT-LMM | chr1:125079549, chr1:121062032 |
| Gout | 1.008 | 1.336 | 0.847 | 0.838 | 0.024 | Logistic regression | chr1:125079549, chr1:121062032 |
| Hereditary ataxia | 1.019 | 1.017 | 0.262 | 0.154 | 1.142 | Logistic regression | chr19:13207859 |
| Myotonic dystrophy | 1.050 | 1.036 | 0.119 | 0.053 | 1.408 | Logistic regression | chr19:45770205 |

*Table S24 Correction factors and inflation metrics from phenotypes used in this study; LD score intercept, mean chi-squared unadjusted value, unadjusted lambda value, unadjusted lambda value for rare (< 1% MAF) markers and attenuation ratio. Marker represents the ID of the association reported.*

| Marker | $R^2$ imp vs raw | SaM vs. others | SaM vs. SaV | SaV vs. others | deCODE vs. Sa | deC vs. SanM | deC vs. SaV |
|---|---|---|---|---|---|---|---|
| chr1:55063542 | 0.997 | 0.098 | 0.294 | 0.746 | 0.211 | 0.1120 | 0.9887 |
| chr19:13207859 | 0.995 | 0.176 | 0.317 | 0.496 | 0.390 | 0.2105 | 0.6639 |
| chr19:45770205 | 0.879 | 0.292 | 0.583 | 0.731 | 0.394 | 0.3174 | 0.8304 |
| chr11:5225486 | 1.000 | 0.436 | 0.984 | 0.730 | 0.349 | 0.3726 | 0.6142 |
| chr12:57010289 | 1.000 | 0.429 | 0.006 | 0.006 | 0.634 | 0.7400 | 0.0090 |
| chr1:121062032 | 0.997 | 0.060 | 0.413 | 0.896 | 0.080 | 0.0563 | 0.8189 |
| chr1:125079549 | 0.998 | 0.103 | 0.317 | 0.620 | 0.186 | 0.1133 | 0.8276 |
| chr20:3726187 | 0.995 | 0.682 | 0.116 | 0.133 | 0.714 | 0.897 | 0.1720 |
| chr19:3939254 | 0.999 | 0.811 | 0.653 | 0.484 | 0.556 | 0.707 | 0.4582 |
| chr1:55029214 | 1.000 | 0.352 | 0.091 | 0.042 | 0.092 | 0.235 | 0.0318 |
| chr4:73399955 | 1.000 | 0.547 | 0.815 | 0.624 | 0.407 | 0.479 | 0.5579 |
| chr16:88716656 | 0.995 | 0.057 | 0.031 | 0.059 | 0.460 | 0.113 | 0.1034 |
| chr16:81069345 | 1.000 | 0.012 | 0.245 | 0.907 | 0.023 | 0.012 | 0.735 |

*Table S25 $R^2$ between raw genotypes and imputed markers in the XBI cohort. p-value for batch effect in the XBI cohort for markers presented in this study. deC = samples sequenced at deCODE genetics. SaV = samples sequenced using the Sanger Vanguard processing pipeline. SaM = samples sequenced using the Sanger main phase pipeline. Sa = samples sequenced at Sanger. Relationship between marker IDs and phenotypes can be seen in Table S24.*

| QC parameter | Sample level | Batch level |
|---|---|---|
| Sequencer type | Illumina NovaSeq6000 or better with standard 151 base, paired-end chemistry | |
| Sequencing library | PCR-free, uniquely dual-indexed in multiplexed pools | |
| Read-length | >100bp | |
| Proper-pairs | % of mapped read-pairs from the same DNA fragment with appropriate orientation and separation:<br>≥95% PASS<br><95% FAIL | |
| Coverage | % of autosome covered ≥15x:<br>≥95% PASS<br><95% FAIL | The mean sample genome coverage across the monthly sequencing batch is expected to be approximately 30X across the genome with a minimum coverage of 26X. |
| Contamination level 1<br>(Freemix) | Freemix sample contamination level as measured by VerifyBamID[80]:<br>≥5% FAIL<br>>1% and <5% further analyzed with Read_haps[81]<br><1% PASS | ≤4 samples per 96 sample sequencing plate<br>≤1% per monthly sequencing batch |
| Contamination level 2<br>(Read_haps) | For samples with Freemix values 1-5%, contamination is verified by Read_haps | |
| Sample Identity Concordance | Discordance at non-reference genotypes ≥2% FAIL<br><2% PASS | Sample identity concordance failures within each monthly sequencing batch must be <0.05% |
| Montly seq batch overall failure rate | | Repeat Sample requests are no more than 1% of the monthly sequencing batch |
